# Ultraliser: a framework for creating multiscale, high-fidelity and geometrically realistic 3D models for *in silico* neuroscience

**DOI:** 10.1101/2022.07.27.501675

**Authors:** Marwan Abdellah, Juan José García Cantero, Nadir Román Guerrero, Alessandro Foni, Jay S. Coggan, Corrado Calì, Marco Agus, Eleftherios Zisis, Daniel Keller, Markus Hadwiger, Pierre J. Magistretti, Henry Markram, Felix Schürmann

## Abstract

Ultraliser is a neuroscience-specific software framework capable of creating accurate and biologically realistic 3D models of complex neuroscientific structures at intracellular (e.g. mitochondria and endoplasmic reticula), cellular (e.g. neurons and glia) and even multicellular scales of resolution (e.g. cerebral vasculature and minicolumns). Resulting models are exported as triangulated surface meshes and annotated volumes for multiple applications in *in silico* neuroscience, allowing scalable supercomputer simulations that can unravel intricate cellular structure-function relationships. Ultraliser implements a high performance and unconditionally robust voxelization engine adapted to create optimized watertight surface meshes and annotated voxel grids from arbitrary non-watertight triangular soups, digitized morphological skeletons or binary volumetric masks. The framework represents a major leap forward in simulation-based neuroscience, making it possible to employ high-resolution 3D structural models for quantification of surface areas and volumes, which are of the utmost importance for cellular and system simulations. The power of Ultraliser is demonstrated with several use cases in which hundreds of models are created for potential application in diverse types of simulations. Ultraliser is publicly released under the GNU GPL3 license on GitHub (BlueBrain/Ultraliser).

**Significance:** There is crystal clear evidence on the impact of cell shape on its signaling mechanisms. Structural models can therefore be insightful to realize the function; the more realistic the structure can be, the further we get insights into the function. Creating realistic structural models from existing ones is challenging, particularly when needed for detailed subcellular simulations. We present Ultraliser, a neuroscience-dedicated framework capable of building these structural models with realistic and detailed cellular geometries that can be used for simulations.

**Key points:**

- Ultraliser creates spatial models of neuro-glia-vascular (NGV) structures with realistic geometries.
- Ultraliser creates high fidelity watertight manifolds and large scale volumes from centerline descriptions, non-watertight surfaces, and binary masks.
- Resulting models enable scalable *in silico* experiments that can probe intricate structure-function relationships.
- The framework is unrivalled both in ease-of-use and in the accuracy of resulting geometry representing a major leap forward in simulation-based neuroscience.

## 1 Introduction

It has been more than a 100 years since Santiago Ramón y Cajal (1854–1934) commenced his pioneering quest to study the brain by elucidating its anatomical structures and establishing the neuron doctrine^1^. Nevertheless, and so far, our knowledge is still incomplete, particularly at cellular and synaptic levels-of-detail^2^. Even with the broad spectrum of research that followed Cajal’s leading efforts, taking into account the vast amount of resulting data, it has been proven that conventional wet lab experiments alone are insufficient to unravel the underlying function of the brain^3^. The generation of massive amounts of experimental data in addition to the recent quantum leap in computing technologies have led to the renaissance of a complementary approach: simulation neuroscience^4^. Mathematical modeling, computer simulations and terabytes of structural data resulting from a myriad diversity of experiments are being successfully consolidated — with this approach we can test our hypotheses and predict the quantitative behavior of complex biological processes^5^.

Data-driven models integrated with systematic computational methods can dramatically increase efficiency, efficacy and reliability of simulations^6^, particularly when our experimental knowledge is fragmented^5^. An important question is whether we can reuse existing structural data to synthesize detailed, multiscale and biologically plausible three-dimensional (3D) models that can be integrated into simulation contexts to gain insights into the cellular function. The interdependency of structure and function produces a kind of metaplasticity^7^ that requires performing simulations within geometrically realistic subdomains at ultrastructural resolutions in which molecular reactions can be contained^8^.

Structural neuroscientific datasets are either acquired from a wide spectrum of imaging modalities, such as imaging scanners and microscopes^9–11^ or digitally synthesized in supercomputer simulations to yield similar structural characteristics of biological counterparts^12–15^. 3D models of structural data (on the scale of 100 nm - 1 mm) can be classified according to their digital representations into four principal formats: morphology skeletons, surface meshes, volumetric meshes and volumetric grids. Each representation (**Supplementary Fig. S1**) is convenient for a specific category of simulation.

Morphology skeletons are point-and-diameter descriptions of either acyclic or cyclic graphs that can model connectivity of neuronal arborizations^16^, astrocytic processes^15^ and dense vascular networks^17,18^. Aside from their usage for topological and visual analysis^19^, these morphologies are used to conduct one-dimensional (1D) compartmental simulations. Neuronal and glial morphologies are used in [NEURON](https://neuron.yale.edu/)^20^ (neuron.yale.edu) to simulate electrophysiology based on Hodgkin-Huxely ion channel formalism^21^, and vascular morphologies are used to simulate blood flow in cerebral vasculature^22,23^.

Surface meshes are sets of vertices, edges and facets (ideally triangles) that can define the boundaries of 3D structures, such as cellular membranes of neurons^24^ and astrocytes^25^ and tubular membranes of blood vessels^26^. These meshes, if watertight, are extensively used in particle-based stochastic molecular simulations, for example with the [MCell](https://mcell.org/) simulator^27,28^(mcell.org). Volumetric meshes are derived from their surface counterparts to model their interior volume with convenient discretization, for example, with tetrahedral^29^ or hexahedral^30^ subdomains. Such meshes are primarily used in reaction-diffusion simulations with [STEPS](https://github.com/CNS-OIST/STEPS)^31^ (steps.sourceforge.net) or [Smoldyn](https://www.smoldyn.org/)^32–34^ (smoldyn.org).

Cartesian volumetric grids use, alternatively, cubic discretization to model the interior volume, in which we can account for the variations in optical properties of different regions of the tissue to simulate its interaction with light^35,36^.

Generally, the same 3D model can be converted from one format to another to be used in hybrid or multimodal simulations; however, the principal format with which a given 3D model can be restructured into any other format is a surface mesh that must be watertight^37^ (**Supplementary Figs. S1b, S2**). For instance, the generation of a volumetric mesh from a surface input —using [TetGen](https://wias-berlin.de/software/index.jsp?id=TetGen&lang=1)^38^, [QuarTet](https://github.com/crawforddoran/quartet)^39^, [GMsh](https://gmsh.info/)^40^ or [CGAL](https://www.cgal.org/)^41^— requires the surface mesh to be watertight. [TetWild](https://wildmeshing.github.io/tetwild/) can tetrahedralize non-watertight meshes, but it has limited performance^29^ and fails to handle complex biological models with realistic geometries^42^. Moreover, accurate skeletonization of cerebral vasculature from microscopy stacks requires multimodal algorithms that combine an input volume with its corresponding watertight mesh to create a morphological representation of the network^43–45^. Therefore, to automate and systematize simulation workflows, watertightness is essential, not only for the simulation per se, but also for data conversion from one representation to another.

On one level, and as a consequence of the segmentation challenges of electron microscopy (EM) volumes, existing 3D mesh models of cellular and subcellular brain structures are expected to be fragmented and non-watertight. This applies to manually segmented neuropil structures^11^ or even those segmented with state-of-the-art deep neural networks^46^. Machine learning (ML) has helped automating the process^47^ allowing to generate a massive amount of neuro-glia-vasculature (NGV) reconstructions that are disseminated online as demonstrated by several research programs such as [MICrONS](https://www.microns-explorer.org/)^48^, FAFB-FFN1^49^, and [FlyEM Hemibrain](https://www.janelia.org/project-team/flyem/hemibrain)^50^. Nevertheless, the majority of the resulting reconstructions requires many person-years of effort for proofreading. Online collaborative efforts are introduced to accelerate the process^51^, which could be effective in resolving fragmentation artifacts, but watertightness remains lacking.

On another level, there is a huge diversity of NGV morphological models that have been released publicly to central databases, for instance, NeuroMorpho.Org^16^ and [Brain Vasculature (BraVa)](http://cng.gmu.edu/brava/home.php?s=1&name_browser=false)^18^. These models are only appropriate for conducting 1D compartmental simulations; there are no existing frameworks that can convert them, taking into consideration their structural artifacts, into watertight mesh models for applications in other types of simulations. The key question therefore is, given an input 3D model in any of aforementioned formats, can we reconstruct it in another format that can be systematically plugged into simulation environments for *in silico* experimentation?

A few relevant *re-meshing* frameworks are capable of handling geometric topology and optimization issues for relatively small scale structures (minuscule segments of spiny dendrites), such as [GAMer](https://github.com/ctlee/gamer)^52^ and [VolRoverN](https://cvcweb.oden.utexas.edu/cvcwp/software/volumerover-neuron/)^53^, but they are incapable of accomplishing watertightness and they cannot process any kind of morphology skeletons. Other applications presented applicable solutions to construct polygonal meshes from morphology skeletons, for (i) neurons such as [NeuroTessMesh](https://github.com/vg-lab/neurotessmesh)^54^, NeuroMorphoVis^19^, [Neuronize](https://github.com/vg-lab/Neuronize)^55^, [AnaMorph](https://github.com/NeuroBox3D/AnaMorph)^56^, (ii) vasculature, such as [VessMorphoVis](https://github.com/BlueBrain/VessMorphoVis)^26^ and (iii) astrocytes^25^ (summarized in **Supplementary Tables S1** and **S2**). Nonetheless, the resulting meshes are neither optimized nor watertight (**Supplementary Section 3**), furthermore they might have unrealistic geometries and structural deficits. We present Ultraliser to eliminate the gap and address those challenges, all within a unique and efficient framework.

## 2 Results

### 2.1 Ultraliser

Ultraliser is a neuroscience-specific framework capable of creating multiscale (from the subcellular scale up to mesoscale circuits) and high fidelity 3D models of neuroscientific datasets that can be integrated in the context of simulation-based experiments, aiming to understand the function. Ultraliser is consistent and unconditionally robust, as it can systematically build adaptively optimized watertight triangular meshes and large-scale annotated volumes from input data models with multiple formats including: (i) ill-conditioned, fragmented and self-intersecting polygonal meshes with irregular topologies, (ii) cyclic and acyclic morphological graphs, (iii) single-channel volumetric stacks given with user-specified isovalues, (iv) binary volume masks segmented from microscopy stacks, and (v) tetrahedral volumetric meshes.

Contrary to traditional remeshing applications that use geometry-based methods^57^ to repair the geometric topology of non-watertight meshes, the core engine of Ultraliser is designed based on efficient voxelization kernels that create intermediate high resolution proxy volumes, with which we can extract continuous surface meshes that are adaptively optimized and watertight, refer to **Figure 1** and **Supplementary Figure S2**. The core library of Ultraliser is exploited to develop several applications that can be part of a large-scale software ecosystem for establishing fully automated neuroscientific pipelines involving multimodal simulation systems. The current version of Ultraliser integrates the following applications:

**Figure 1.**
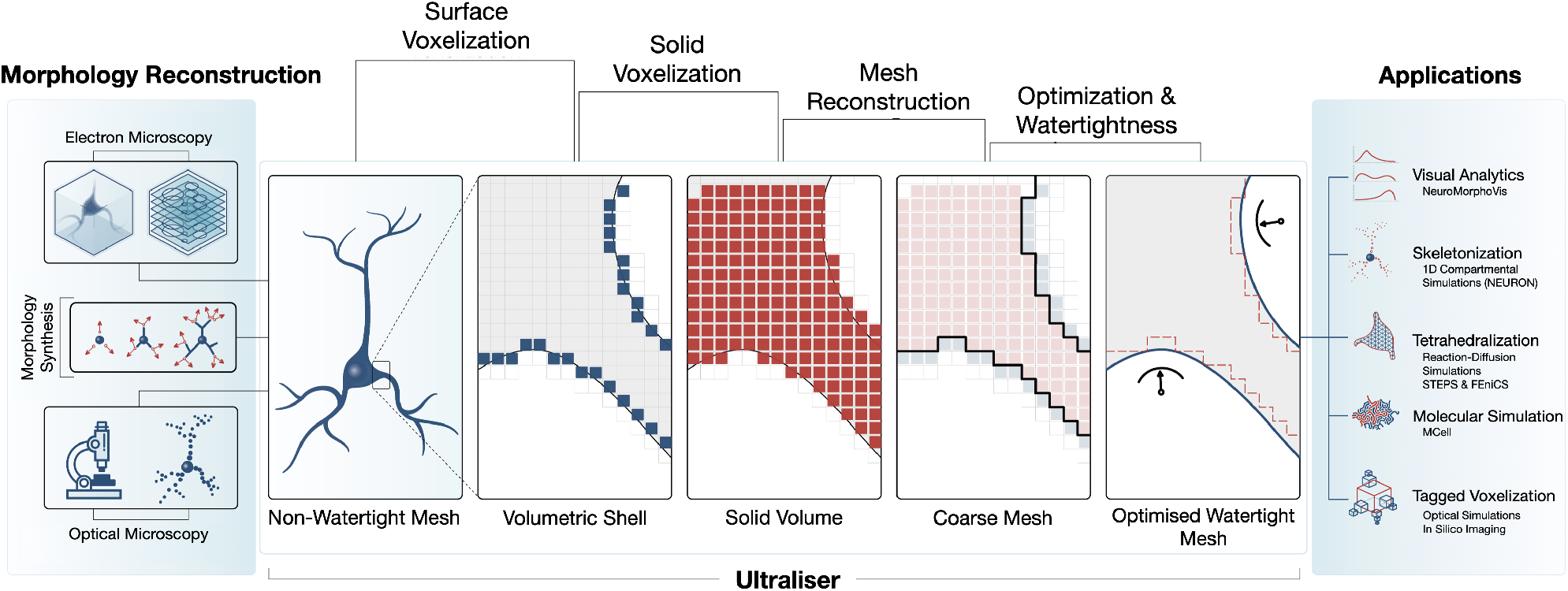
Ultraliser workflow. Ultraliser implements a voxelization-based remeshing engine to create annotated volume models and watertight surface manifolds from input morphology skeletons, non-watertight triangular soups, gray scale volumes and segmented binary masks. The workflow has five essential stages: surface and solid voxelization, triangular mesh reconstruction from uniformly sampled volume grids, surface optimization, and watertight verification. Detailed workflow is illustrated in **Supplementary Figure S2**.

1. ultraMesh2Mesh, restructures an input polygonal surface mesh composed of a set of unorganized triangles with no defined connectivity—i.e. a triangle soup—into a smooth, optimized, two-manifold and watertight triangular surface mesh.
2. ultraMeshes2Mesh, a similar application to ultraMesh2Mesh, but it produces a single output watertight mesh from a list of non-watertight input meshes that have existing spatial relationship.
3. ultraMesh2Volume, reconstructs 1-bit (one bit per voxel) and 8-bit (one byte per voxel) volumes from an input mesh that is not necessarily watertight and might have severe geometric deficits including self-intersecting facets, fragmented partitions and even floating vertices.
4. ultraVolume2Mesh, generates a watertight surface mesh from a single-channel volume stack. The surface of the resulting mesh is established based on a user-specified isovalue or range of isovalues.
5. ultraMask2Mesh, a similar application to ultraVolume2Mesh, but it takes an input binary mask that is typically segmented from microscopy stacks, where the isosurface is already reconstructed as a set of voxels.
6. ultraNeuroMorpho2Mesh, converts an input acyclic graph representing a neuronal morphology skeleton into an optimized and continuous membrane with a biologically realistic 3D somatic profile reconstructed on a physically plausible basis. Spines can be also integrated along the dendritic surface if the morphology is reconstructed in a digital microcircuit^2^, where synaptic locations are determined.
7. ultraAstroMorpho2Mesh, converts an input astrocytic morphology containing branching processes and endfeet surfaces into an optimized and continuous membrane surface.
8. ultraVessMorpho2Mesh, converts an input cyclic graph representing a large-scale vascular network into an optimized, watertight and multi-partitioned mesh model with accurate branching geometries.
9. ultraCircuit2Volume, takes an input microcircuit and a configuration file, and produces a volumetric tissue model that is tagged with multiple optical properties. The configuration file describes the annotation details of the circuit^10^.

In the following sections, we demonstrate the significance of Ultraliser for *in silico* neuroscience taking into account several use cases that involve multiple types of input structural data including: non-watertight surface meshes of cellular and subcellular structures segmented from EM volumes and NGV (neuronal, astrocytic and vascular) morphologies that are either segmented from optical microscopy stacks or generated synthetically.

### 2.2 Remeshing cellular and subcellular structures of the NGV ensemble^58^

In the context of a recent collaboration between EPFL and KAUST, we developed a multi-stage framework (imaging, segmentation, modeling, simulation and visualization) to re-create the NGV ensemble *in silico*^58^. This framework aims to advance our understanding of the substantial role of cytoscale structures and functions in controlling brain energy metabolism. Our fundamental objective is to digitally reconstruct 3D structural models of NGV structures with realistic geometries to fuel detailed subcellular simulations^59,60^, allowing us to investigate the biochemical and biophysical properties of oligocellular networks^61^. Realizing the objectives of this framework, however, has been impeded by the lack of availability of topologically accurate watertight meshes with which we can conduct the simulations or even skeletonize the meshes extracted in the segmentation stage (**Fig. 2a**).

**Figure 2.**
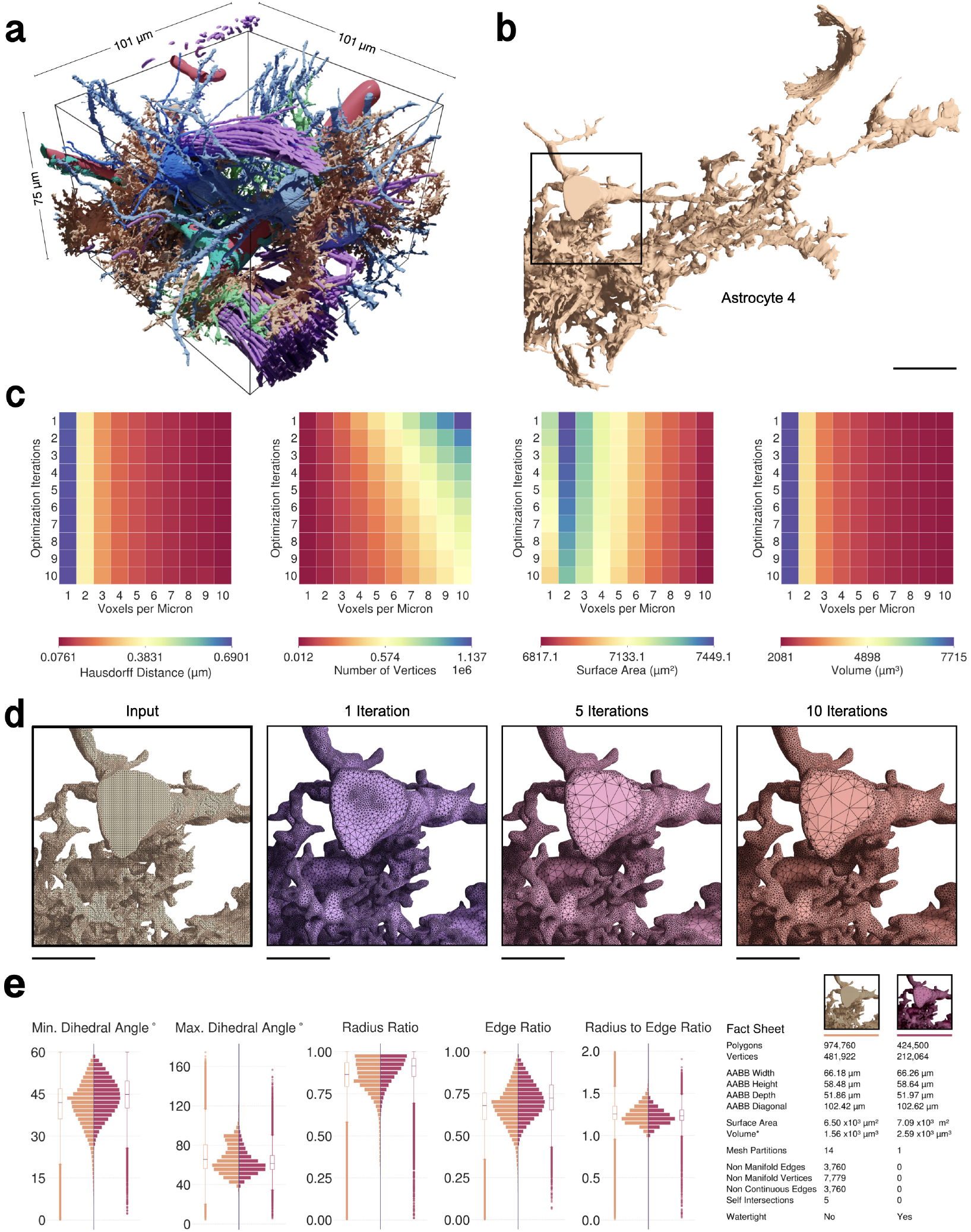
Ultraliser creates high-fidelity watertight surface manifolds of neuropil structures from non-watertight inputs. (**a**) The EM volume block is segmented semi-automatically to extract 3D mesh models of individual cell morphologies and other structures (**Supplementary Fig. S3**). (**b**) An exemplar mesh (Astrocyte 4) is selected for evaluating the optimum values of the remeshing parameters. (**c**) The effect of varying voxelization resolution (in voxels per micron) and the number of optimization iterations on— from left to right—the Hausdorff distance (in μm), number of vertices, total surface area (in μm^2^) and volume (μm^3^) of the mesh. Astrocyte 4 has been re-meshed at multiple voxelization resolutions (0.1 - 1.0 μm) and optimized with different optimization iterations (1 - 10) to determine the most optimum values for these parameters as a reference to be used to remesh all the other segmented structures from the neuropil volume. (**d**) The input mesh of Astrocyte 4 is not watertight and is also over tessellated with ∼ 1.7 million triangles. This mesh has been re-meshed with a voxelization resolution of 5 voxels per micron (200 nm), and optimized with one, five and ten optimization iterations. (**e**) ultraMesh2Mesh creates an adaptively optimized watertight mesh with only ∼ 425 thousand triangles. The distributions show a comparison between the qualitative analysis metrics of the input mesh and resulting one. Scale bars, 5*μ*m (**b**), 2.5*μ*m (**d**).

We acquired a 750,000 cubic micron volume from layer IV of the somatosensory cortex of a two-week-old rat. Within this volume, a total of 186 structures were labeled, segmented, and classified into (i) cellular structures including: astrocytes, neurons, microglia, pericytes and oligodendrocytes, (ii) subcellular structures including: nuclei, mitochondira and endoplasmic reticula (ER), (iii) other fragmented structures including a few blood vessel segments and a group of myelinated axons and (iv) other non-identifiable structures. From this collection, the following *complete* structures — that are of central significance to our modeling objectives — were segmented: four astrocytes, four neurons, four microglia, four pericytes and a single oligodendrocyte in addition to the mitochondria of all the cells and the ER of the astrocytes (**Supplementary Section 5**).

A surface mesh model corresponding to each structure is exported into a Wavefront OBJ file for qualitative and quantitative analysis (**Supplementary Tables S3** and **S4**). From those 17 cells and their subcellular structures, only the meshes of three pericytes and astrocytic ER were verified to be watertight, but they had poor geometric topologies. Meshes of the remaining cells were confirmed to be non-watertight; containing thousands of non-manifold edges and vertices in addition to tens of self-intersecting polygons.

We then re-meshed all the cellular and subcellular meshes using ultraMesh2Mesh to reconstruct corresponding watertight manifolds that are accurate and geometrically optimized (Methods). Geometric accuracy, tessellation and topology of the resulting meshes are subject to two principal parameters: voxelization resolution and number of optimization iterations. Resolution is defined by the number of voxels per micron used to rasterize the input mesh. Geometric accuracy is primarily measured by the Hausdorff distance and difference in volume between the input and output meshes.

To estimate the optimum values of these two parameters, we constructed five analysis matrices for an exemplar mesh (Astrocyte 4, shown in **Fig. 2b**) in which we can apply those optimum values to other cellular meshes in the block. **Figure 2c** illustrates four quantitative matrices showing the effect of varying the two principal parameters on the Hausdorff distance, number of vertices, surface area and volume of the output mesh. The number of vertices determines the footprint of the resulting mesh. Essentially, the lower this footprint is, the better for processing. However, the lower it becomes, the values of the difference in volume and Hausdorff distance between resulting and original meshes are increasing, which implies altering shape or losing detail — for example, of a spine located along a dendritic branch. Therefore, we must combine the volume analysis and Hausdorff distance matrices to determine convenient values of voxelization resolution and optimization iterations that could preserve the geometry and volume of the original mesh **(Supplementary Section 4). Figure 2d** shows the effect of using one, five and ten optimization iterations on the tessellation and topology of the resulting mesh. The full analysis matrix is shown in **Supplementary Figure S4**. From this analysis, the optimum values were estimated to be five voxels per micron and five optimization iterations. Those values are then used to re-mesh all the cellular meshes in the block. **Supplementary Figures S5 - S26** (summary in **Table S6**) reveal detailed quantitative, qualitative and visual comparisons (the results for Astrocyte 2 are shown in **Figure 2e**) between the input and output meshes of the complete cellular structures shown in **Figure 2a**.

Subcellular meshes were re-meshed with higher voxelization resolution (10 voxels per micron) to ensure resolving their fragmented and minuscule segments. Complete comparative analysis of the subcellular meshes is shown in **Supplementary Figures S29 - S53** (**Table S7**).

Resulting astrocytic meshes were therefore applicable for skeletonization, with which we have successfully developed a novel pipeline to synthesize a digital reconstruction of the NGV ensemble at micrometer anatomical resolution^15^. Moreover, all the resulting meshes were tetrahedralized using [TetGen](https://wias-berlin.de/software/tetgen/features.html)^38^ and [Gmsh](https://gmsh.info/)^40^ to create corresponding volume, or simulation-ready, meshes for [STEPS](https://github.com/CNS-OIST/STEPS)^31^ simulations with which we have successfully completed the objectives of the collaboration. Simulation results are beyond the scope of this sequel.

### 2.3 Remeshing poorly segmented meshes with fragmented partitions and slicing artifacts

Dense reconstructions of brain circuits are made available with volume EM and advanced ML-based segmentation solutions, allowing us to render hundreds of thousands of cortical structures — including complete cells, cell parts, cytoplasmic organelles and blood vessels — that are shared^62^ and made freely available online^49,50,63,64^. A decent amount of the segmented structures are proofread to resolve false-split (fragmentation) and false-merge (connectivity) errors. Nevertheless, pipelines involved in the segmentation process yield triangular mesh models characterized by sharp features, rough surfaces and high tessellation rates, leading to triangle soups with giant numbers of geometric deficits. The meshes might have large discontinued partitions with overlapping geometries (**Figs. 3a, b, f**), gaps (**Fig. 3j**) and tiny disconnected fragments (**Supplementary Fig. S75**) due to common slicing and misalignment artifacts^65^. Those poorly reconstructed and fragmented meshes cannot be effectively repaired — or remeshed to produce watertight counterparts — relying on geometric-based solutions^57^. Thanks to its voxelization engine, Ultraliser can transparently handle these deficits and build a continuous, adaptively tessellated and high resolution manifold of the entire structure with superior topology.

**Figure 3.**
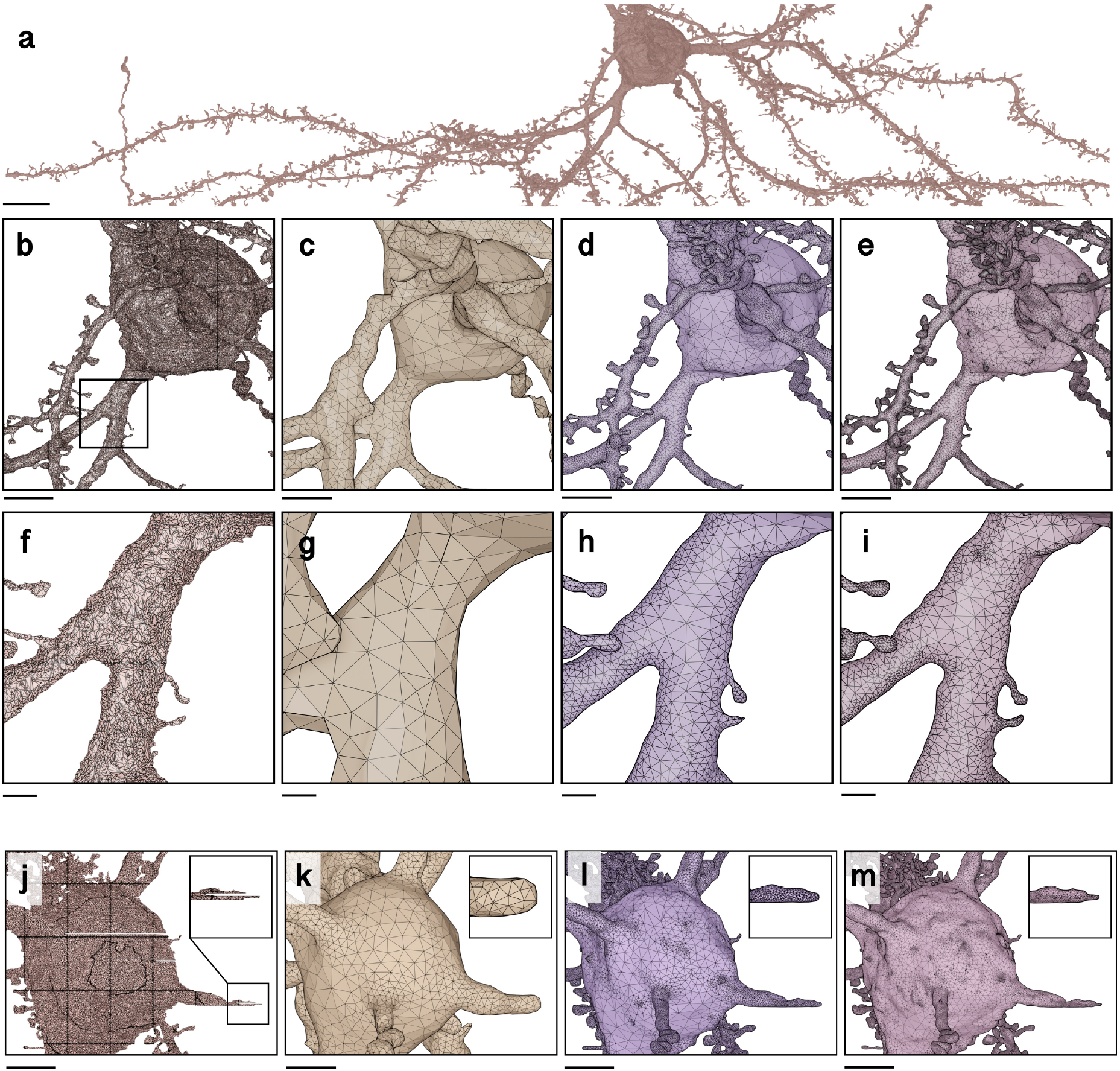
Remeshing fragmented mesh model with multiple mesh partitions, self intersecting faces and slicing artifacts. (**a**) Segmented mesh model of a neuron extracted from layer II/III of the visual cortex of the rat. The closeup in (b) shows the messy fragmented structure of the mesh. This triangle soup has been processed with Ultraliser to generate an optimized, connected and two-manifold mesh model with high quality topology at three different resolutions: (**c**) one, (**d**) five and (**e**) 10 voxels per micron. The closeups in (**f**) - (**i**) focus on a small spiny branch to highlight the effect of varying the reconstruction resolution on the ultrastructure of the mesh. Supplementary Figure S55 shows comparative analysis between the input mesh and the ultralised one. (**j**) The mesh contains a few gaps due to the misalignment and slicing artifacts. The 3-way solid voxelization is used to reconstruct a continuous manifold with optimized topology at different resolutions (**k, l, m**). Scale bars, 10 *μ*m (**a**), 5 *μ*m (**b, c, d, e**), 1 *μ*m (**f, g, h, i**).

To accomplish this objective, we used ultraMesh2Mesh, with which the polygons of an input surface mesh are independently rasterized into a volume grid, where self-intersecting and duplicate facets are implicitly eliminated; they are rasterized to the same voxel or a neighboring one. This grid is then processed by isosurface extraction kernels to produce an intermediate highly tessellated surface, which is smoothed, adaptively optimized and finally verified to be watertight (Methods). **Figure 3** demonstrates how effective Ultraliser can be in processing an exemplar mesh —segmented from layer II/III of the visual cortex of a young rat— with thousands of mesh partitions, self-intersections and other geometric artifacts. **Supplementary Figures S55 - S74** show comparative remeshing results for a small subset of meshes of pyramidal neurons (the full block is shown in **Supplementary Fig. S54**) that were publicly provided by [MICrONS program](https://www.microns-explorer.org/)^48,66^. All the resulting meshes are watertight and have a single mesh partition with a continuous manifold.

Segmented meshes suffering from slicing or alignment artifacts are, in certain cases, characterized by thin gaps across their surfaces. Such gaps lead to surface discontinuity (**Fig. 3j**) or disconnected fragments of the mesh (shown in red in **Supplementary Fig. S75**). Those gaps cannot be easily detected and might require advanced and computationally intensive ML-based algorithms to identify and repair them. In our implementation, we extended the voxelization stage and integrated a 3-way solid voxelization algorithm, that is seamlessly taking into account repairing those gaps, in which the interiors bounded by cell membranes are voxelized along each axis independently and then merged into a single volume (Methods). **Figures 3k, l, m** illustrate the results of remeshing a pyramidal neuron mesh with three obvious slicing artifacts (two of them exist on the soma and one is located along a dendritic branch) into a continuous surface mesh at three different voxelization resolutions. **Supplementary Figure S75** illustrates a side-by-side comparison between an input mesh with several artifacts and the resulting one from ultrasMesh2Mesh, in which all the gaps are filled to connect the disconnected fragments to the surface of the mesh to produce a continuous surface.

### 2.4 Generating biologically realistic neuronal meshes from digitized morphologies

There is a huge diversity of neuronal morphologies (mainly in SWC format) that is routinely used for simulating electrical activity in [NEURON](https://neuron.yale.edu/)^20^ and its modern extensions^67,68^. Such diversity can be a significant resource for performing meso-scale hybrid simulations combining electrophysiology with intracellular calcium dynamics simultaneously^69^. However, this objective entails the development of a robust technique that can construct biologically detailed and watertight neuronal mesh models consistent with their morphological counterparts. Several applications were developed to create neuronal surface meshes from their corresponding morphologies (**Supplementary Table S1**). The majority was focused on building low-tessellated, non-watertight and visually appealing mesh models that can be rapidly generated and efficiently used for visualization or in visual analytics^19,24,54,55,70^. Only a few ones addressed the watertightness challenge, however, using simplified geometries and without support to integrate spine models^56,71^. We therefore implemented an application (ultraNeuroMorpho2Mesh) capable of consolidating neuronal meshes combining both features, biological realism and watertightness, irrespective to the conditions or topology of the input morphology.

To create an integrated mesh model of a spiny neuronal morphology, our implementation addressed four principal challenges: (i) creating a plausible 3D somatic surface that can simulate the biological growth of the soma relying merely on the initial segments of the neurites, (ii) creating neuritic arborization membranes with non-overlapping geometries around their bifurcation points, irrespective to their branching angles, (iii) integrating realistic spine models along the membranes of their dendritic branches, and finally (iv) optimization and watertightness verification (Methods).

Digitized neuronal morphologies are composed of three distinct structural components: somata, neurities and dendritic spines (**Supplementary Fig. S76**). Somata are typically approximated with geometric primitives, mainly a sphere^71^, whose radius is computed based on the relative locations of the initial segments of each neurite, or in some cases, cylinders^72^. Advanced traces digitize the soma into a two-dimensional (2D) contour representing its projection along the optical axis. To reconstruct a plausible 3D somatic profile, we apply the finite element method (FEM) approach^73^ to deform a volumetric model of an elastic sphere by pulling towards each neurite in the morphology. This approach preserves the initial volume of the sphere, resulting into more realistic somatic surface (**Fig. 4a**). Neurites are represented by directed acyclic graphs (DAGs) as a set of interconnected nodes, each of which defines a 3D Cartesian position and radius (**Fig. 4b**), with which we can reproduce cross-sectional variations and orientation of each segment in the morphology. A depth-first scheme is used to construct a set of connected paths from the root node (or the soma) to the terminal ones. For each path, a cross-sectional geometry is created as an independent proxy mesh (**Fig. 4c**) (Methods). The integration of the spines along dendritic membranes is optional; spines are not comprised by default within the morphological descriptions of individual neurons loaded from SWC files. Spiny neurons are modeled after circuit building^2^, where we can localize synapses and characterize their spine attributes. We designed a set of realistic spine geometries (**Supplementary Fig. S79**) based on a few reconstructions of interneurons segmented from the somatosensory cortex^11^. All the proxy meshes — of the soma, neurites and spines — are agglomerated and rasterized in parallel to create a corresponding solid volume, with which the final mesh is generated (Methods). **Figure 4** illustrates the steps of building a mesh model of a spiny neuron from its morphology skeleton. The resulting mesh is shown in detail in **Supplementary Figure S80**. Our implementation has been tested with a group of 25 neurons with various morphological types^16^. Morphology files and resulting meshes are available in the Supplementary Data.

**Figure 4.**
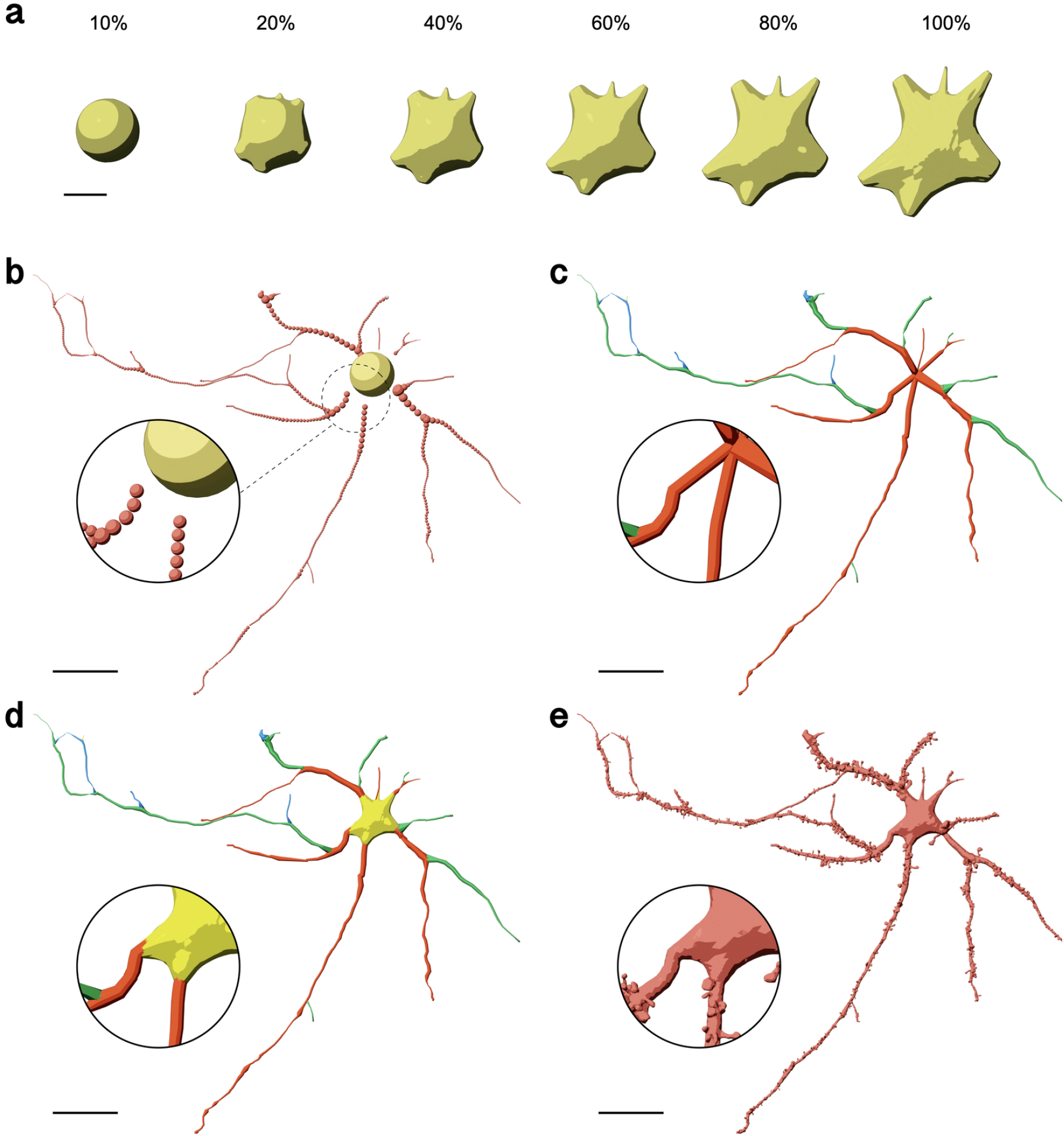
Creating biologically realistic spiny neuronal surface mesh from its morphology. (**a**) Progressive reconstruction of the soma from an initial icosphere into a 3D plausible profile based on the FEM approach^73^. (**b**) The morphology is rendered as a set of samples. (**c**) For each neurite, we reconstruct a list of proxy-geometries linking a set of principal sections from the root node and until the leaf: level 1 is in red, level 2 is in green, and level 3 is in blue. Note that the proxy geometries start from the origin of the soma to avoid any gaps when the soma mesh is added afterwards (**d**). (**e**) Spine proxy geometries are added along the surface of the proxy-mesh. All the section geometries, spines and somatic mesh are rasterized to create a continuous and watertight manifold. Renderings of multiple closeups of this mesh are shown in **Supplementary Figure S80**. Scale bars, 5 *μ*m (**a**) and 20 *μ*m (**b, c, d, e**).

### 2.5 Generating astroglial meshes from complete synthetic morphologies

Complete astrocytic morphologies with endfeet reconstructions are sparse. NeuroMorpho.Org has ∼5,500 astrocytes, but none of them contains any endfeet descriptions; only the arborizations of perisynaptic and perivascular processes. Based on a few reconstructions of complete astroglial morphologies^11^, we presented in a recent study^15^ an effective method to synthesize biologically inspired, and complete, astrocytic morphologies including endfeet patches (the skeleton of an astroglial morphology is shown **Supplementary Fig. S77**). The objective of the study is to allow the creation of a huge diversity of astrocytic morphologies that can be used to understand their structure-function relationship using molecular simulations.

We therefore complemented this effort and integrated a specific application (ultraAstroMorpho2Mesh) to create simulation-ready astrocytic meshes from their morphological counterparts. The mesh generation algorithm is similar to that used to create neuronal meshes, but in addition, endfeet proxy geometries are created using implicit surface modeling (Methods). Figure 5 illustrates the results for reconstructing a watertight astrocytic mesh that is consistent with its morphological description. We also tested the implementation with a group of 25 astrocytes sampled from different cortical regions and created their corresponding watertight manifolds (Supplementary Data).

**Figure 5.**
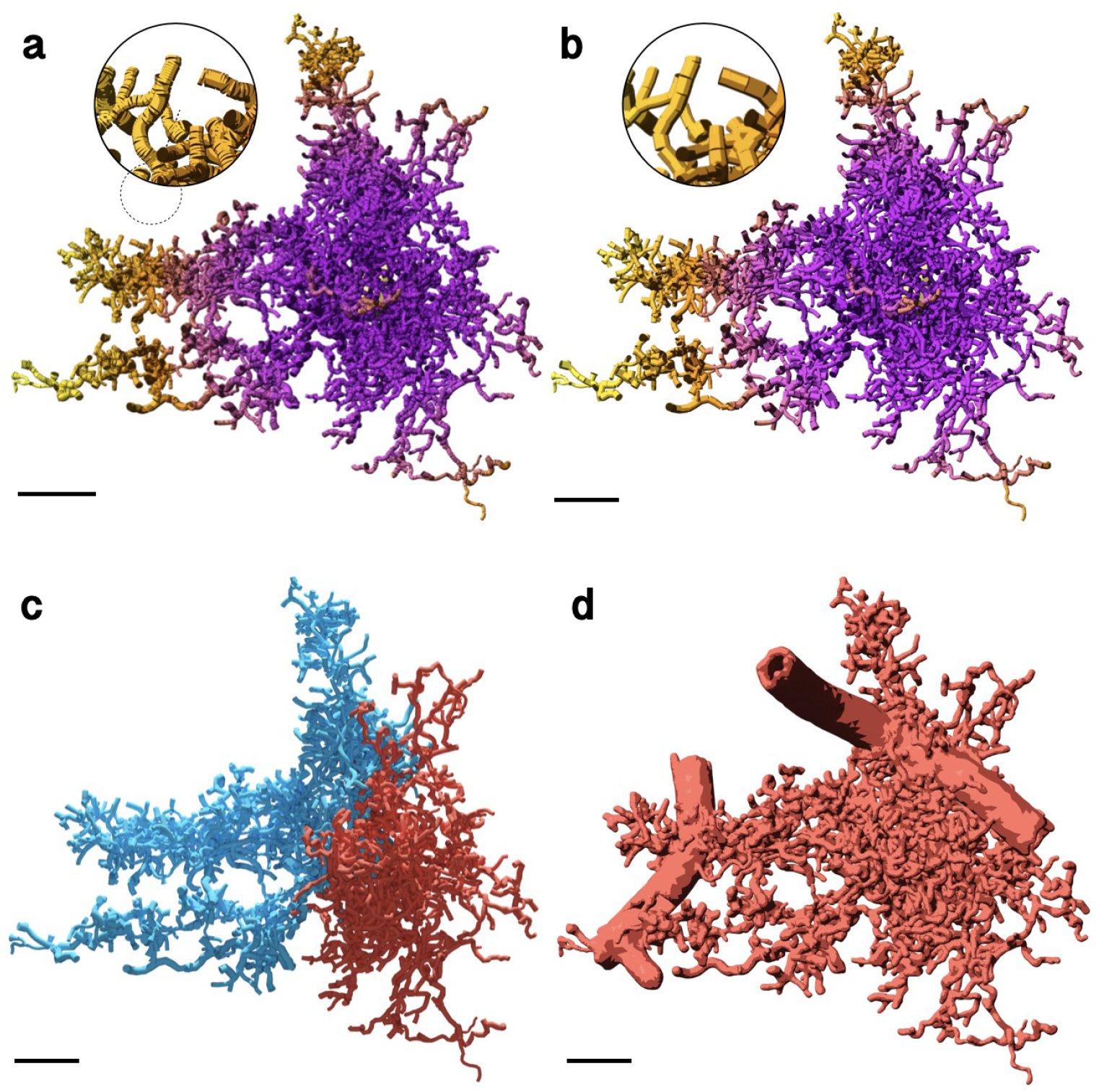
Creating a realistic and watertight astroglial mesh from synthetic astrocytic morphology with two endfeet. (**a**) Synthetic astrocytes are excessively oversampled. (**b**) The processes are adaptively resampled to avoid any reconstruction artifacts during the mesh generation process. (**b**) The processes are adaptively resampled for convenience. (**d**) Reconstruction of the astrocytic surface mesh with endfeet included. A high resolution reconstruction of this mesh is illustrated in **Supplementary Figure S82**. Scale bars, 10 *μ*m (**a, b, c, d**).

### 2.6 Generating continuous cellular meshes from fragmented components

We extended the re-meshing pipeline and integrated another application (ultraMeshes2Mesh) that agglomerates a list of fragmented meshes —that are spatially overlapping— into a single and continuous watertight mesh. This extension, however seamless, enables the reconstruction of ultrarealistic cellular models based on existing meshing implementations (**Supplementary Table S1**), in which we can assemble 3D mesh models of different cellular components generated independently by multiple third-party applications into a single watertight mesh with continuous surface (Methods). For instance, ultraNeuroMorpho2Mesh uses the FEM to reconstruct a plausible 3D surface of the soma, but meanwhile, there are other advanced implementations that use Hooke’s law and mass-spring models to reconstruct 3D somatic profiles with more realistic shapes, which could ultimately improve the realism of resulting neuronal meshes. This can be demonstrated with the [Soma Reconstruction Toolbox](https://github.com/BlueBrain/NeuroMorphoVis/wiki/Soma-Reconstruction) in NeuroMorphoVis^19^, where users can fine tune the parameters of the soma reconstruction algorithm and validate the resulting profile with respect to a segmented ground-truth mesh. **Supplementary Figure S81** provides an example of creating a watertight mesh of a spiny neuron from a set of input meshes, each represents a single component of the neuron (soma, arbor, or even a spine). ultraMeshes2Mesh can therefore be seen as a complementary or post-processing application that can ensure the watertightness of resulting mesh models of NGV cellular structures created by other software applications.

### 2.7 Generating vasculature meshes from corresponding graph networks

Another application is developed to create ‘multi-partitioned’ watertight mesh models of large-scale cerebral vascular networks. This application is intended to confront the rising trend to automate the reconstruction of accurate 3D models of brain vasculature, with which we can analyze their structural angioarchitecture and characterize their dynamic behavior^17,75^. The networks are typically segmented into vectorized graphs, i.e. centerlines with point-and-diameter representations (the structure of a vascular morphology graph is shown in **Supplementary Fig. S78**). Those graphs are becoming imperative for performing vascular simulations, whether used in vectorized format in 1D compartmental simulations or being converted into alternative formats, for example, into tetrahedral meshes that can be applied in reaction-diffusion simulations. Such simulations pave the way to understand how our brains meet the energy demands of neuronal computations. Nevertheless, creating such ‘simulation-ready’ vascular models from segmented data is challenging.

The first challenge is the fragmentation of the network. Even with current state-of-the-art imaging and segmentation protocols, it is near impossible to reconstruct a full – and accurate – high resolution cerebral vascular network segmented into a single and connected graph^76^. Resulting graphs are typically composed of multiple disconnected partitions, which complicates the creation of watertight meshes if the partitions are self-intersecting. The second challenge is the scale of segmented networks, which has been exponentially growing due to the recent advances in lightsheet imaging and Clarity^77,78^, allowing to create whole-brain vascular maps down to capillary level^79^. The third challenge is the segmentation quality of the vessel network; in particular for small vessels, the quality is not optimum, and the resulting skeletons might have severe topological artifacts around branching terminals. Therefore, typical meshing algorithms of branching structures would fail to build watertight meshes of such complex geometries.

There are plenty of tools that can be used to visualize the graphs, but only a few are capable of visualizing large-scale graphs^26,80^. Some tools^26,54^ can also create surface meshes from their corresponding morphologies, but usually the results are not watertight, particularly for dense graphs. To fill this gap, we designed ultraVessMorpho2Mesh, a vasculature-specific application that can efficiently convert large-scale networks of vascular morphologies into multi-partitioned and adaptively optimized watertight meshes with smooth branching geometries. Our algorithm handles an input graph as a linear list of sections without the necessity to have predefined connectivity information. Initially, the morphology skeleton is analyzed, where high frequency variations in cross-sectional radii are filtered and short sections are eliminated (**Fig. 6b**). Each section in the graph is independently converted into a proxy mesh. However, and to guarantee the connectivity of branches along the surface of the final mesh, we add multiple sphere meshes (icospheres) at both terminals of the section. All the proxy meshes are then rasterized and converted into a solid volume, with which the final mesh is generated (Methods).

**Figure 6.**
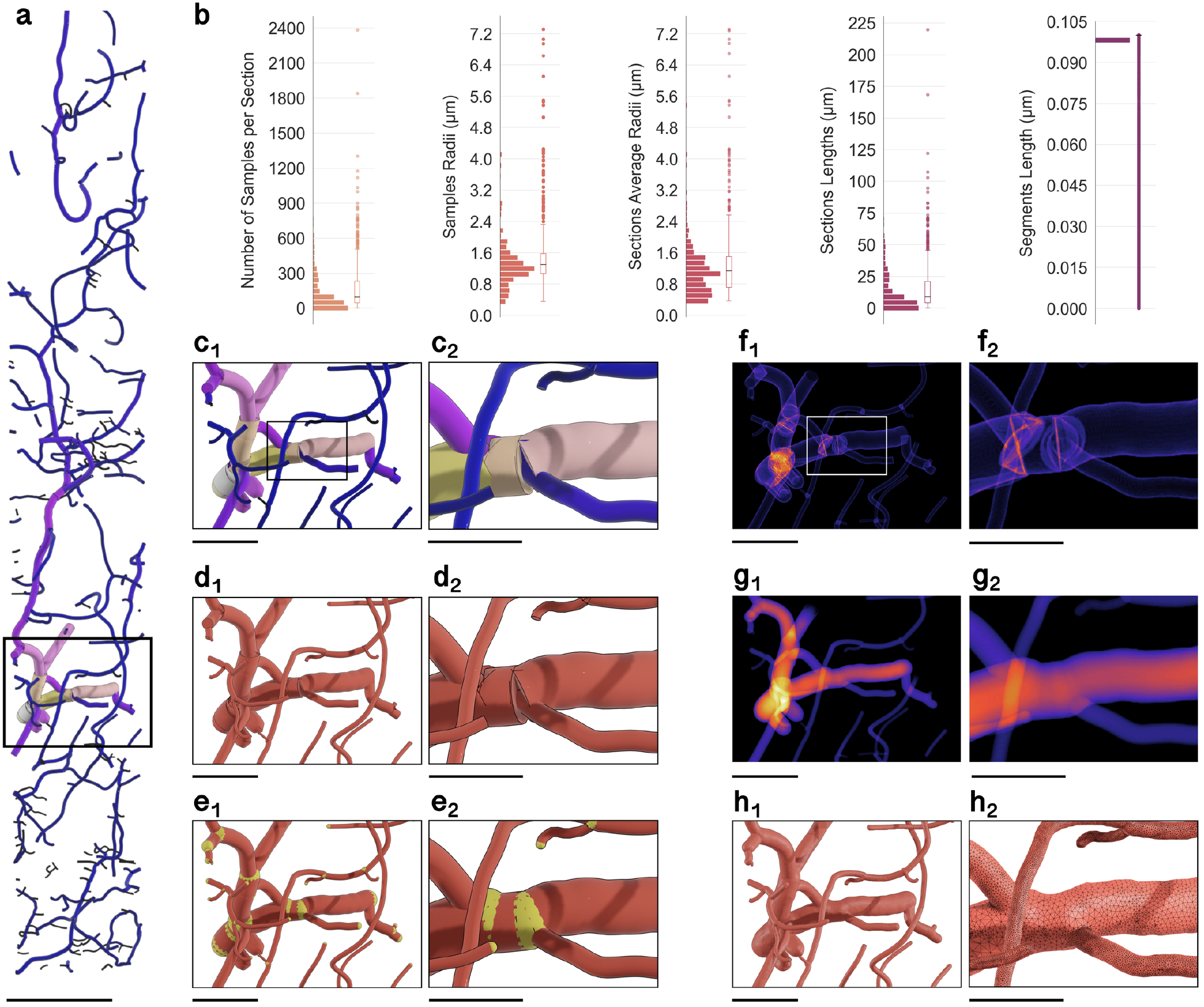
Reconstruction of a watertight mesh of a cerebral vascular network from its corresponding vectorized graph. (a) The data set is sliced from a larger cortical network with hundreds of millions of vertices to demonstrate how Ultraliser is effective in building mesh models with clean geometric topology and accurate branching structures from raw vectorized morphologies. (b) The data set is qualitatively and quantitatively analyzed to evaluate its local geometry and topology (**Supplementary Table S9**). (c) A closeup revealing the overlap between the different sections at a common branching point, where each section is assigned a different color based on its average radius. (d) Each section of morphology is converted into a tubular proxy mesh with a circular cross-section interpolated at every vertex along the section. (e) We add packing spheres (in yellow) at the terminal samples of each section to ensure the smoothness and continuity of the proxy mesh. (f) The proxy mesh is rasterized into a volumetric grid with a resolution of five microns per voxel, where the overlap between the different sections is obvious. (g) Applying the 3-way solid voxelization algorithm fills the intravascular space and removes any intersections, in which we can extract a continuous manifold of every partition in the volume. (h) A watertight mesh is reconstructed with clean geometric topology and accurate branching. This mesh can be then used in several simulation experiments. Scale bars, 100 *μ*m (**a**), 50 *μ*m (**c**_1_, **d**_1_, **e**_1_, **f**_1_, **g**_1_, **h**_1_), 25 *μ*m (**c**_2_, **d**_2_, **e**_2_, **f**_2_, **g**_2_, **h**_2_).

Due to the cyclic nature of vascular graphs, there is a high possibility that our slice-based solid voxelization algorithm will fail. We therefore use 3-way solid voxelization instead of 1-way voxelization, in which the flood-filling algorithm is applied on every principal axis independently. Prior to the optimization stage, the different partitions in the polygonized mesh object are split and optimized individually. After optimization, these partitions are regrouped again in a single mesh object (Methods). **Figure 6** depicts the stages of processing a fragmented vascular network towards reconstructing a high fidelity watertight surface mesh with multiple partitions. **Supplementary Figure S83** shows the results of converting a more complex vascular network into a watertight mesh with ultraVessMorpho2Mesh and ultimately into a tetrahedral one using TetGen^38^.

### 2.8 Generating annotated 3D tissue volumes from digital circuits

We also extended Ultraliser to create annotated voxel-based tissue models from surface meshes of neuronal morphologies. Voxel-based models are becoming essential, not only for visual analytics, but also for performing *in silico* optical imaging experiments that simulate light interaction with brain tissue using physically plausible Monte Carlo visualization^10^. Simulation applicability and even its accuracy are subject to two factors. First, the derivation of advanced mathematical formalism of the radiative transfer equation (RTE) that could account for absorption, scattering and, in certain cases, fluorescence. Second, the existence of biologically realistic 3D models that account for (i) the optical properties of brain tissue across its different regions at microscopic levels and (ii) the spectral properties of fluorescent dyes used in wet lab experiments for cell labeling.

What is missing then? RTE was extended to model light interaction with low- and high-scattering fluorescent participating media^35,81^, recent experiments were able to build accurate 3D brain atlases mapping the different optical properties of the tissue^82^, and fluorescence databases are available online, where spectral properties of common fluorescent dyes are provided. The only missing element is a robust application capable of creating detailed biologically and optically accurate volume models of cortical circuits. To address this issue, we implemented ultraCircuit2Volume.

Our implementation uses the information retrieved from the circuits that are digitally reconstructed by the Blue Brain Project^2,74^. These circuits identify neuron types, their coordinates and orientation. Starting from raw morphologies, corresponding surface meshes are generated either with Ultraliser directly or relying on third-party applications, such as NeuroMorphoVis^19^. Meshes are then rasterized in voxel grids, where each voxel is annotated with location-specific optical properties. In case of fluorescence, voxels corresponding to intracellular spaces are annotated with the spectral parameters of fluorescent dyes (Methods). The resulting volumes are used with recently developed *in silico* imaging simulators^10^ to create synthetic optical sections of cortical tissue models, on physically plausible basis (shown in **Fig. 7**).

**Figure 7.**
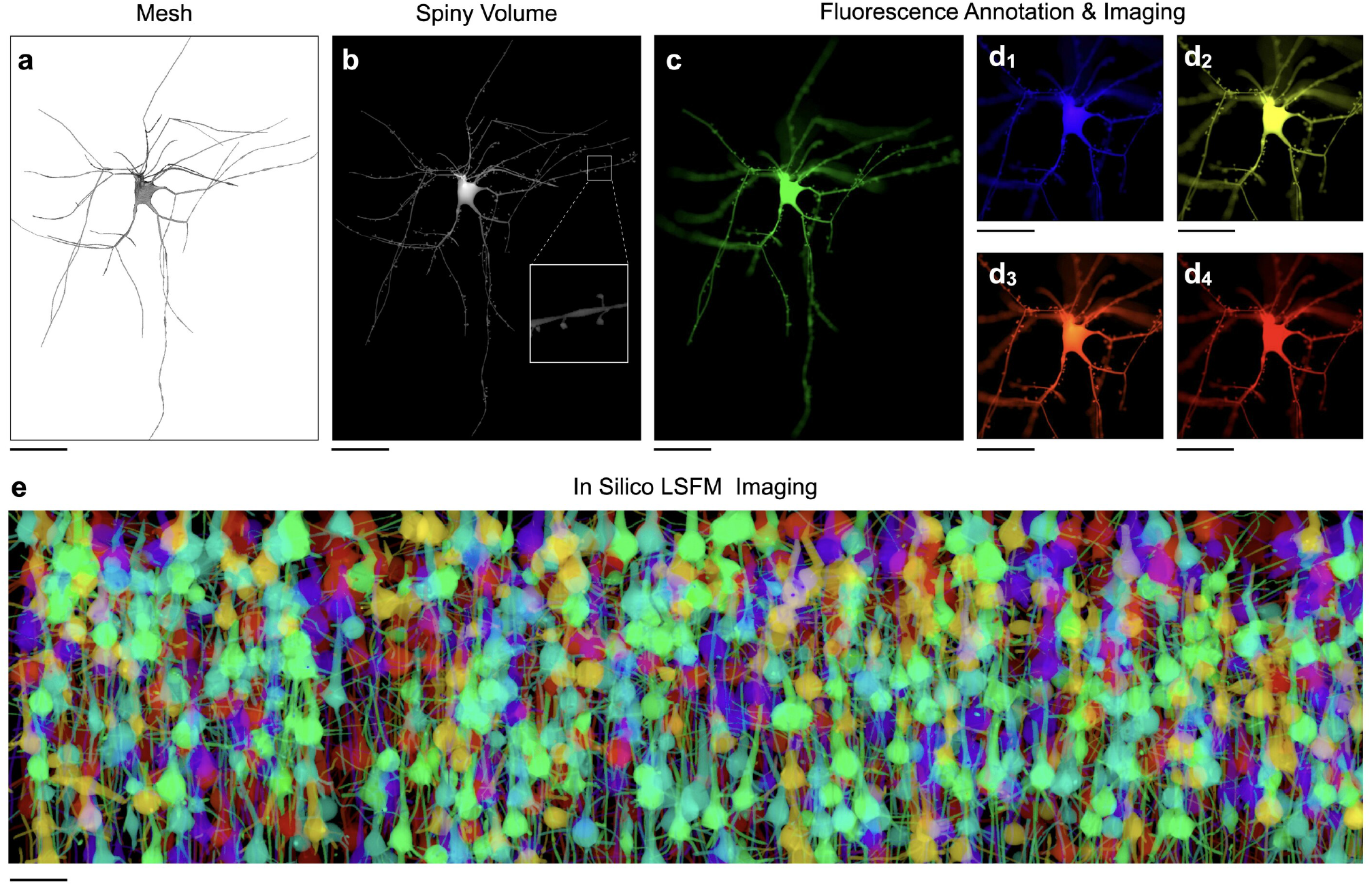
Creating annotated volumetric tissue models for *in silico* imaging. (a) The mesh is created using NeuroMorphoVis^19^ from a neuronal morphology reconstructed from the somatosensory cortex of a P14 rat^74^. (b) The neuron is placed into a digital circuit^2^ to determine its connectivity in which we can identify the synaptic locations and integrate the spines along its dendritic arborizations. (b) The neuron mesh is used to reconstruct a high resolution annotated volume in which its intracellular space is tagged with optical properties of multiple fluorescent dyes. The volume is then used for simulating the imaging of a single cell with fluorescence microscope. The following dyes are used Alexa Fluor 488, 405, 532, 658 and 610 in (c), (d_1_), (d_2_), (d_3_) and (d_4_) respectively. The simulated tissue block is illuminated with collimated laser beams at a wavelength that corresponds to the maximum excitation of each respective dye. (e) *In silico* brainbow optical section of a digitally reconstructed neocortical slice (920 × 640 × 1740 cubic microns) simulating the imaging of lightsheet fluorescence microscope. The slice is tagged with six fluorescent labels (GFP, CFP, eCFP, mBannan, mCheery and mPlum) and illuminated at the maximum excitation wavelength of each respective dye. Scale bars 25 *μ*m (**a-d**), 50 *μ*m (**e**).

### 2.9 Comparative analysis with existing frameworks

To demonstrate the critical significance of Ultraliser and its accompanying applications, we performed detailed quantitative and qualitative comparisons with relevant open source frameworks that are used for remeshing and mesh reconstruction from morphologial skeletons of neurons (**Supplementary Table S1**) and vascular networks (**Supplementary Table S2**). Comparative results and their analysis are discussed in detail in **Supplementary Section 13**. From the comparisons presented in **Supplementary Figures S85 - S89**, Ultraliser has demonstrated obvious superiority in terms of topology, tessellation, watertightness and its robust performance.

## 3 Conclusion

Biologically realistic simulations are indispensable for revealing the structure-function relationships within and among brain cells. Driven by a quantum leap in computing technologies, *in silico* brain research is complementing *in vivo* and *in vitro* methods. Meanwhile, advances in imaging technologies and automated ML-based segmentation algorithms are boosting the creation of detailed and anatomically realistic 3D neuroscientific models that are fueling simulation-based research. The goal of Ultraliser is to provide a systematic and robust framework for creating accurate watertight meshes and high resolution annotated volumes of 3D brain structures that can be integrated in multimodal supercomputer simulations. On one level, Ultraliser unconditionally rectifies non-watertight mesh models that cannot be repaired with existing remeshing solutions. On another level, it has native support to create ultrarealistic watertight meshes and annotated volumes of NGV models from their morphological descriptions. The framework has a modular and extensible architecture, making it possible to integrate further relevant applications that are of paramount importance in structural systems neuroscience.

## Methods

### Ultraliser: an overview

Ultraliser is an unconditionally robust and optimized framework dedicated primarily to *in silico* neuroscience research, allowing to generate high fidelity and multiscale (from subcellular and up to multicellular scales of resolution: 100 nm - 1 mm) 3D neuroscientific models — such as: nuclei, mitochondria, endoplasmic reticula, neurons, astrocytes, pericytes, neuronal branches with dendritic spines, minicolumns with thousands of neurons and large networks of cerebral vasculature — with realistic geometries. Ultraliser implements an effective voxelization-based remeshing engine that can rasterize non-watertight surface meshes —in the form of triangular soups— into high resolution volumes, with which we can reconstruct topologically accurate, adaptively optimized and watertight surface manifolds (**Supplementary Fig. S1**). In addition to their importance for accurate quantitative analysis, resulting models are primarily intended to automate the process of conducting supercomputer simulations of neuroscience experiments; complementing *in vivo* and *in vitro* techniques. Watertight triangular meshes are used for (i) performing 3D particle simulations, (ii) mesh-based skeletonization, in which accurate morphologies of cellular structures are obtained for performing 1D compartmental simulations and (iii) tetrahedralization, where we can generate tetrahedral volume meshes for 3D reaction-diffusion simulations. Annotated volumetric tissue models are also used in *in silico* imaging studies, where we can simulate optical imaging experiments with brightfield or fluorescence microscopy^10^. Ultraliser’s workflow is graphically illustrated in Figure 1 and a detailed schematic showing the ecosystem and relationship between its different modules is illustrated in **Supplementary Figure S2**.

The code is open sourced under the [GNU General Public License version 3.0 (GPL)](https://www.gnu.org/licenses/gpl-3.0.en.html), and is available for free on GitHub at https://github.com/BlueBrain/Ultraliser. Documentation and tutorials are available online at https://github.com/BlueBrain/Ultraliser/wiki. Quantitative and qualitative analysis scripts used in this study are also open sourced and integrated into NeuroMorphoVis^19^ (github.com/BlueBrain/NeuroMorphoVis). Images, movies and datasets produced in this study are publicly available on Zenodo (10.5281/zenodo.7105941).

### Data structures

Ultraliser is a C++ based library accompanied by several usecase-specific applications that can generate 3D models in two principal formats: watertight triangular surface manifolds and annotated, or tagged, volumes from a diverse set of input data formats including (i) digitized morphology skeletons, (ii) fragmented, self-intersecting and non-watertight polygonal surface meshes, (iii) binary volume masks, (iv) grayscale volumes, and (v) tetrahedral volume meshes (**Supplementary Fig. S1a**). Resulting watertight meshes can be further processed and converted into morphology skeletons or volume meshes relying on existing mesh-based skeletonization or tetrahedralization applications respectively.

#### Morphology skeletons

Neuroscientific 3D models with branching topologies —that are traced from optical microscopy stacks, for example: neurons, astrocytes and vasculature— are segmented, digitized and typically stored as morphology skeletons. 3D models acquired with electron microscopy^11,58^ are further processed and converted into morphology skeletons using skeletonization^83,84^. Neuronal morphologies are commonly stored in the standardized [SWC](http://www.neuronland.org/NLMorphologyConverter/MorphologyFormats/SWC/Spec.html) file format^16^. This format is also used to store processes of astroglial cells, but it does not account for any endfeet information, which is typically stored as trianglular patches and requires custom file formats that can combine branching and surface data^15,25^. The [SWC](http://www.neuronland.org/NLMorphologyConverter/MorphologyFormats/SWC/Spec.html) format is adopted by the NeuroMorpho.Org^16^ database, which contains hundreds of thousands of neuronal and astrocytic morphologies collected from a huge diversity of experiments. The [SWC](http://www.neuronland.org/NLMorphologyConverter/MorphologyFormats/SWC/Spec.html) format has been also adapted to store cerebral arterial arborizations, for instance the datasets of the [Brain Vasculature (BraVa)](http://cng.gmu.edu/brava/help.php)^18^ database (cng.gmu.edu/brava). Ultraliser has full support to load [SWC](http://www.neuronland.org/NLMorphologyConverter/MorphologyFormats/SWC/Spec.html) morphologies of neurons, astrocytes and cerebral vasculature. Moreover, it supports loading customized file formats such as (i) H5 morphologies that were defined within the scope of the Human Brain Project^85^ and (ii) the [VMV](https://github.com/BlueBrain/VessMorphoVis/wiki/File-Formats#vmv-file-format) format that is used to store vascular morphologies supported by [VessMorphoVis](https://github.com/BlueBrain/VessMorphoVis)^26^. Morphology skeletons are represented by a list of connected morphological samples, where each sample has a unique identifier, 3D Cartesian coordinate, cross-sectional radius, and optionally an index characterizing the type of the branch it belongs to. A pair of adjacent samples defines a segment or an edge, and a concatenated list of adjacent edges between two branching points defines a section or strand. In memory, morphologies are stored as a linear list of sections, where each section has a unique index and references to its parent and child sections. Those references are used to reconstruct the hierarchical organization of the morphology when required. The structures of neuronal, astrocytic and vascular morphologies are illustrated in **Supplementary Figures S76, S77** and **S78** respectively.

#### Polygonal surface meshes

Ultraliser has native support to import and export polygonal surface meshes with multiple file formats including those that are commonly used for visual analytics with Blender^86^ (blender.org) or MeshLab^87^ (meshlab.net), such as OBJ and PLY, and also those required for 3D printing, molecular simulations and conversion into tetrahedral meshes by tetrahedralization applications^38,39^ such as STL and OFF file formats. The meshes downloaded from the [MICrONS program](https://www.microns-explorer.org/)^48^ are stored in [H5](https://www.microns-explorer.org/phase1) files based on the HDF5^88^ library. Although this format is not standard, it is straightforward to reconstruct the triangulation of the mesh. Ultraliser implements two complementary data structures to store ‘triangular’ surface meshes. The first data structure Ultraliser::Mesh is a light version that only stores vertices and triangles. It has low memory footprint, allowing to perform most of the operations that do not necessitate any details of surface normals, edges or patch connectivity. The other data structure Ultraliser::AdvancedMesh is much more advanced and stores further information including surface normals, edges, connectivity between vertices, edges and triangles. This structure has been adopted from [MeshFix](https://github.com/MarcoAttene/MeshFix-V2.1)^89^ and adapted to address the essential requirements needed to accomplish the objectives of the framework. It is mainly used for repairing any geometric deficits in the mesh, detecting fragmented mesh partitions, removing self-intersecting triangles and also for watertighntess verification.

#### Volumetric models sampled on 3D Cartesian grids

Ultraliser processes and creates volume models sampled uniformly on 3D Cartesian grids. Ultraliser can import and export several volume formats including: 1-bit binary volumes (in BIT/HDR format), 8-, 16-, 32- and 64-bit volumes (in RAW/HDR and NRRD formats) and TIFF image stacks. The HDR file is an ASCII file that contains the volume dimensions, i.e. the number of voxels along each axis, and the precision of the data. Ultraliser::Volume implements three different types of grids. Ultraliser::BitVolumeGrid uses bit-arrays^90^ (Ultraliser::BitArray) to represent every voxel in the volume by a single bit in memory. The voxel is either set or unset; this is sufficient to voxelize the interior of a polygonal mesh. By default and unless otherwise specified, this data structure is used by the voxelization kernels, making it possible to process large scale volumes efficiently with reduced memory footprint. Ultraliser::UnsignedVolumeGrid stores every voxel in the volume in either 1, 2, 3, or 4 byte(s) in memory, allowing to define a grayscale volume. Ultraliser::VoxelGrid stores a list of attributes per voxel, for example: its value, an index representing the optical properties of a participating medium or a sub-region in the volume it belongs too. This data structure is mainly used for creating tagged volumes that are needed to create physically plausible visualizations that can simulate optical imaging experiments^35,81^.

### Voxelization and volume reconstruction

Voxlization is the process of creating 3D volumes of geometric models either from their parametric representations or from polygonal or polyhedral, tetrahedral and hexahedral meshes. Voxelization is classified into two categories: surface and solid voxelization. Surface voxelization creates volumetric shells representing boundaries of surface manifolds as a series of connected voxels–if no holes exist on the surface, while solid voxelization fills their interiors^91^.

#### Surface voxelization

We implemented a fast data-parallel surface voxelization algorithm, which sets all the voxels that overlap with any triangle in a given mesh using conservative rasterization^92^. In contrary to standard rasterization, the conservative criterion guarantees that a voxel is filled if it is partially overlapping or even touching a triangle. The algorithm can reconstruct a volumetric shell corresponding to the extent of a given triangle soup, even in the presence of self-intersecting triangles, non-manifold edges and non-manifold vertices^93^. The bounding box of each voxel is computed from its three-dimensional index and side length. For every triangle in the mesh, a box-triangle intersection test is performed to rasterize all the polygons in the mesh and create a volumetric shell that reflects the surface of the mesh^94^. Note that all the n-gons (n > 3) in the input mesh are automatically split into triangles prior to voxelization.

#### Solid voxelization

Conventional solid voxelization algorithms in computer graphics require a watertight manifold to successfully voxelize its interior into occupancy grids. By definition, a watertight mesh consists of a compact manifold that has clearly defined inside and does not contain any holes across its surface; that is if the surface is punctured with a hypodermic needle trying to fill it with water, it will not leak. A triangular mesh is guaranteed to be watertight – if and only if – it has no self-intersecting triangles, zero non-manifold edges, zero nonmanifold vertices, and no boundary edges (**Supplementary Section 4**). In reality, and unfortunately, neuroscientific mesh models segmented from microscopy stacks have ill topologies with hundreds or even thousands of self-intersections, non-manifold edges and vertices and even fragmented mesh partitions. Major contributions have been introduced to fix the topology of these meshes using geometric mesh conditioning^52,95^, nevertheless, their solutions are neither robust nor scalable, based on trials. Therefore, existing solid voxelization algorithms would fail to handle detailed mesh reconstructions with realistic geometries. Contrary to traditional methods, we present an efficient data-parallel, slice-based solid voxelization algorithm that does not entail an input watertight mesh. Initially, the surface voxelization algorithm converts a given triangular mesh into a volumetric shell in a uniformly sampled 3D Cartesian grid. The rasterization is binary, where each voxel in the grid is either set or cleared. The interior of the shell can be filled using 3D flood-filling. However, this algorithm is accompanied with extensive computational loads and cannot be easily parallelized. Our algorithm is based on 2D flood-filling that can be implemented in parallel. The 2D flood-filling kernel is applied independently to each slice in the volume. The aggregate result is exactly similar to what can be accomplished with 3D flood-filling, but in much less time.

#### 3-way solid voxelization

By default, the solid voxelization algorithm is applied on a per-slice-basis along the *Z-axis* of a given volume grid, where each slice is processed, or flood-filled, in a separate thread, independently. Certain structures, for example vascular morphologies – represented with cyclic graphs – have loops. In the general case, the flood-filling algorithm is unable to identify any internal boundaries beyond the first one detected. Therefore, running the flood-filling kernel along a single axis will fail to capture the entire geometry of an input mesh. To resolve this constraint, we implemented a 3-way solid voxelization algorithm which processes the volume along the X, Y and Z axes to produce three volume grids that are combined later with a logical *ANDing* operation to obtain the final grid. This approach makes it possible to resolve all the loops in a given cyclic structure.

### Mesh reconstruction

The binary volume resulting from the voxelization operation is then processed to reconstruct a smooth, optimized and watertight triangular mesh in four principal steps: (i) isosurface polygonization, (ii) Laplacian smoothing, (iii) adaptive or non-adaptive mesh optimization and (iv) watertightness verification.

#### Isosurface polygonization

Ultraliser integrates efficient implementations of two popular isosurface extraction algorithms: the default marching cubes (MC) algorithm^96^ and its superior, dual marching cubes (DMC)^97^. MC is relatively faster than DMC, but in certain cases it cannot reproduce rough surfaces with high frequency structures or sharp edges, while DMC can preserve thin surface features without excessive tessellation. The DMC algorithm reconstructs quadrilateral patches, but for consistency, every tetragon –or quadrilateral– created is divided and stored as two triangles with a shared edge. Moreover, MC cannot handle complex triangulation configurations, leading to self-intersecting faces and consequently non-watertight meshes. Adaptive optimization and watertightness verification are, however, implemented in subsequent stages whether any of the two algorithms is used for surface reconstruction. Therefore, using MC or DMC would ultimately yield an optimized and watertight mesh.

#### Surface smoothing

Due to the finite resolution of volumes, and their discretized nature, polygonal meshes reconstructed from those volumes exhibit zigzagged or ‘staircase’ artifacts on their surfaces (**Supplementary Fig. S1a**). Such artifacts distort the organic appearance of the resulting meshes, unless surface smoothing is applied. We therefore use the Laplacian operator to remedy those staircase artifacts in an iterative scheme. Initially, we build a list of neighbor vertices and faces for each vertex. In each iteration, it uses the aforementioned lists to compute a smoothing kernel for each vertex, applies it and then updates the mesh surface to prepare it for the next iteration. To compute the kernel, we identify the difference between the vertex and the arithmetic sum of the neighbour vertices, each weighted by their average cotangent. To compute the average cotangent, we use the two cotangents calculated from the two neighboring vertices of the edge formed by the vertex to the neighbour. To obtain the smoothed vertex, we linearly interpolate between the original vertex, and the original vertex weighted by the kernel. The interpolating parameter, also called the smoothing value, is an input to the algorithm chosen by the user, which must be greater than zero to have an effect. Additionally, an inflate parameter –also provided by the user– can be specified to dampen the shrinking effect on thin parts of the mesh. This parameter is used in the same form as the smoothing value, but with the opposite effect, and its value must be less than zero to have an effect.

#### Mesh optimization

Irrespective to the applied surface extraction algorithm, the tessellation of the reconstructed surface depends primarily on the resolution of the volume grid that is used to sample and voxelize the input mesh. For convenience, the resolution is set in terms of number of voxels per microns. The resolution is a free parameter that is either controlled by the user or automatically set based on the axis-aligned bounding box (AABB) of the input mesh, the size of the finest detail in the mesh and the scale or focus of the potential experiment in which the resulting mesh will be plugged in. Ultraliser takes advantage of binary volume grids, in which each voxel is represented in memory with a single bit; it is therefore capable of creating large scale volumes which can resolve the finest features of an object, for example: the ultrastructure of a dendritic spine in tall-tufted layer 5 pyramidal neurons^98^. With such resolutions, the reconstructed mesh is excessively tessellated, possibly with tens of millions of polygons, whose practicality is undoubtedly questionable. Ultraliser integrates a mesh optimization module that can adaptively refine highly tessellated meshes to create optimized counterparts with preserved features. The mesh optimization module extends an existing implementation^99^ that uses an angle-based approach for adaptive tessellation and normal-based smoothing to guarantee the quality of the resulting surface. The optimization process includes: surface smoothing, normal smoothing, flat coarsening, dense coarsening and also adaptive optimization.

#### Watertightness verification

The optimized meshes are guaranteed to have good topology and convenient polygonization, nevertheless, their watertightness is not guaranteed; the optimizer might introduce self-intersecting triangles depending on surface complexity and roughness. Watertightness has been addressed by extending an existing solution based on [MeshFix](https://github.com/MarcoAttene/MeshFix-V2.1)^89^ that uses a heuristic iterative approach that strives to reconstruct a single compact manifold with neither degeneracies nor self-intersections from a low quality input.

#### Meshes with multiple partitions

In certain cases, 3D models of cellular structures are not composed of a single and continuous object (or partition), but rather of multiple fragmented objects that could be spatially overlapping. This fragmentation is common due to labeling or tracing artifacts that arise during the segmentation of cellular models characterized with complex or thin structures^100^ such as astrocytes, neurons or microglia (refer to **Supplementary Table S3** - Partitions). Applying mesh reconstruction kernels to segmented volume stacks of such cellular models will result in polygonal surface meshes with multiple partitions, in which each partition is isolated as an independent set of polygons, nevertheless, and is still part of the mesh. Mesh analysis applications expect a watertight mesh to have a single partition represented as a set of connected vertices, edges and faces on a continuous manifold. Processing fragmented meshes with multiple partitions requires special handling to avoid generating incomplete or even non-watertight meshes. We allow the user to choose either to process the largest partition in the mesh and remove the other ones or to preserve all the partitions in the mesh. In the latter case, each partition is split and treated as an independent mesh object during the optimization. Afterwards, all the partitions are grouped together in a single mesh objects.

### Watertight mesh generation from input triangle soup

Triangular soups of fragmented non-watertight meshes are processed to create watertight counterparts in two steps: voxelization and isosurface polygonization. Initially, the input mesh is triangulated, in which each polygon (n-gon: n > 3) in the mesh is split into a list of corresponding triangles. The AABB of the mesh is then computed. Based on the dimensions of this AABB and the voxelization resolution (in voxels per micron) defined by the user, a binary 3D volume grid is created to cover the spatial extent of this AABB. The mesh is converted into a volumetric shell using surface voxelization, where every triangle along the surface of the mesh is independently rasterized in the volume grid. The interior of the resulting volumetric shell is then filled using solid voxelization; a 2D flood-filling kernel is applied independently to every slice in the grid along the Z-axis. If the input mesh was suspected to have loops, 3-way solid voxelization is then performed, in which the flood-filling kernel is applied along the X, Y and finally the Z axis. In the second stage, the binary volume grid is processed by a user-selected isosurface polygonization kernel (MC or DMC) to reconstruct a low-quality triangular mesh. This mesh is then post-processed to generate an optimized watertight manifold in three steps: (i) surface smoothing using the Laplacian operator, (ii) re-tesselation to remove unnecessary small triangles resulting from the polygonization process, and (iii) watertightness verification, ensuring that all self-intersecting triangles, boundary edges, floating vertices and non-manifold edges and vertices are removed.

### Watertight mesh generation from input morphology skeletons

The conversion of morphology skeletons into watertight meshes is performed in two steps: (i) creation of intermediate proxy meshes that can be accurately rasterized into volumetric grids and (ii) applying the remeshing routine used to create watertight meshes from triangle soups. These proxy meshes are known to be spatially overlapping and self-intersecting, but they are only used to rasterize the geometry of every section in the morphology into the volume grid. We implemented two algorithms to build these proxy-meshes. The first one converts every section in the morphology into an independent mesh. Each mesh is overlapping with its adjacent ones that correspond to parent and child sections in the morphology. To guarantee the continuity between the neighboring sections at their branching points, packing spheres are added. The radius of every sphere is computed based on the largest terminal sample of all the sections connected at the respective branching point. This algorithm is optimum to reconstruct vasculature meshes from their corresponding morphologies and, in general, can be applied to handle structures with cyclic graphs. The other algorithm computes the longest connected paths along the graph of an input morphology to create a proxy mesh, not on a per-section-basis, but rather on a per-path-basis. Every path is a continuous list of samples that can represent an individual section or an aggregate of two adjacent sections or more. This algorithm is well suited to handle morphologies with directed acyclic graphs, including neuronal arborizations and astroglial processes. As illustrated in **Supplementary Figures S76, S77** and **S78**, each section in the morphology is composed of a sequence of samples, each defines a position and radius. For each section, or path, a mesh is reconstructed by resampling the corresponding segments using the cubic Hermite spline interpolation. The positions and tangents of the new samples are defined by **Supplementary Equations 1** and **2**. To avoid loops or self-intersections between the different sections in the reconstructed mesh, the tangent at each original segment point is computed using the Centripetal Catmull-Rom spline formulation, that uses the positions of the previous and next samples to the current segment, as shown by **Supplementary Equations 3** and **4**. Once all new samples of the section are computed, a sectional geometry in the form of circumference is used to interpolate along the path to construct a connected vertex assembly in the form of a tubular mesh (**Supplementary Section 9**).

### Watertight mesh generation from input volumes

Input volumes are directly converted into watertight meshes using isosurface polygonization followed by watertightness verification. 1-bit volumes, binary volumes or segmented masks are directly processed to reconstruct a surface mesh. However, n-bit grayscale volumes, where n is 8, 16, 32 or 64, require specifying an additional parameter to complete the process: the isovalue, with which an isosurface is segmented and used for surface reconstruction.

### Volume generation from input meshes or morphologies

Volume generation is implicit; it is automatically implemented within the remeshing pipeline during the voxelization stage. Unless specified, resulting volumes are binary, in which every voxel is represented by a single bit, and therefore, these volumes are not annotated to account for any variations across the spatial extent of the volume. To create annotated volumes, Ultraliser::VoxelGrids are used for voxelization, in which we can assign annotation indices to every voxel in the grid. Volumes can be exported into BIN (1-bit), RAW (8-bit) and NRRD (8-bit) files.

### Tetrahedralization

Ultraliser reads tetrahedral meshes for the purpose of data conversion between formats, i.e. to create watertight surface meshes and volumes from tetrahedral inputs using ultraTet2Surface. However, it does not implement any tetrahedral mesh generators within its pipeline to create meshes in a direct manner. For this purpose, we rely on existing implementations, mainly [TetGen](https://wias-berlin.de/software/tetgen/features.html)^38^ (tetgen.org) and [Gmsh](https://gmsh.info/)^40^ (gmsh.info), which can complement our pipeline to create tetrahedral volumetric meshes from the watertight meshes created by Ultraliser.

### Generating biologically realistic neuronal meshes from digitized morphologies

A 3D somatic profile is created based on a finite element method (FEM) approach^73^. The algorithm takes into account the coordinates of the initial segments of the neurites that only emanate from to the soma. The connected neurites are identified in a pre-processing step, in which the distance between their initial segments to the center of the soma is evaluated and relatively compared with respect to its average radius. The soma is initially modeled by a tetrahedral icosahedron (or icosphere) approximating the mean radius of the soma. Projective mapping is then applied, where cross-sectional areas of the initial segments of the connected neurites are projected onto the ico-sphere. Vertices located within every projection surface are selected and grouped together to identify their center. Simultaneously, a pulling force is applied at every center to deform the ico-sphere towards the neurites, giving it a realistic profile.

Every tree corresponding to an individual neurite in the morphology is then processed and converted into a proxy mesh. As aforementioned, proxy meshes are not watertight, but they are essential to reconstruct a volumetric shell in the following voxelization stage. Mesh branching at bifurcation (or trifurcation) points is not explicitly implemented; most mesh branching algorithms at small branching angles (less than 30°) fail to reconstruct an organic and accurate bifurcation geometry. To guarantee continuous branching, whatever the conditions at the bifurcation point, we implemented an exhaustive algorithm that builds all the possible paths starting from the soma and until the terminal segments. At every section in the morphology, the algorithm reconstructs all the path combinations between the section itself, its parent section and the child sections. These formed paths are considered as independent polylines with thickness. Each segment of the path is resampled using the cubic Hermite spline interpolation to compute the positions and tangents of the new generated intermediate nodes. Once all new nodes of the path are computed, to mesh the complete polyline, a sectional geometry in the form of circumference is placed at the origin of coordinates of the plane defined by the position and the tangent at each node. The tangent of the node is taken as the normal vector of the plane. Finally, since the sectional geometry keeps the same number of vertices along the path, the vertex assembly consists in a simple connection between the already sorted vertices.

To improve the realism of the resulting mesh models, we extracted ∼50 spine meshes (**Supplementary Fig. S79**) from the four neurons shown in **Supplementary Figures S10 - S13**. Each neuronal mesh is loaded in Blender, where spine geometries are visually identified. Afterwards, and for each spine, we created a bounding box covering its spatial extent and overlapping with the dendritic section it emanates from. We then applied, per spine, a mesh intersection operator to extract its geometry as an independent object. The spines are oriented along the Y-axis after identifying their base and apex. Spine geometries are then processed to clean any self-intersecting facets along their surfaces, optimized and finally exported as independent mesh objects.

### Generating astroglial meshes from complete synthetic morphologies

In terms of representation, processes of astroglial cells are similar to neuronal arborizations, except that they have relatively compact extents and star-shaped structure, and are excessively oversampled in certain cases^15^. Moreover, astrocytic morphologies contain triangular surface patches that represent their endfeet geometries (**Supplementary Fig. S77**). Therefore, the same routine used to reconstruct neuronal meshes is applied to build astrocytic somata and processes. We then extended the implementation to generate endfeet proxy meshes using implicit surfaces. Endfeet patches are composed of a set of connected triangles and their respective vertices, and each vertex has a specific diameter that accounts for thickness at this particular vertex. Implicit surface modeling requires sufficient vertex density to avoid fragmented mesh partitions. Accordingly, we resample the surface of every endfoot patch, in which the distance between any two connected vertices across the patch is greater than the thickness of the endfoot. Following to the rasterization of the somatic and processes proxy meshes, resulting endfeet proxy meshes are rasterized to create a continuous volume shell of the entire astrocyte morphology. Surface reconstruction routines (MC or DMC) are directly applied on the resulting volume shell without applying solid voxelization. During the surface optimization process, all the internal mesh partitions are automatically removed. The partition with largest surface area (or tessellation), which represents the astrocytic membrane, remains.

### Generating continuous cellular meshes from fragmented components

In general, ultraMeshes2Mesh can compile a group of individual meshes into a single mesh object. If the input meshes are not spatially overlapping at all, then the resulting mesh will be composed of multiple partitions. In case that all the input meshes are overlapping, the output mesh will have a single partition with a continuous manifold. We therefore can use this application to generate ultrarealistic structural mesh models of neurons or astrocytes relying on a combination of already existing methods that are summarized in **Supplementary Table S1**. For neurons, we can use the [Soma Reconstruction Toolbox](https://github.com/BlueBrain/NeuroMorphoVis/wiki/Soma-Reconstruction) in NeuroMorphoVis^19^ and create a plausible somatic mesh. Skin modifiers in Blender (blender.org) can also be used to generate neuronal arborizations with organic-looking or realistic branching structure^24^. These individual meshes —of the soma and arborizations— can be combined together to generate a single mesh object with a continuous cellular surface. If spine meshes are available^101^, even at a later stage, we can also integrate them along the dendritic surface of the resulting mesh to create an integrated spiny mesh model of the neuron (as shown in **Supplementary Figure S81**). Nevertheless, it is the responsibility of the user to ensure that all the meshes of the cellular components (soma, neurites and spines) are spatially overlapping without having any gaps either between parent and child sections or between the soma and the all branches that originate from it, to be able to establish a single continuous surface manifold. ultraMeshes2Mesh loads a list of meshes grouped in a single input directory and computes an aggregate bounding volume, with which we can identify the spatial extent of the resulting mesh. Based on the voxelization resolution specified by the user, a binary volume grid is created, where the input meshes are rasterized, in parallel. The interior of the grid is filled with solid voxelization to create a homogeneous volume, with which the surface mesh is reconstructed, optimized and verified to be watertight.

### Creating vasculature meshes from corresponding graph networks

Frequently, vascular skeletons vectorized from optical microscopy stacks are excessively oversampled. We accordingly apply adaptive resampling to every section in the morphology to remove any unnecessary samples while preserving its structure. In certain cases, resampling reduces the total number of samples by 60-70%, thus lessening the tessellation of the proxy meshes created on a per-section-basis. To avoid fragmentation artifacts, we identify the samples with the least radii across the entire morphology, with which we can identify the most convenient voxelization resolution needed to preserve the integrity of the final vascular mesh, avoiding the structural fragmentation that arise due to surface smoothing and optimization. Samples with comparatively small radii –or zero-radius samples– are interpolated, and short sections with zero-length edges are eliminated. To ensure continuity between interconnected sections, i.e. smooth and accurate branching geometries, we add packing spheres (explicit ico-spheres) at the terminal samples of each section. Proxy geometries are then created on a per-section-basis and rasterized in a volume grid. Packing spheres are also rasterized to yield a continuous shell of voxels for every partition in the morphology. To avoid flood-filling vascular loops, 3-way solid voxelization is applied. The resulting volume grid is finally used for mesh reconstruction and optimization.

## Data sources

Cellular and subcellular NGV meshes segmented from the volume shown in **Figure 2** are provided by the collaborating co-authors affiliated with KAUST. Neuronal meshes shown in **Figure 3, Supplementary Figures S55 - S75** and **Supplementary Figures S85** are publicly available from the [MICrONS](https://www.microns-explorer.org/) program^48^. Neuronal morphologies shown in **Figure 4, Supplementary Figures S80 - S81** and **Supplementary Figure S86** are publicly available from NeuroMorpho.Org^16^. Astrocytic morphologies (**Figure 5** and **Supplementary Figure S82**) are provided by Eleftherios Zisis^15^. Vascular morphologies (rat’s cerebral microvasculature) shown in **Figure 6** and **Supplementary Figures S83 - S84** are courtesy of [Bruno Weber](https://www.neuroscience.uzh.ch/en/about/people/steering/Weber.html)^22^, University of Zürich (UZH). The vascular morphology of the arterial arborizations shown in **Supplementary Figure S88** is available from the [Brain Vasculature (BraVa)](http://cng.gmu.edu/brava/home.php?s=1&name_browser=false) database^18^ (cng.gmu.edu/brava).

## Supplementary Data

**Supplementary Data 1** contains the input (non-watertight) surface meshes of the block (shown in **Figure 2a**) reconstructed within the context of the EPFL-KAUST collaboration, and the corresponding output (watertight) meshes generated by Ultraliser. **Supplementary Data 2** contains a set of 20 non-watertight meshes that were randomly selected from the block shown in **Supplementary Figure S54** and another set of the their watertight counterparts. **Supplementary Data 3** contains a set of 25 neuronal morphologies with different morphological types and their corresponding watertight meshes. **Supplementary Data 4** contains a set of 25 synthetic astroglial morphologies^15^ and their corresponding watertight meshes. **Supplementary Data 5** contains the vascular morphology (shown in **Supplementary Fig. S83**) and a corresponding multi-partitioned watertight mesh. **Supplementary Data 6** contains the datasets used for the comparative analysis shown in **Supplementary Section 13**. Neuronal, astrocytic and vascular morphologies are stored in [SWC](http://www.neuronland.org/NLMorphologyConverter/MorphologyFormats/SWC/Spec.html), H5 and [VMV](https://github.com/BlueBrain/VessMorphoVis/wiki/File-Formats) file formats respectively. The file structures of the SWC and VMV formats are publicly available online. The H5 files of the complete astrocyte cells can be made available from corresponding authors upon request. All the surface meshes are stored either in Wavefront OBJ or in PLY files. Additional STL meshes are generated to be used for [TetGen](https://wias-berlin.de/software/tetgen/features.html) to create corresponding tetrahedral meshes. All the input and generated data files are publicly available on Zenodo (10.5281/zenodo.7105941).

## Software availability

Ultraliser is developed entirely in C++. The data-parallel sections of the code are parallelized using OpenMP^102^. The code is released to public as an open-source software (OSS) in accordance with the regulations of the [Blue Brain Project, École polytechnique fédérale de Lausanne (EPFL)](https://www.epfl.ch/research/domains/bluebrain) for open sourcing under the [GNU GPL3](https://www.gnu.org/licenses/gpl-3.0.en.html) license. The code is freely available online at https://github.com/BlueBrain/Ultraliser. The version of the code used to create all the results demonstrated in this study is available in the **Supplementary Software**.

## Acknowledgments

We thank Grigori Chevtchenko and Samuel Lapere for the impactful discussions on watertight meshing, Pawel Podhajski on technical assistance to deploy the software on Blue Brain 5. We also thank Karin Holm and Judit Planas for their valuable comments on the manuscript. We acknowledge the artistic touch of Elvis Boci to improve the quality of the figures.

## Authors’ contributions

M.A. and F.S. co-conceived the study. M.A., M.H., H.M, P.M., F.S. co-led the study. M.A. designed and implemented the framework, reconstructed and analyzed the resulting models and wrote the manuscript with input and critique from all authors. JJ.G.C. implemented the neuronal mesh reconstruction algorithm with FEM simulation, implemented the astrocytic endfeet proxy mesh reconstruction using implicit surfaces and contributed to the manuscript. N.R.G implemented the surface smoothing filters and contributed to the manuscript. A.F. implemented the morphology and mesh analysis code and provided technical assistance to load the H5 meshes downloaded from [MICrONS program](https://www.microns-explorer.org/). JS.C. contributed to the simulation discussions and the manuscript. C.C. reconstructed the NGV datasets within the EPFL-KAUST collaboration, analyzed them and contributed to the manuscript. M.Ag. contributed to the meshing discussions. E.Z. processed and provided the vasculature morphologies based on the original data set that was reconstructed by [Bruno Weber (UZH)](https://www.pharma.uzh.ch/en/research/functionalimaging.html), created the synthetic astrocytic morphologies and provided assistance to load them in Python. D.K. contributed to the discussions on the simulation requirements and the watertight meshing performance. All authors reviewed the manuscript.

## Authors’ Biography

**Marwan Abdellah** is a senior visualization research engineer at the Blue Brain Project of the École polytechnique fédérale de Lausanne (EPFL). He received his Ph.D. in neuroscience from EPFL in 2017.

**Juan José García Cantero** is a post-doctroal fellow at the Blue Brain Project of the École polytechnique fédérale de Lausanne (EPFL).

**Nadir Román Guerrero** is a visualization engineer at the Blue Brain Project of the École polytechnique fédérale de Lausanne (EPFL).

**Alessandro Foni** is a system specialist at the Blue Brain Project of the École polytechnique fédérale de Lausanne (EPFL).

**Jay S. Coggan** is a senior scientist in the Molecular Systems group in the Simulation Neuroscience division at the Blue Brain Project of the École polytechnique fédérale de Lausanne (EPFL).

**Corrado Calì** is a group leader and assistant professor RTDB in Human Anatomy at the Neuroscience Institute Cavalieri Ottolenghi (NICO) - Unito.

**Marco Agus** is an assistant professor at the College of Science and Engineering, Hamad Bin Khalifa University (HBKU).

**Eleftherios Zisis** is a software engineer at the Blue Brain Project of the École polytechnique fédérale de Lausanne (EPFL).

**Daniel Keller** is the group leader of the Molecular Systems group at the Blue Brain Project of the École polytechnique fédérale de Lausanne (EPFL).

**Markus Hadwiger** is an associate professor in computer science and the Visual Computing Center (VCC) at King Abdullah University of Science and Technology (KAUST), leading the High-Performance Visualization research group at VCC.

**Pierre J. Magistretti** is a full professor at the Brain Mind Institute of the École polytechnique fédérale de Lausanne (EPFL), director of the Center for Psychiatric Neuroscience, Department of Psychiatry / CHUV, a distinguished professor and vice president for research and the director of the KAUST Smart Health Initiative at King Abdullah University of Science and Technology (KAUST).

**Henry Markram** is a professor of neuroscience at the École polytechnique fédérale de Lausanne (EPFL), director of the Laboratory of Neural Microcircuitry (LNMC) and the founder and director of the Blue Brain Project.

**Felix Schürmann** is adjunct professor at the École Polytechnique Fédérale de Lausanne, co-director of the Blue Brain Project and involved in several research challenges of the European Human Brain Project of the École polytechnique fédérale de Lausanne (EPFL).

## Additional information

**Supplementary information** is available in the attached **Supplementary Document.**

**Correspondence and requests for materials** should be addressed to M.A and F.S.

## References

1. Y Cajal, S. R. Histologie du système nerveux de l’homme & des vertébrés: Cervelet, cerveau moyen, rétine, couche optique, corps strié, écorce cérébrale générale & régionale, grand sympathique (A. Maloine, 1911).

2. Markram, H., Muller, E., Ramaswamy, S., Reimann, M. W., Abdellah, M., et al. Reconstruction and simulation of neocortical microcircuitry. Cell 163, 456–492 (2015).

3. Di Ventura, B., Lemerle, C., Michalodimitrakis, K. & Serrano, L. From in vivo to in silico biology and back. Nature 443, 527–533 (2006).

4. Markram, H. The Blue Brain Project. Nature Reviews Neuroscience 7, 153–160 (2006).

5. Gerstner, W., Sprekeler, H. & Deco, G. Theory and simulation in neuroscience. science 338, 60–65 (2012).

6. Hunt, C. A. et al. At the biological modeling and simulation frontier. Pharmaceutical research 26, 2369–2400 (2009).

7. Mahajan, G. & Nadkarni, S. Intracellular calcium stores mediate metaplasticity at hippocampal dendritic spines. The Journal of physiology 597, 3473–3502 (2019).

8. Coggan, J. S. et al. Evidence for ectopic neurotransmission at a neuronal synapse. Science 309, 446–451 (2005).

9. Donohue, D. E. & Ascoli, G. A. Automated reconstruction of neuronal morphology: an overview. Brain research reviews 67, 94–102 (2011).

10. Abdellah, M. In Silico Brain Imaging: Physically-plausible Methods for Visualizing Neocortical Microcircuitry, 400. http://infoscience.epfl.ch/record/232444 (2017).

11. Cali, C. et al. 3D cellular reconstruction of cortical glia and parenchymal morphometric analysis from Serial Block-Face Electron Microscopy of juvenile rat. Progress in neurobiology 183, 101696 (2019).

12. Ascoli, G. A. & Krichmar, J. L. L-Neuron: a modeling tool for the efficient generation and parsimonious description of dendritic morphology. Neurocomputing 32, 1003–1011 (2000).

13. Cuntz, H., Forstner, F., Borst, A. & Häusser, M. One rule to grow them all: a general theory of neuronal branching and its practical application. PLoS computational biology 6, e1000877 (2010).

14. Kanari, L. et al. Computational synthesis of cortical dendritic morphologies. Cell Reports 39, 110586 (2022).

15. Zisis, E. et al. Digital Reconstruction of the Neuro-Glia-Vascular Architecture. Cerebral Cortex 31, 5686–5703 (2021).

16. Ascoli, G. A., Donohue, D. E. & Halavi, M. NeuroMorpho. Org: a central resource for neuronal morphologies. Journal of Neuroscience 27, 9247–9251 (2007).

17. Blinder, P. et al. The cortical angiome: an interconnected vascular network with noncolumnar patterns of blood flow. Nature neuroscience 16, 889–897 (2013).

18. Wright, S. N. et al. Digital reconstruction and morphometric analysis of human brain arterial vasculature from magnetic resonance angiography. Neuroimage 82, 170–181 (2013).

19. Abdellah, M. et al. NeuroMorphoVis: a collaborative framework for analysis and visualization of neuronal morphology skeletons reconstructed from microscopy stacks. Bioinformatics 34, i574–i582 (2018).

20. Hines, M. L. & Carnevale, N. T. The NEURON simulation environment. Neural computation 9, 1179–1209 (1997).

21. McCormick, D. A., Shu, Y. & Yu, Y. Hodgkin and Huxley model—still standing? Nature 445, E1–E2 (2007).

22. Reichold, J. et al. Vascular graph model to simulate the cerebral blood flow in realistic vascular networks. Journal of Cerebral Blood Flow & Metabolism 29, 1429–1443 (2009).

23. Feiger, B. et al. Determining the impacts of venoarterial extracorporeal membrane oxygenation on cerebral oxygenation using a one-dimensional blood flow simulator. Journal of biomechanics 104, 109707 (2020).

24. Abdellah, M., Favreau, C., Hernando, J., Lapere, S. & Schürmann, F. Generating High Fidelity Surface Meshes of Neocortical Neurons using Skin Modifiers in Computer Graphics and Visual Computing (CGVC) (eds Vidal, F. P., Tam, G. K. L. & Roberts, J. C.) (The Eurographics Association, 2019). ISBN: 978-3-03868-096-3.

25. Abdellah, M. et al. Metaball skinning of synthetic astroglial morphologies into realistic mesh models for visual analytics and in silico simulations. Bioinformatics 37, i426–i433 (2021).

26. Abdellah, M. et al. Interactive visualization and analysis of morphological skeletons of brain vasculature networks with VessMorphoVis. Bioinformatics 36, i534–i541 (2020).

27. Stiles, J. R., Bartol, T. M., et al. Monte Carlo methods for simulating realistic synaptic microphysiology using MCell. Computational neuroscience: realistic modeling for experimentalists, 87–127 (2001).

28. Khan, D., Yan, D.-M., Gui, S., Lu, B. & Zhang, X. Molecular surface Remeshing with local region refinement. International journal of molecular sciences 19, 1383 (2018).

29. Hu, Y. et al. Tetrahedral meshing in the wild. ACM Trans. Graph. 37, 60–1 (2018).

30. Li, X., Zhou, Z. & Kleiven, S. An anatomically detailed and personalizable head injury model: significance of brain and white matter tract morphological variability on strain. Biomechanics and modeling in mechanobiology 20, 403–431 (2021).

31. Hepburn, I., Chen, W., Wils, S. & De Schutter, E. STEPS: efficient simulation of stochastic reaction–diffusion models in realistic morphologies. BMC systems biology 6, 1–19 (2012).

32. Andrews, S. S., Addy, N. J., Brent, R. & Arkin, A. P. Detailed simulations of cell biology with Smoldyn 2.1. PLoS Comput Biol 6, e1000705 (2010).

33. Andrews, S. S. in Bacterial Molecular Networks 519–542 (Springer, 2012).

34. Robinson, M., Andrews, S. S. & Erban, R. Multiscale reaction-diffusion simulations with Smoldyn. Bioinformatics 31, 2406–2408 (2015).

35. Abdellah, M., Bilgili, A., Eilemann, S., Markram, H. & Schürmann, F. Physically-based in silico light sheet microscopy for visualizing fluorescent brain models. BMC bioinformatics 16, S8 (2015).

36. Pharr, M., Jakob, W. & Humphreys, G. Physically based rendering: Fromtheory to implementation Third edition. ISBN: 0128006455. http://pbrt.org (Morgan Kaufmann, 2016).

37. Botsch, M., Kobbelt, L., Pauly, M., Alliez, P. & Lévy, B. Polygon mesh processing (CRC press, 2010).

38. Si, H. TetGen, a Delaunay-based quality tetrahedral mesh generator. ACM Transactions on Mathematical Software (TOMS) 41, 1–36 (2015).

39. Labelle, F. & Shewchuk, J. R. in ACM SIGGRAPH 2007 papers 57–es (2007).

40. Geuzaine, C. & Remacle, J.-F. Gmsh: A 3-D finite element mesh generator with built-in pre-and post-processing facilities. International journal for numerical methods in engineering 79, 1309–1331 (2009).

41. Fabri, A. & Pion, S. CGAL: The computational geometry algorithms library in Proceedings of the 17th ACM SIGSPATIAL international conference on advances in geographic information systems (2009), 538–539.

42. Hu, Y., Schneider, T., Wang, B., Zorin, D. & Panozzo, D. Fast tetrahedral meshing in the wild. ACM Transactions on Graphics (TOG) 39, 117–1 (2020).

43. Narayanaswamy, A. et al. Robust adaptive 3-D segmentation of vessel laminae from fluorescence confocal microscope images and parallel GPU implementation. IEEE transactions on medical imaging 29, 583–597 (2009).

44. Tagliasacchi, A., Alhashim, I., Olson, M. & Zhang, H. Mean curvature skeletons in Computer Graphics Forum 31 (2012), 1735–1744.

45. Damseh, R. et al. Automatic graph-based modeling of brain microvessels captured with two-photon microscopy. IEEE journal of biomedical and health informatics 23, 2551–2562 (2018).

46. Januszewski, M. et al. High-precision automated reconstruction of neurons with flood-filling networks. Nature methods 15, 605–610 (2018).

47. Konishi, K. et al. Practical method of cell segmentation in electron microscope image stack using deep convolutional neural network. Microscopy 68, 338–341 (2019).

48. Bae, J. A. et al. Functional connectomics spanning multiple areas of mouse visual cortex. bioRxiv (2021).

49. Zheng, Z. et al. A complete electron microscopy volume of the brain of adult Drosophila melanogaster. Cell 174, 730–743 (2018).

50. Xu, C. S. et al. A connectome of the adult drosophila central brain. BioRxiv (2020).

51. Dorkenwald, S. et al. FlyWire: online community for whole-brain connectomics. Nature methods 19, 119–128 (2022).

52. Lee, C. T. et al. 3D mesh processing using GAMer 2 to enable reaction-diffusion simulations in realistic cellular geometries. PLoS computational biology 16, e1007756 (2020).

53. Edwards, J. et al. VolRoverN: enhancing surface and volumetric reconstruction for realistic dynamical simulation of cellular and subcellular function. Neuroinformatics 12, 277–289 (2014).

54. Garcia-Cantero, J. J., Brito, J. P., Mata, S., Bayona, S. & Pastor, L. NeurotessMesh: A tool for the Generation and Visualization of Neuron Meshes and Adaptive on-the-Fly Refinement. Frontiers in neuroinformatics 11, 38 (2017).

55. Brito, J. P. et al. Neuronize: a tool for building realistic neuronal cell morphologies. Frontiers in neuroanatomy 7 (2013).

56. Mörschel, K., Breit, M. & Queisser, G. Generating neuron geometries for detailed three-dimensional simulations using anamorph. Neuroinformatics 15, 247–269 (2017).

57. Lee, C. T. et al. An open-source mesh generation platform for biophysical modeling using realistic cellular geometries. Biophysical journal 118, 1003–1008 (2020).

58. Coggan, J. S. et al. A Process for Digitizing and Simulating Biologically Realistic Oligocellular Networks Demonstrated for the Neuro-Glio-Vascular Ensemble. Frontiers in neuroscience 12 (2018).

59. Wils, S. & De Schutter, E. STEPS: modeling and simulating complex reaction-diffusion systems with Python. Frontiers in neuroinformatics 3, 15 (2009).

60. Chen, W. & De Schutter, E. Parallel STEPS: large scale stochastic spatial reaction-diffusion simulation with high performance computers. Frontiers in Neuroinformatics 11, 13 (2017).

61. Coggan, J. S., Bittner, S., Stiefel, K. M., Meuth, S. G. & Prescott, S. A. Physiological dynamics in demyelinating diseases: unraveling complex relationships through computer modeling. International journal of molecular sciences 16, 21215–21236 (2015).

62. Ascoli, G. A., Maraver, P., Nanda, S., Polavaram, S. & Armañanzas, R. Win–win data sharing in neuroscience. Nature methods 14, 112–116 (2017).

63. Takemura, S.-y. et al. Synaptic circuits and their variations within different columns in the visual system of Drosophila. Proceedings of the National Academy of Sciences 112, 13711–13716 (2015).

64. Kasthuri, N. et al. Saturated reconstruction of a volume of neocortex. Cell 162, 648–661 (2015).

65. Dorkenwald, S. et al. Automated synaptic connectivity inference for volume electron microscopy. Nature methods 14, 435–442 (2017).

66. Dorkenwald, S. et al. Binary and analog variation of synapses between cortical pyramidal neurons. bioRxiv (2019).

67. Kumbhar, P. et al. CoreNEURON: an optimized compute engine for the NEURON simulator. Frontiers in neuroinformatics 13, 63 (2019).

68. Awile, O. et al. Modernizing the NEURON Simulator for Sustainability, Portability, and Performance. Frontiers in Neuroinformatics 16 (2022).

69. Grein, S., Stepniewski, M., Reiter, S., Knodel, M. M. & Queisser, G. 1D-3D hybrid modeling—from multi-compartment models to full resolution models in space and time. Frontiers in neuroinformatics 8, 68 (2014).

70. Lasserre, S. et al. A neuron membrane mesh representation for visualization of electrophysiological simulations. IEEE Transactions on Visualization and Computer Graphics 18, 214–227 (2012).

71. McDougal, R. A., Hines, M. L. & Lytton, W. W. Water-tight membranes from neuronal morphology files. Journal of neuroscience methods 220, 167–178 (2013).

72. McDougal, R. A. & Shepherd, G. M. 3D-printer visualization of neuron models. Frontiers in neuroinformatics 9, 18 (2015).

73. Erleben, K., Sporring, J., Henriksen, K. & Dohlmann, H. Physics-based animation (Charles River Media Hingham, 2005).

74. Ramaswamy, S. et al. The neocortical microcircuit collaboration portal: a resource for rat somatosensory cortex. Frontiers in neural circuits 9 (2015).

75. Schmid, F., Barrett, M. J., Jenny, P. & Weber, B. Vascular density and distribution in neocortex. Neuroimage 197, 792–805 (2019).

76. Mihelic, S. A. et al. Segmentation-Less, Automated, Vascular Vectorization. PLoS computational biology 17, e1009451 (2021).

77. Chung, K. & Deisseroth, K. CLARITY for mapping the nervous system. Nat Meth 10, 508–513 (June 2013).

78. Feng, Y. et al. CLARITY reveals dynamics of ovarian follicular architecture and vasculature in three-dimensions. Scientific Reports 7, 1–13 (2017).

79. Di Giovanna, A. P. et al. Whole-brain vasculature reconstruction at the single capillary level. Scientific reports 8, 1–11 (2018).

80. Miyawaki, T. et al. Visualization and molecular characterization of whole-brain vascular networks with capillary resolution. Nature communications 11, 1–11 (2020).

81. Abdellah, M., Bilgili, A., Eilemann, S., Markram, H. & Schürmann, F. Physically-based rendering of highly scattering ?uorescent solutions using path tracing in Proceedings of the 37th Annual Conference of the European Association for Computer Graphics: Posters (2016), 17–18.

82. Azimipour, M. et al. Extraction of optical properties and prediction of light distribution in rat brain tissue. Journal of Biomedical Optics 19 (2014).

83. Dima, A., Scholz, M. & Obermayer, K. Automatic segmentation and skeletonization of neurons from confocal microscopy images based on the 3-D wavelet transform. IEEE Transactions on image processing 11, 790–801 (2002).

84. He, W. et al. Automated three-dimensional tracing of neurons in confocal and brightfield images. Microscopy and microanalysis 9, 296–310 (2003).

85. Markram, H. The human brain project. Scientific American 306, 50–55 (2012).

86. Foundation, B. Blender - 3D modelling and rendering package Blender Foundation (Blender Institute, Amsterdam, 2018). https://www.blender.org.

87. Cignoni, P., Corsini, M. & Ranzuglia, G. Meshlab: an Open-Source 3D mesh processing system. Ercim news 73, 6 (2008).

88. Folk, M., Heber, G., Koziol, Q., Pourmal, E. & Robinson, D. An overview of the HDF5 technology suite and its applications in Proceedings of the EDBT/ICDT 2011 Workshop on Array Databases (2011), 36–47.

89. Attene, M. A lightweight approach to repairing digitized polygon meshes. The visual computer 26, 1393–1406 (2010).

90. Osborn, W. Exploring bit arrays for join processing in spatial data streams in International Conference on Network-Based Information Systems (2019), 73–85.

91. Schwarz, M. & Seidel, H.-P. Fast parallel surface and solid voxelization on GPUs in ACM Transactions on Graphics (TOG) 29 (2010), 179.

92. Hasselgren, J., Akenine-Möller, T. & Ohlsson, L. Conservative rasterization. GPU Gems 2, 677–690 (2005).

93. Shin, H., Park, J. C., Choi, B. K., Chung, Y. C. & Rhee, S. Efficient topology construction from triangle soup in Geometric Modeling and Processing, 2004. Proceedings (2004), 359–364.

94. Akenine-Möllser, T. Fast 3D triangle-box overlap testing. Journal of graphics tools 6, 29–33 (2001).

95. Du, C.-J., Hawkins, P. T., Stephens, L. R. & Bretschneider, T. 3D time series analysis of cell shape using Laplacian approaches. BMC bioinformatics 14, 1–16 (2013).

96. Lorensen, W. E. & Cline, H. E. Marching cubes: A high resolution 3D surface construction algorithm. ACM siggraph computer graphics 21, 163–169 (1987).

97. Nielson, G. M. Dual marching cubes in IEEE Visualization 2004 (2004), 489–496.

98. Ramaswamy, S. & Markram, H. Anatomy and physiology of the thick-tufted layer 5 pyramidal neuron. Frontiers in cellular neuroscience 9, 233 (2015).

99. Yu, Z., Holst, M. J., Cheng, Y. & McCammon, J. A. Feature-preserving adaptive mesh generation for molecular shape modeling and simulation. Journal of Molecular Graphics and Modelling 26, 1370–1380 (2008).

100. Shapson-Coe, A. et al. A connectomic study of a petascale fragment of human cerebral cortex. bioRxiv (2021).

101. Al-Absi, A.-R., Christensen, H. S., Sanchez, C. & Nyengaard, J. R. Evaluation of semi-automatic 3D reconstruction for studying geometry of dendritic spines. Journal of Chemical Neuroanatomy 94, 119–124. ISSN: 0891-0618 (2018).

102. Chandra, R. Parallel programming in OpenMP ISBN: 9780262533027 (Morgan kaufmann, 2001).

