## Supplementary Document for "Ultraliser: a framework for creating multiscale, high-fidelity and geometrically realistic 3D models for *in silico* neuroscience"

#### Supplementary Material

Marwan Abdellah[1], Juan José García Cantero[1], Nadir Román Guerrero[1], Alessandro Foni[1], Jay S. Coggan[1], Corrado Calì[2,4], Marco Agus[3,5], Eleftherios Zisis[1], Daniel Keller[1], Markus Hadwiger[3], Pierre J. Magistretti[2], Henry Markram[1] and Felix Schürmann[1*]

[1]Blue Brain Project
École Polytecnique Fédérale de Lausanne (EPFL)
Geneva, Switzerland

[2]Biological and Environmental Sciences and Engineering Division
King Abdullah University of Science and Technology (KAUST)
Thuwal, Saudi Arabia

[3]Visual Computing Center
King Abdullah University of Science and Technology (KAUST)
Thuwal, Saudi Arabia

[4]Neuroscience Institute Cavalieri Ottolenghi (NICO)
Department of Neuroscience, University of Torino
Orbassano, Italy

[5]College of Science and Engineering
Hamad Bin Khalifa University (HBKU)
Doha, Qatar

2022

112 Pages, 89 Figures

**8 Spine geometries extracted from cortical EM volumes** 94

**9 Generating biologically realistic neuronal meshes from digitized morphologies** 95

**10 Generating continuous cellular meshes from fragmented components** 97

**11 Generating astroglial meshes from complete synthetic morphologies** 98

**12 Generating vasculature meshes from corresponding graph networks** 99

**13 Comparative analysis with existing frameworks and applications** 102

### 1 Digital representations and formats of neuroscientific 3D models

Neuroscientific spatial models (100 nm - 1 mm) can be represented by several structural formats, mainly: morphological skeletons, polygonal surface meshes, tetra- and hexa-hedral volume meshes and voxel grids. Figure. S1 illustrates each representation and the processes used to convert between the different representations.

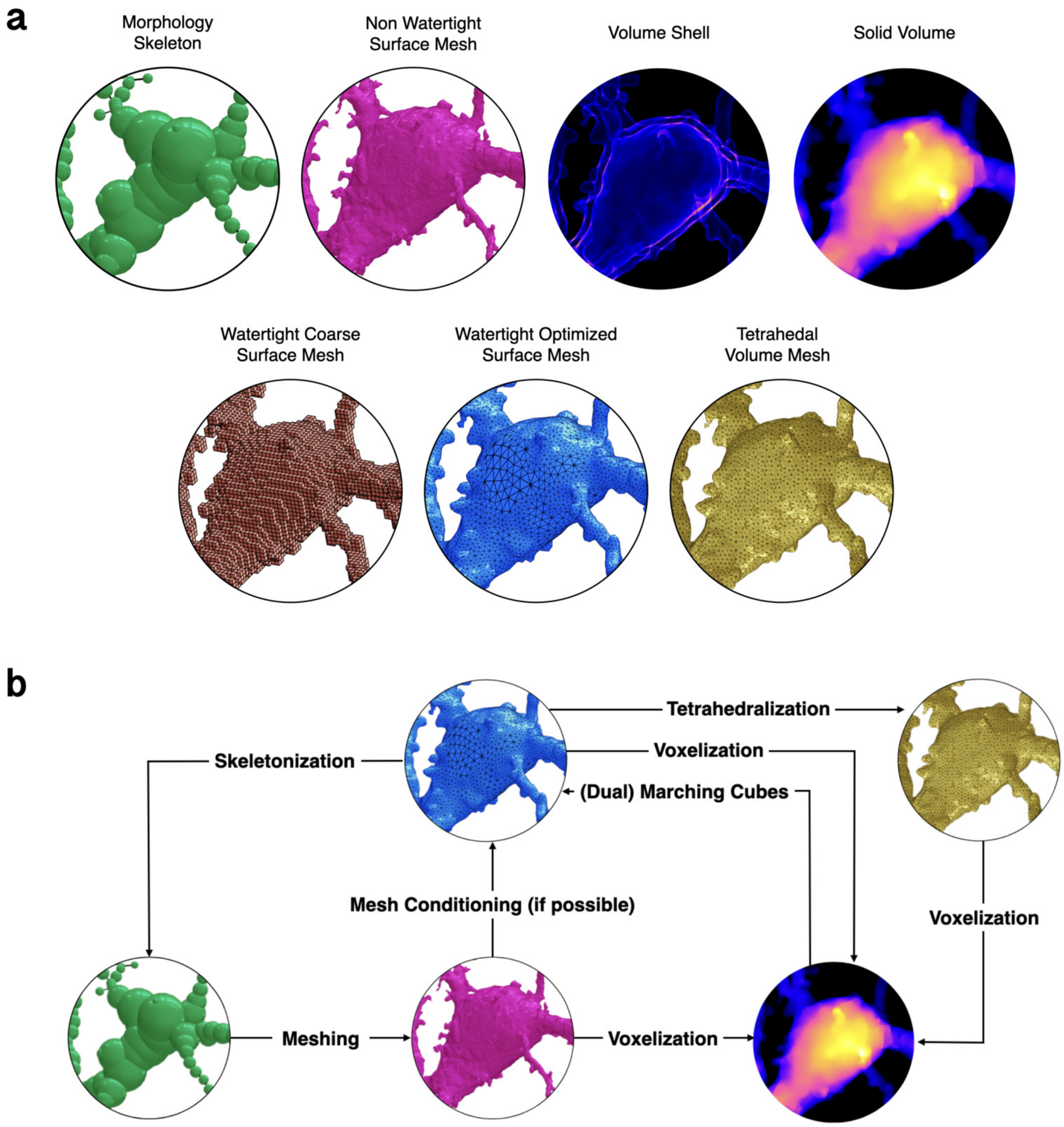

Figure S1: **Summary of the different representations and formats of neuroscientific spatial data (100 nm - 1 mm).**
**(a)** The structure of neuroscientific 3D models can be digitally represented by several formats: morphological skeletons (point-and-diameter representation), polygonal surface meshes, tetrahedral volumetric meshes and volumes (composed of voxels) sampled on 3D Cartesian grids. **(b)** Watertight meshes are required to convert a model to other formats, in which we can perform several kinds of simulations including compartmental, particle, reaction-diffusion and optical imaging simulations. A spiny pyramidal neuron is used for demonstration purposes only, but the idea applies to other cellular and subcellular structures at various spatial scales.

#### 2 Ultraliser workflow

Ultraliser is an unconditionally robust and optimized framework dedicated primarily to *in silico* neuroscience research, allowing to generate high fidelity and multiscale (from subcellular and up to multicellular scales of resolution: 100 nm - 1 mm) 3D neuroscientific models — such as: nuclei, mitochondria, endoplasmic reticula, neurons, astrocytes, pericytes, neuronal branches with dendritic spines, minicolumns with thousands of neurons and large networks of cerebral vasculature — with realistic geometries. Ultraliser implements an effective voxelization-based remeshing engine that can rasterize non-watertight surface meshes —in the form of triangular soups— into high resolution binary volumes (each voxel is represented by a single bit rather than 1 byte in typical voxelization applications), with which we can reconstruct topologically accurate, adaptively optimized and watertight surface manifolds (Figure S1).

In addition to their importance for accurate quantitative analysis, resulting models are primarily intended to automate the process of conducting supercomputer simulations of neuroscience experiments; complementing *in vivo* and *in vitro* techniques. Watertight triangular meshes are used for (i) performing 3D particle simulations, (ii) mesh-based skeletonization, in which accurate morphologies of cellular structures are obtained for performing 1D compartmental simulations and (iii) tetrahedralization, in which we can generate tetrahedral volume meshes for 3D reaction-diffusion simulations. Annotated volumetric tissue models are also used in *in silico* imaging studies, where we can simulate optical imaging experiments with brightfield or fluorescence microscopy[1]. Ultraliser's workflow is graphically illustrated in Figure S2.

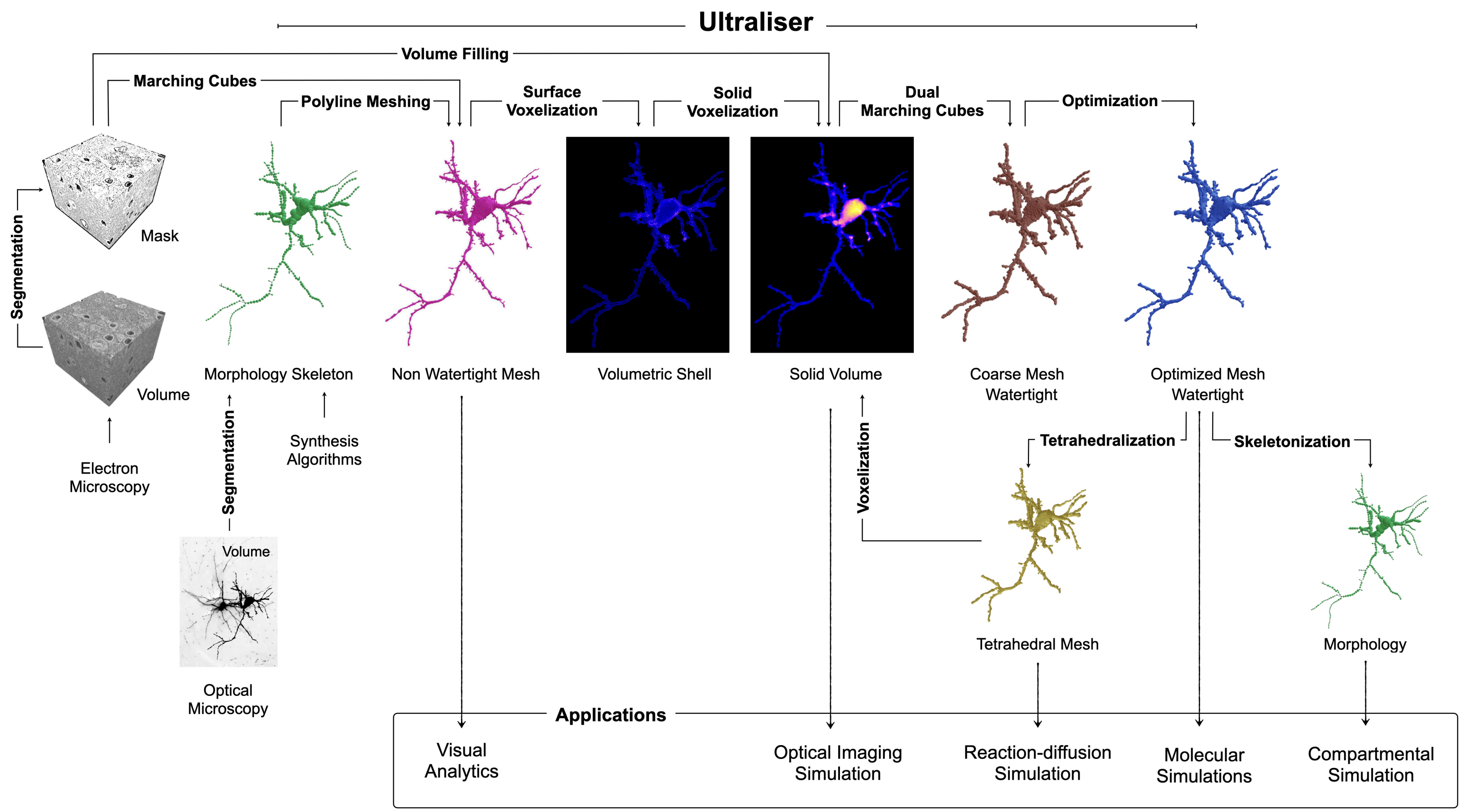

Figure S2: **Ultraliser Workflow.** Related to Fig. 1 and Fig. S1a (Data Formats).
A high-level overview of Ultraliser's workflow showing its subsequent stages (surface and solid voxelization, and mesh reconstruction and optimization) including the different data formats that can be processed within its workflow (masks, gray-scale volumes, polygonal surface meshes, morphological skeletons and tetrahedral volumetric meshes). Ultraliser creates high-fidelity, adaptively optimized and watertight surface manifolds and large-scale annotated volumes. The surface meshes are used directly for molecular simulations, and can be further processed to create volumetric (tetrahedral or hexahedral) meshes for performing reaction-diffusion simulation. They can be also used by mesh-based skeletonization applications for skeletonize accurate morphological counterparts. The annotated volumes are used in *in silico* optical imaging experiments[1].

### 3 Summary and limitations of similar existing frameworks

Ultraliser is a neuroscience-dedicated and *multi-functional* framework; the core is designed to create optimized and topologically accurate surface meshes and annotated volumes, however, its *novelty* comes from the following features:

1. the capability to load multiple structures including (i) non-watertight meshes, (ii) 8-, 16-, 32- and 64-bit volumes, (iii) binary masks, (iv) neuronal, (v) astrocytic and (vi) vascular morphology skeletons, that are principally defined by neuroscience-specific file formats,
2. the scalability to operate on large-scale models,
3. the applicability to create watertight mesh models of neurons, astrocytes, microglia, blood vessels, subcellular structures and cortical networks either from their morphological representations or from previously segmented non-watertight meshes,
4. the integration in a single unifying framework that is dedicated to *in silico* neuroscience research.

The literature does not have similar existing frameworks combining the same functional aspects in a single framework. But, there are other frameworks and standalone applications that can handle individual use cases.

#### 3.1 Remeshing of non-watertight meshes

A few relevant *re-meshing* frameworks – that are open source – are capable of handling geometric topology and optimization issues for relatively small scale structures (minuscule segments of spiny dendrites), such as GAMer[2] and VolRoverN[3], but they are incapable of accomplishing watertightness, unable to process highly tessellated meshes and even can destruct the topology if input meshes are merely triangular soups. Other generic re-meshing frameworks such as ManifoldPlus[4] can create watertight meshes for some CAD/CAM models, but it fails to preserve the spatial structure and topology of complex structures such as neuronal or astroglial meshes. TriMesh has recently integrated a mesh repair module, but when tested, it could not resolve the artifacts of a neuronal triangular soup and the resulting mesh was not watertight.

#### 3.2 Meshing of neuronal and astroglial morphologies

There are several implementations for creating neuronal mesh models from their corresponding morphological skeletons (point-and-diameter representations, Figure S76). In summary, these implementations can be classified into two categories. The first one focuses on creating visually appealing and low-tessellated meshes for visual analysis[5] and rendering[6] applications; this class does not address watertightness. The second category includes a set of application that try to guarantee the watertightness of the resulting meshes, but the geometric realism of the generated models can be questionable. Table S1 provides a summary of these various meshing applications, their availability and features.

Table S1: A list of open source neuronal and astrocytic mesh reconstruction frameworks and applications

| Framework or Method | Soma | Branching Geometry | Watertightness |
|---|---|---|---|
| AnaMorph[7] | Spheres | Not accurate | Possible |
| CTNG[8] | 2D Profile-based | Not accurate | Possible |
| Neuronize[9,10] | Mass-spring | Not accurate | Not watertight |
| NeuroTessMesh[11] | FEM | Intersections exist | Not watertight |
| NeuroMorphoVis[12] | Mass-Spring | Overlapping | Not watertight |
| Metaball Skinning[13] | Implicit Geometry | Organic and accurate | Not guaranteed |
| Union-operators Skinning[14] | Implicit Geometry | Organic but not accurate | Not Watertight |

##### 3.3 Meshing of vascular morphologies

Table S2 summarizes a list of methods for meshing vascular morphologies and their features.

Table S2: A list of open source vascular mesh reconstruction frameworks

| Framework | Method | Branching | Watertightness |
|---|---|---|---|
| VessMorphoVis[15] | Implicit geometry (Metaballs) | Organic and accurate | Possible |
| VessMorphoVis[15] | Intersecting polylines | Not accurate | Not watertight |

### 4 Mesh quality and analysis metrics

#### 4.1 Watertightness

A watertight mesh is a manifold that consists of one closed surface, i.e. it does not contain any gaps or holes and have a clearly defined boundary and inside. This criteria is essential in CAD/CAM design and in the context of computational modeling. By definition, a mesh is watertight if the following conditions are met: (i) it has no self-intersecting faces, (ii) it is two-manifold, i.e. does not contain any non-manifold edges and non-manifold vertices, and (iii) has no boundary edges. A self-intersection is an intersection of two facets belonging to the same mesh. A non manifold edge is an edge that has more than two incident faces. To understand what a non-manifold vertex is, we define the *star of a vertex* to be the union of all its incident faces. A non-manifold vertex is a vertex where the corresponding star is not any further connected after the removal of the vertex. A two-manifold mesh is a mesh that has zero non-manifold edges and non-manifold vertices. A watertight mesh is then a two-manifold mesh that has no self-intersecting faces and zero boundary edges[16].

#### 4.2 Triangular mesh quality metrics

For a given triangular mesh $\mathcal{M}$, and for a given triangle $T$ that has three Cartesian points $P_0$, $P_1$ and $P_2$, the edge vectors of T can be named by the vertex opposite the edge such that the edges $E_0$, $E_1$ and $E_2$ can be constructed as follows:

$$\begin{aligned} \vec{E_0} &= \vec{P_1} - \vec{P_0} \\ \vec{E_1} &= \vec{P_2} - \vec{P_1} \\ \vec{E_2} &= \vec{P_0} - \vec{P_2} \end{aligned} \tag{1}$$

The triangle edges lengths are denoted $L_{E_0}$, $L_{E_1}$ and $L_{E_2}$ and defined as follows:

$$L_{E_0} = \mid \vec{P_0} \mid \qquad L_{E_1} = \mid \vec{P_1} \mid \qquad E_2 = \mid \vec{P_2} \mid \tag{2}$$

Meanwhile, the smallest and largest lengths of the three edges are defined as follows:

$$L_{\min} = \min(L_{E_0}, L_{E_1}, L_{E_2}) \qquad L_{\max} = \max(L_{E_0}, L_{E_1}, L_{E_2}) \tag{3}$$

The area $A$, inner radius $r$ and circumradius $R$ of $T$ can therefore be computed according to Equations 4, 5 and 6 respectively:

$$A = \frac{1}{2} \mid \vec{E_0} \times \vec{E_1} \mid = \frac{1}{2} \mid \vec{E_1} \times \vec{E_2} \mid = \frac{1}{2} \mid \vec{E_2} \times \vec{E_0} \mid \tag{4}$$

$$r = \frac{2A}{\mid \vec{E_0} \mid + \mid \vec{E_1} \mid + \mid \vec{E_2} \mid} \tag{5}$$

$$R = \frac{| \vec{E_0} || \vec{E_1} || \vec{E_2} |}{2r(| \vec{E_0} | + | \vec{E_1} | + | \vec{E_2} |)} \tag{6}$$

*Mesh area* The surface area of the mesh (composed of $N$ triangles) can be computed as follows:

$$A_{\mathcal{M}} = \sum_{i=0}^{i=N-1} A_i \tag{7}$$

*Mesh volume* The volume of the surface mesh can be computed as follows, where $n$ is the normal on the triangle:

$$V_{\mathcal{M}} = \frac{1}{6} \sum_{i=0}^{i=N-1} A_i \cdot n_i \tag{8}$$

*Dihedral angles* The minimum $\theta_{\text{min}}$ and maximum $\theta_{\text{max}}$ dihedral angles are computed according to Equations 9 and 10 respectively. The acceptable ranges for $\theta_{\text{min}}$ and $\theta_{\text{max}}$ are $[\,30^\circ, 60^\circ\,]$ and $[\,60^\circ, 90^\circ\,]$ respectively.

$$\theta_{\text{min}} = \min_{n \in \{0,1,2\}} \left\{ \arccos\left( \frac{\vec{E_n} \cdot \vec{E_{n+1}}}{| \vec{E_n} || \vec{E_{n+1}} |} \right) \left( \frac{180^\circ}{\pi} \right) \right\} \tag{9}$$

$$\theta_{\text{max}} = \max_{n \in \{0,1,2\}} \left\{ \arccos\left( \frac{\vec{E_n} \cdot \vec{E_{n+1}}}{| \vec{E_n} || \vec{E_{n+1}} |} \right) \left( \frac{180^\circ}{\pi} \right) \right\} \tag{10}$$

*Edge ratio* The edge ratio of the triangle is computed according to Equation 11. The acceptable range of $\mathcal{E}_{\mathcal{M}}$ is $[\,0, 1.0\,]$.

$$\mathcal{E}_{\mathcal{M}} = \frac{L_{\text{max}}}{L_{min}} \tag{11}$$

*Radius ratio* The radius ratio is computed according to Equation 12. The acceptable range of $\mathcal{E}_{\mathcal{M}}$ is $[\,0, 1.0\,]$.

$$\mathcal{R} = \frac{2r}{R} \tag{12}$$

*Radius to edge ratio* The radius to edge ratio is computed according to Equation 13. The acceptable range of $\mathcal{E}_{\mathcal{M}}$ is $[\,0, 1.3\,]$.

$$\mathcal{R}\mathcal{E} = \frac{\mathcal{R}}{\mathcal{E}} \tag{13}$$

##### 4.3 Hausdorff distance

Hausdorff distance (**Eq. 14**) measures the error between 3D discrete surfaces represented by triangular patches.[17]. We will use the Hausdorff distance metric to determine how close the resulting meshes are compared to the input ones.

*Definition* Let $X$ and $Y$ be two non-empty subsets of a metric space (i.e. 3D triangular meshes) $(\mathcal{M}_1, \mathcal{M}_2)$. We define the Hausdorff distance between the two subsets $d_{\mathrm{H}}(X, Y)$ by

$$d_{\mathrm{H}}(X,Y) = \max\left\{ \sup_{x \in X} d(x,Y), \sup_{y \in Y} d(X,y) \right\} \tag{14}$$

where $sup$ represents the *supremum*, $inf$ the *infimum*, and where $d(a,B) = \inf_{b \in B} d(a,b)$ quantifies the distance from a point $a \in X$ to the subset $B \subseteq X$.

##### 4.4 Mesh volume

In computational biology, volume preservation of structural models is essential to guarantee the results of simulations. During the remeshing procedure, the volume of the resulting mesh can significantly change with respect to the actual volume of the input mesh. This depends on multiple parameters, mainly the voxelization resolution and number of smoothing iterations. We therefore measure and compare the volumes of the resulting meshes from Ultraliser with respect to the volumes of input meshes, morphologies or volumes, with which we can validate the results of the simulations. The volume of a given mesh (must be watertight) can be computed according to Equation 8.

### 5 Analysis of segmented models from the NGV ensemble

The tissue block, shown in Figure S3 (101×101×75 cubic microns), is acquired from layer VI of the somatosensory cortex of a two-weeks-old rat (P14) using serial block-face scanning electron microscopy at 20 nm resolution[18]. The acquired volume stack contains 1513 images, where each image has 4096×4096 pixels. The stack is segmented with a hybrid pipeline that combined TrakEM2[19,20] and iLastik[21] (ilastik.org) for manual and automated segmentation respectively. Further details on the imaging procedure, segmentation pipeline and the morphological analysis of the reconstructed morphologies are available[22].

A total of 186 cells were visually identified and classified into 124 neurons, 22 astrocytes, 17 microglia, 11 pericytes, 6 endothelium and 6 other unknown structures. Nevertheless, the complete, or *almost* complete, reconstructions were limited to four astrocytes, four neurons, four microglia, four pericytes and an oligodendrocyte including their mitochondria and nuclei. We also segmented two blood vessels, 213 myelinated axons and a group of endoplasmic reticula of one astrocyte. The segmented mesh models are initially stored in a single Blender file in which we can verify their spatial relationship in comparison to the acquire volume stack. Afterwards, every individual mesh is exported to a separate Wavefreont object file for subsequent mesh analysis and visual analytics. Quantitative analysis results of cellular (astrocytes, neurons, microglia, pericytes and oligodendrocytes) and subcellular (mitochondria and endoplasmic reticula) meshes are detailed in Tables S3 and S4.

Except for the pericytes that relatively have very simple surfaces, the analysis demonstrates that none of the segmented cellular meshes is watertight, and most of cells are fragmented into multiple partitions, with thousands of self-intersecting facets, non-manifold edges, non-manifold vertices and boundary edges. A summary of the small and incomplete structures including blood vessels, RBCs and myelinated axons is shown in Table S5. The analysis shows also a significant amount of non watertight meshes. Nevertheless, they were supposed to be used for reaction-diffusion simulations for understanding the kinetics. We therefore used ultraMesh2Mesh to reconstruct watertight counterparts that can be tetrahedralized and employed in STEPS simulations. Note that all the cellular and subcellular meshes have been resulting with voxelization resolutions of 5 and 10 voxels per micron respectively. Reference to a summary of the visual, quantitative and qualitative analytical comparisons between original and resulting meshes is listed in Tables S6 and S7.

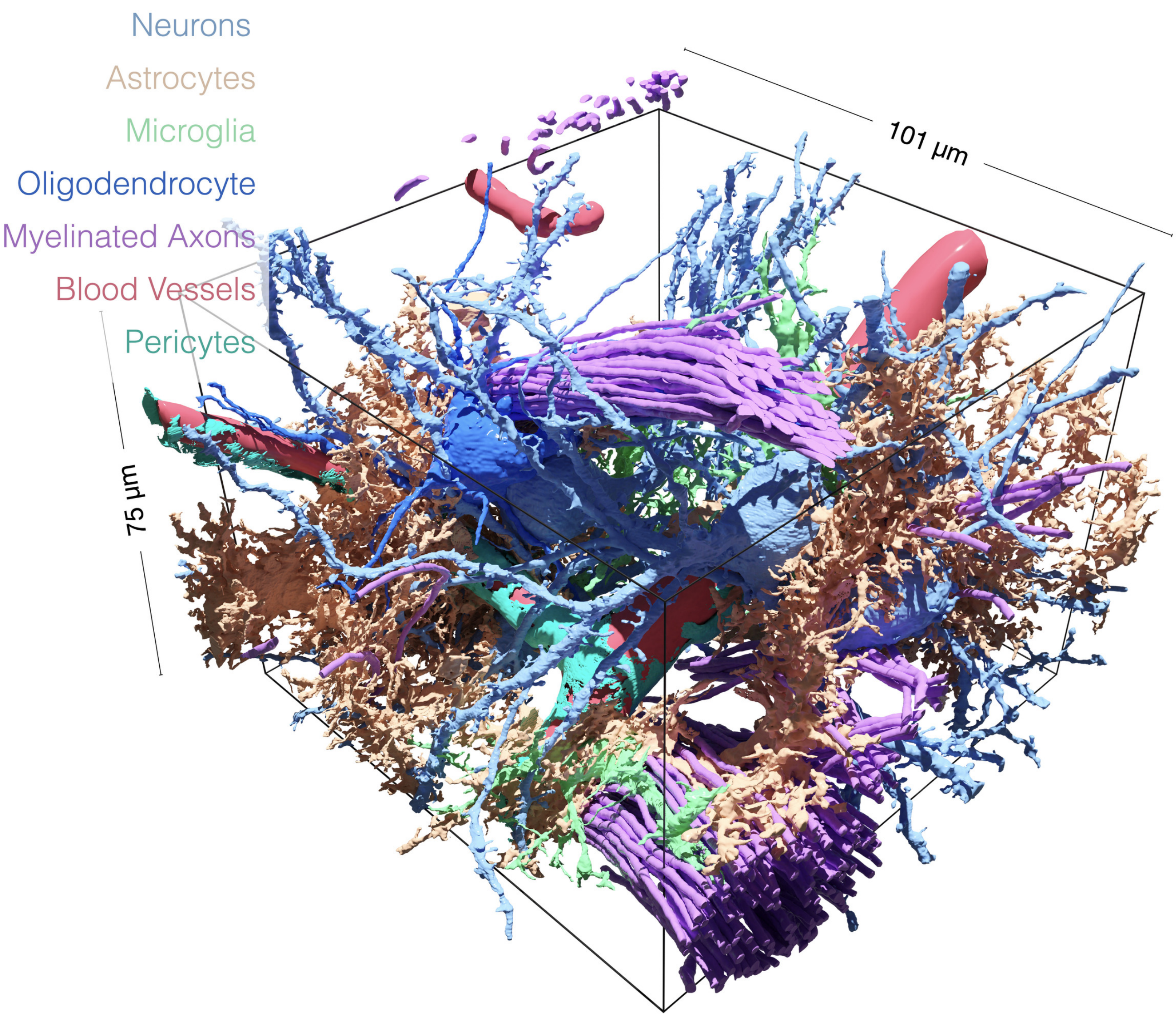

Figure S3: **Segmented NGV Structures**, Magnification of Fig. 2a.
3D reconstruction of NGV structures from Layer IV somatosensory cortex brain parenchyma of a juvenile rat (101 × 101 × 75 $\mu m^3$). Neurons are in light blue, astrocytes are in light orange, microglia are in light green, pericytes are in turquoise, oligodendrocyte is in darkblue, myelinated axons are in violet, and blood vessels are in red. Intracellular structures such as RBCs, nuclei, mitochondria and endoplasmic reticula are not shown. Morphometric analysis of all the segmented structures is detailed in a previous study[22].

Table S3: Quantitative analysis of the cellular meshes segmented from the neuropil volume shown in Figure S3.

| Mesh | Polygons | Vertices | Intersections[1] | NME [2] | NMV[3] | Partitions[4] | Watertight[5] |
|---|---|---|---|---|---|---|---|
| Astrocyte 1 | 2,048,700 | 1,036,667 | 2 | 4,774 | 9,353 | 20 | **No** |
| Astrocyte 2 | 2,116,720 | 1,040,410 | 47 | 13,967 | 25,558 | 89 | **No** |
| Astrocyte 3 | 1,373,888 | 676,467 | 15 | 10,257 | 19,418 | 116 | **No** |
| Astrocyte 4 | 974,760 | 481,922 | 6 | 3,760 | 7,779 | 14 | **No** |
| Neuron 1 | 308,988 | 153,902 | 5 | 579 | 1,116 | 15 | **No** |
| Neuron 2 | 1,194,532 | 595,631 | 2 | 1,601 | 3,058 | 16 | **No** |
| Neuron 3 | 1,747,684 | 872,226 | 0 | 1,411 | 2,686 | 1 | **No** |
| Neuron 4 | 1,303,396 | 650,459 | 0 | 1,090 | 2,128 | 1 | **No** |
| Microglia 1 | 245,240 | 121,384 | 0 | 996 | 1,819 | 12 | **No** |
| Microglia 2 | 345,620 | 172,421 | 0 | 349 | 677 | 1 | **No** |
| Microglia 3 | 657,220 | 327,623 | 0 | 881 | 1,692 | 7 | **No** |
| Microglia 4 | 93,624 | 46,687 | 3 | 222 | 436 | 22 | **No** |
| Pericyte 1 | 20,624 | 10,314 | 0 | 0 | 0 | 1 | Yes |
| Pericyte 2 | 11,908 | 5,956 | 0 | 0 | 0 | 1 | Yes |
| Pericyte 3 | 14,056 | 7,030 | 0 | 0 | 0 | 1 | Yes |
| Pericyte 4 | 10,465 | 5,283 | 0 | 1 | 2 | 1 | **No** |
| Oligodendrocyte 1 | 272,056 | 135,129 | 0 | 873 | 1,722 | 6 | **No** |

1 Intersections in this context indicate self intersecting faces in the mesh even if the mesh contains more than a single partition.
2 Non manifold edges, where an edge in a surface mesh is defined to be non-manifold if it has more than two incident faces.
3 Non-manifold vertices, where a vertex in a polygonal surface mesh is defined to be non-manifold if its corresponding star is not connected when removing the vertex.
4 A mesh partition in a polygonal surface mesh is defined to be an independent set of vertices, edges and faces that clearly define a partial surface manifold of the entire mesh. A single mesh could consist of multiple mesh partitions or islands.
5 A polygonal surface mesh is defined to be watertight if has no self intersections and is two-manifold, i.e. if it does contain neither non-manifold edges nor non-manifold vertices.

Table S4: Quantitative analysis of the subcellular meshes segmented from the neuropil volume shown in Figure S3. MT and ER stand for Mitochondria and Endoplasmic Reticulum respectively.

| Mesh | Polygons | Vertices | Intersections | NME | NMV | Partitions | Watertight |
|---|---|---|---|---|---|---|---|
| Astrocyte 1 MT | 391,104 | 194,756 | 99 | 1,406 | 1,684 | 127 | **No** |
| Astrocyte 2 MT | 322,320 | 160,435 | 142 | 1,516 | 220 | 220 | **No** |
| Astrocyte 3 MT | 953,721 | 473,987 | 195 | 3,734 | 5,111 | 292 | **No** |
| Astrocyte 4 MT | 143,656 | 71,850 | 261 | 1,119 | 1,003 | 32 | **No** |
| Neuron 1 MT | 523,480 | 261,966 | 353 | 3,037 | 3,168 | 817 | **No** |
| Neuron 2 MT | 474,360 | 236,173 | 183 | 2,027 | 2,501 | 223 | **No** |
| Neuron 3 MT | 686,164 | 342,159 | 612 | 4,021 | 3,508 | 336 | **No** |
| Neuron 4 MT | 243,422 | 121,691 | 0 | 978 | 870 | 156 | **No** |
| Microglia 1 MT | 14,508 | 7,413 | 0 | 384 | 103 | 38 | **No** |
| Microglia 2 MT | 50,144 | 25,115 | 0 | 577 | 432 | 102 | **No** |
| Microglia 3 MT | 169,112 | 84,243 | 0 | 356 | 641 | 78 | **No** |
| Microglia 4 MT | 142,896 | 71,199 | 191 | 1,337 | 1,016 | 94 | **No** |
| Pericyte 1 MT | 52,260 | 26,080 | 121 | 624 | 530 | 81 | **No** |
| Pericyte 2 MT | 180,060 | 91,044 | 283 | 1,245 | 469 | 409 | **No** |
| Pericyte 3 MT | 180,564 | 89,923 | 55 | 685 | 898 | 103 | **No** |
| Pericyte 4 MT | 81,792 | 40,898 | 51 | 404 | 347 | 72 | **No** |
| Astrocyte 1 ER | 293,052 | 146,530 | 0 | 0 | 0 | 472 | Yes |
| Astrocyte 2 ER | 326,492 | 170,500 | 0 | 0 | 0 | 4,128 | Yes |
| Astrocyte 3 ER | 140,972 | 73,380 | 0 | 0 | 0 | 1,960 | Yes |
| Astrocyte 4 ER | 33,040 | 17,552 | 0 | 0 | 0 | 654 | Yes |

Table S5: Summary of the number of non-watertight meshes of small or incomplete structures in the neuropil volume, Figure S3.

| Meshes | Total Count | Number of Non-Watertight Meshes |
|---|---|---|
| Blood Vessels | 2 | 1 |
| Myelinated Axons | 213 | 174 |
| RBCs | 33 | 26 |

Table S6: References to the figures showing full comparative analysis between input and ultralized meshes of cellular structures listed in Table S3 and rendered in Figure S3.

| Neuropil Structure | Wireframe Visualization | Qualitative and Quantitative Analysis |
|---|---|---|
| Astrocyte 1 | Fig. S5 | Fig. S9 (a) |
| Astrocyte 2 | Fig. S6 | Fig. S9 (b) |
| Astrocyte 3 | Fig. S7 | Fig. S9 (c) |
| Astrocyte 4 | Fig. S8 | Fig. S9 (d) |
| Neuron 1 | Fig. S10 | Fig. S14 (a) |
| Neuron 2 | Fig. S11 | Fig. S14 (b) |
| Neuron 3 | Fig. S12 | Fig. S14 (c) |
| Neuron 4 | Fig. S13 | Fig. S14 (d) |
| Microglia 1 | Fig. S15 | Fig. S19 (a) |
| Microglia 2 | Fig. S16 | Fig. S19 (b) |
| Microglia 3 | Fig. S17 | Fig. S19 (c) |
| Microglia 4 | Fig. S18 | Fig. S19 (d) |
| Pericyte 1 | Fig. S20 | Fig. S24 (a) |
| Pericyte 2 | Fig. S21 | Fig. S24 (b) |
| Pericyte 3 | Fig. S22 | Fig. S24 (c) |
| Pericyte 4 | Fig. S23 | Fig. S24 (d) |
| Oligodendrocyte | Fig. S25 | Fig. S26 |

Table S7: References to the figures showing full comparative analysis between input and ultralized meshes of subcellular and incomplete cellular structures listed in Table S4 and rendered in Figure S3. MT and ER stand for Mitochondria and Endoplasmic Reticulum.

| Neuropil Structure | Wireframe Visualization | Qualitative and Quantitative Analysis |
|---|---|---|
| Blood Vessels | Fig. S27 | Fig. S28 |
| Astrocyte 1 MT | Fig. S29 | Fig. S33 (a) |
| Astrocyte 2 MT | Fig. S30 | Fig. S33 (b) |
| Astrocyte 3 MT | Fig. S31 | Fig. S33 (c) |
| Astrocyte 4 MT | Fig. S32 | Fig. S33 (d) |
| Neuron 1 MT | Fig. S34 | Fig. S38 (a) |
| Neuron 2 MT | Fig. S35 | Fig. S38 (b) |
| Neuron 3 MT | Fig. S36 | Fig. S38 (c) |
| Neuron 4 MT | Fig. S37 | Fig. S38 (d) |
| Microglia 1 MT | Fig. S39 | Fig. S43 (a) |
| Microglia 2 MT | Fig. S40 | Fig. S43 (b) |
| Microglia 3 MT | Fig. S41 | Fig. S43 (c) |
| Microglia 4 MT | Fig. S42 | Fig. S43 (d) |
| Pericyte 1 MT | Fig. S44 | Fig. S48 (a) |
| Pericyte 2 MT | Fig. S45 | Fig. S48 (b) |
| Pericyte 3 MT | Fig. S46 | Fig. S48 (c) |
| Pericyte 4 MT | Fig. S47 | Fig. S48 (d) |
| Astrocyte 1 ER | Fig. S49 | Fig. S53 (a) |
| Astrocyte 2 ER | Fig. S50 | Fig. S53 (b) |
| Astrocyte 3 ER | Fig. S51 | Fig. S53 (c) |
| Astrocyte 4 ER | Fig. S52 | Fig. S53 (d) |

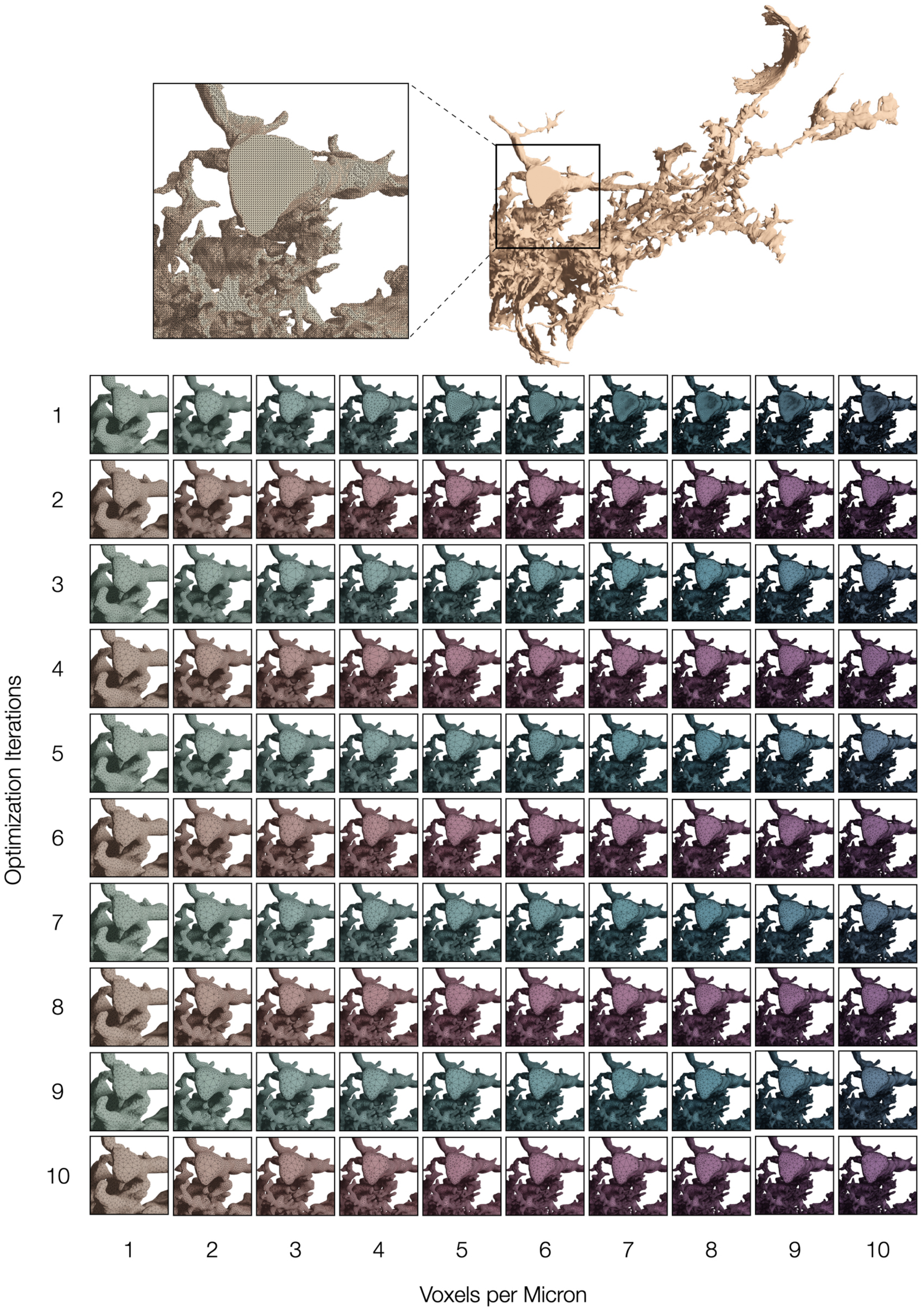

Figure S4: **Visual Analysis Matrix**, Related to Fig. 2d.
Visual mesh analysis that compares the resulting meshes of the same astrocytic 3D model at different voxelization resolutions (in voxels per micron) with respect to the number of optimization iterations that is proportional to the tessellation level of the resulting mesh.

#### 5.1 Astrocytes

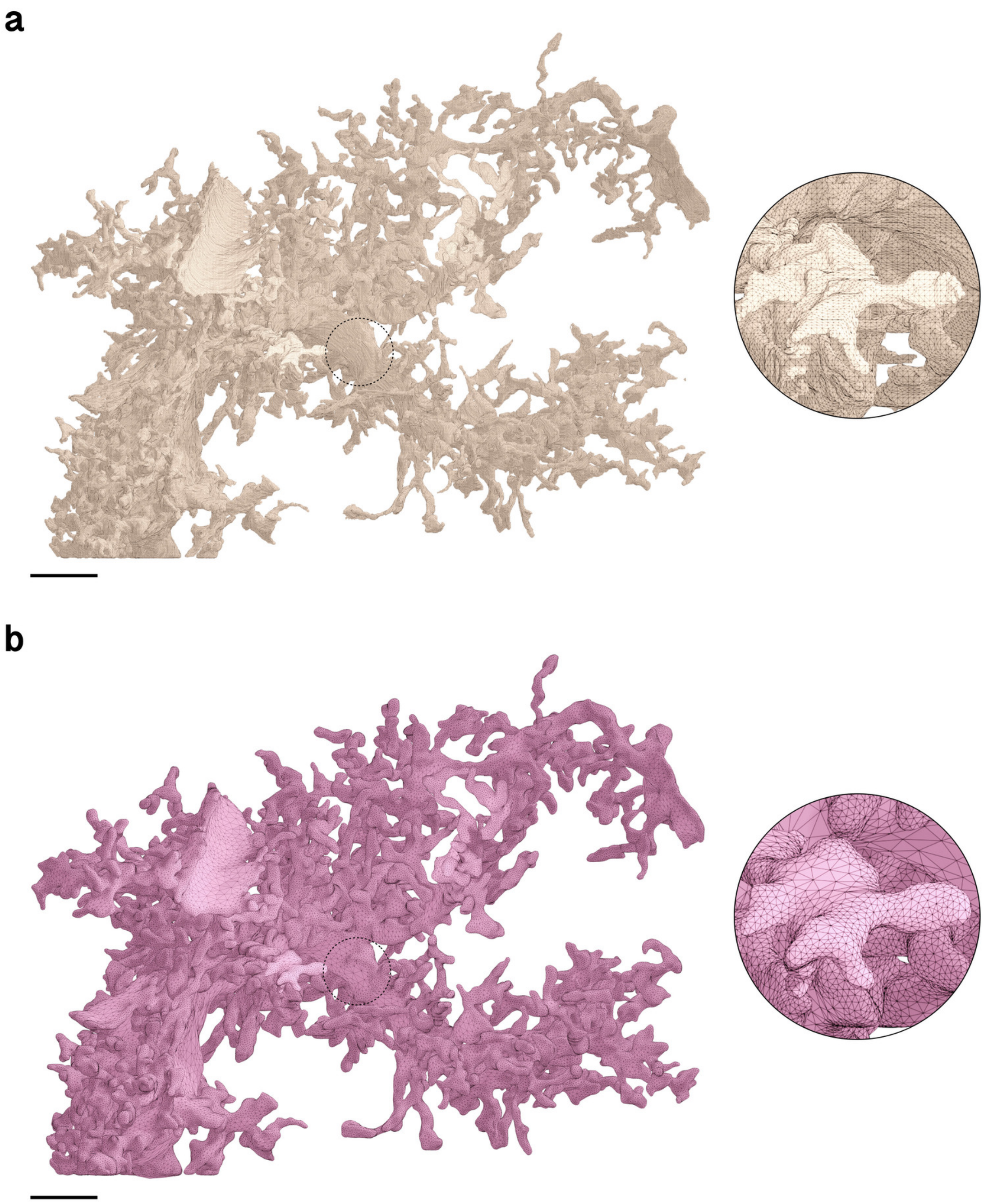

Figure S5: Wireframe visualizations comparing input (**a**) and resulting (**b**) surface meshes of Astrocyte 1. Fragmented partitions and floating vertices are automatically removed to yield a single mesh partition with continuous membrane manifold. The closeups highlight the contrast in surface roughness, topology and tessellation. Comparative quantitative and qualitative analysis is shown in Figure S9a. Scale bars, 5 $\mu$m (**a, b**).

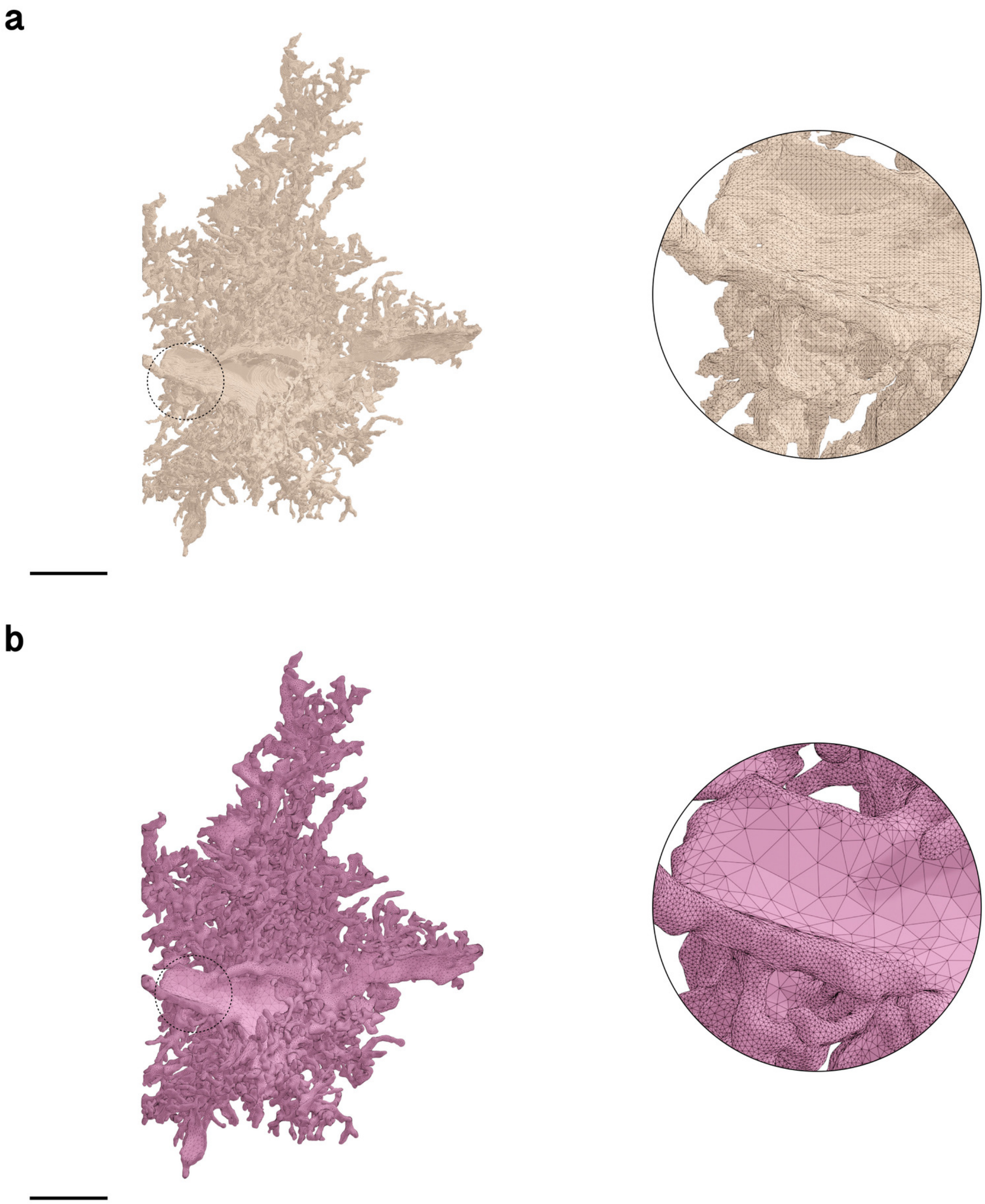

Figure S6: Wireframe visualizations comparing input (**a**) and resulting (**b**) surface meshes of Astrocyte 2. Fragmented partitions and floating vertices are automatically removed to yield a single mesh partition with continuous membrane manifold. The closeups highlight the contrast in surface roughness, topology and tessellation. Comparative quantitative and qualitative analysis is shown in Figure S9b. Scale bars, 10 $\mu$m (**a, b**).

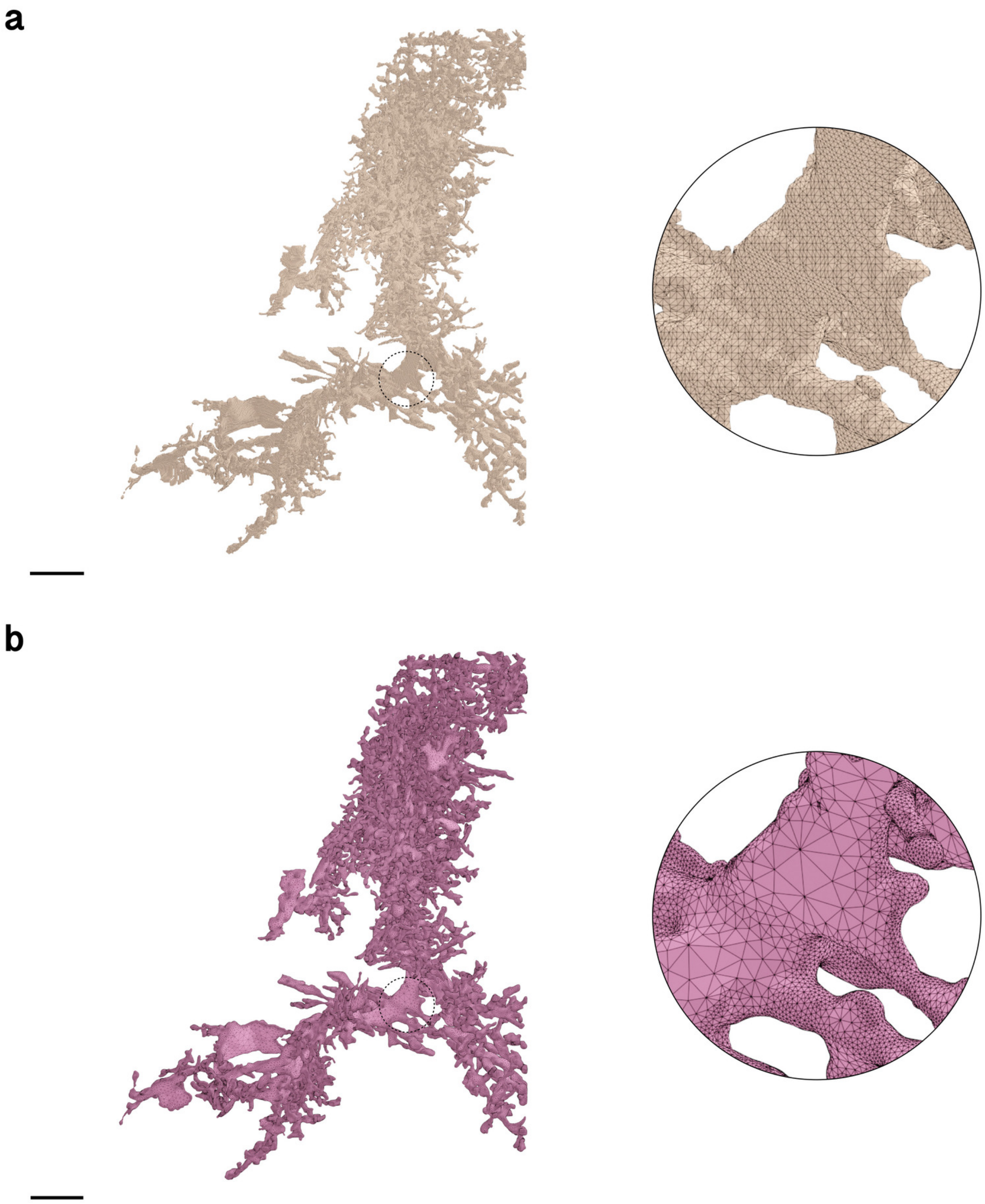

Figure S7: Wireframe visualizations comparing input (**a**) and resulting (**b**) surface meshes of Astrocyte 3. Fragmented partitions and floating vertices are automatically removed to yield a single mesh partition with continuous membrane manifold. The closeups highlight the contrast in surface roughness, topology and tessellation. Comparative quantitative and qualitative analysis is shown in Figure S9c. Scale bars, 10 $\mu$m (**a, b**).

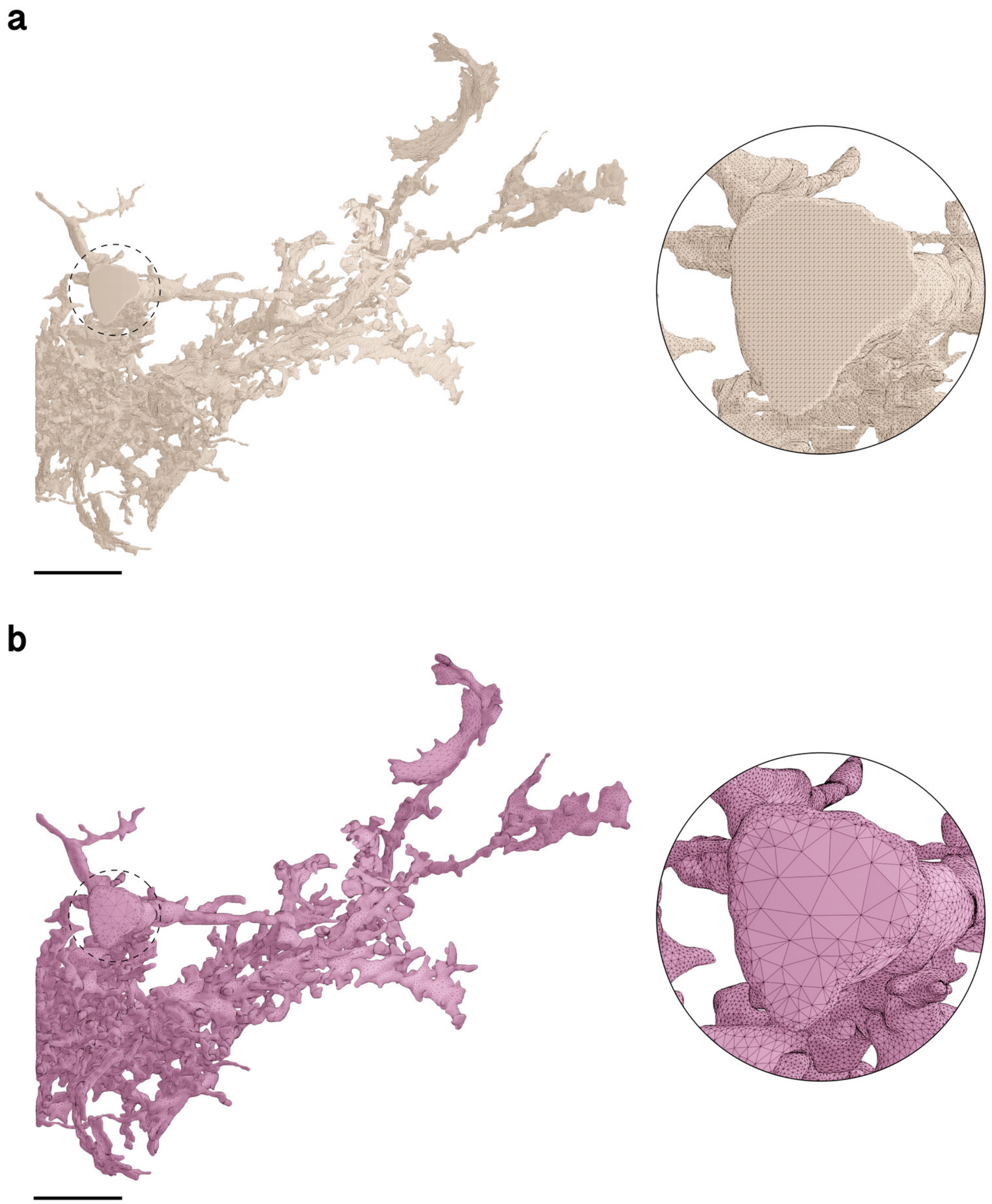

Figure S8: Wireframe visualizations comparing input (**a**) and resulting (**b**) surface meshes of Astrocyte 4. Fragmented partitions and floating vertices are automatically removed to yield a single mesh partition with continuous membrane manifold. The closeups highlight the contrast in surface roughness, topology and tessellation. Comparative quantitative and qualitative analysis is shown in Figure S9d. Scale bars, 10 $\mu$m (**a, b**).

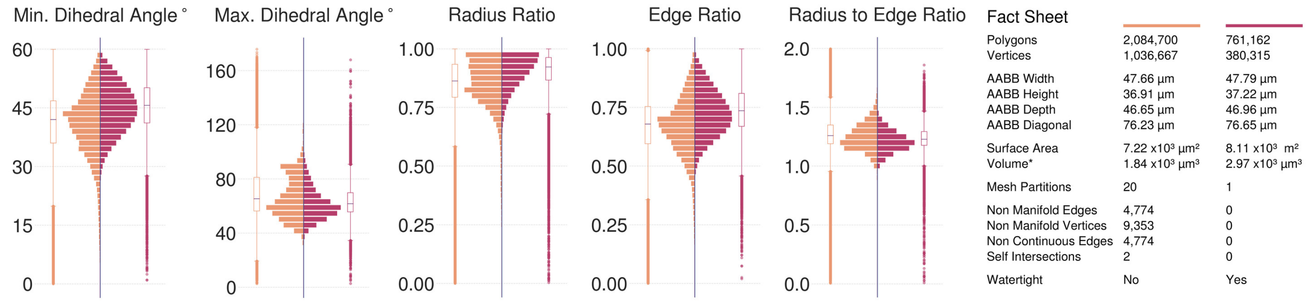

(**a**) Astrocyte 1, Visualization in Figure S5.

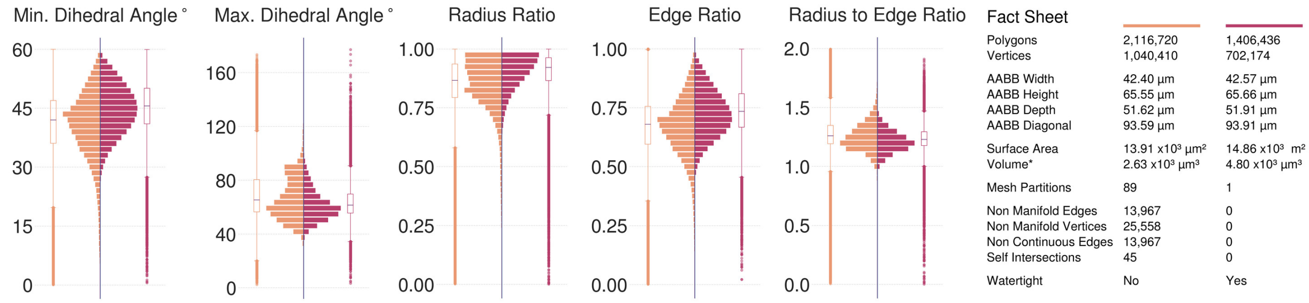

(**b**) Astrocyte 2, Visualization in Figure S6.

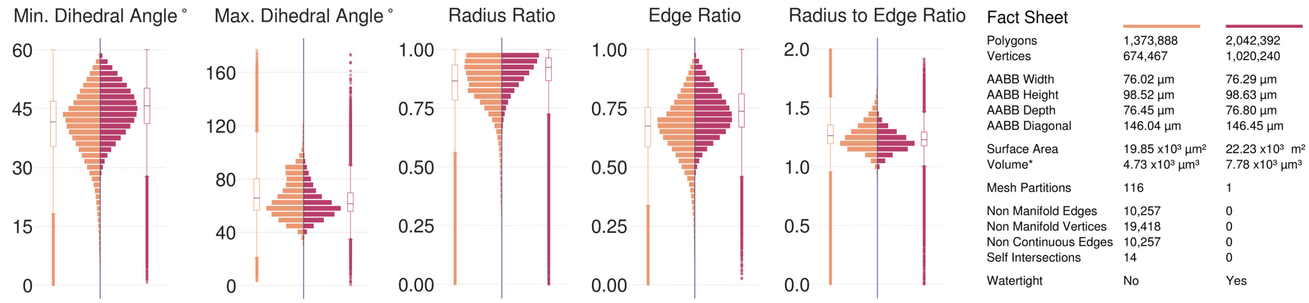

(**c**) Astrocyte 3, Visualization in Figure S7.

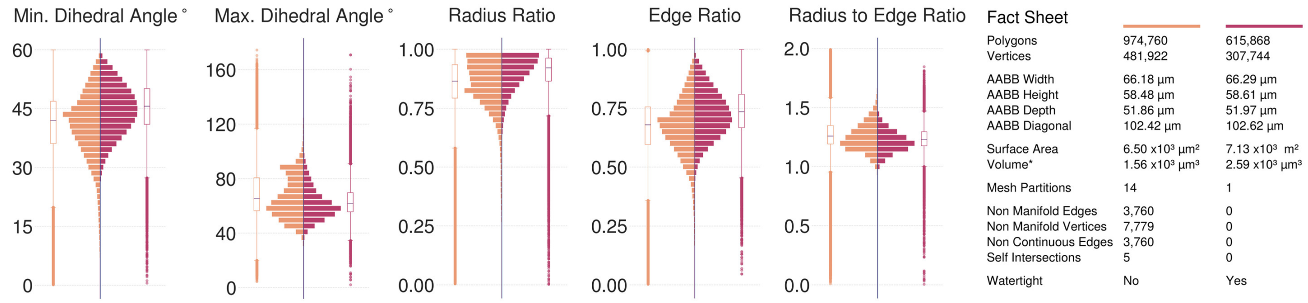

(**d**) Astrocyte 4, Visualization in Figure S8.

Figure S9: Comparative analysis between input (left in orange) and resulting (right in dark salmon) meshes of the complete astrocytes segmented from the volume shown in Figure S3. All the input meshes are not watertight; they have multiple fragmented partitions and severe geometric artifacts. The resulting meshes are reconstructed at 5 voxels per micron resolution (200 nm) yielding single, continuous and adaptively optimized manifolds with no artifacts.

#### 5.2 Neurons

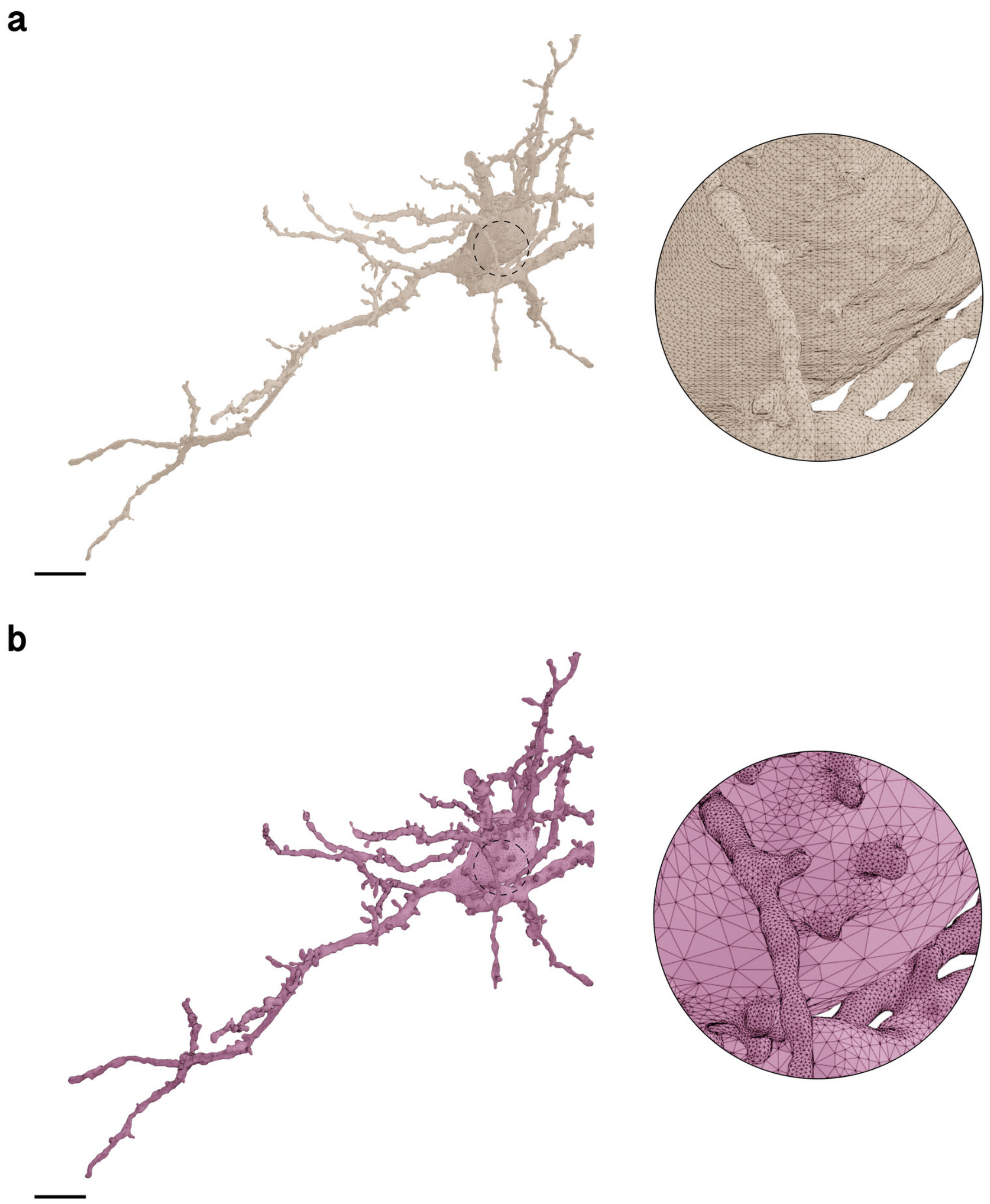

Figure S10: Wireframe visualizations comparing input (**a**) and resulting (**b**) surface meshes of Neuron 1. Fragmented partitions and floating vertices are automatically removed to yield a single mesh partition with continuous membrane surface. The closeups highlight the contrast in surface roughness, topology and tessellation. Comparative quantitative and qualitative analysis is shown in Figure S14a. Scale bars, 5 $\mu$m (**a, b**).

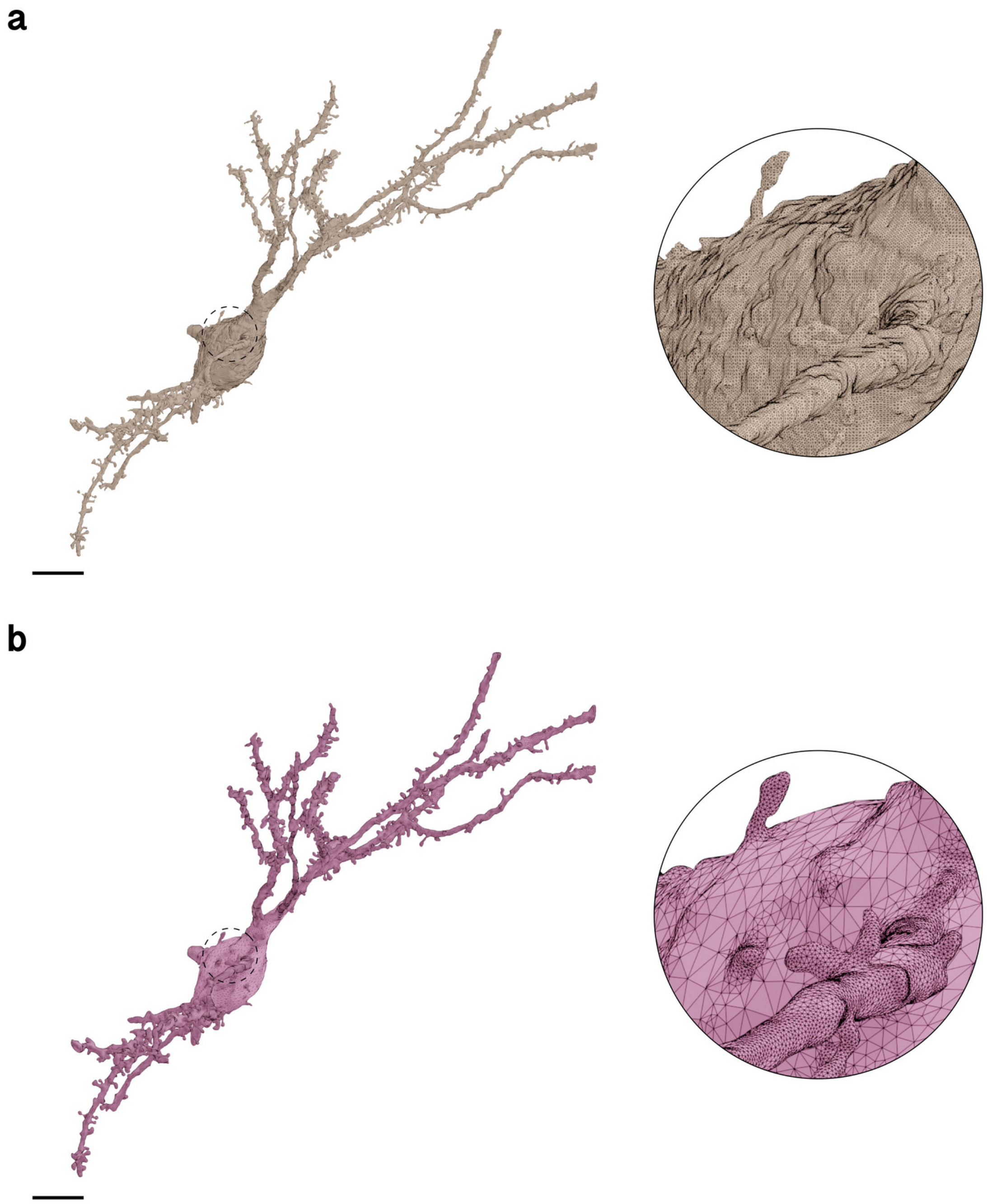

Figure S11: Wireframe visualizations comparing input (**a**) and resulting (**b**) surface meshes of Neuron 2. Fragmented partitions and floating vertices are automatically removed to yield a single mesh partition with continuous membrane surface. The closeups highlight the contrast in surface roughness, topology and tessellation. Comparative quantitative and qualitative analysis is shown in Figure S14b. Scale bars, 5 $\mu$m (**a, b**).

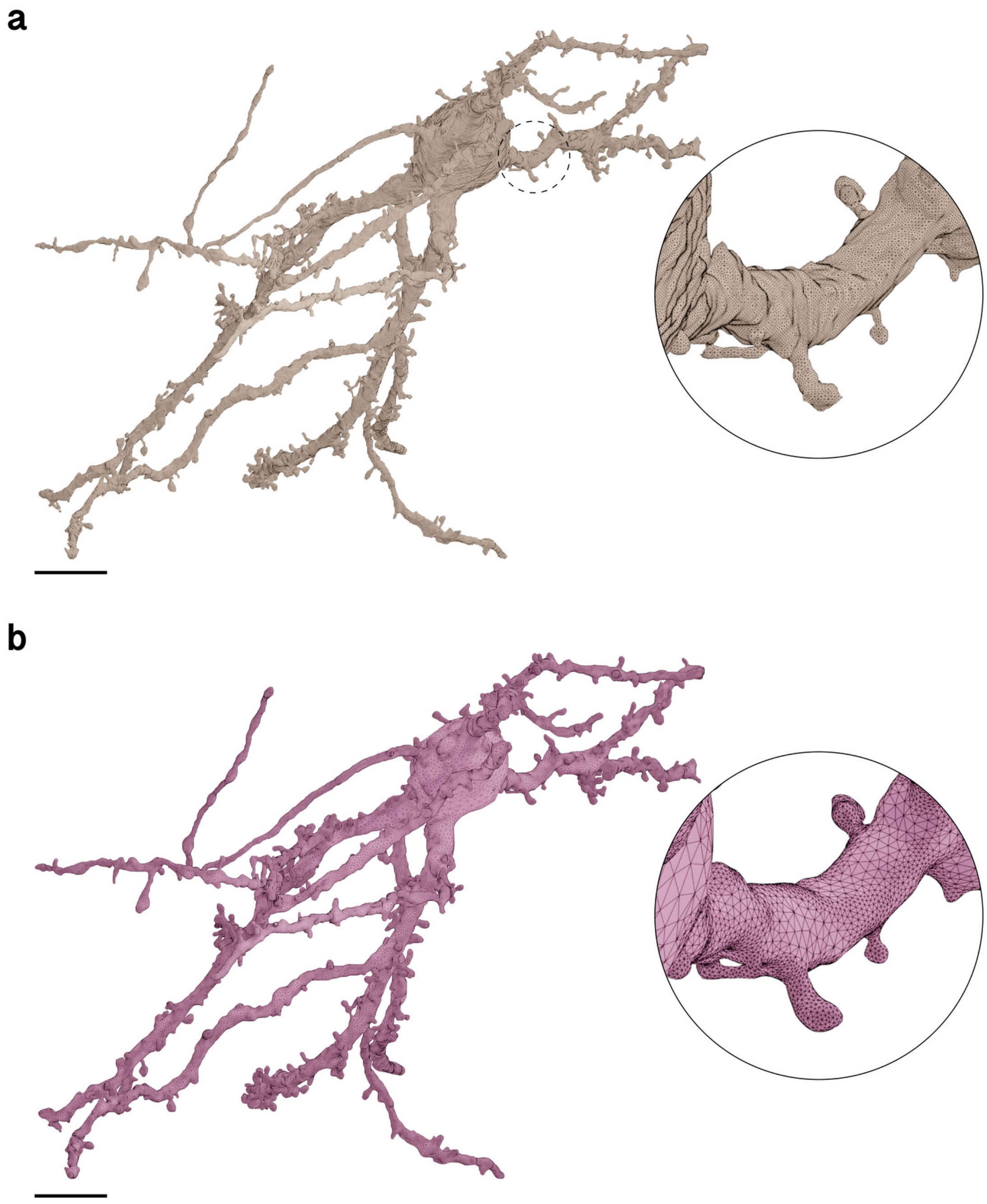

Figure S12: Wireframe visualizations comparing input (**a**) and resulting (**b**) surface meshes of Neuron 3. Fragmented partitions and floating vertices are automatically removed to yield a single mesh partition with continuous membrane surface. The closeups highlight the contrast in surface roughness, topology and tessellation. Comparative quantitative and qualitative analysis is shown in Figure S14c. Scale bars, 5 $\mu$m (**a, b**).

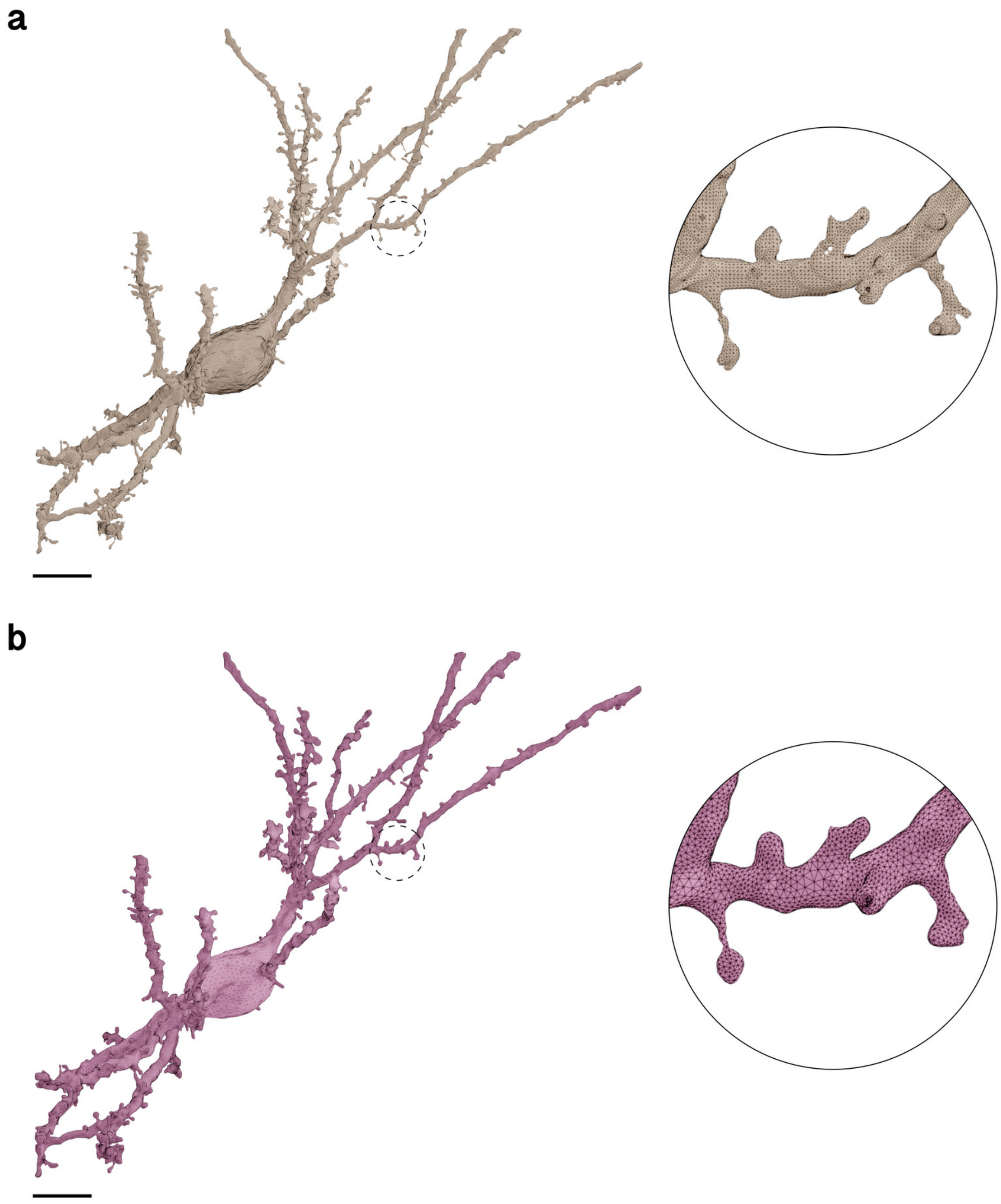

Figure S13: Wireframe visualizations comparing input (**a**) and resulting (**b**) surface meshes of Neuron 2. Fragmented partitions and floating vertices are automatically removed to yield a single mesh partition with continuous membrane surface. The closeups highlight the contrast in surface roughness, topology and tessellation. Comparative quantitative and qualitative analysis is shown in Figure S14d. Scale bars, 5 $\mu$m (**a, b**).

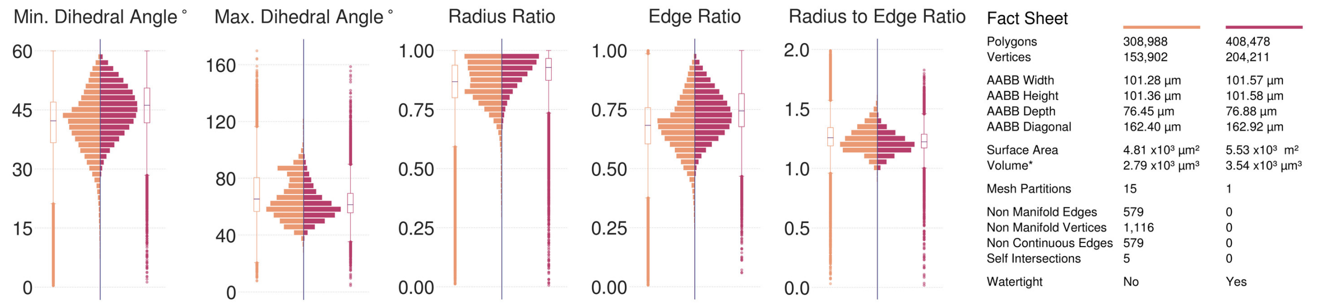

(**a**) Neuron 1, Visualization in Figure S10.

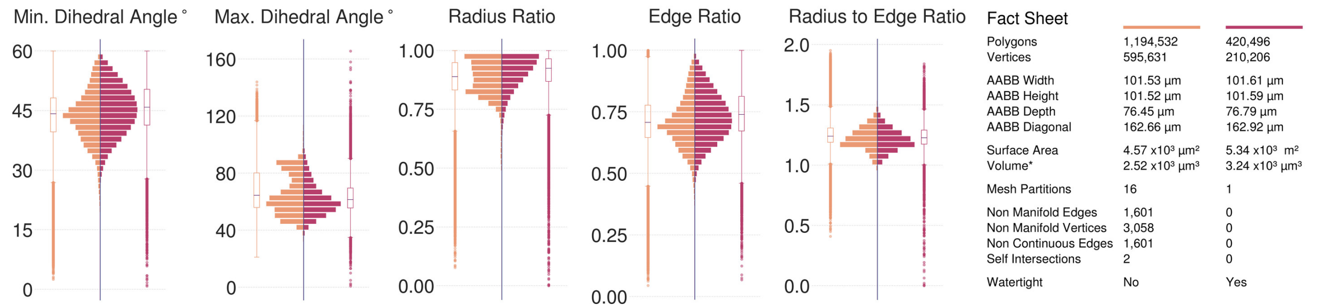

(**b**) Neuron 2, Visualization in Figure S11.

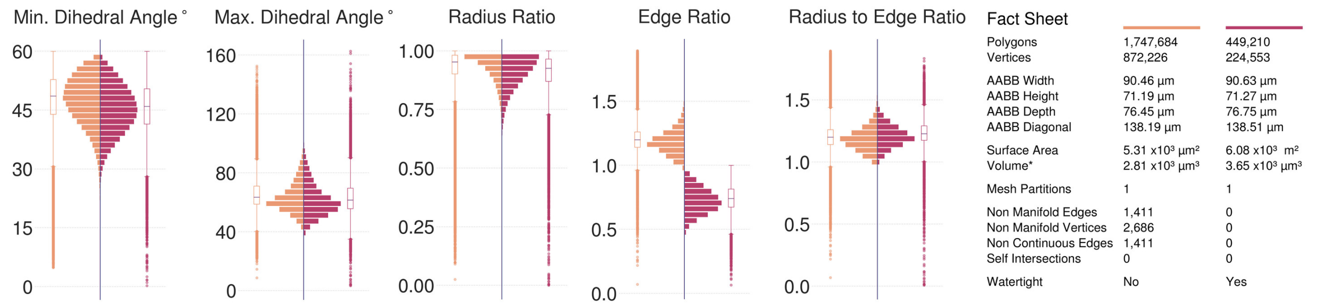

(**c**) Neuron 3, Visualization in Figure S12.

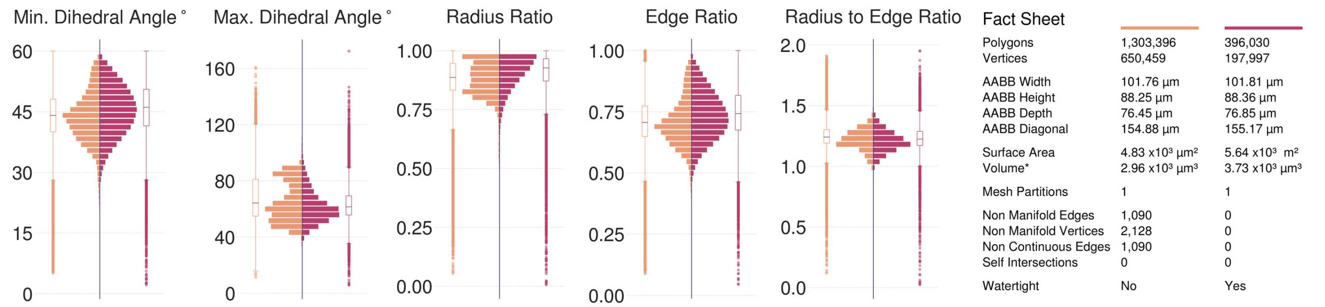

(**d**) Neuron 4, Visualization in Figure S13.

Figure S14: Comparative analysis between input (left in orange) and resulting (right in dark salmon) meshes of the complete neurons segmented from the volume shown in Figure S3. All the input meshes are not watertight; they have multiple fragmented partitions and severe geometric artifacts. The resulting meshes are reconstructed at 5 voxels per micron resolution (200 nm) yielding single, continuous and adaptively optimized manifolds with no artifacts. Note that the resulting mesh of Astrocyte 3 is more tessellated than the input one as a result of its relatively large AABB.

#### 5.3 Microglia

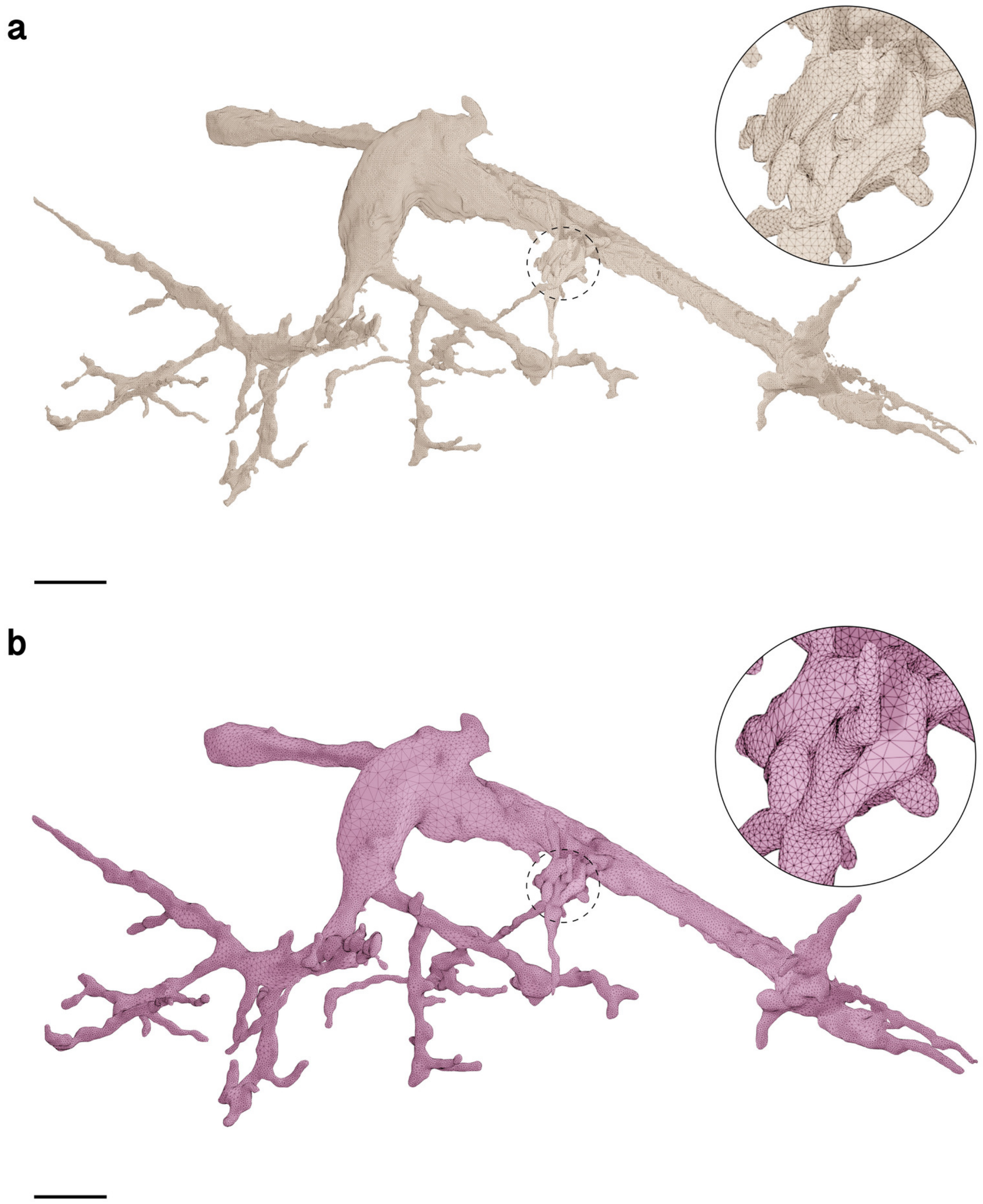

Figure S15: Wireframe visualizations comparing input (**a**) and resulting (**b**) surface meshes of Microglia 1. Fragmented partitions and floating vertices are automatically removed to yield a single mesh partition with continuous membrane surface. The closeups highlight the contrast in surface roughness, topology and tessellation. Comparative quantitative and qualitative analysis is shown in Figure S19a. Scale bars, 5 $\mu$m (**a, b**).

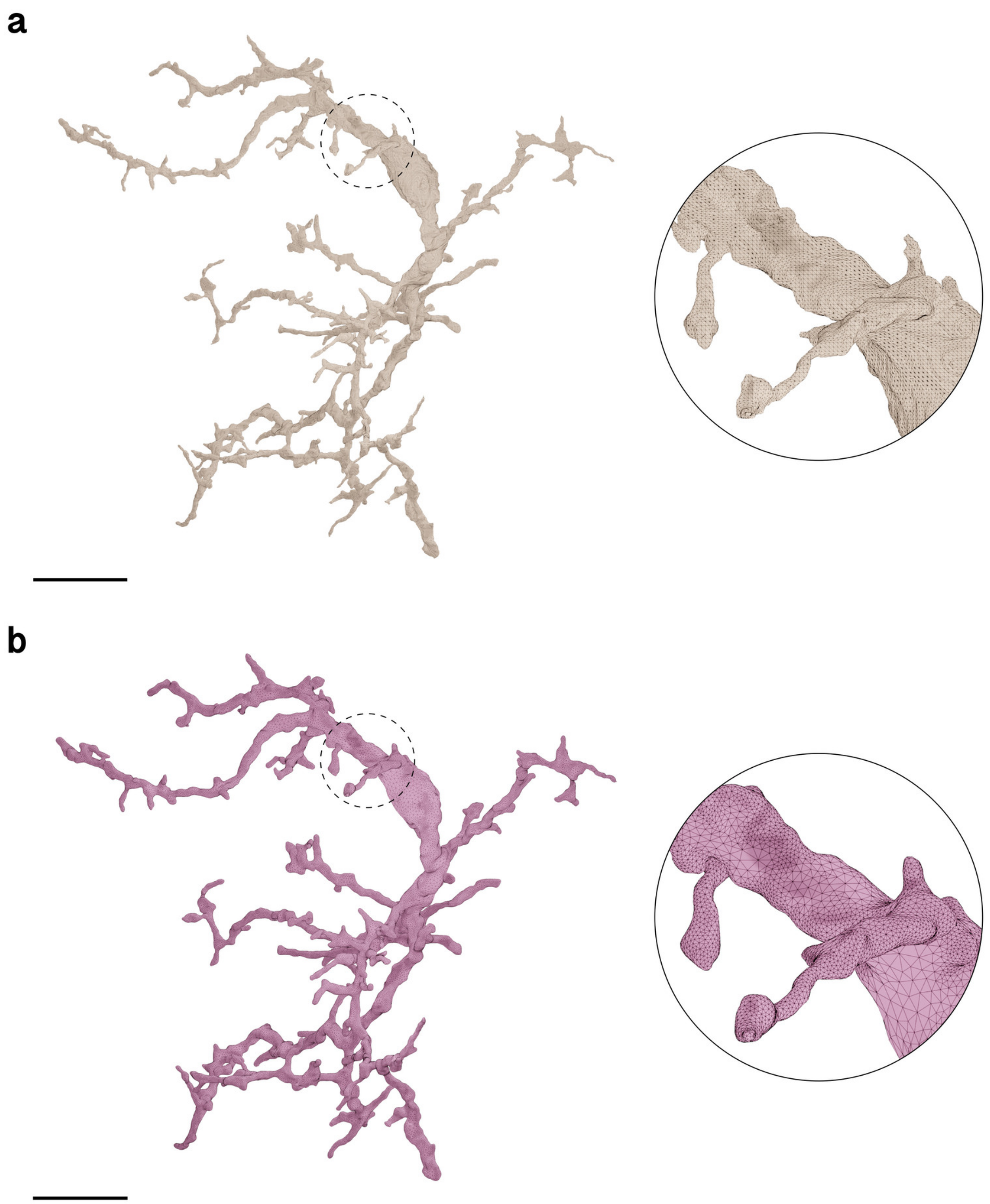

Figure S16: Wireframe visualizations comparing input (**a**) and resulting (**b**) surface meshes of Microglia 2. Fragmented partitions and floating vertices are automatically removed to yield a single mesh partition with continuous membrane surface. The closeups highlight the contrast in surface roughness, topology and tessellation. Comparative quantitative and qualitative analysis is shown in Figure S19b.
Scale bars, 10 $\mu$m (**a, b**).

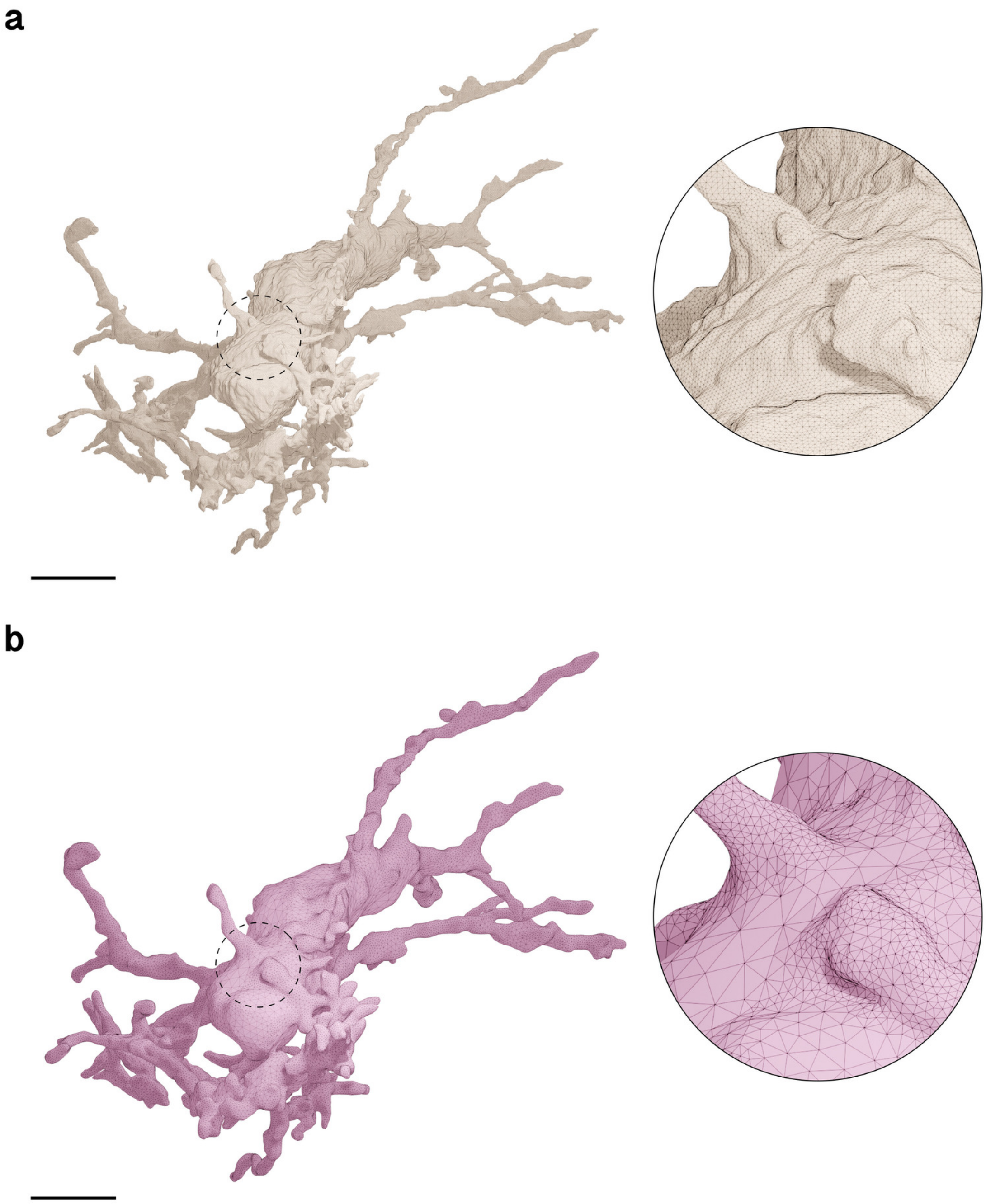

Figure S17: Wireframe visualizations comparing input (**a**) and resulting (**b**) surface meshes of Microglia 3. Fragmented partitions and floating vertices are automatically removed to yield a single mesh partition with continuous membrane surface. The closeups highlight the contrast in surface roughness, topology and tessellation. Comparative quantitative and qualitative analysis is shown in Figure S19c. Scale bars, 5 $\mu$m (**a, b**).

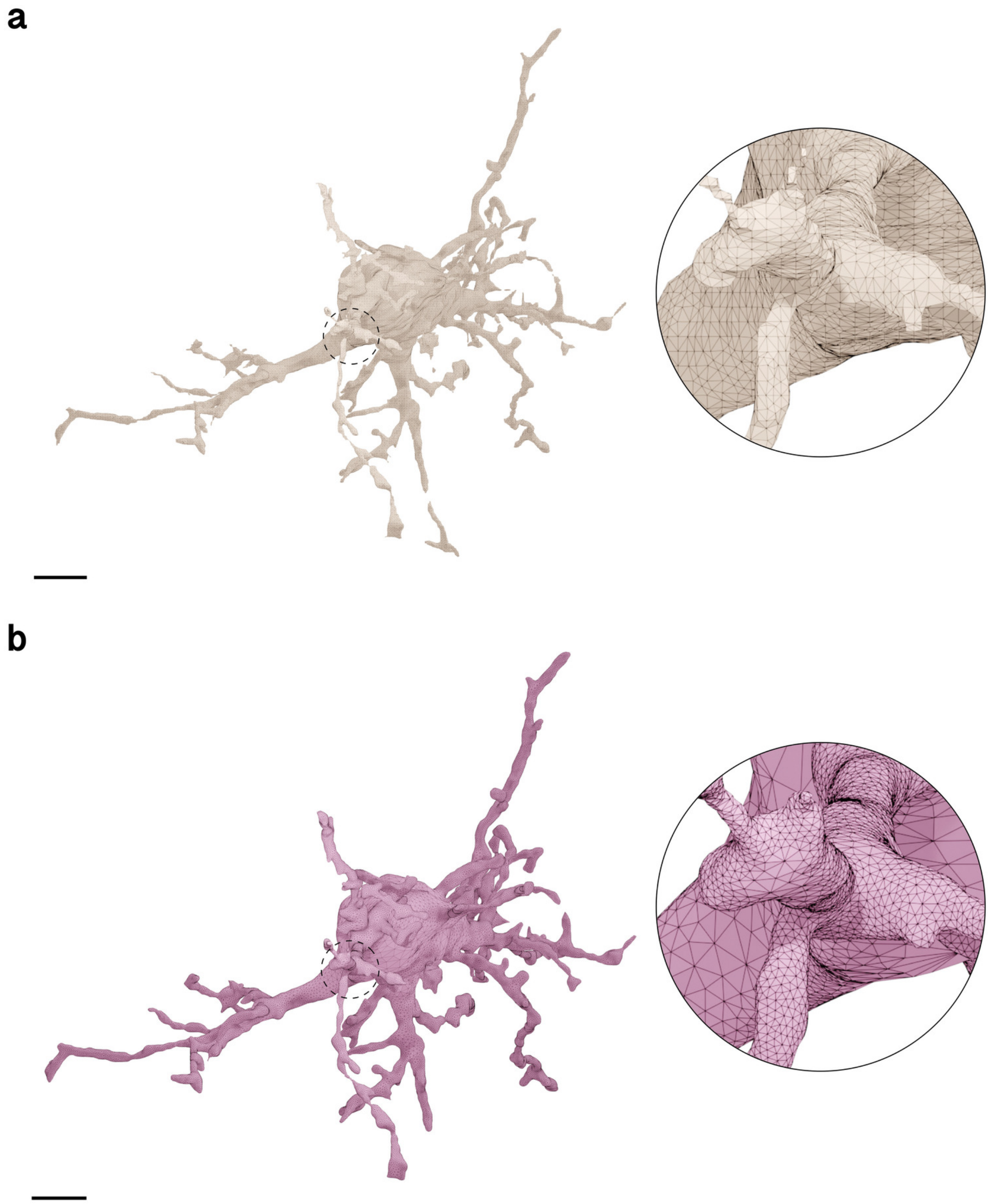

Figure S18: Wireframe visualizations comparing input (**a**) and resulting (**b**) surface meshes of Microglia 4. Fragmented partitions and floating vertices are automatically removed to yield a single mesh partition with continuous membrane surface. The closeups highlight the contrast in surface roughness, topology and tessellation. Comparative quantitative and qualitative analysis is shown in Figure S19d.
Scale bars, 5 $\mu$m (**a, b**).

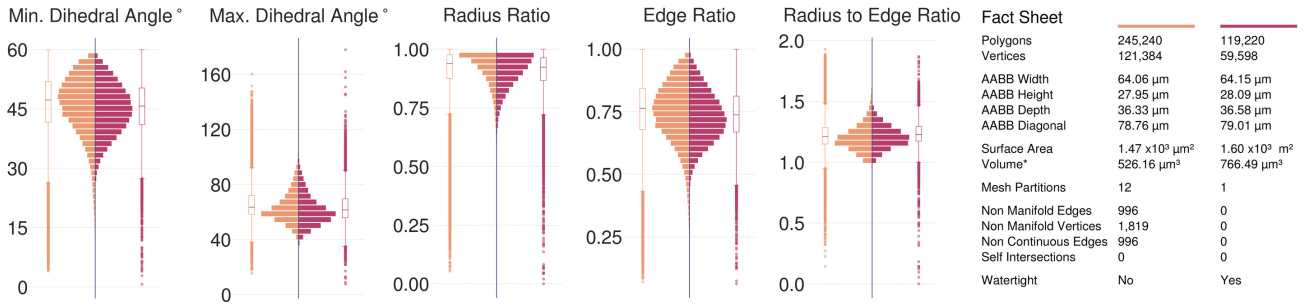

(**a**) Microglia 1, Visualization in Figure S15.

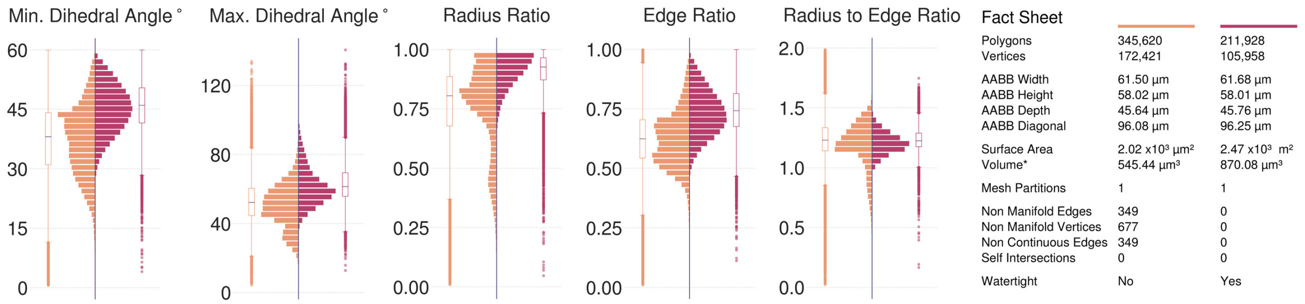

(**b**) Microglia 2, Visualization in Figure S16.

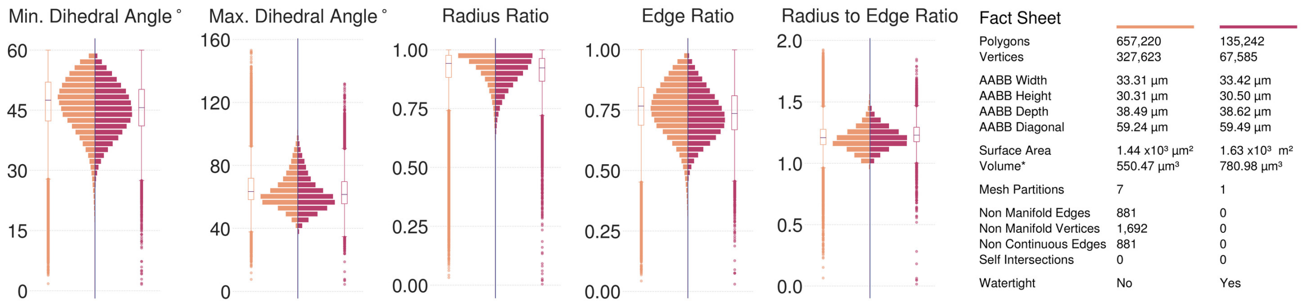

(**c**) Microglia 3, Visualization in Figure S17.

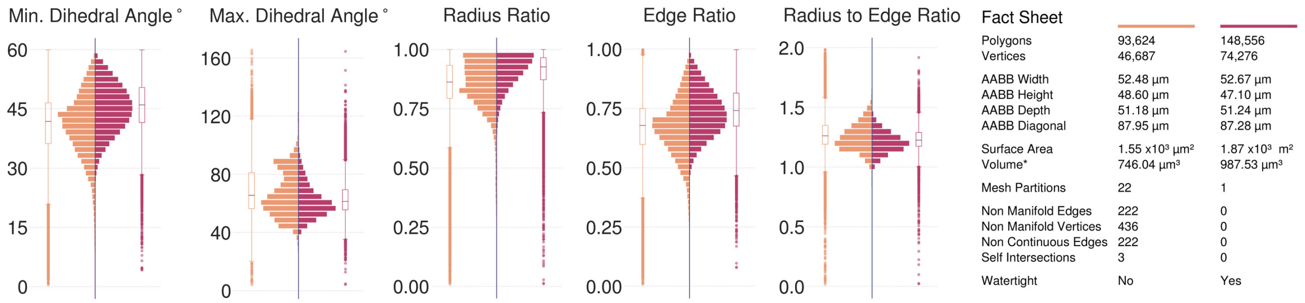

(**d**) Microglia 4, Visualization in Figure S18.

Figure S19: Comparative analysis between input (left in orange) and resulting (right in dark salmon) meshes of the complete microglia segmented from the volume shown in Figure S3. All the input meshes are not watertight. The resulting meshes are reconstructed at 5 voxels per micron resolution (200 nm) yielding single, continuous and adaptively optimized manifolds with no artifacts.

##### 5.4 Pericytes

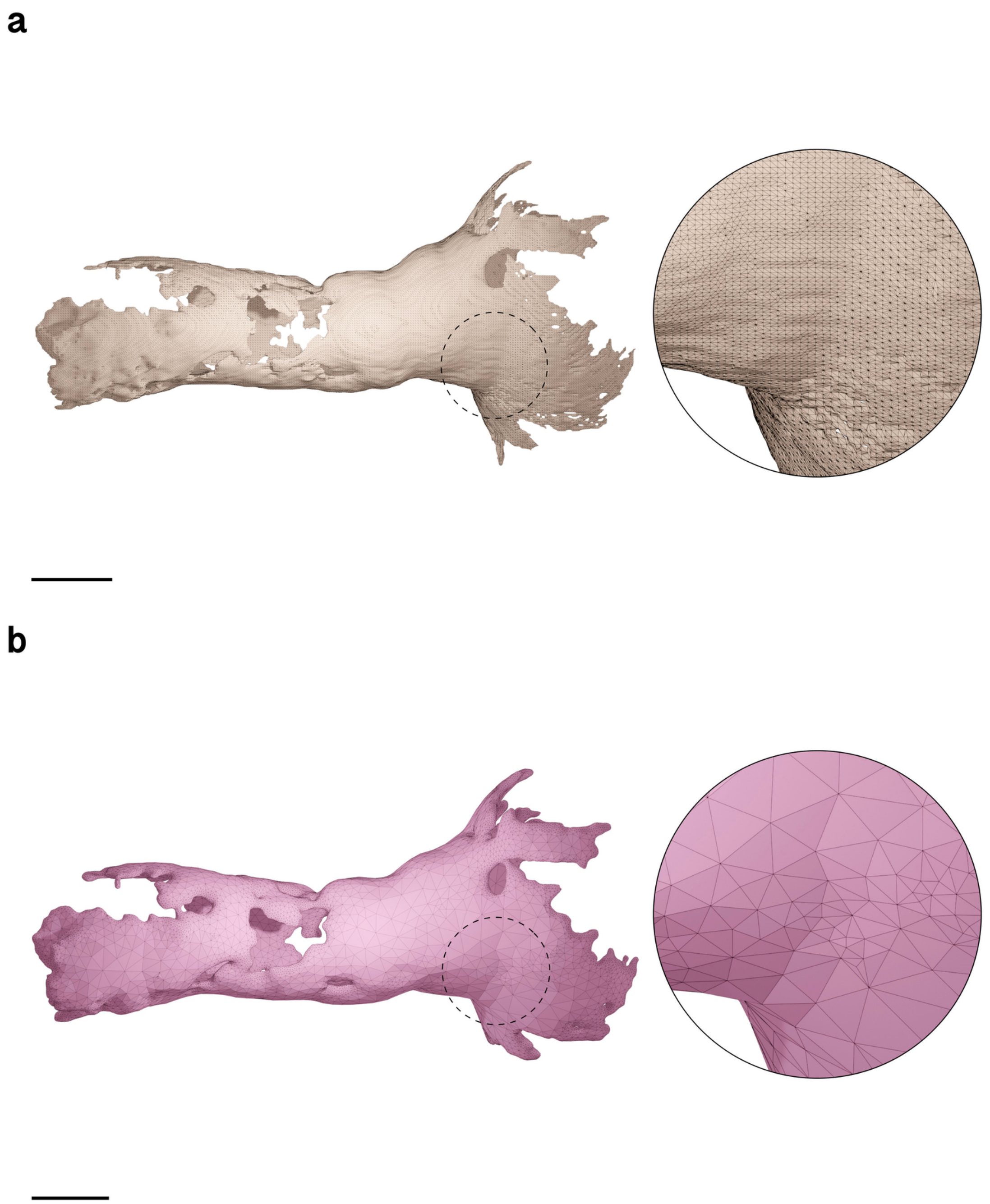

Figure S20: Wireframe visualizations comparing input (**a**) and resulting (**b**) surface meshes of Pericyte 1. The closeups highlight the contrast in surface roughness, topology and tessellation. Comparative quantitative and qualitative analysis is shown in Figure S24a. Scale bars, 5 $\mu$m (**a, b**).

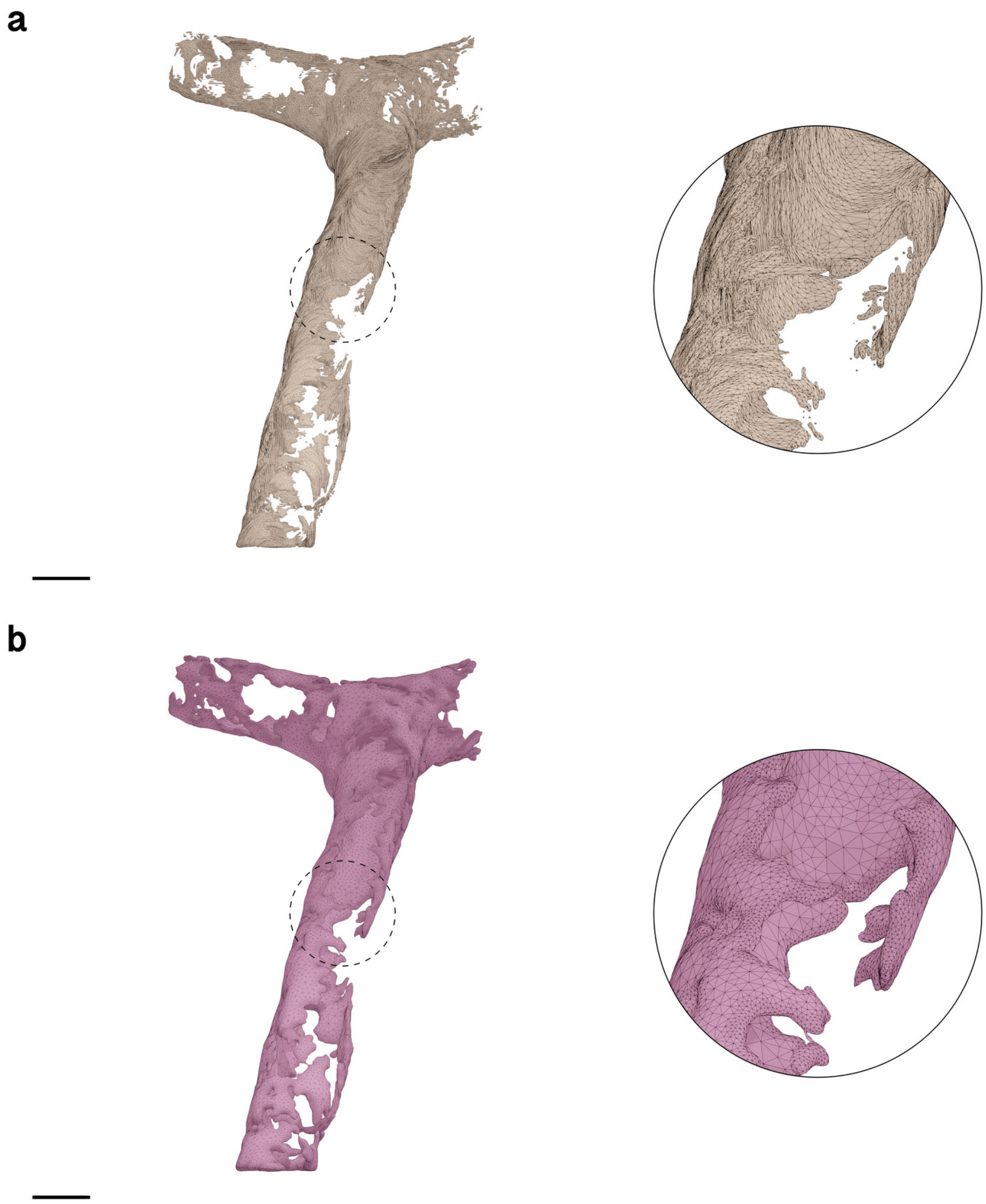

Figure S21: Wireframe visualizations comparing input (**a**) and resulting (**b**) surface meshes of Pericyte 2. The closeups highlight the contrast in surface roughness, topology and tessellation. Comparative quantitative and qualitative analysis is shown in Figure S24b. Scale bars, 10 $\mu$m (**a, b**).

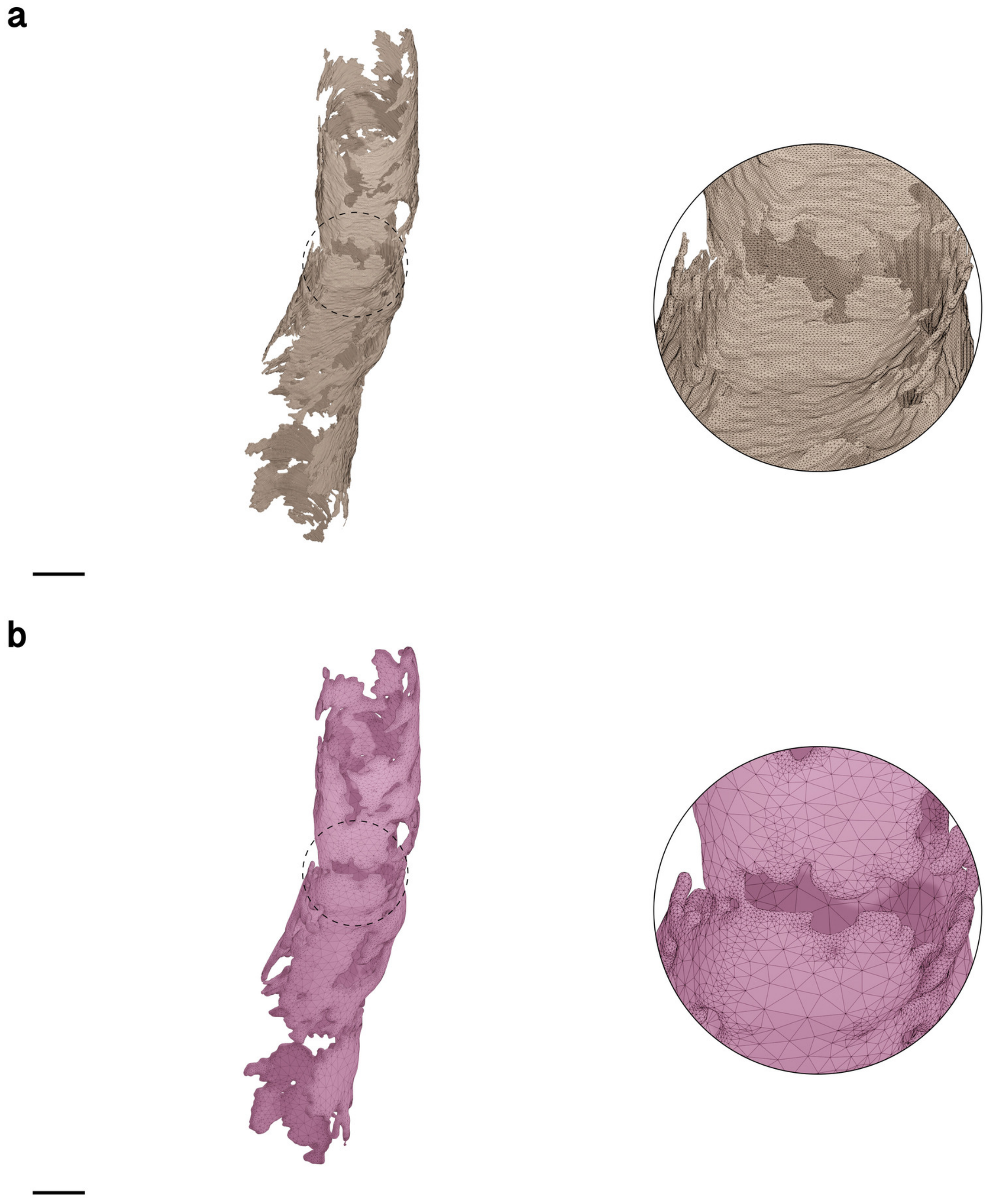

Figure S22: Wireframe visualizations comparing input (**a**) and resulting (**b**) surface meshes of Pericyte 3. The closeups highlight the contrast in surface roughness, topology and tessellation. Comparative quantitative and qualitative analysis is shown in Figure S24c. Scale bars, 5 $\mu$m (**a, b**).

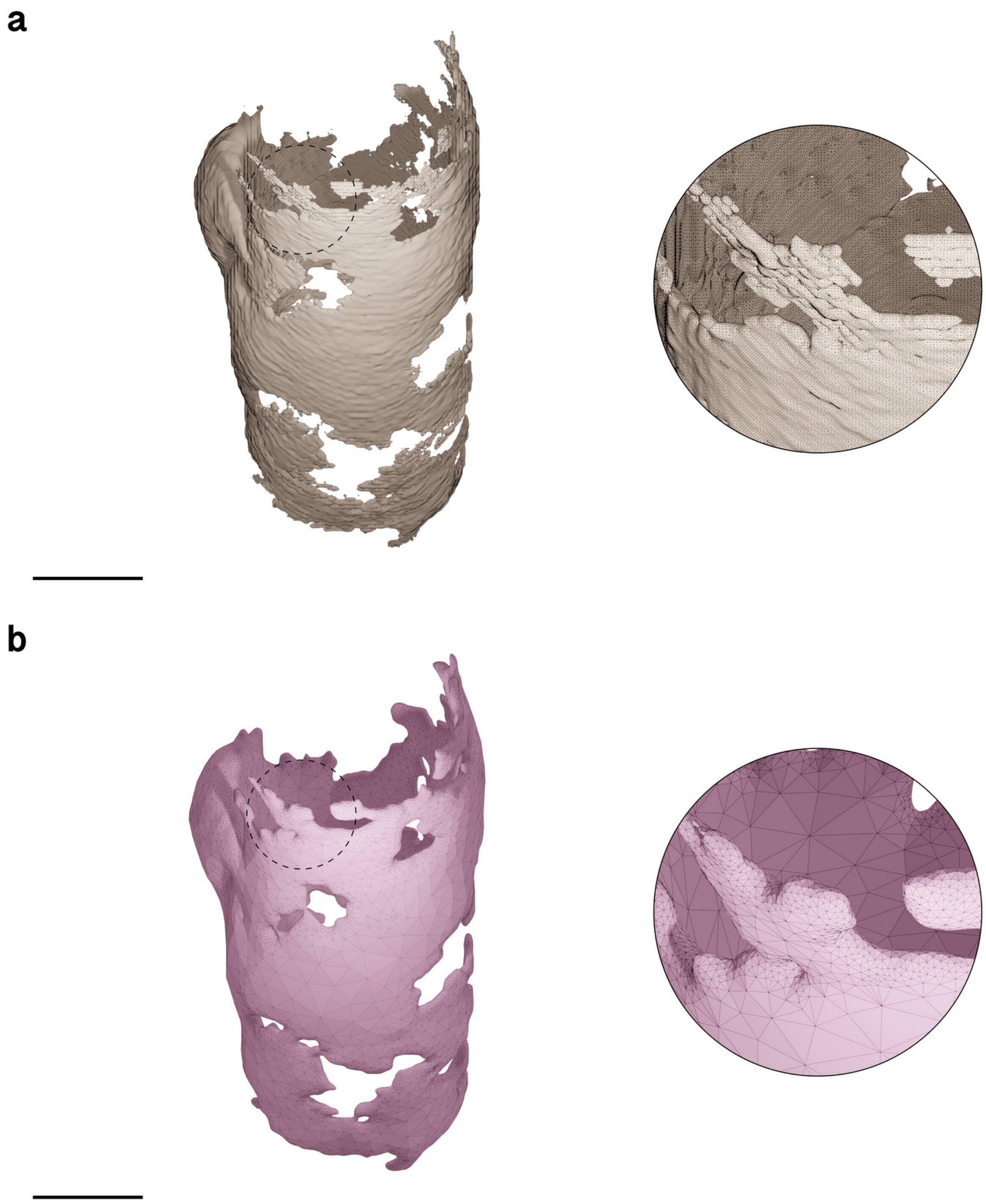

Figure S23: Wireframe visualizations comparing input (**a**) and resulting (**b**) surface meshes of Pericyte 4. The closeups highlight the contrast in surface roughness, topology and tessellation. Comparative quantitative and qualitative analysis is shown in Figure S24d. Scale bars, 5 $\mu$m (**a, b**).

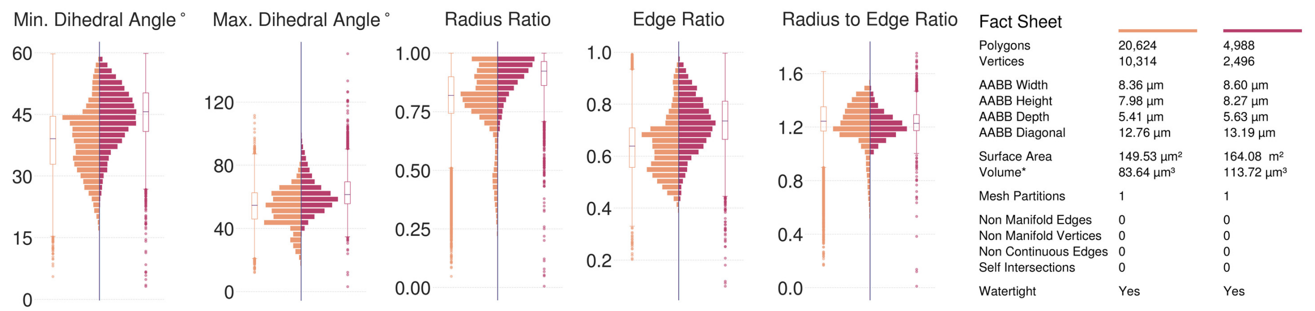

(**a**) Pericyte 1, Visualization in Figure S20.

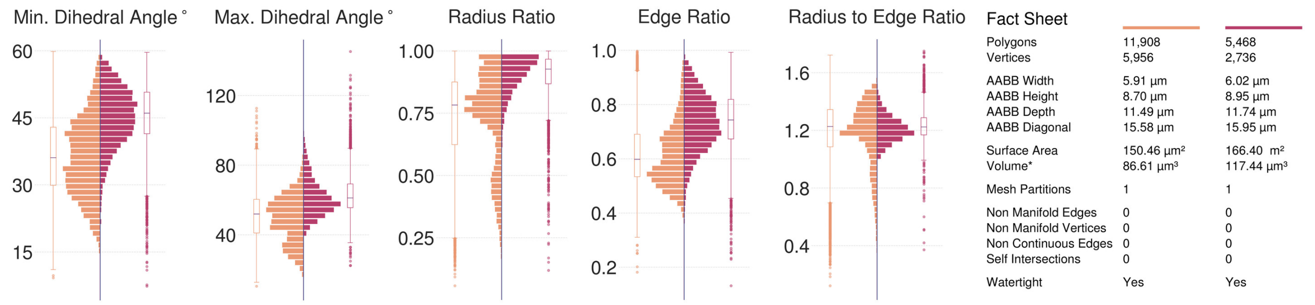

(**b**) Pericyte 2, Visualization in Figure S21.

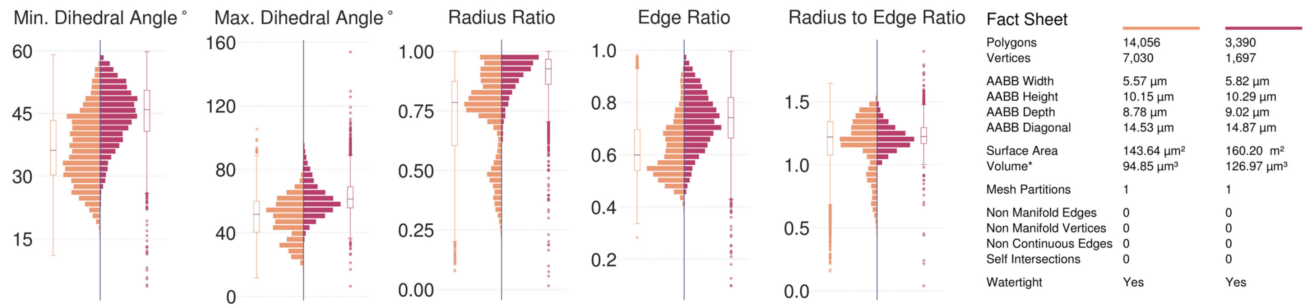

(**c**) Pericyte 3, Visualization in Figure S22.

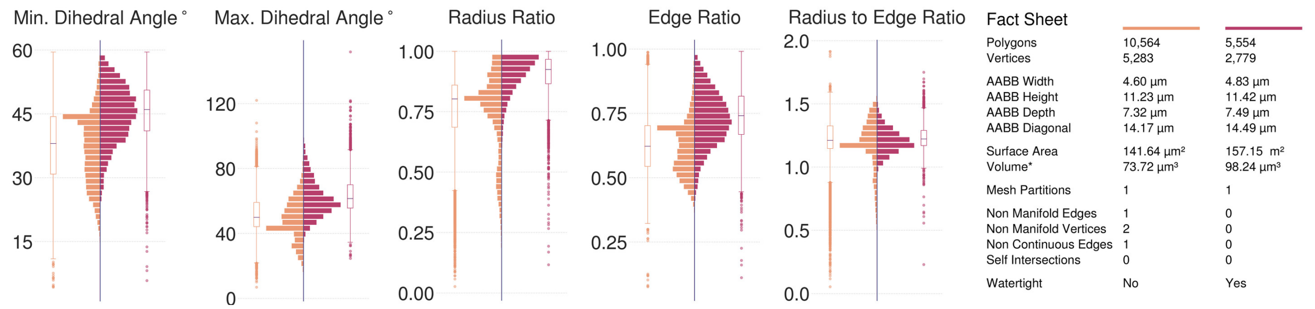

(**d**) Pericyte 4, Visualization in Figure S23.

Figure S24: Comparative analysis between input (left in orange) and resulting (right in dark salmon) meshes of the complete pericytes segmented from the volume shown in Figure S3. Although two input pericyte meshes are watertight, but their geometric qualitative analysis show their poor topological quality compared to the resulting ones. The resulting meshes are reconstructed at 5 voxels per micron resolution (200 nm) yielding single, continuous and adaptively optimized manifolds with no artifacts.

#### 5.5 Oligodendrocyte

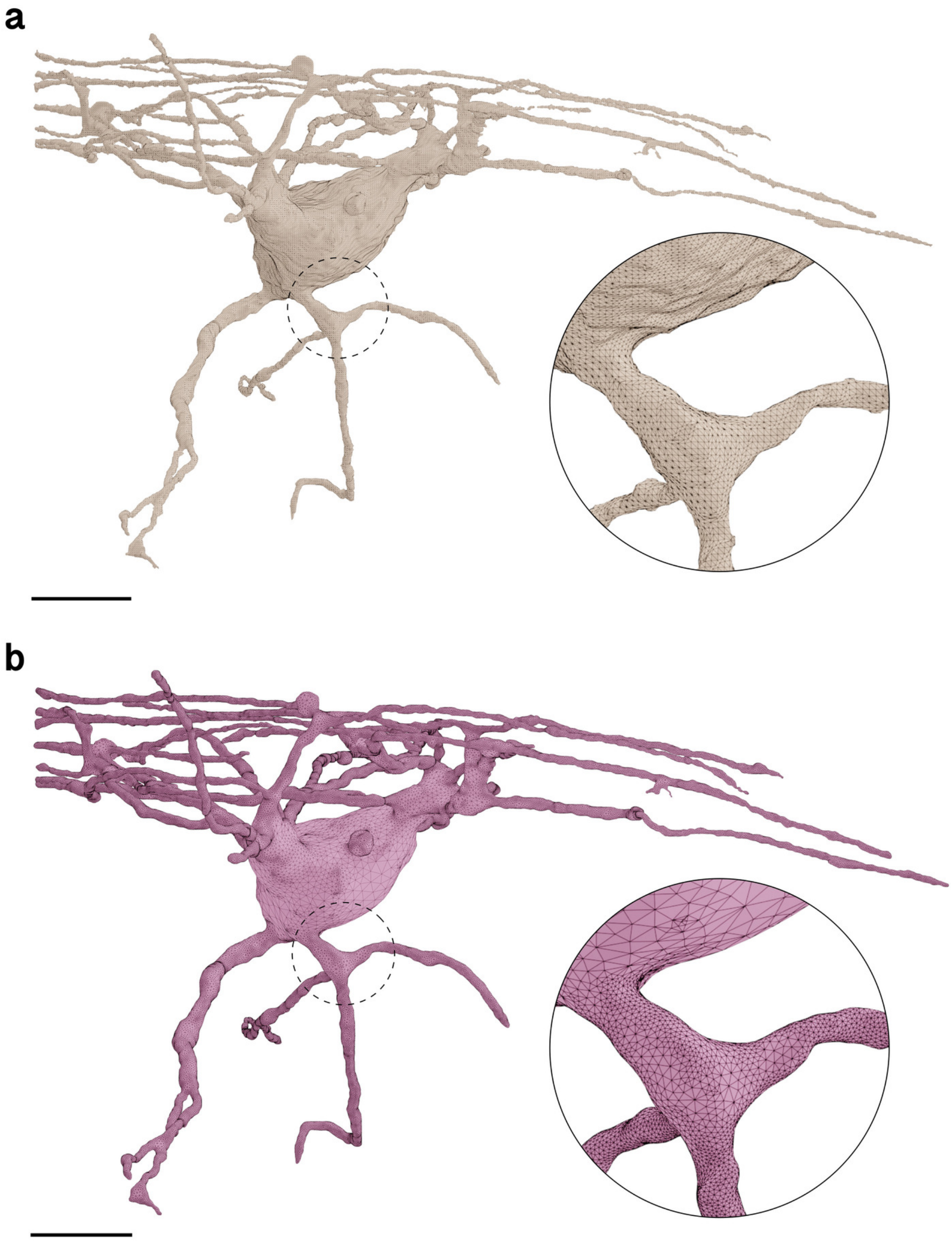

Figure S25: Wireframe visualizations comparing input (**a**) and resulting (**b**) surface meshes of the Oligodendrocyte. The closeups highlight the contrast in surface roughness, topology and tessellation. Comparative quantitative and qualitative analysis is shown in Figure S26. Scale bars, 5 $\mu$m (**a, b**).

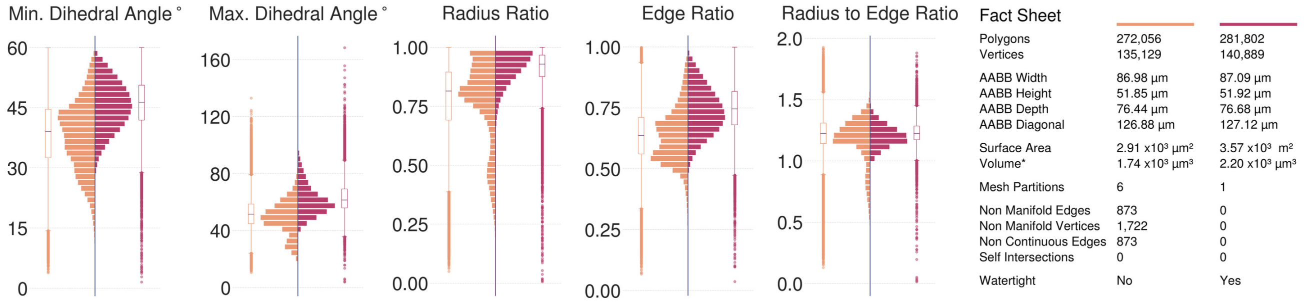

Figure S26: Comparative analysis between input (left in orange) and resulting (right in dark salmon) meshes of the oligodendrocyte segmented from the volume shown in Figure S3. The input mesh is not watertight due to the presence of non-manifold edges and vertices, but it has no self-intersections. Visualization in Figure S25.

#### 5.6 Blood Vessel Segments

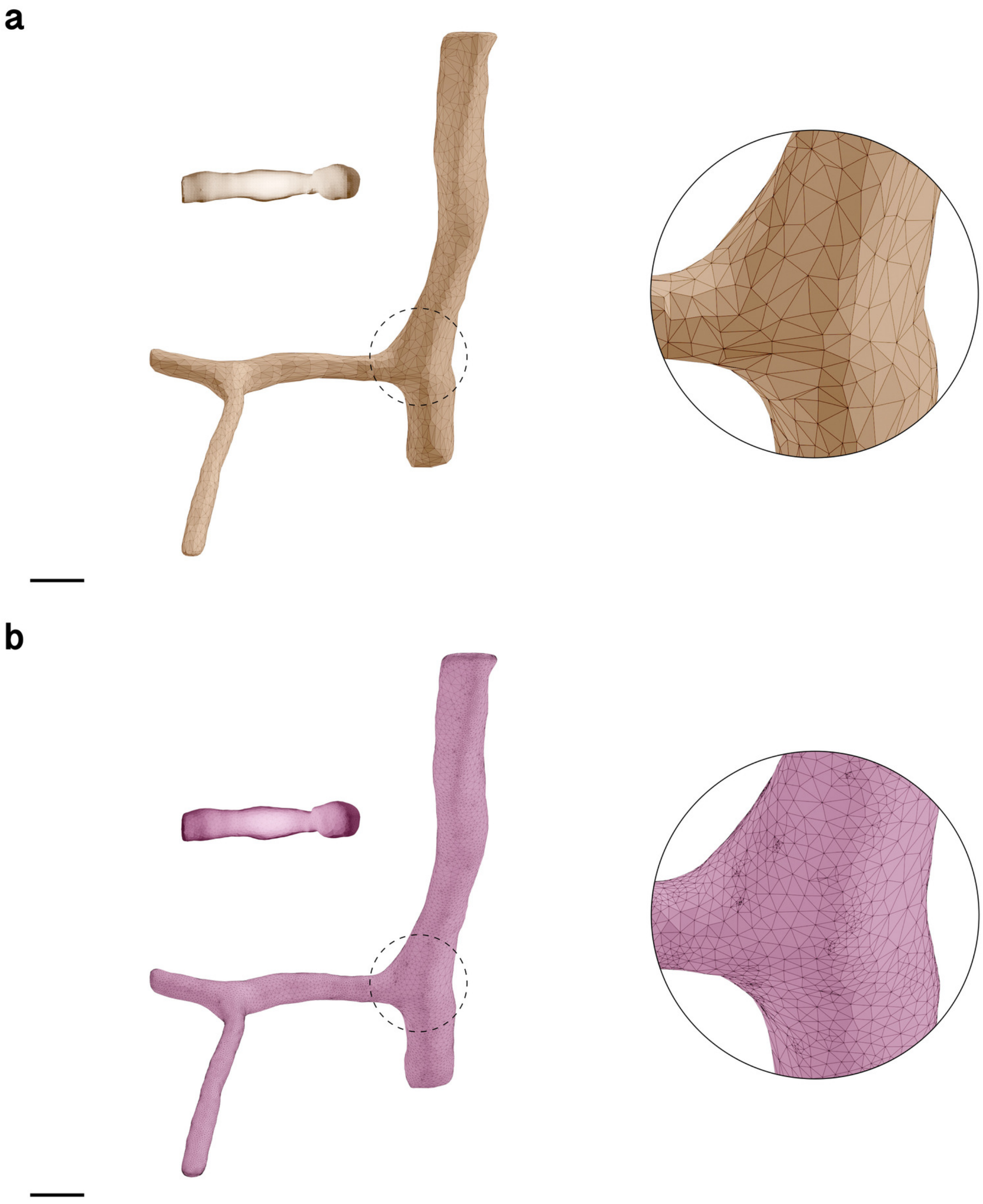

Figure S27: Wireframe visualizations comparing input (**a**) and resulting (**b**) surface meshes of the blood vessels segmented from the volume shown in Figure S3. Scale bars, 5 $\mu$m (**a, b**).

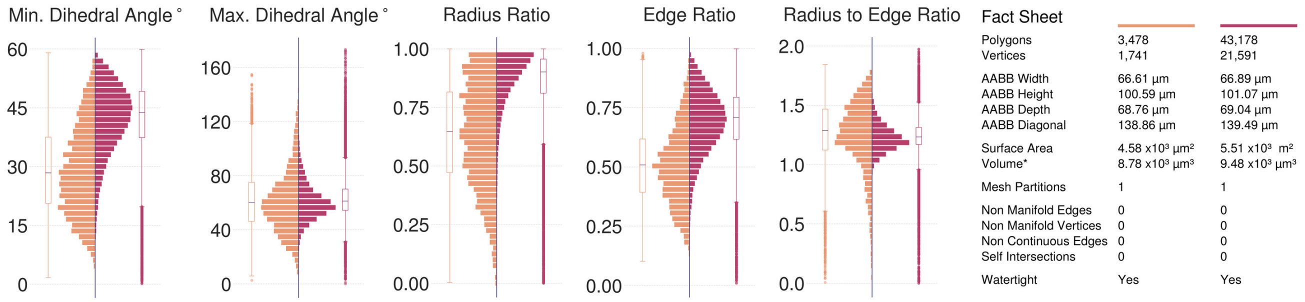

(a) Blood Vessel 1

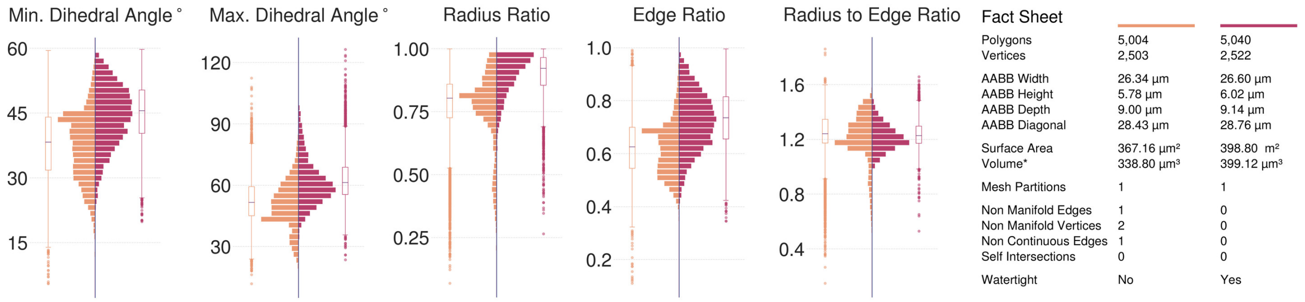

(b) Blood Vessel 2

Figure S28: Comparative analysis between input (left in orange) and resulting (right in dark salmon) meshes of the blood vessels segmented from the volume shown in Figure S3. Note that the input mesh representing Blood Vessel 1 is watertight, but has poor geometric quality compared to the resulting one. Visualization in Figure S27.

##### 5.7 Astrocytes Mitochondria

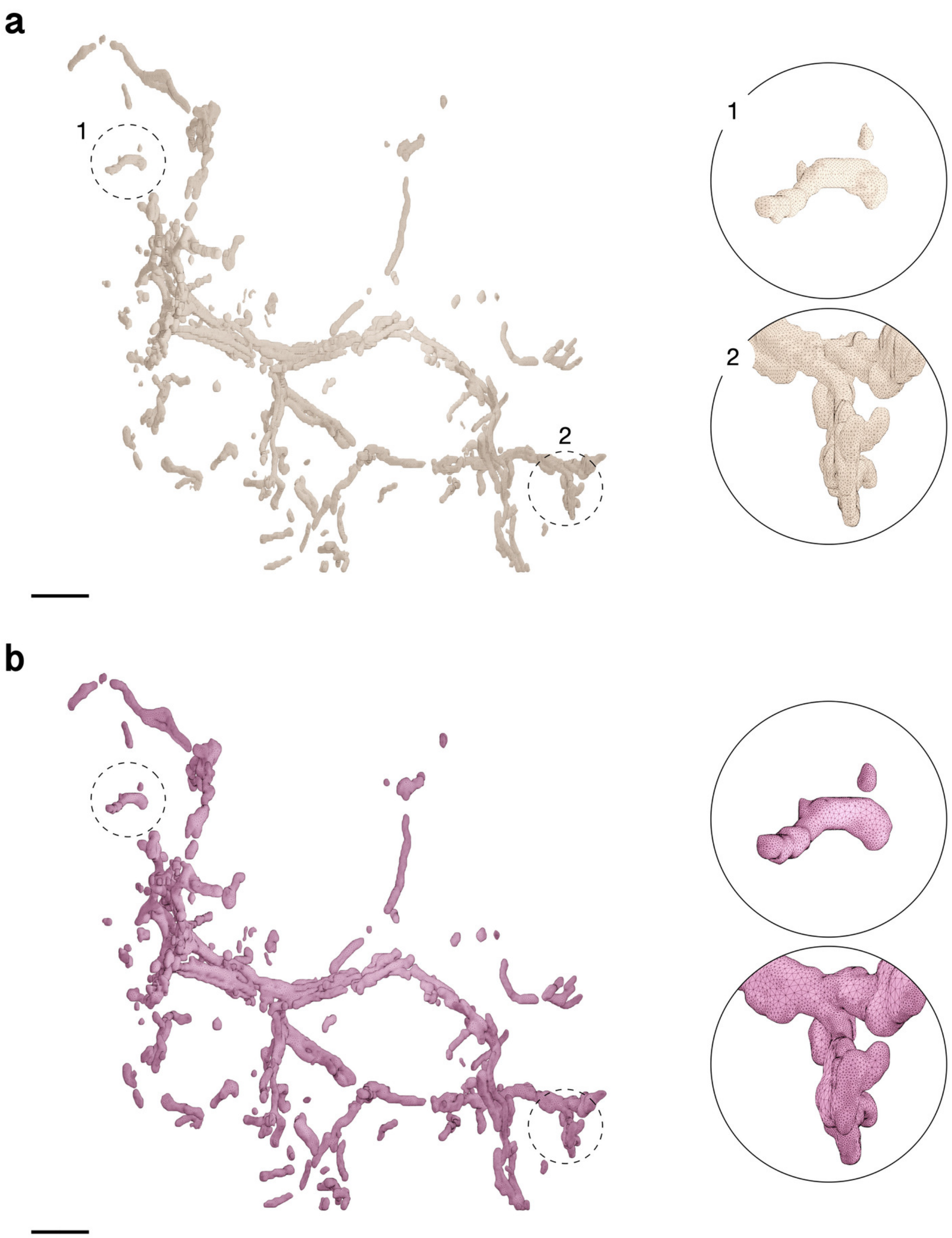

Figure S29: Comparative visual analysis between input (a) and reconstructed (b) meshes of the mitochondria of Astrocyte 1. The closeups highlight the contrast in surface roughness, topology and tessellation. Comparative quantitative and qualitative analysis is shown in Figure S33a. Scale bars, 5 $\mu$m (**a, b**).

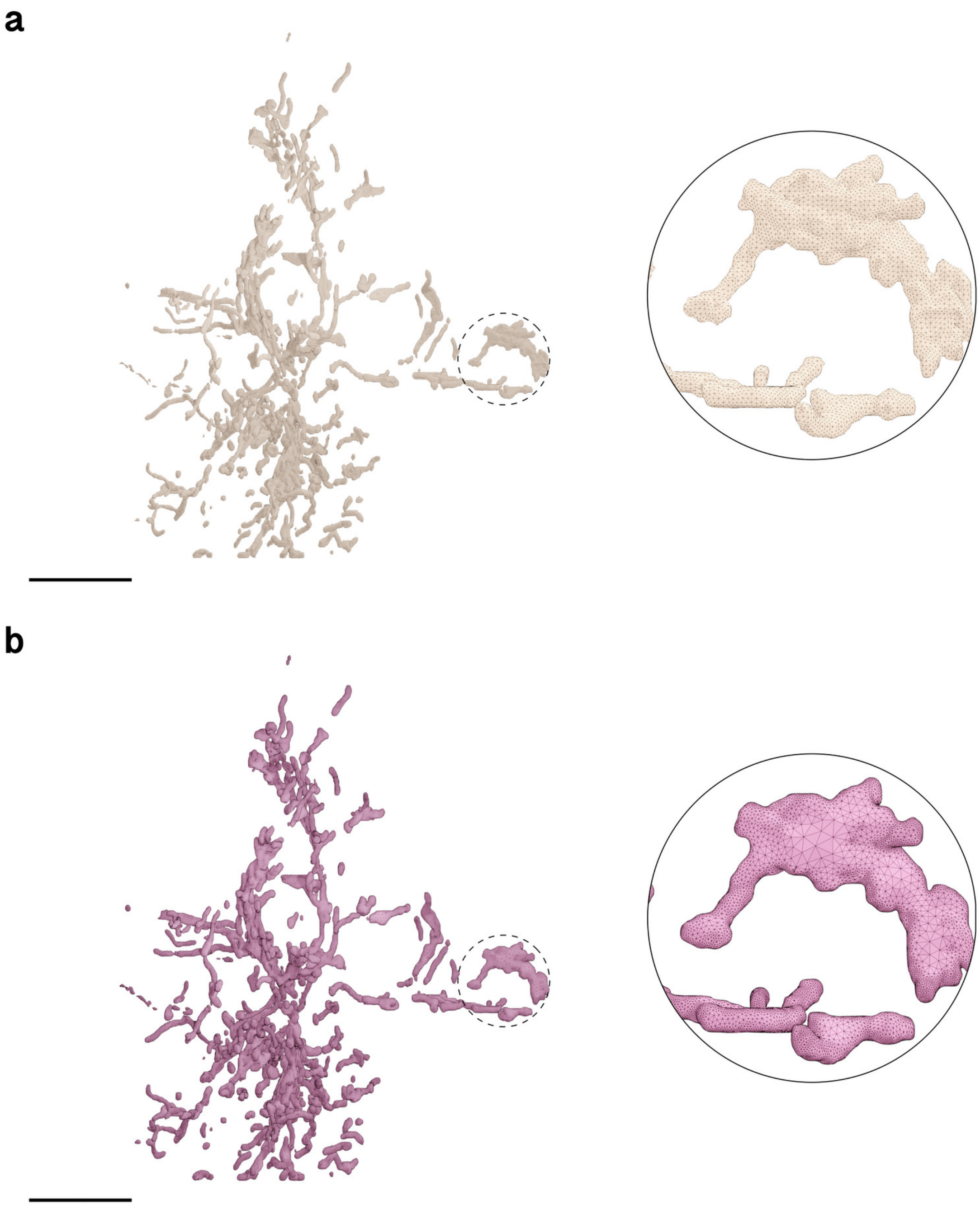

Figure S30: Comparative visual analysis between input (a) and reconstructed (b) meshes of the mitochondria of Astrocyte 2. The closeups highlight the contrast in surface roughness, topology and tessellation. Comparative quantitative and qualitative analysis is shown in Figure S33b. Scale bars, 10 $\mu$m (**a, b**).

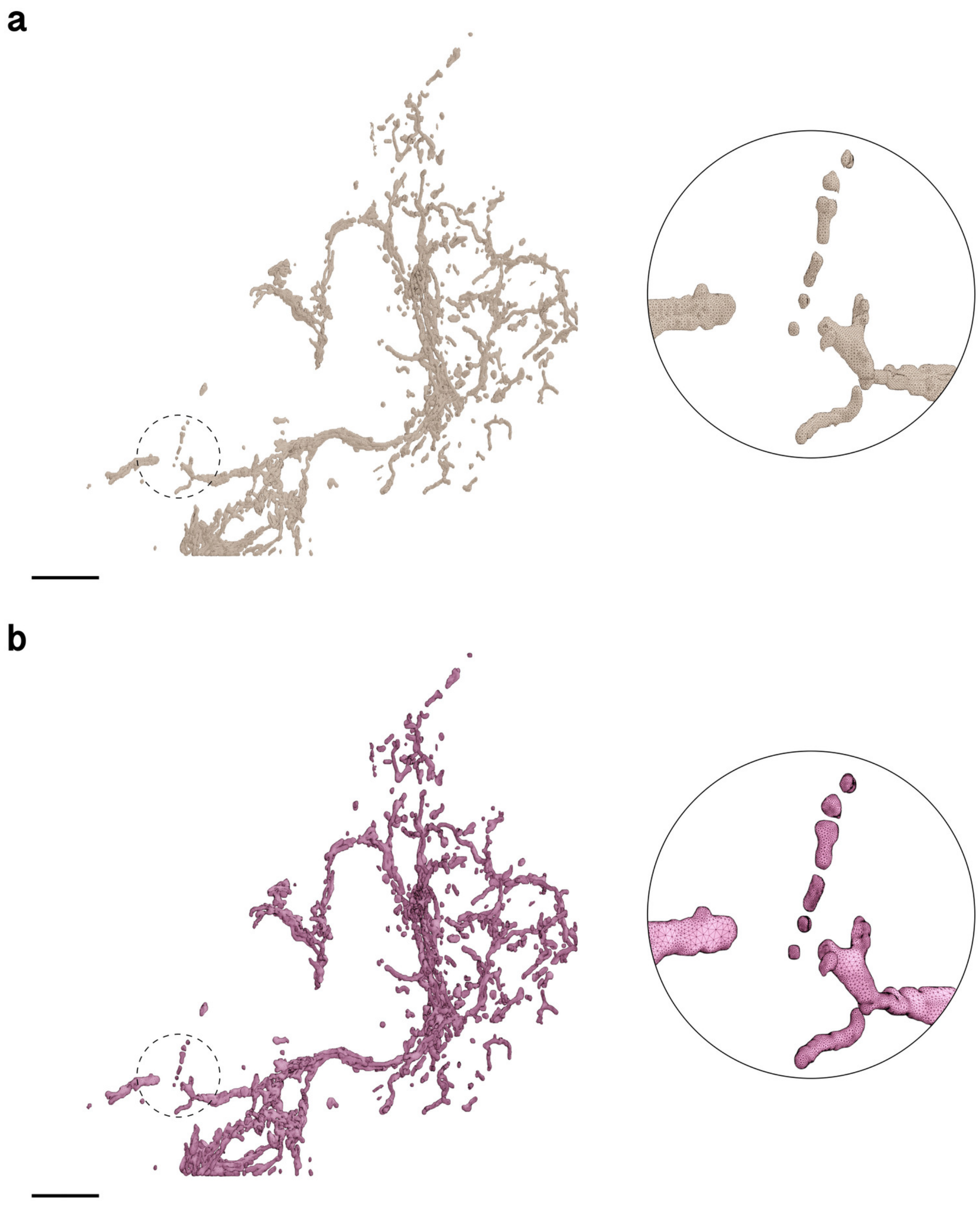

Figure S31: Comparative visual analysis between input (a) and reconstructed (b) meshes of of the mitochondria of Astrocyte 3. The closeups highlight the contrast in surface roughness, topology and tessellation. Comparative quantitative and qualitative analysis is shown in Figure S33c. Scale bars, 10 $\mu$m (**a, b**).

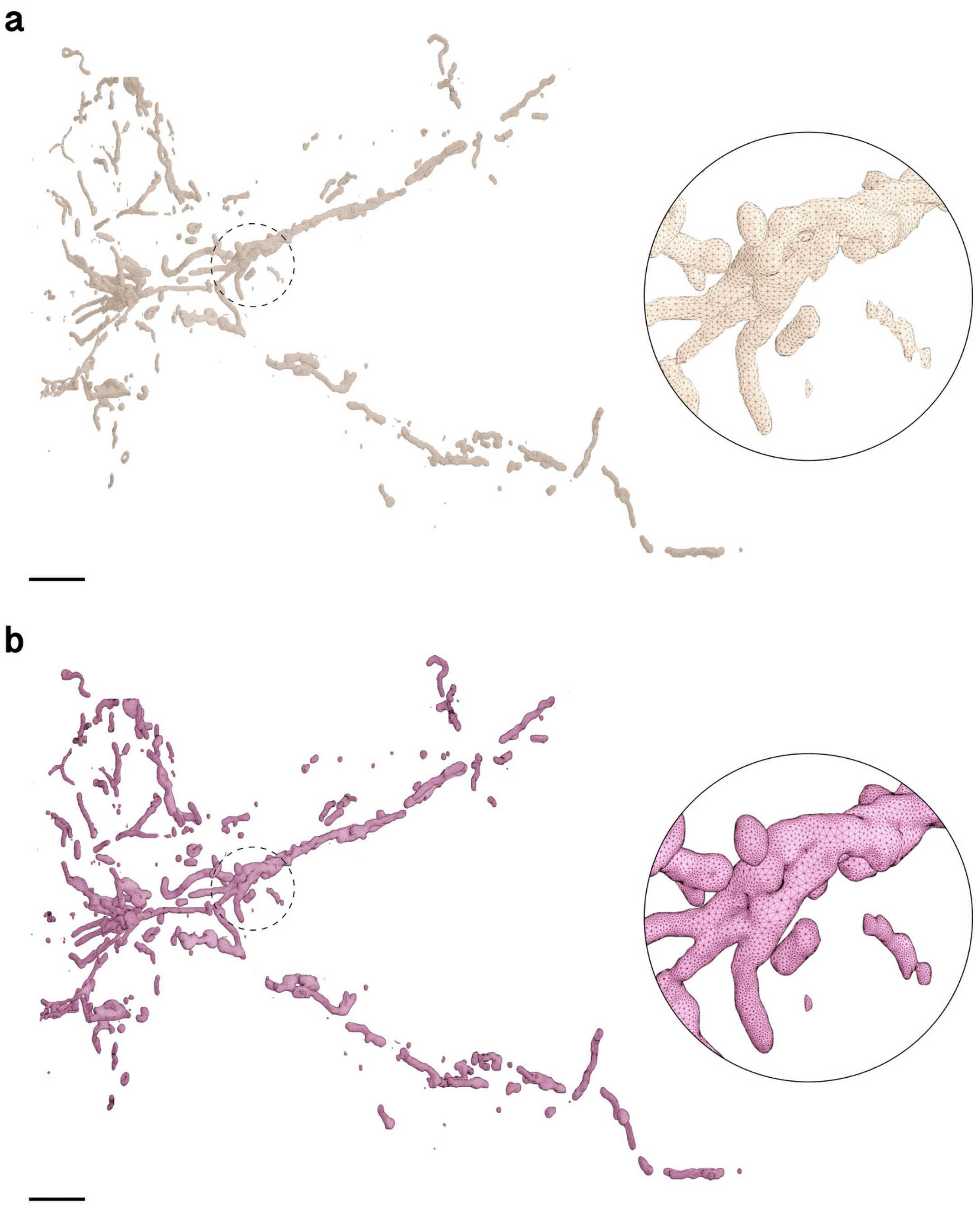

Figure S32: Comparative visual analysis between input (a) and reconstructed (b) meshes of of the mitochondria of Astrocyte 4. The closeups highlight the contrast in surface roughness, topology and tessellation. Comparative quantitative and qualitative analysis is shown in Figure S33d. Scale bars, 5 $\mu$m (**a, b**).

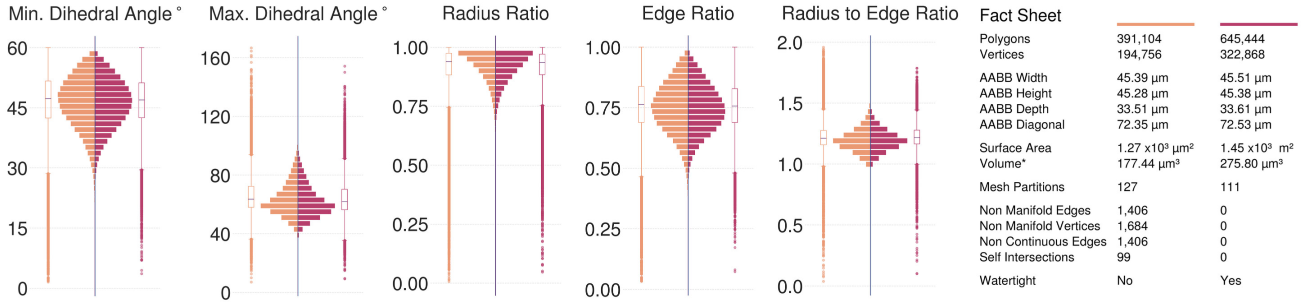

(a) Astrocyte 1 Mitochondria

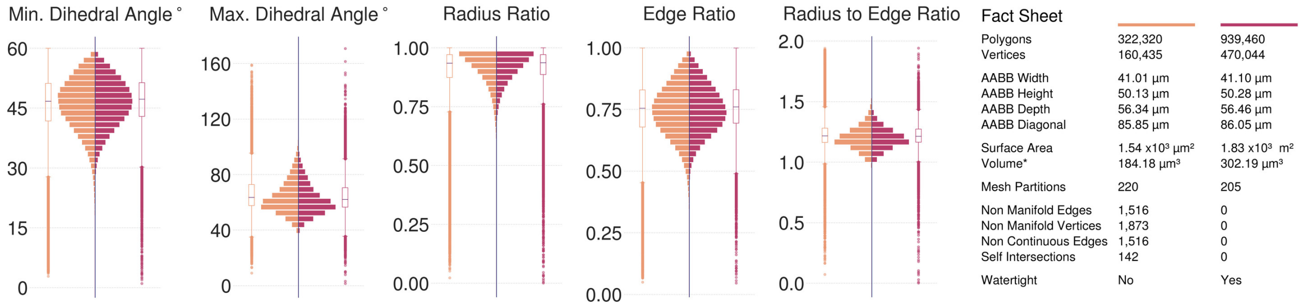

(b) Astrocyte 2 Mitochondria

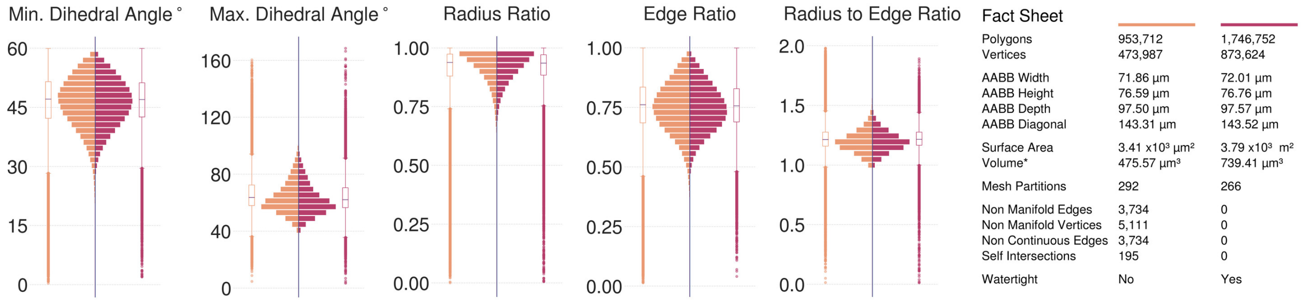

(c) Astrocyte 3 Mitochondria

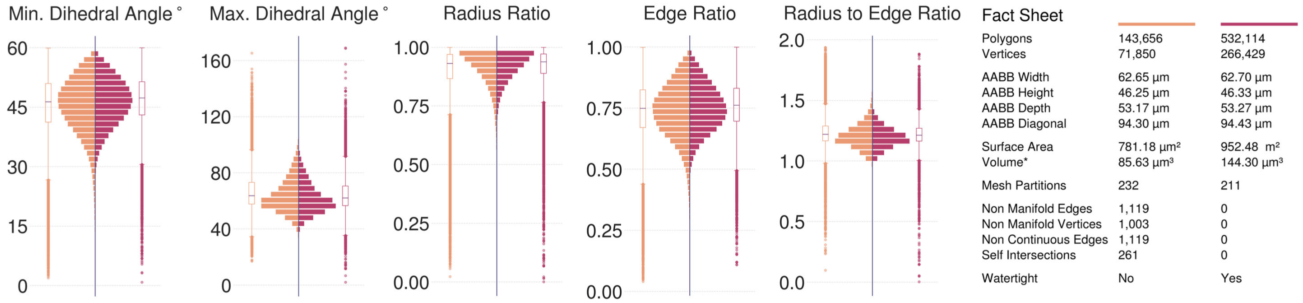

(d) Astrocyte 4 Mitochondria

Figure S33: Comparative analysis between input (left in orange) and resulting (right in dark salmon) meshes of the mitochondira of the four astrocytes segmented from the volume shown in Figure S3. All the input meshes are not watertight; they have self-intersections and non-manifold edges and vertices. The resulting meshes are reconstructed at 15 voxels per micron resolution (∼67 nm) yielding multi-partitioned, adaptively optimized and watertight manifolds with no artifacts.

#### 5.8 Neurons Mitochondria

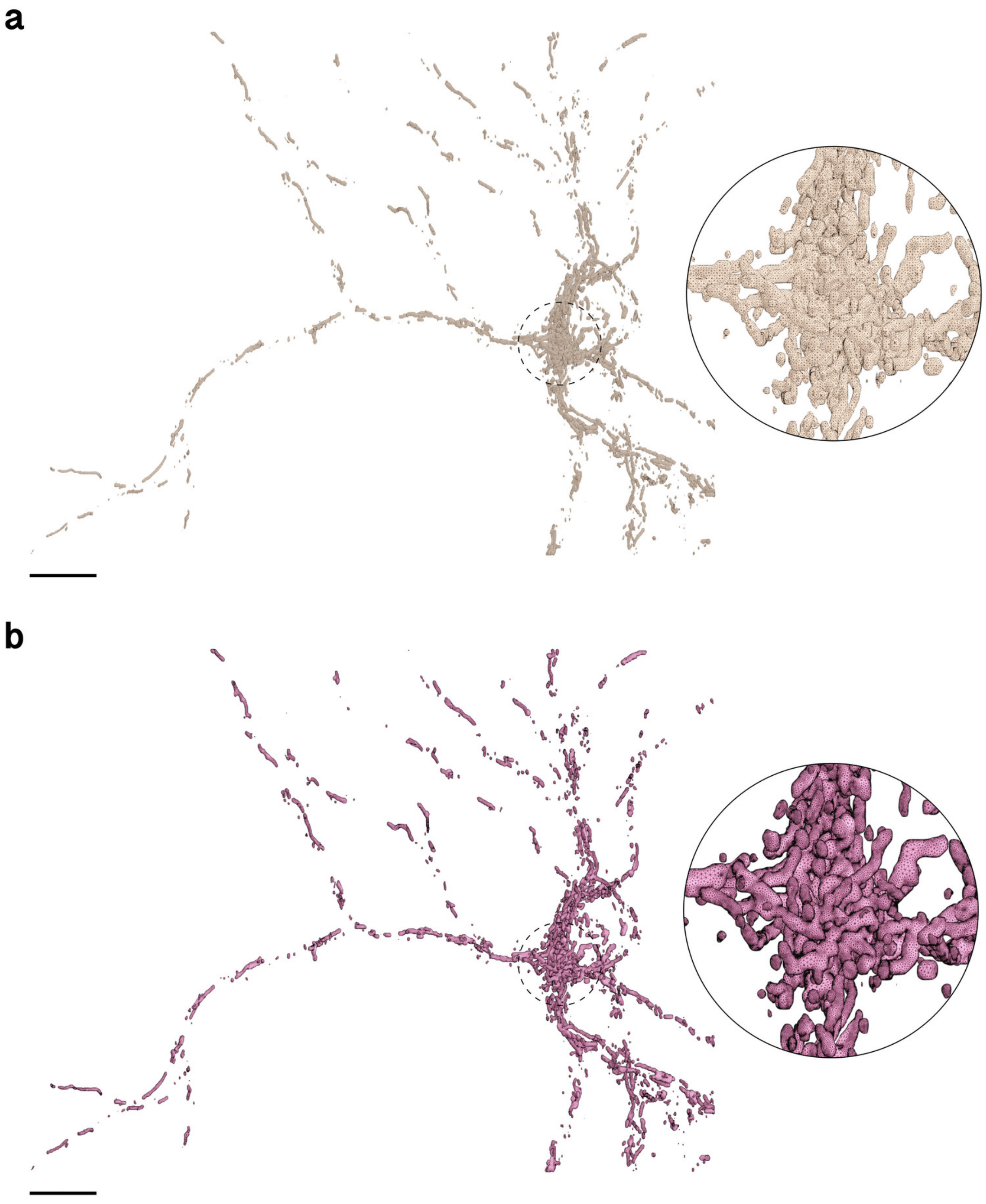

Figure S34: Comparative visual analysis between input (a) and reconstructed (b) meshes of of the mitochondria of Neuron 1. Scale bars, 10 $\mu$m (**a, b**).

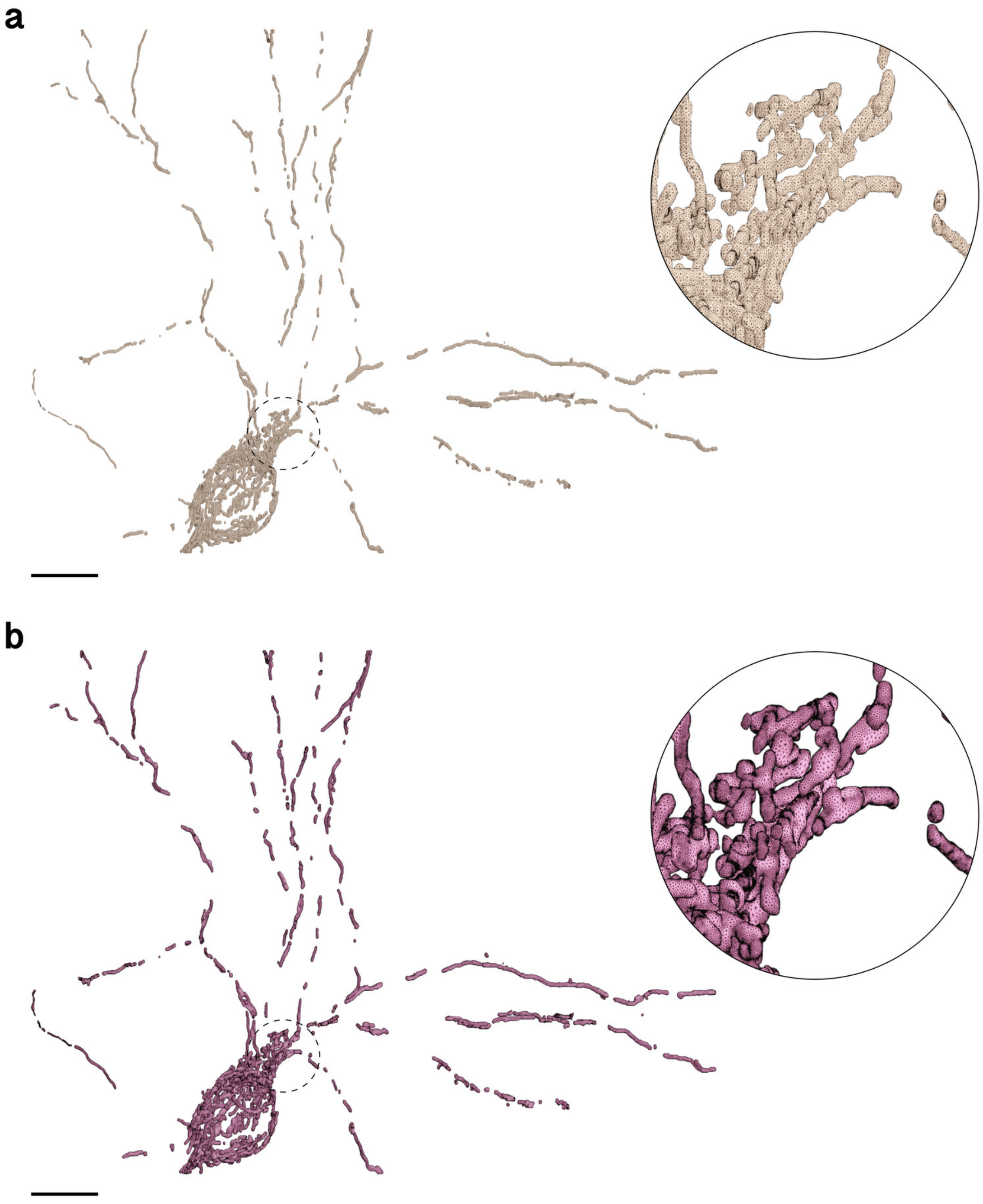

Figure S35: Comparative visual analysis between input (a) and reconstructed (b) meshes of of the mitochondria of Neuron 2. Scale bars, 10 $\mu$m (**a, b**).

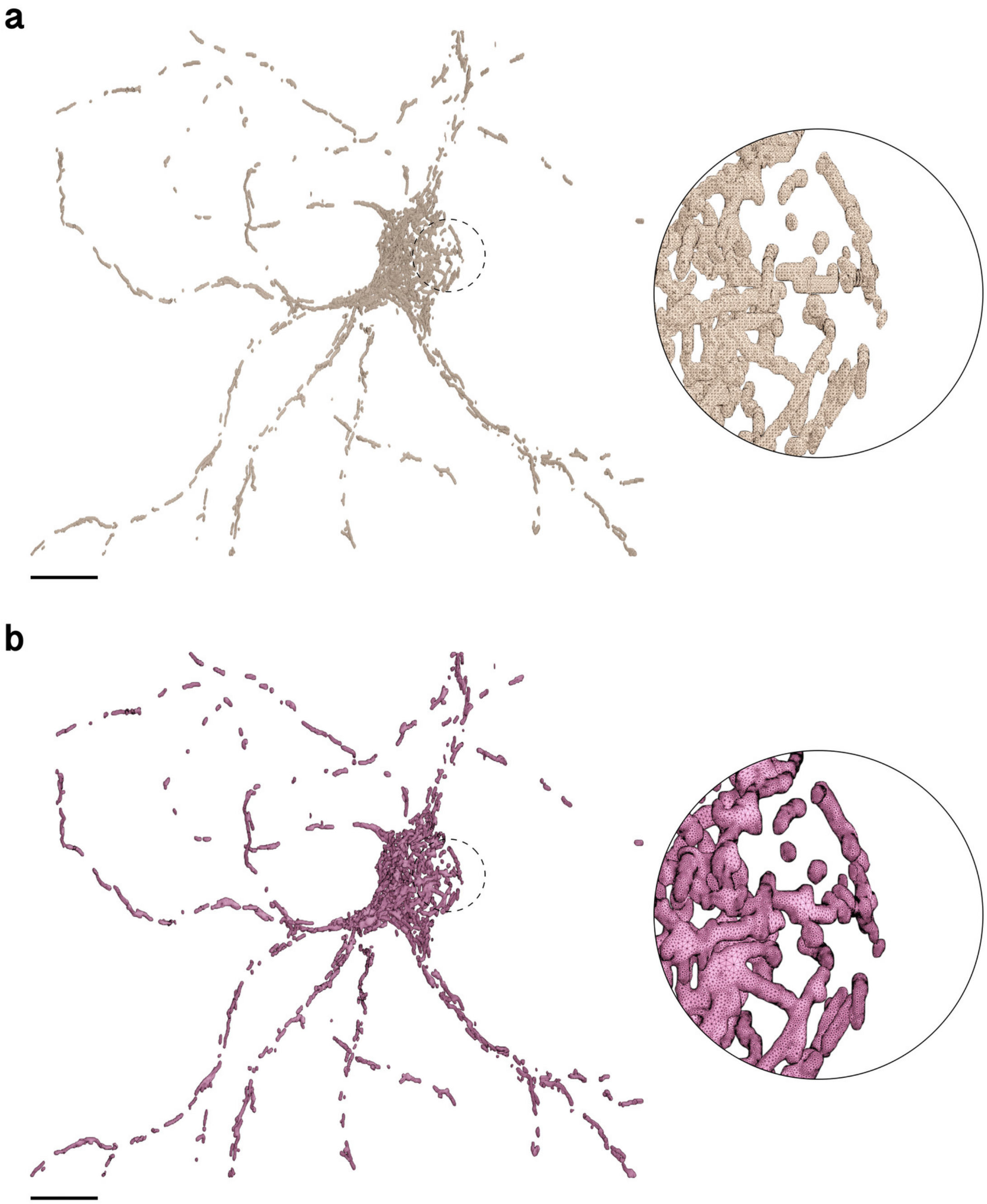

Figure S36: Comparative visual analysis between input (a) and reconstructed (b) meshes of of the mitochondria of Neuron 3. Scale bars, 10 $\mu$m (**a, b**).

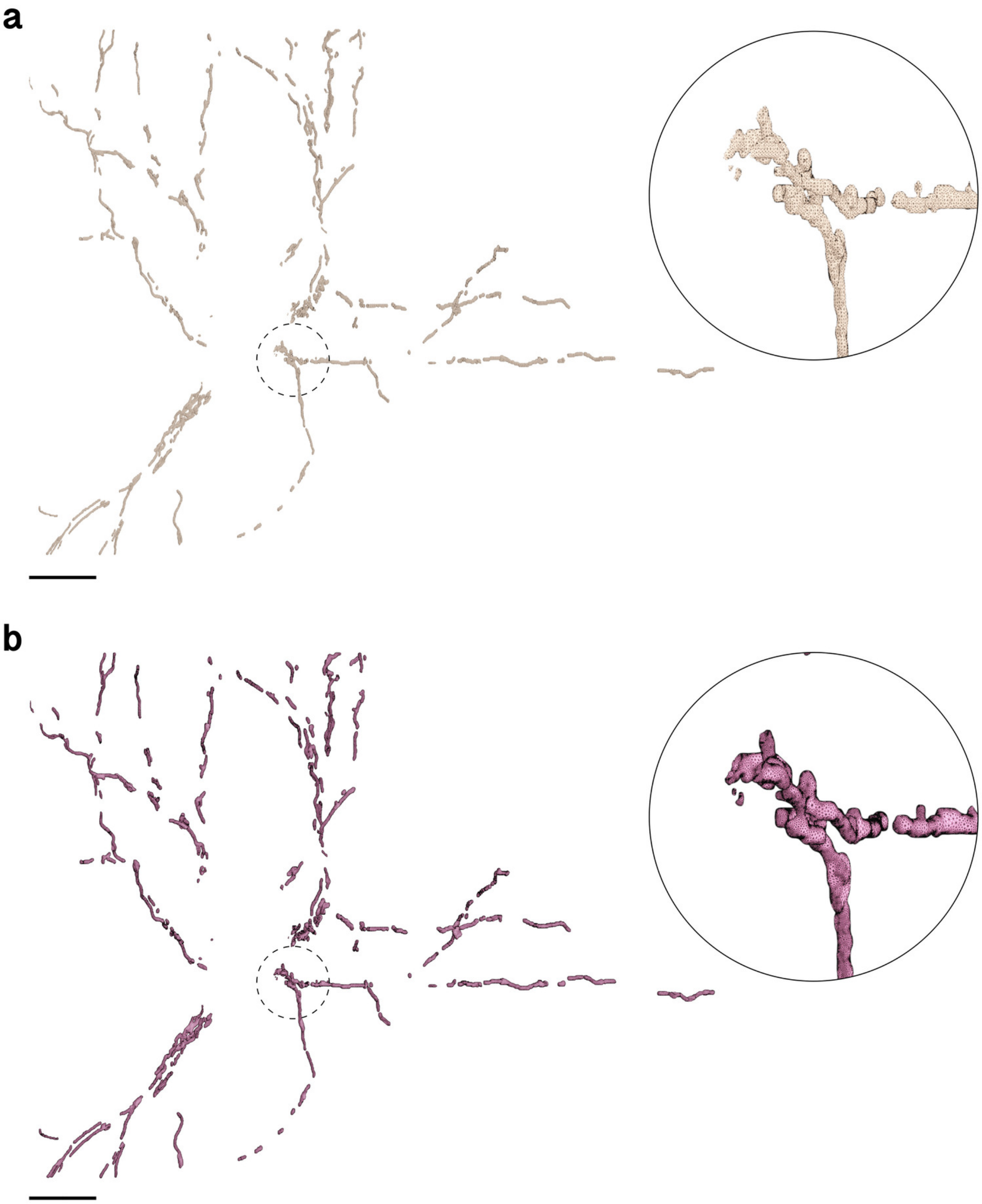

Figure S37: Comparative visual analysis between input (a) and reconstructed (b) meshes of of the mitochondria of Neuron 4. Scale bars, 10 $\mu$m (**a, b**).

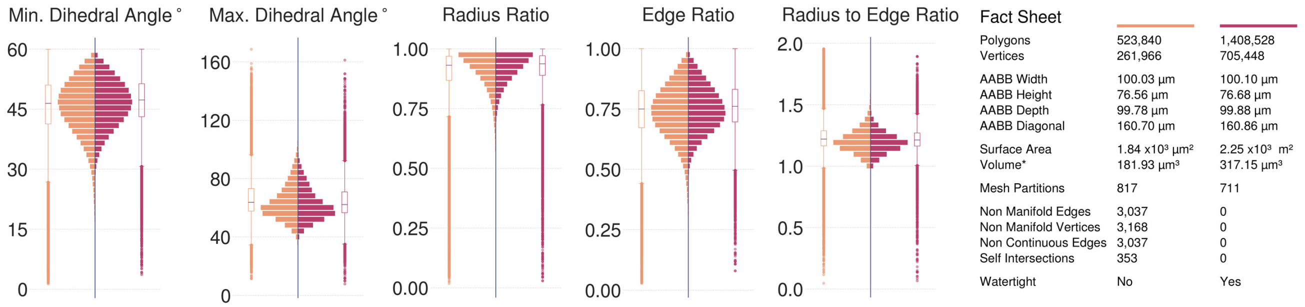

(a) Neuron 1 Mitochondria

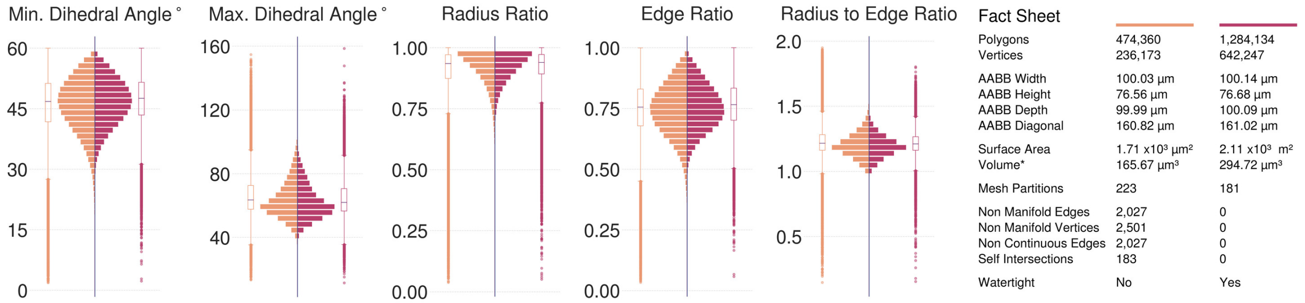

(b) Neuron 2 Mitochondria

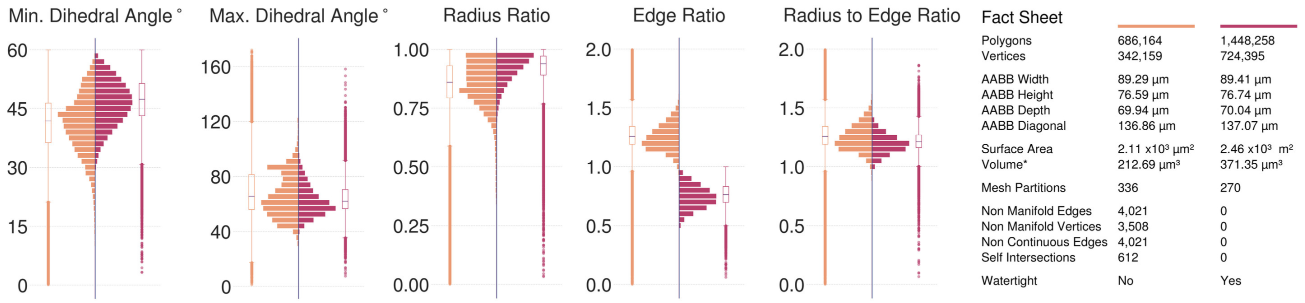

(c) Neuron 3 Mitochondria

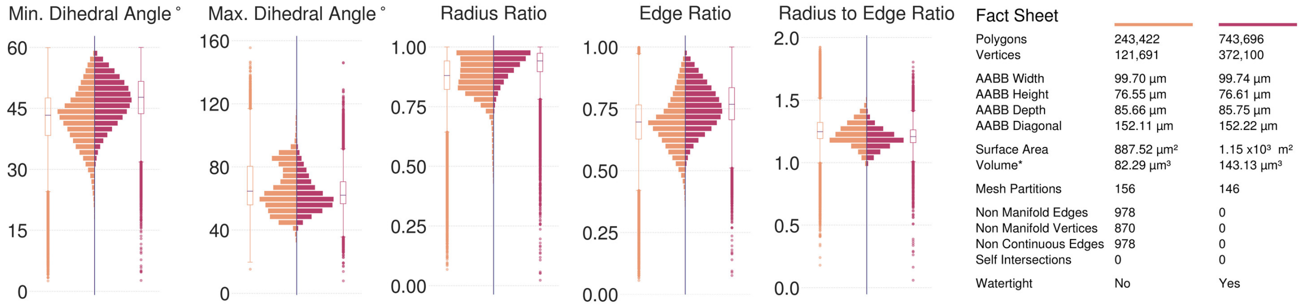

(d) Neuron 4 Mitochondria

Figure S38: Comparative analysis between input (left) and resulting (right) meshes of the mitochondria of the neurons segmented from the tissue block. Note that all the input meshes are not watertight. The resulting meshes are reconstructed with a voxelization resolution of 15 voxels per micron (66.7 nm).

#### 5.9 Microglia Mitochondria

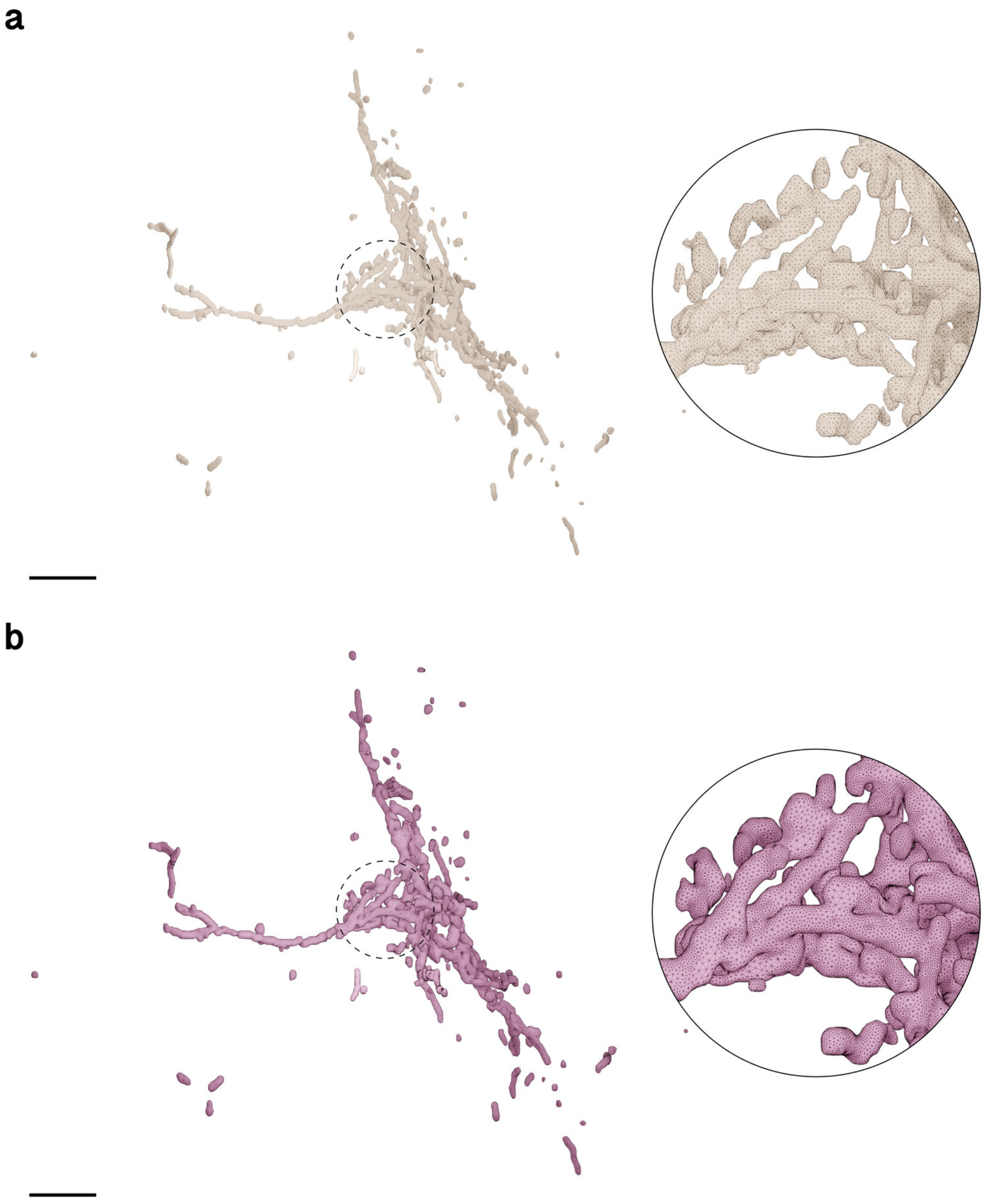

Figure S39: Comparative visual analysis between input (a) and reconstructed (b) meshes of of the mitochondria of Microglia 1. Scale bars, 5 $\mu$m (**a, b**).

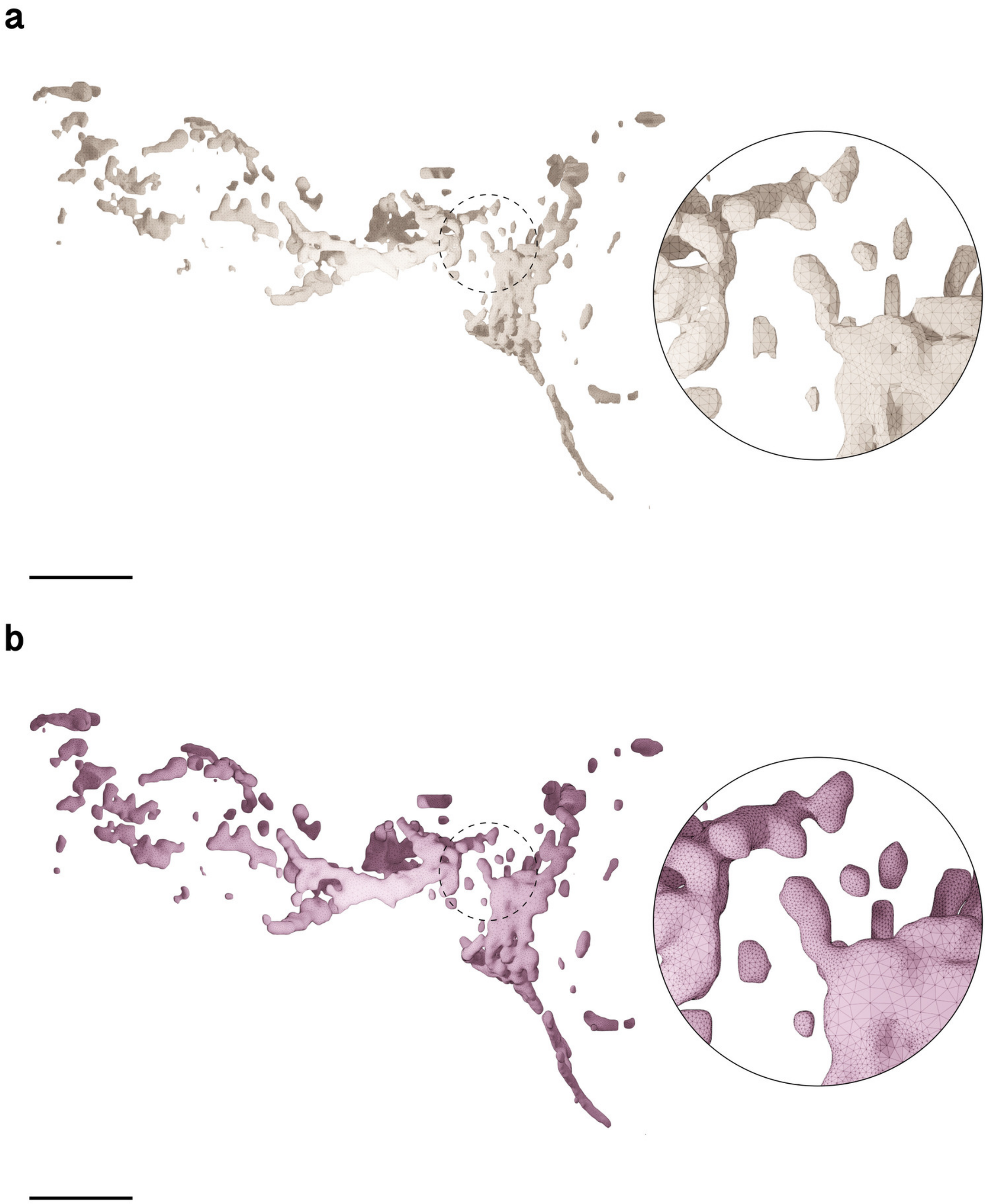

Figure S40: Comparative visual analysis between input (a) and reconstructed (b) meshes of of the mitochondria of Microglia 2. Scale bars, 5 $\mu$m (**a, b**).

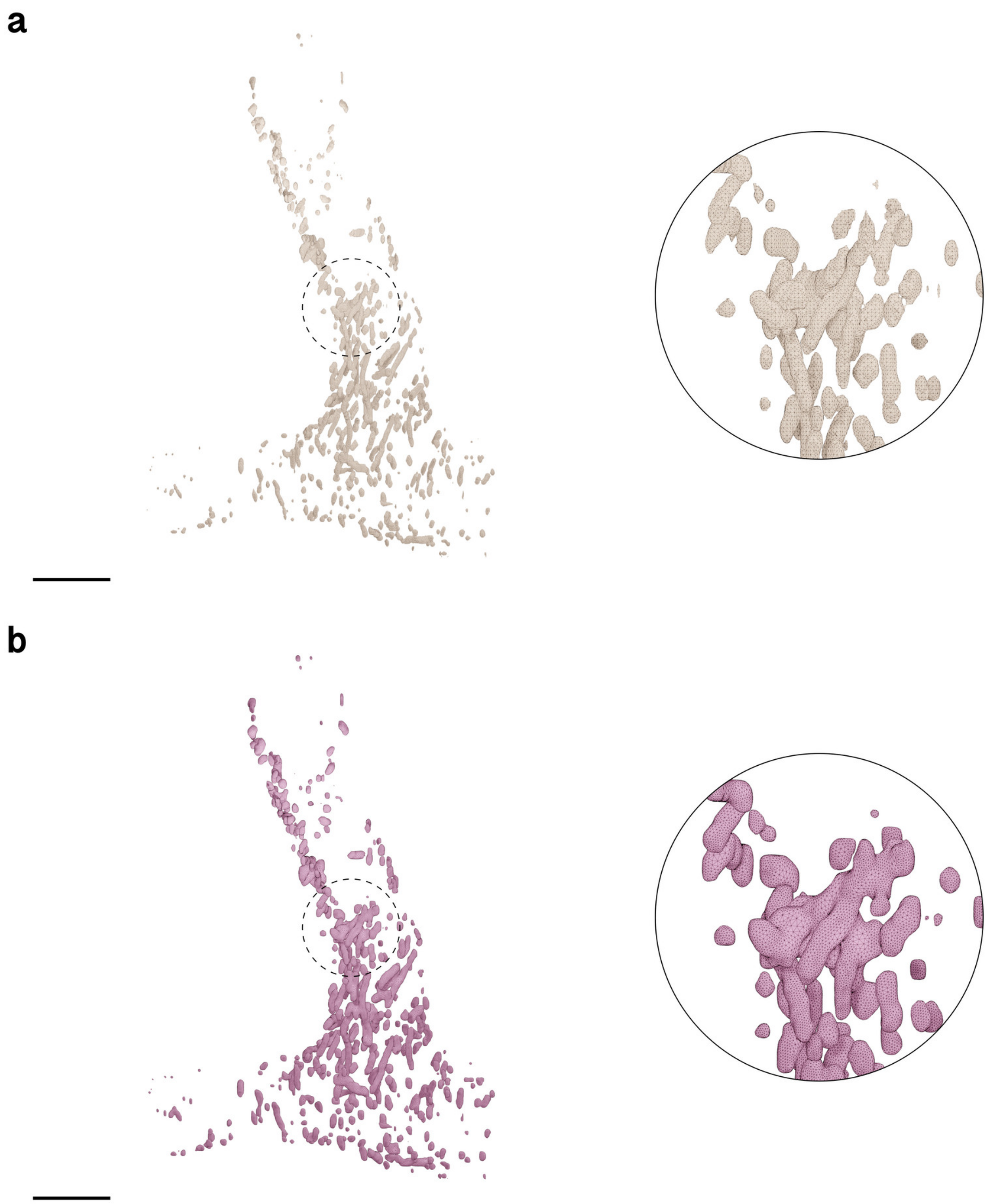

Figure S41: Comparative visual analysis between input (a) and reconstructed (b) meshes of of the mitochondria of Microglia 3. Scale bars, 5 $\mu$m (**a, b**).

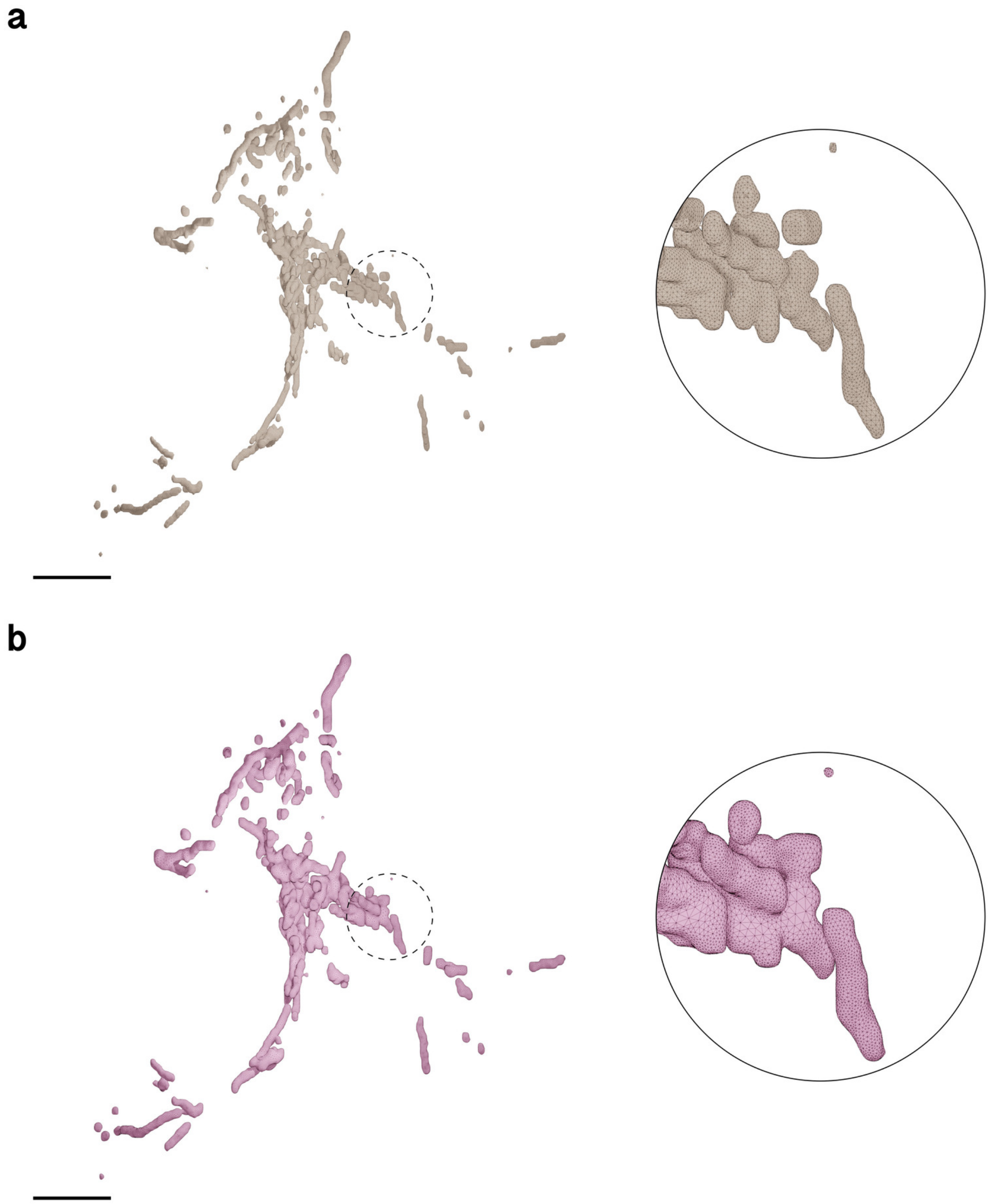

Figure S42: Comparative visual analysis between input (a) and reconstructed (b) meshes of of the mitochondria of Microglia 4. Scale bars, 5 $\mu$m (**a, b**).

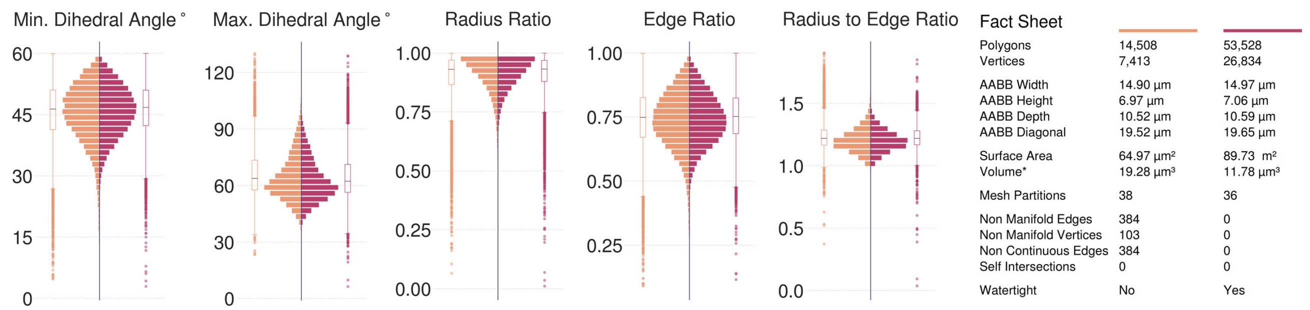

(**a**) Microglia 1 Mitochondria, Visualization in Figure S39.

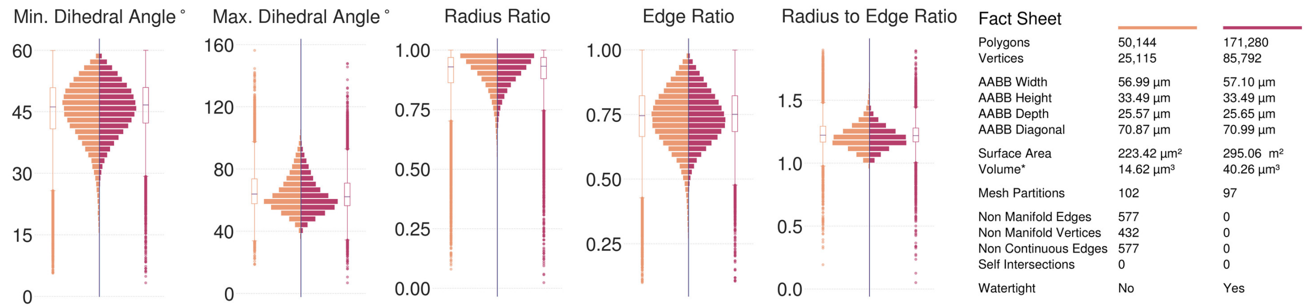

(**b**) Microglia 2 Mitochondria, Visualization in Figure S40.

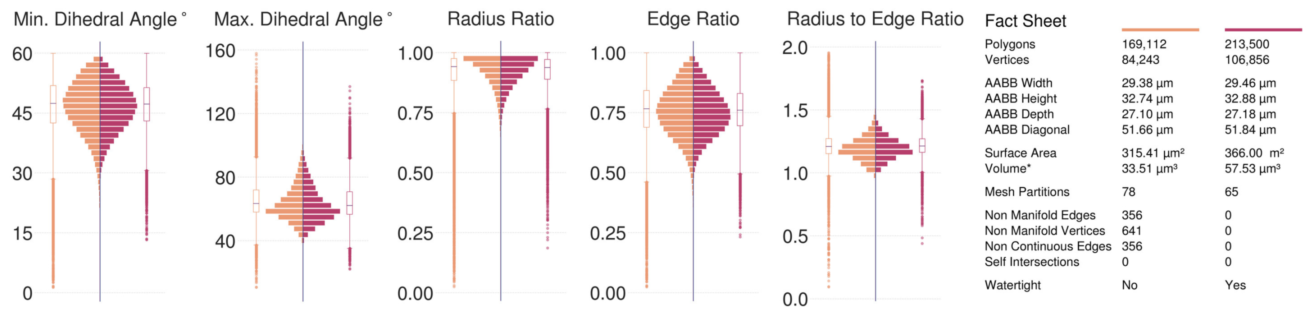

(**c**) Microglia 3 Mitochondria, Visualization in Figure S41.

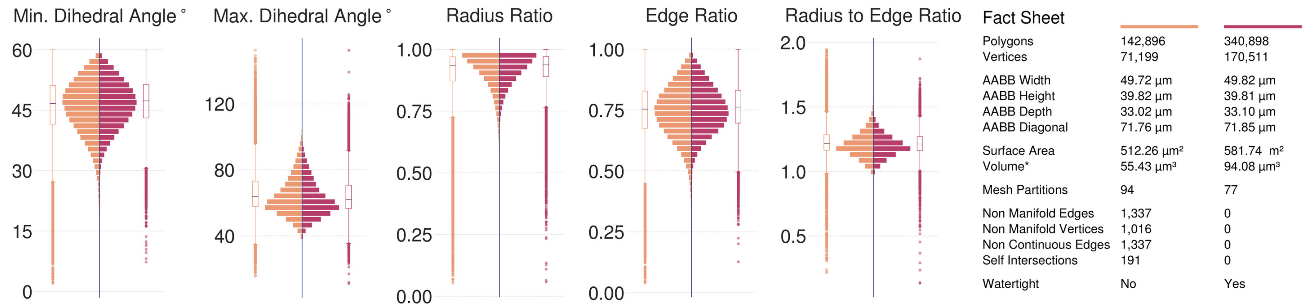

(**d**) Microglia 4 Mitochondria, Visualization in Figure S42.

Figure S43: Comparative analysis between input (left in orange) and resulting (right in dark salmon) meshes of the mitochondira of the four pericytes segmented from the volume shown in Figure S3. All the input meshes are not watertight; they have self-intersections and non-manifold edges and vertices. The resulting meshes are reconstructed at 15 voxels per micron resolution (67 nm) yielding multi-partitioned, adaptively optimized and watertight manifolds with no artifacts.

#### 5.10 Pericytes Mitochondria

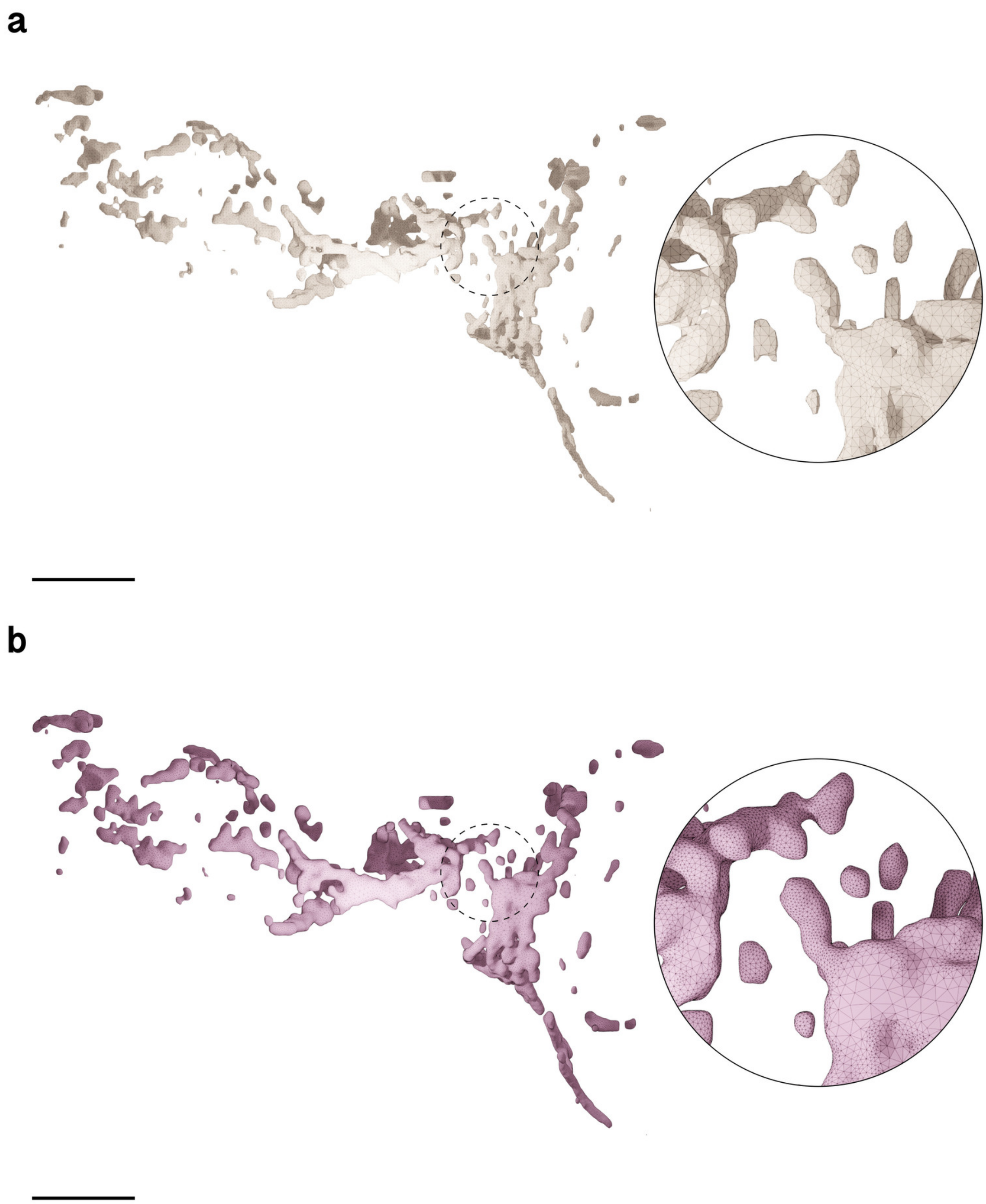

Figure S44: Wireframe visualizations comparing input (**a**) and resulting (**b**) surface meshes of the mitochondria of Pericyte 1. The closeups highlight the contrast in surface roughness, topology and tessellation. Comparative quantitative and qualitative analysis is shown in Figure S48a. Scale bars, 5 $\mu$m (**a, b**).

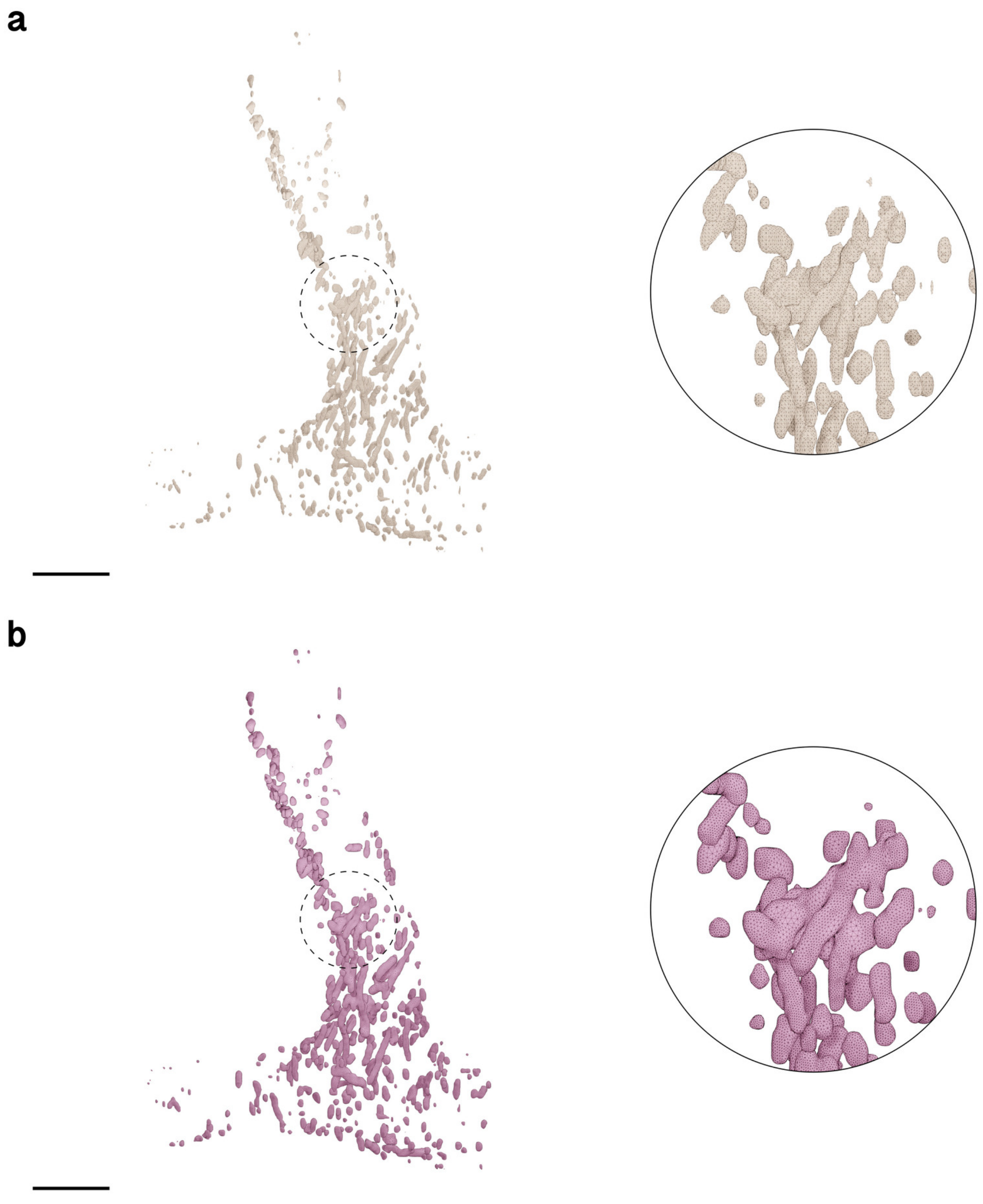

Figure S45: Wireframe visualizations comparing input (**a**) and resulting (**b**) surface meshes of the mitochondria of Pericyte 2. The closeups highlight the contrast in surface roughness, topology and tessellation. Comparative quantitative and qualitative analysis is shown in Figure S48b. Scale bars, 5 $\mu$m (**a, b**).

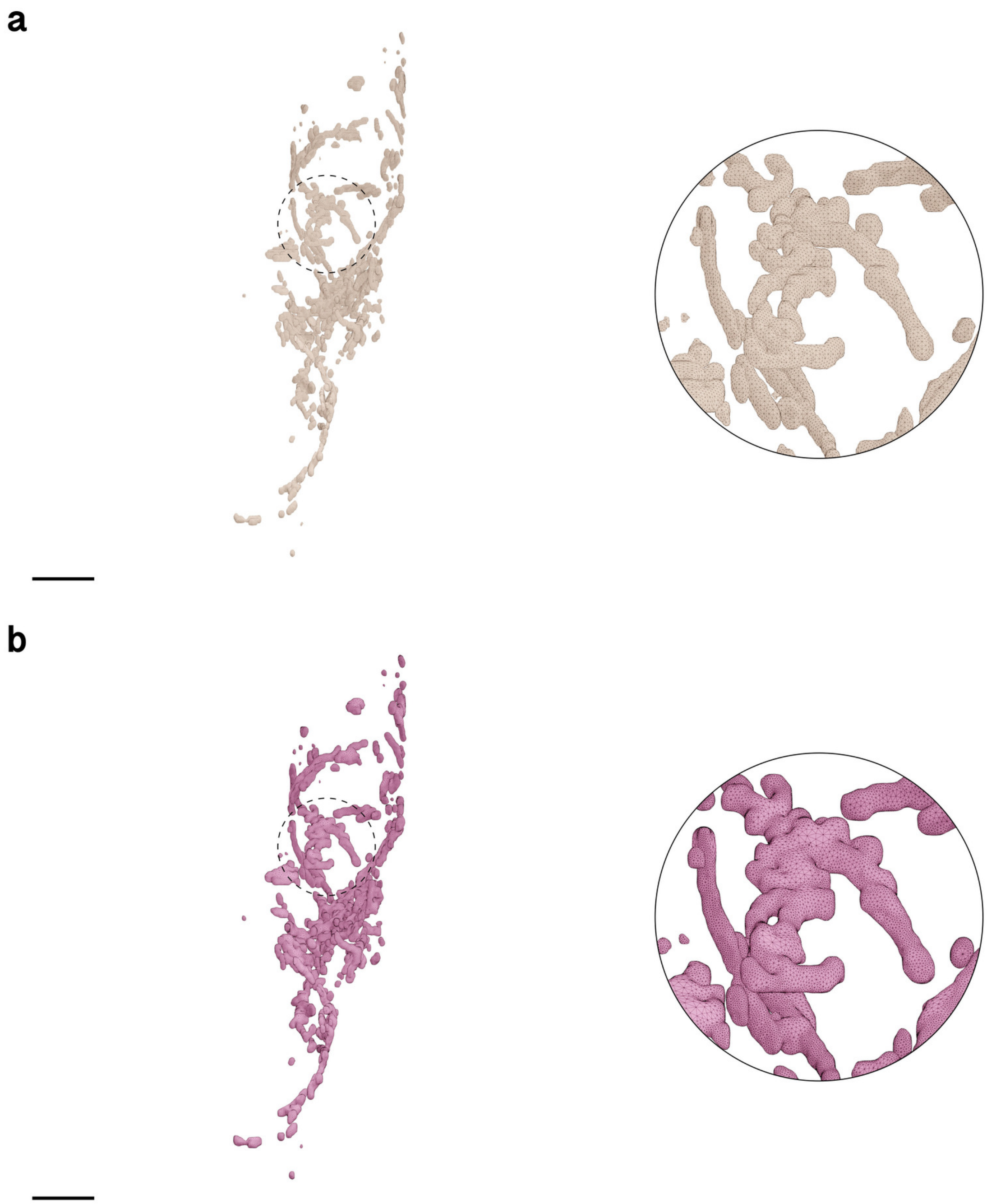

Figure S46: Wireframe visualizations comparing input (**a**) and resulting (**b**) surface meshes of the mitochondria of Pericyte 3. The closeups highlight the contrast in surface roughness, topology and tessellation. Comparative quantitative and qualitative analysis is shown in Figure S48c. Scale bars, 5 $\mu$m (**a, b**).

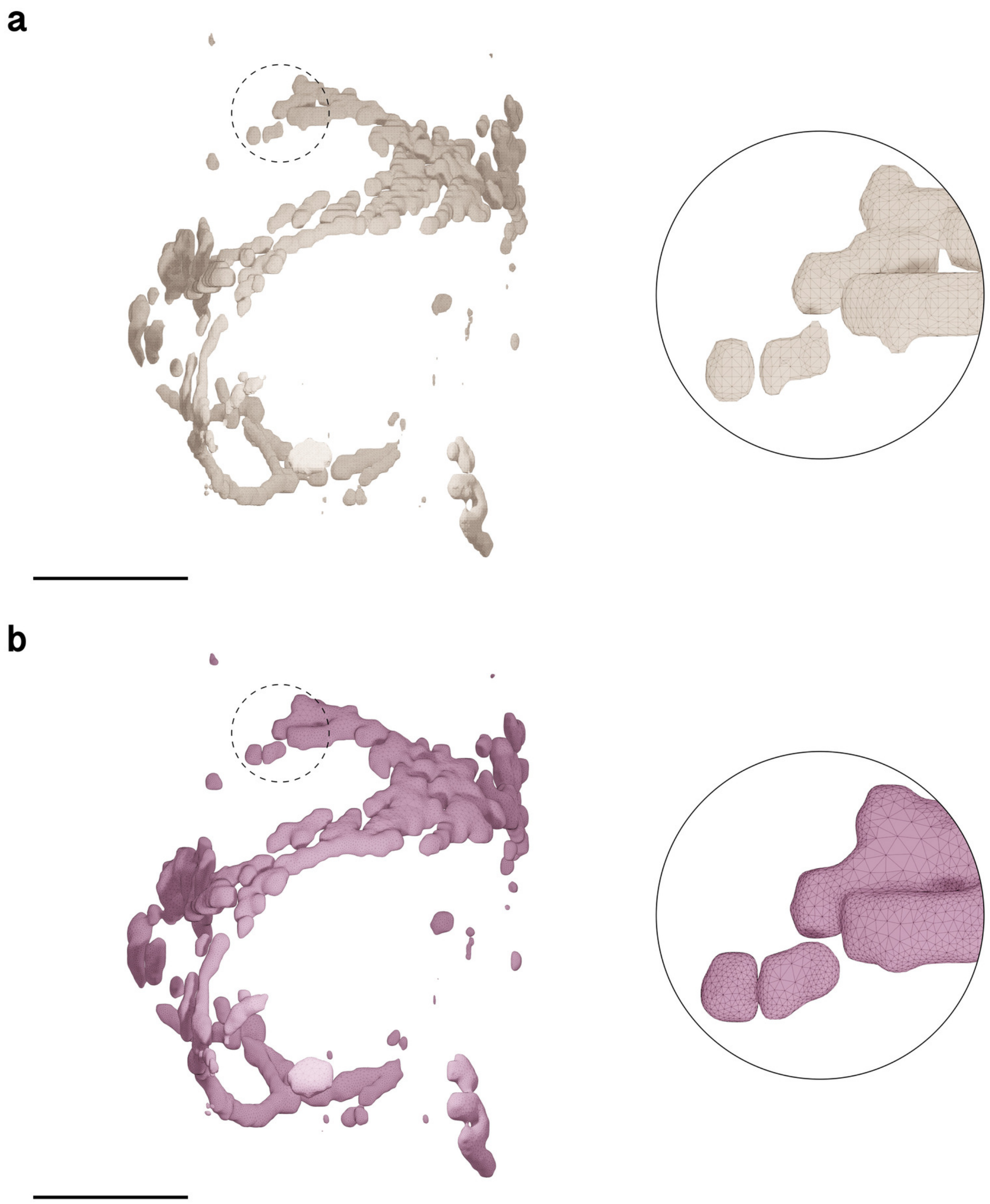

Figure S47: Wireframe visualizations comparing input (**a**) and resulting (**b**) surface meshes of the mitochondria of Pericyte 4. The closeups highlight the contrast in surface roughness, topology and tessellation. Comparative quantitative and qualitative analysis is shown in Figure S48d. Scale bars, 5 $\mu$m (**a, b**).

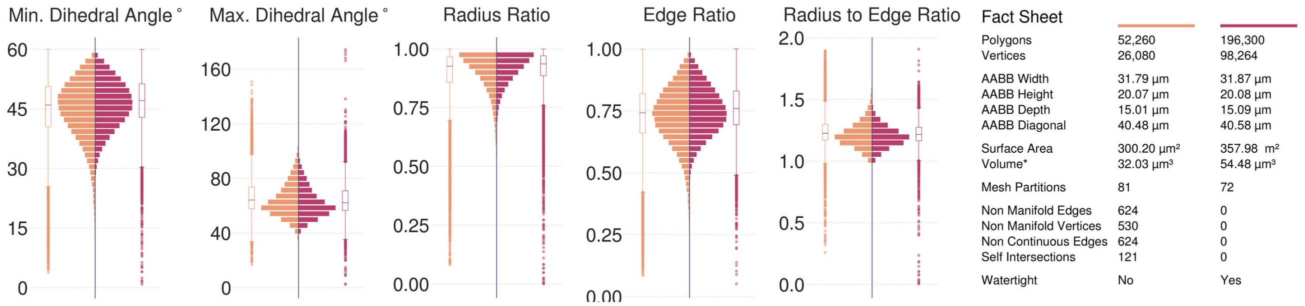

(**a**) Pericyte 1 Mitochondria, Visualization in Figure S44.

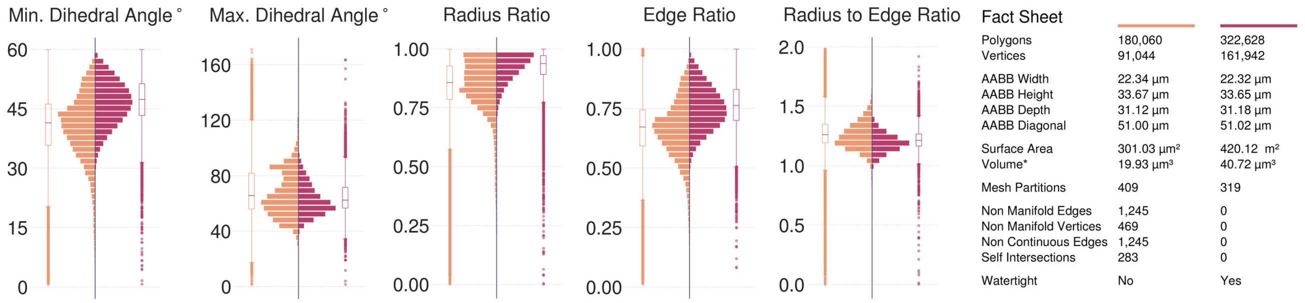

(**b**) Pericyte 2 Mitochondria, Visualization in Figure S45.

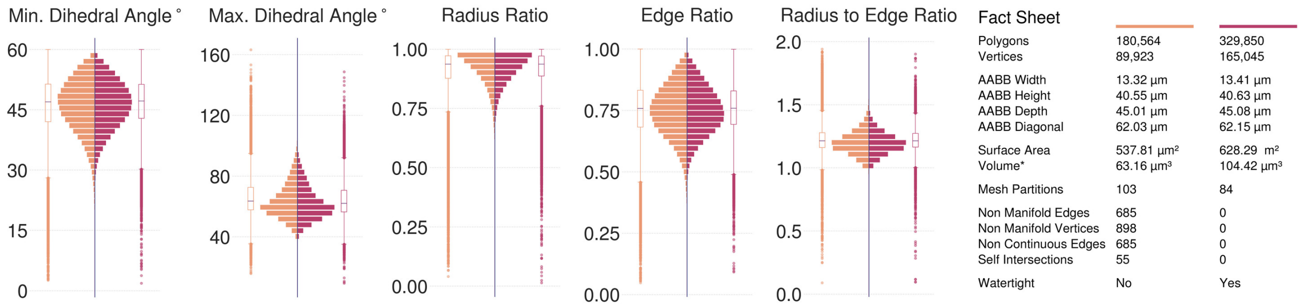

(**c**) Pericyte 3 Mitochondria, Visualization in Figure S46.

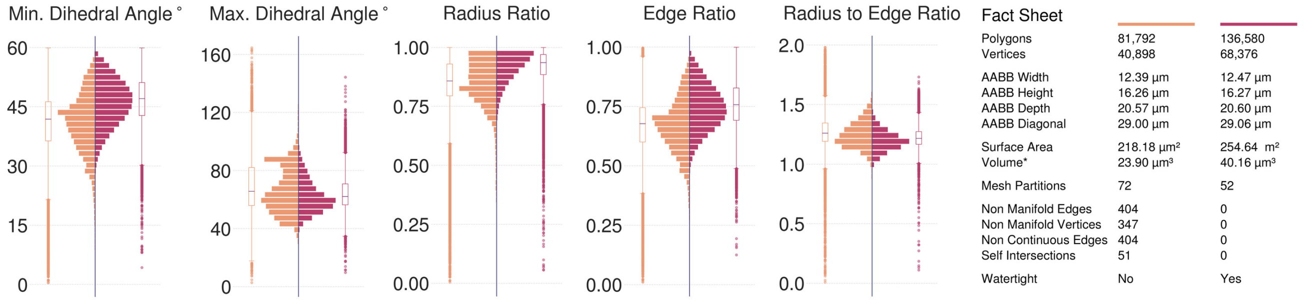

(**d**) Pericyte 4 Mitochondria, Visualization in Figure S47.

Figure S48: Comparative analysis between input (left in orange) and resulting (right in dark salmon) meshes of the mitochondira of the four pericytes segmented from the volume shown in Figure S3. All the input meshes are not watertight; they have self-intersections and non-manifold edges and vertices. The resulting meshes are reconstructed at 15 voxels per micron resolution (67 nm) yielding multi-partitioned, adaptively optimized and watertight manifolds with no artifacts.

#### 5.11 Astrocytes ER

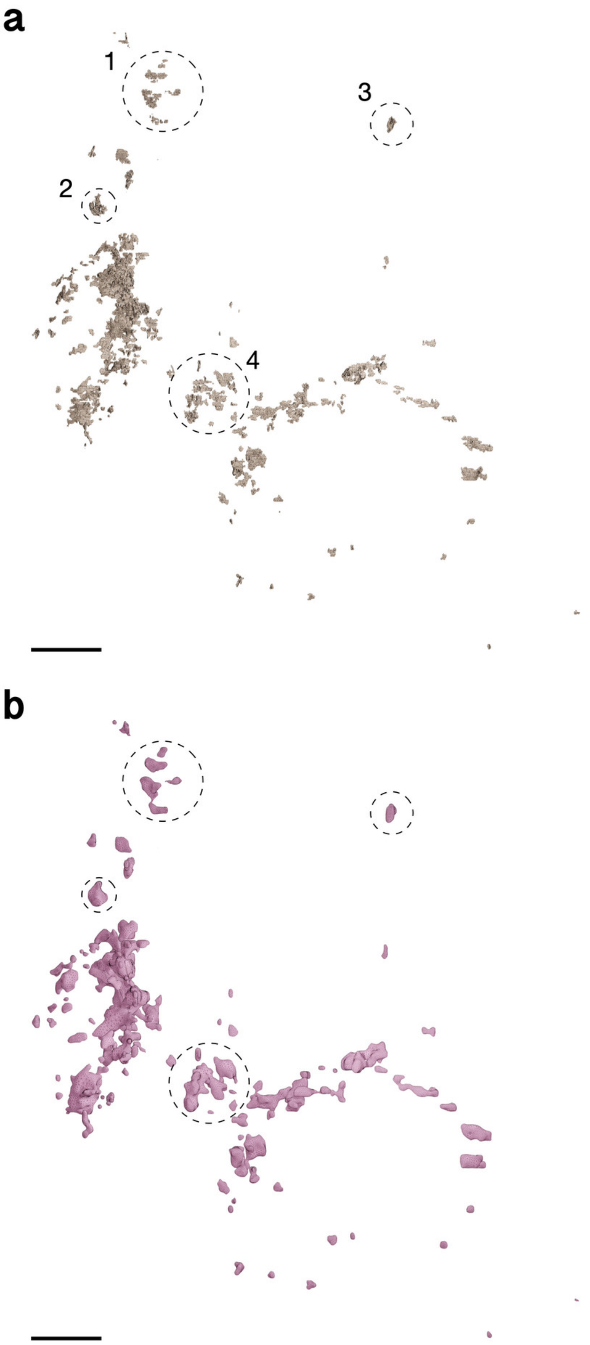

Figure S49: Wireframe visualizations comparing input (**a**) and resulting (**b**) surface meshes of the ER of Astrocyte 1. The closeups highlight the contrast in surface roughness, topology and tessellation. Comparative quantitative and qualitative analysis is shown in Figure S53a. Scale bars, 10 $\mu$m (**a, b**).

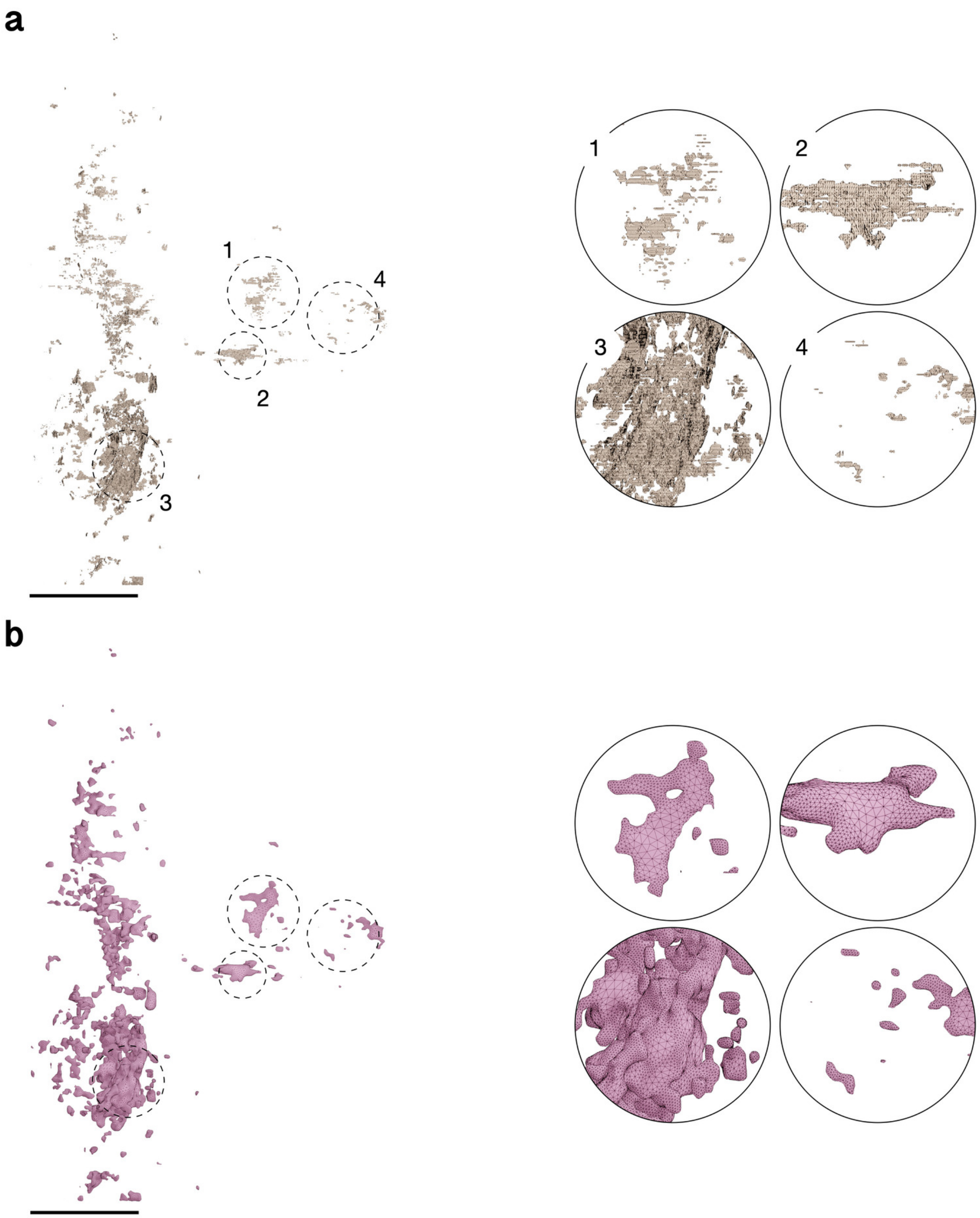

Figure S50: Wireframe visualizations comparing input (**a**) and resulting (**b**) surface meshes of the ER of Astrocyte 2. The closeups highlight the contrast in surface roughness, topology and tessellation. Comparative quantitative and qualitative analysis is shown in Figure S53b. Scale bars, 10 $\mu$m (**a, b**).

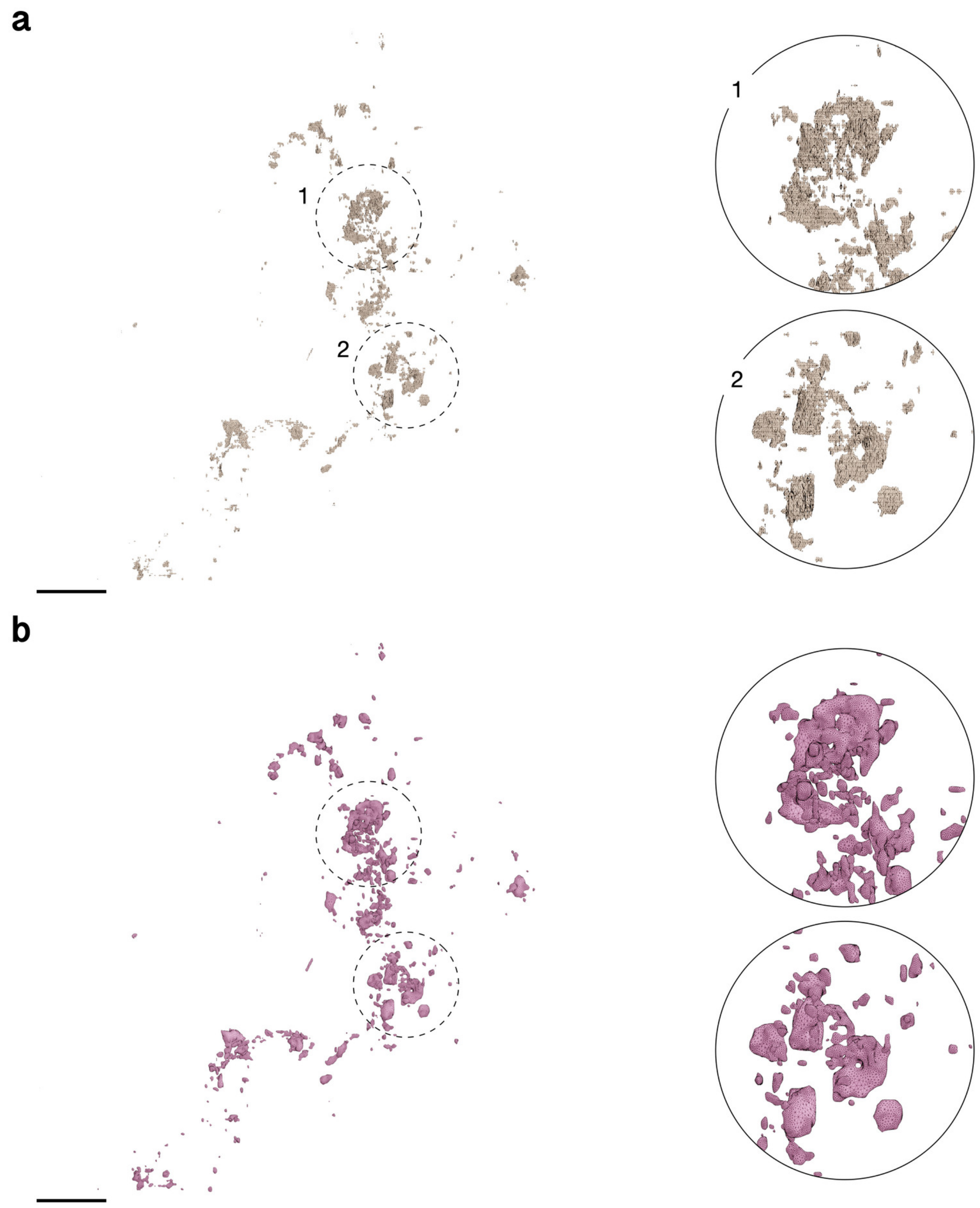

Figure S51: Wireframe visualizations comparing input (**a**) and resulting (**b**) surface meshes of the ER of Astrocyte 3. The closeups highlight the contrast in surface roughness, topology and tessellation. Comparative quantitative and qualitative analysis is shown in Figure S53c. Scale bars, 10 $\mu$m (**a, b**).

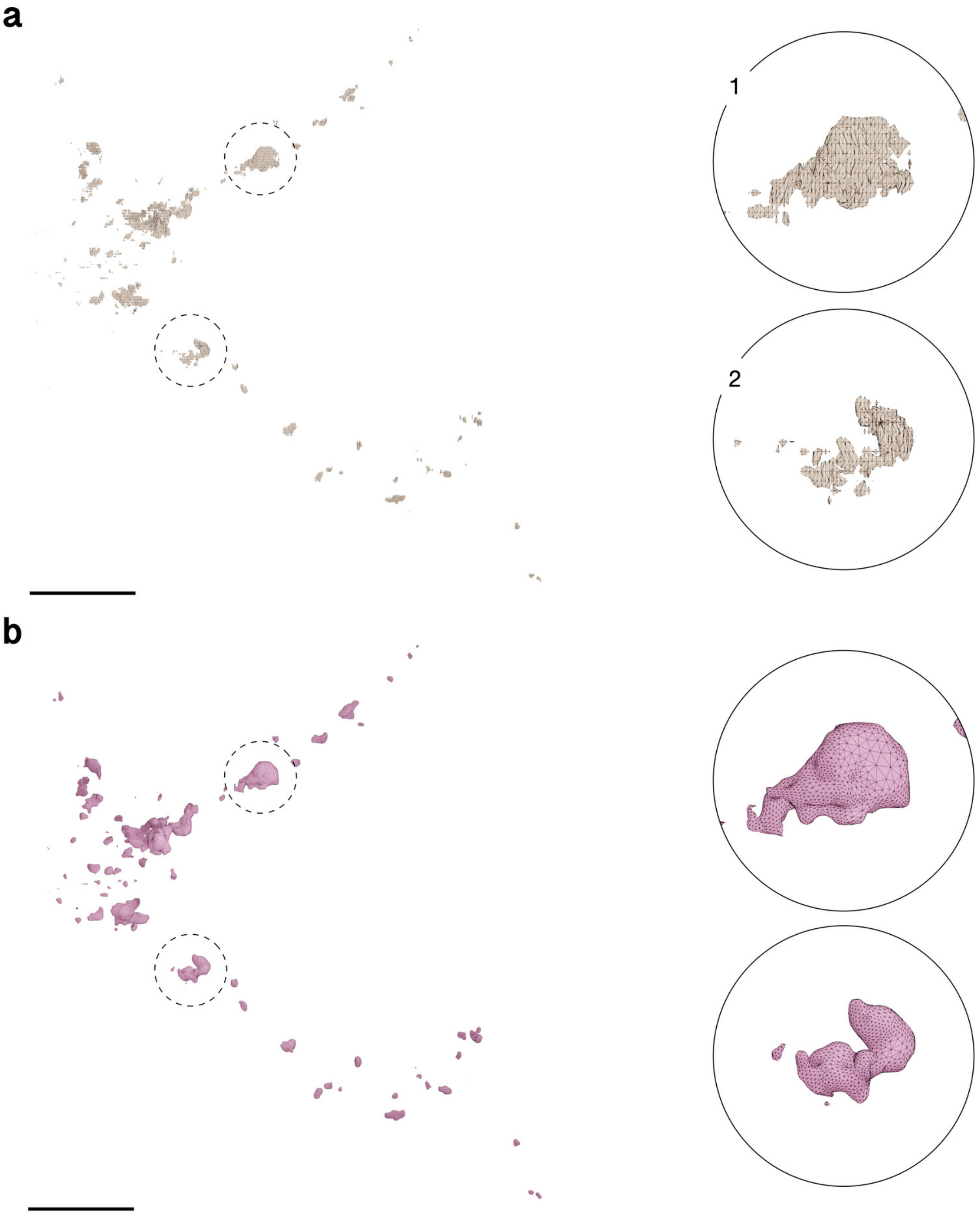

Figure S52: Wireframe visualizations comparing input (**a**) and resulting (**b**) surface meshes of the ER of Astrocyte 4. The closeups highlight the contrast in surface roughness, topology and tessellation. Comparative quantitative and qualitative analysis is shown in Figure S53d. Scale bars, 10 $\mu$m (**a, b**).

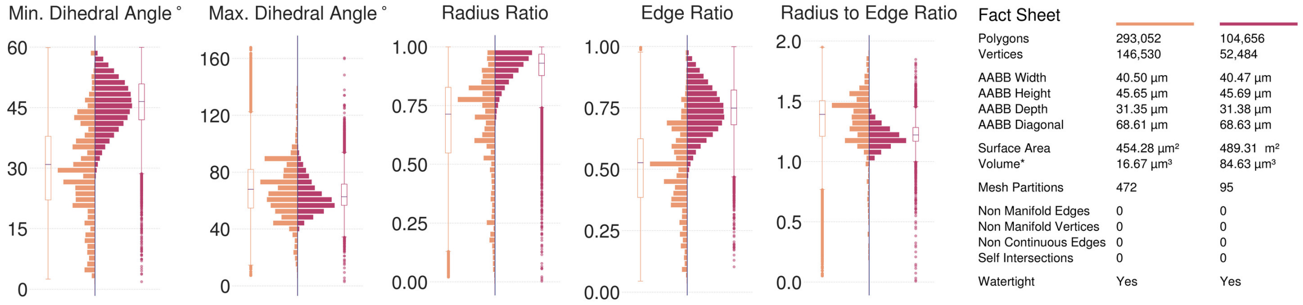

(**a**) Astrocyte 1 ER, Visualization in Figure S49.

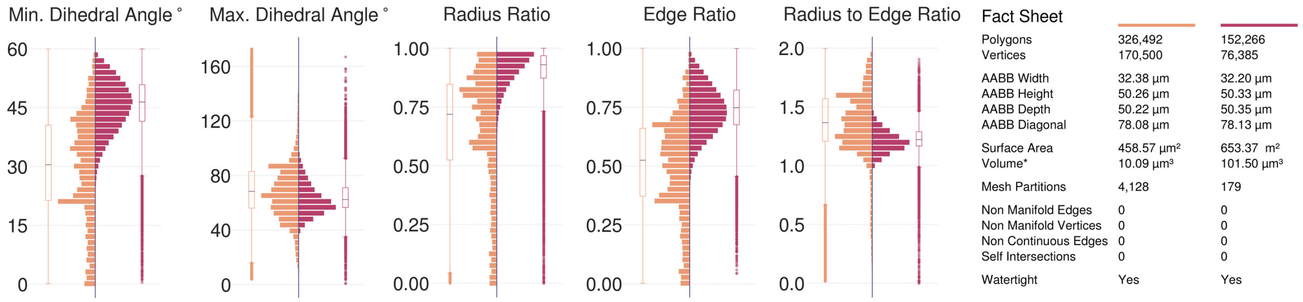

(**b**) Astrocyte 2 ER, Visualization in Figure S50.

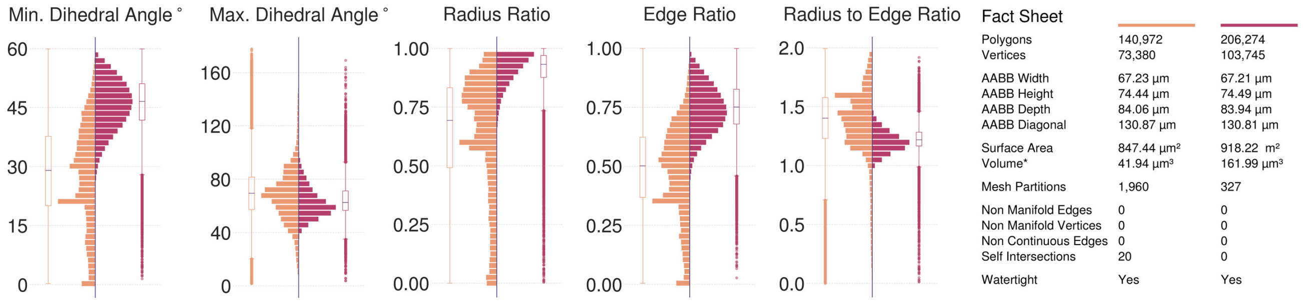

(**c**) Astrocyte 3 ER, Visualization in Figure S51.

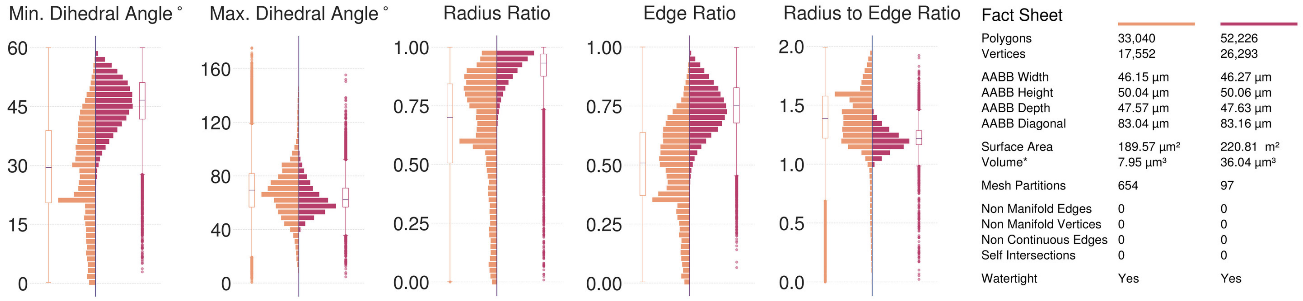

(**d**) Astrocyte 4 ER, Visualization in Figure S52.

Figure S53: Comparative analysis between input (left in orange) and resulting (right in dark salmon) meshes of the ER of the complete astrocytes segmented from the volume shown in Figure S3. Note that all the input meshes are watertight, but their qualitative geometric analysis is comparatively poor. The resulting meshes are reconstructed at 10 voxels per micron resolution (100 nm) yielding multi-partitioned and adaptively optimized manifolds with no artifacts.

### 6 Remeshing poorly segmented neuronal meshes with fragmented partitions and slicing artifacts

#### 6.1 Remeshing multi-partitioned meshes with poor topology

We have used ultraMesh2Mesh to remesh (and create optimized watertight meshes) 20 neuronal meshes segmented from layer II/III of the visual cortex of a P36 male mouse (Figure S54). These segmented meshes contain multiple partitions and low-quality over-tessellated topology with several non-manifold vertices and edges. A comparative analysis between the downloaded and resulting meshes is detailed in Figures S55–S74.

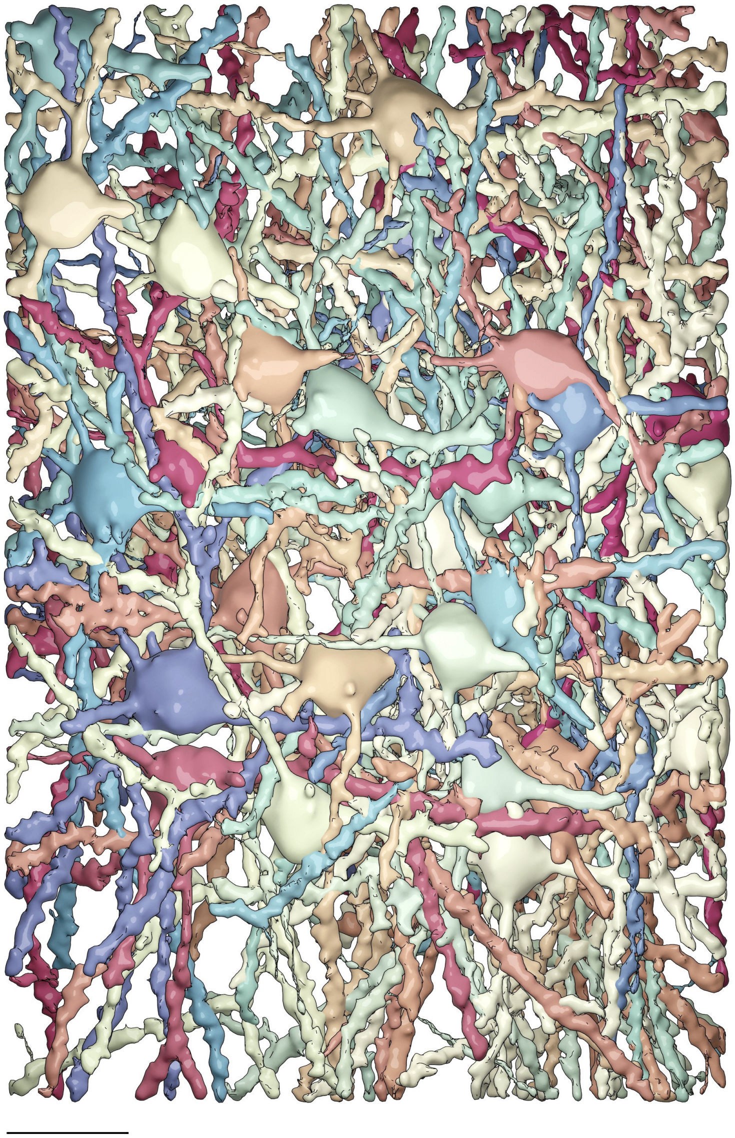

Figure S54: Rendering of a small block of cortical volume containing hundreds of pyramidal neurons that are segmented within the scope of the MICrONS program (microns-explorer.org). Scale bar, 50 micron.

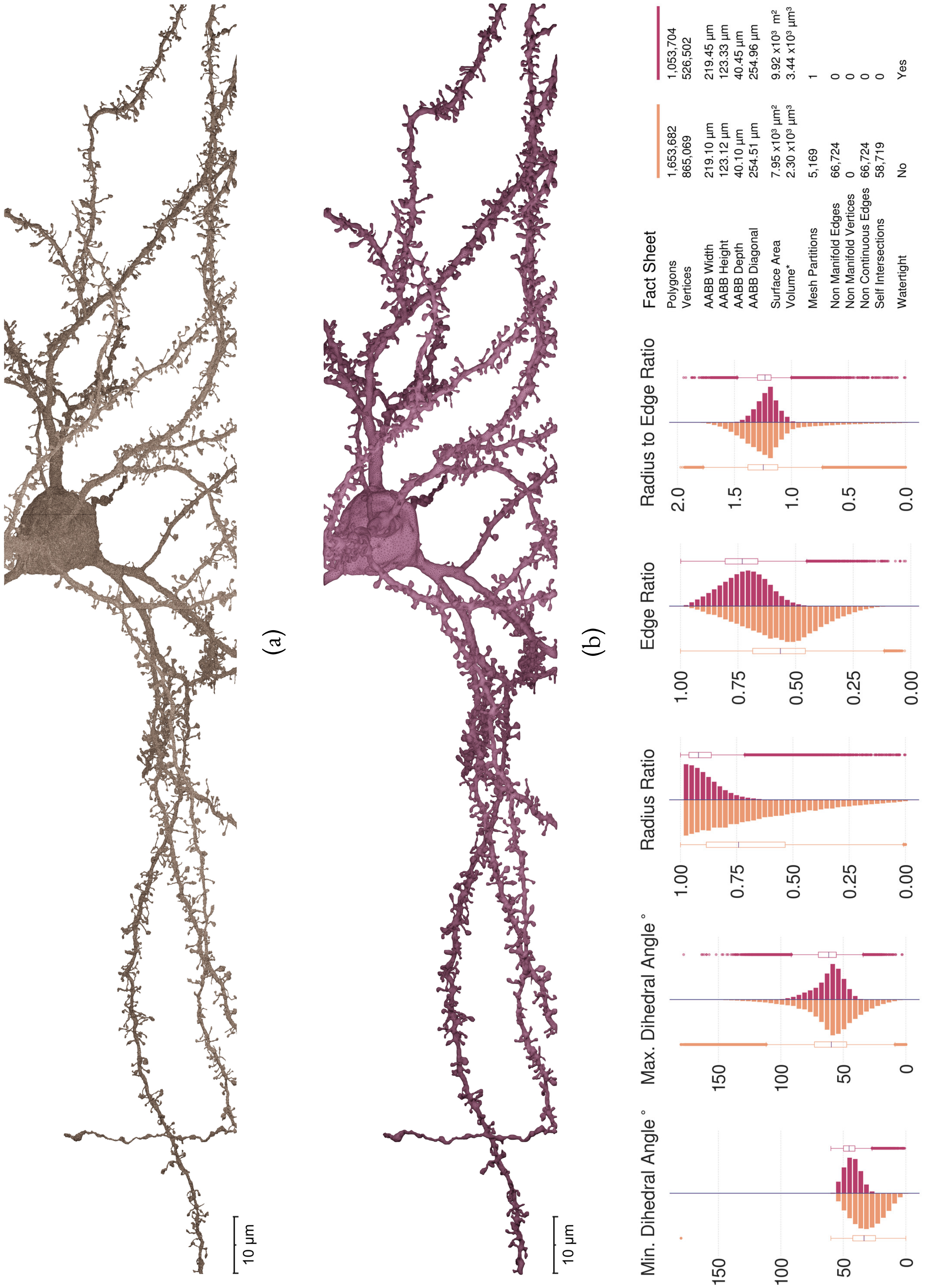

Figure S55: Comparative visual and quantitative analysis between input (in orange) and resulting (in salmon) meshes of Neuron 1.

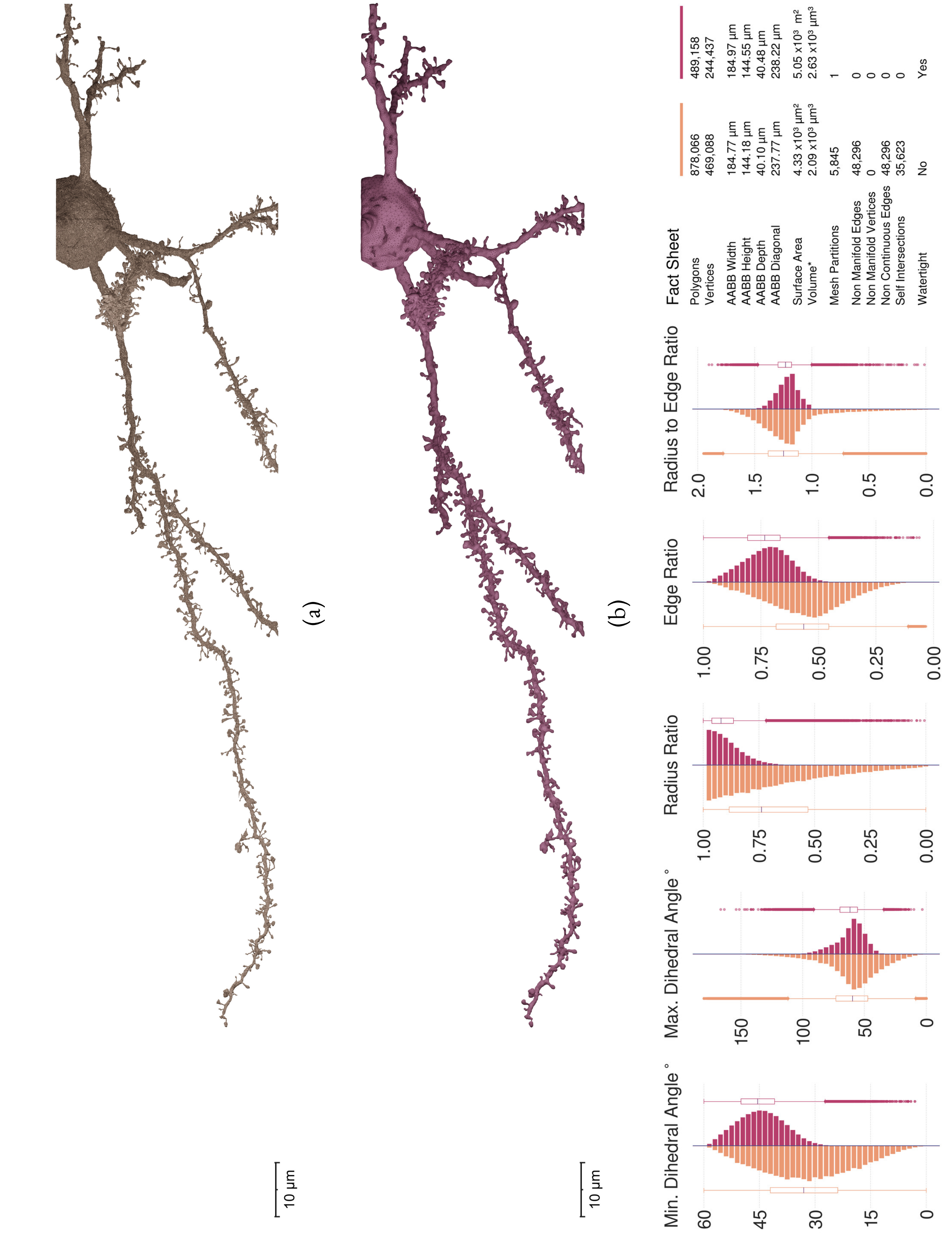

Figure S56: Comparative visual and quantitative analysis between input (in orange) and resulting (in salmon) meshes of Neuron 2.

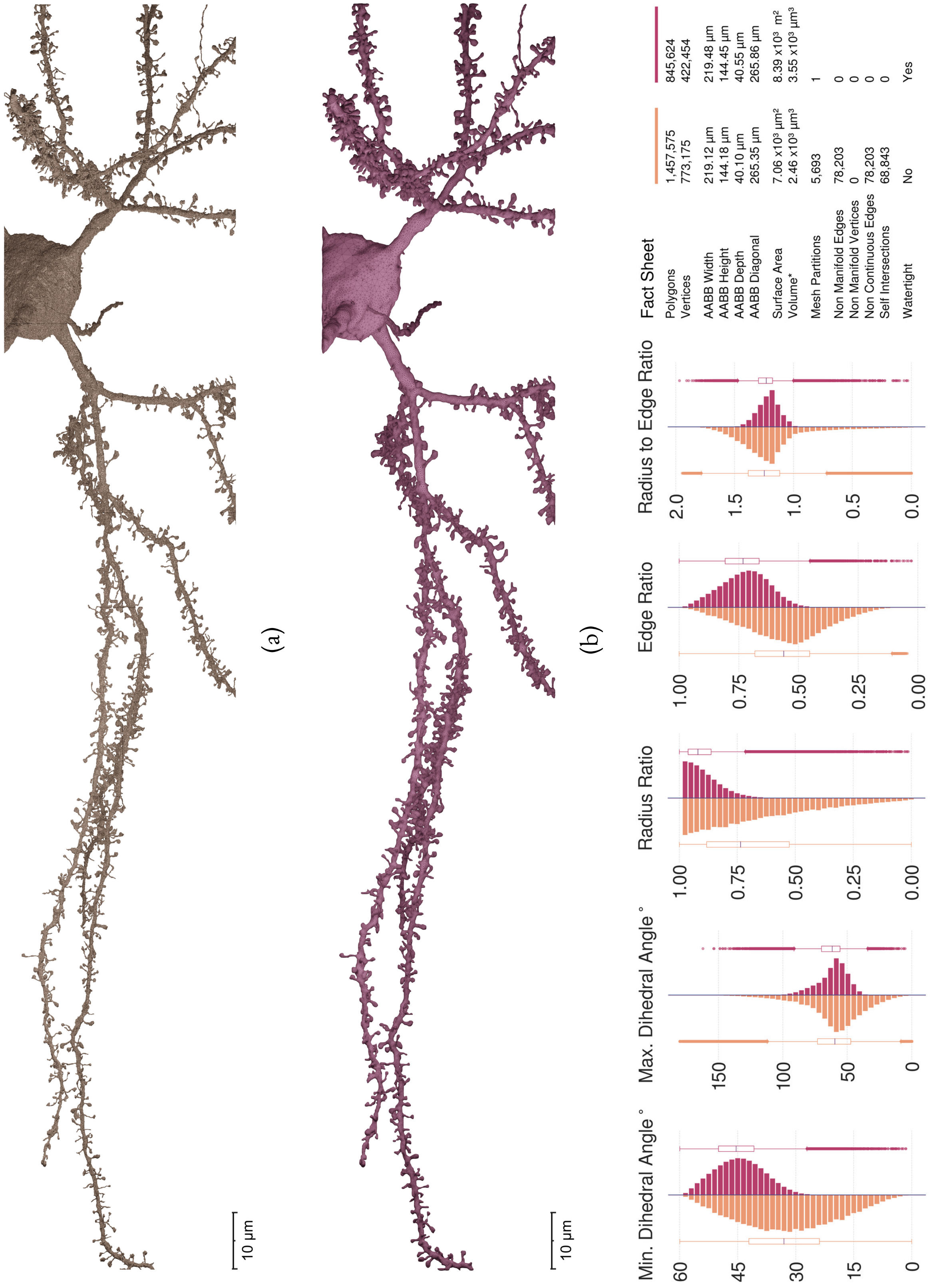

Figure S57: Comparative visual and quantitative analysis between input (in orange) and resulting (in salmon) meshes of Neuron 3.

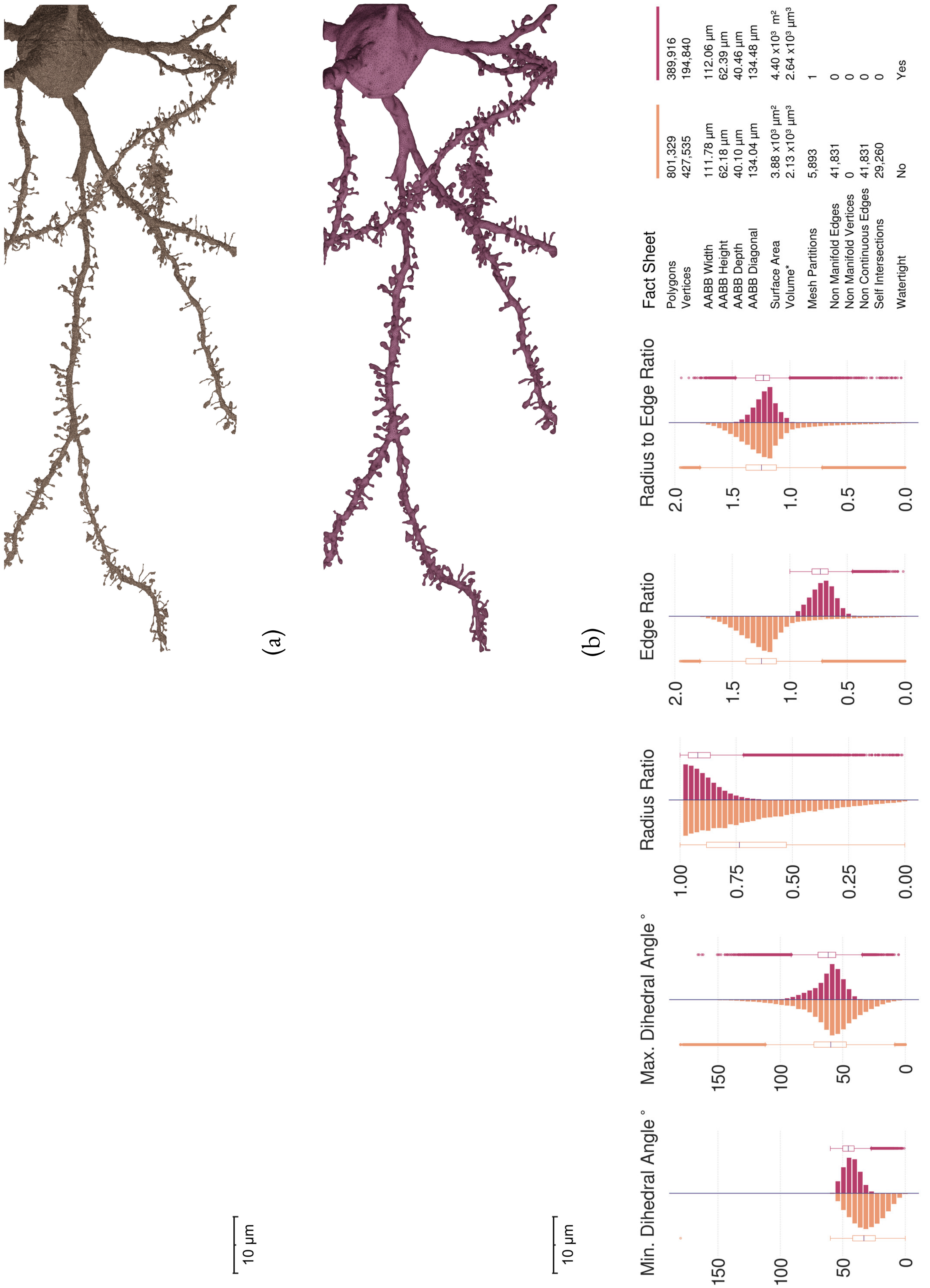

Figure S58: Comparative visual and quantitative analysis between input (in orange) and resulting (in salmon) meshes of Neuron 4.

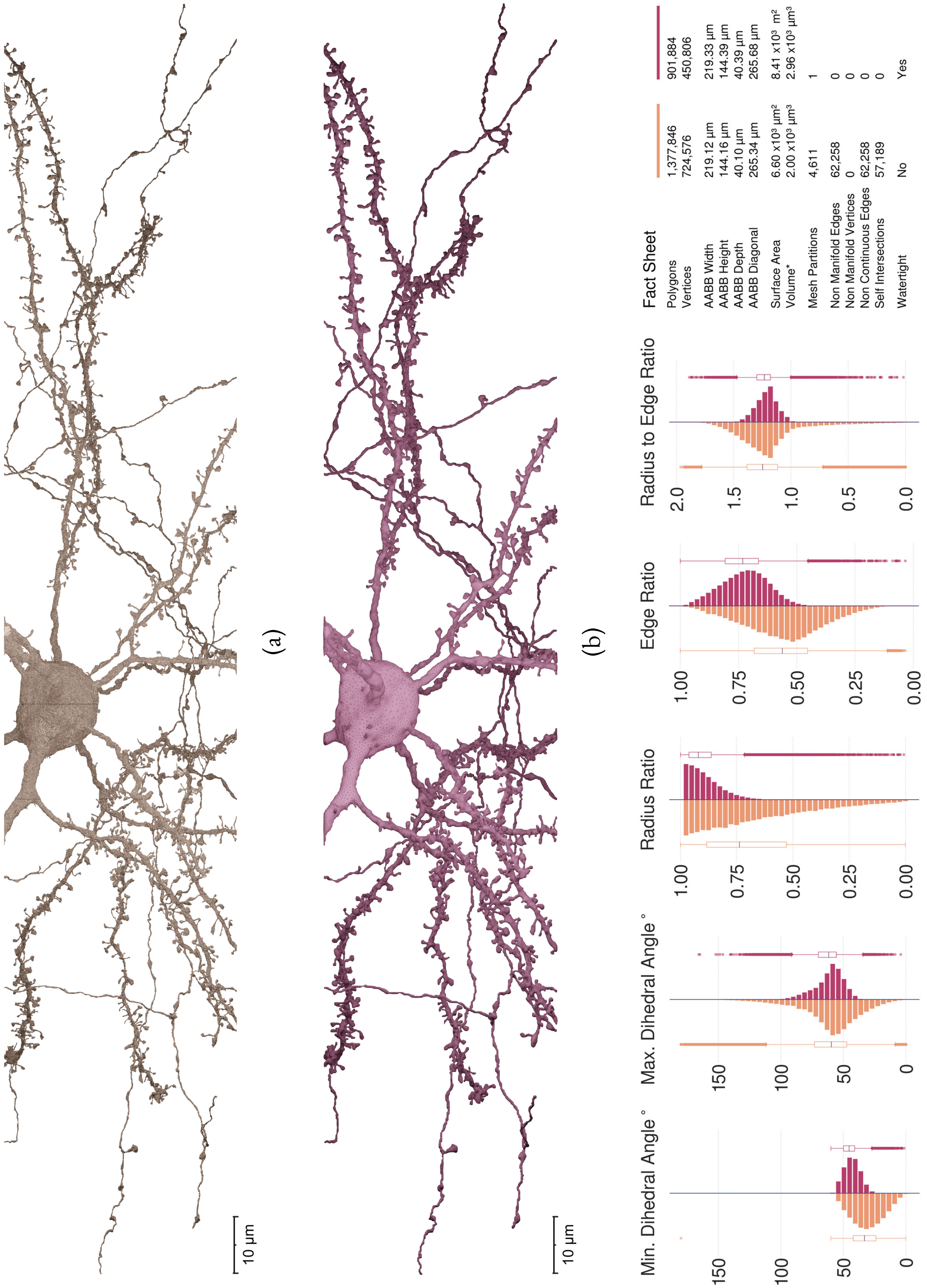

Figure S59: Comparative visual and quantitative analysis between input (in orange) and resulting (in salmon) meshes of Neuron 5.

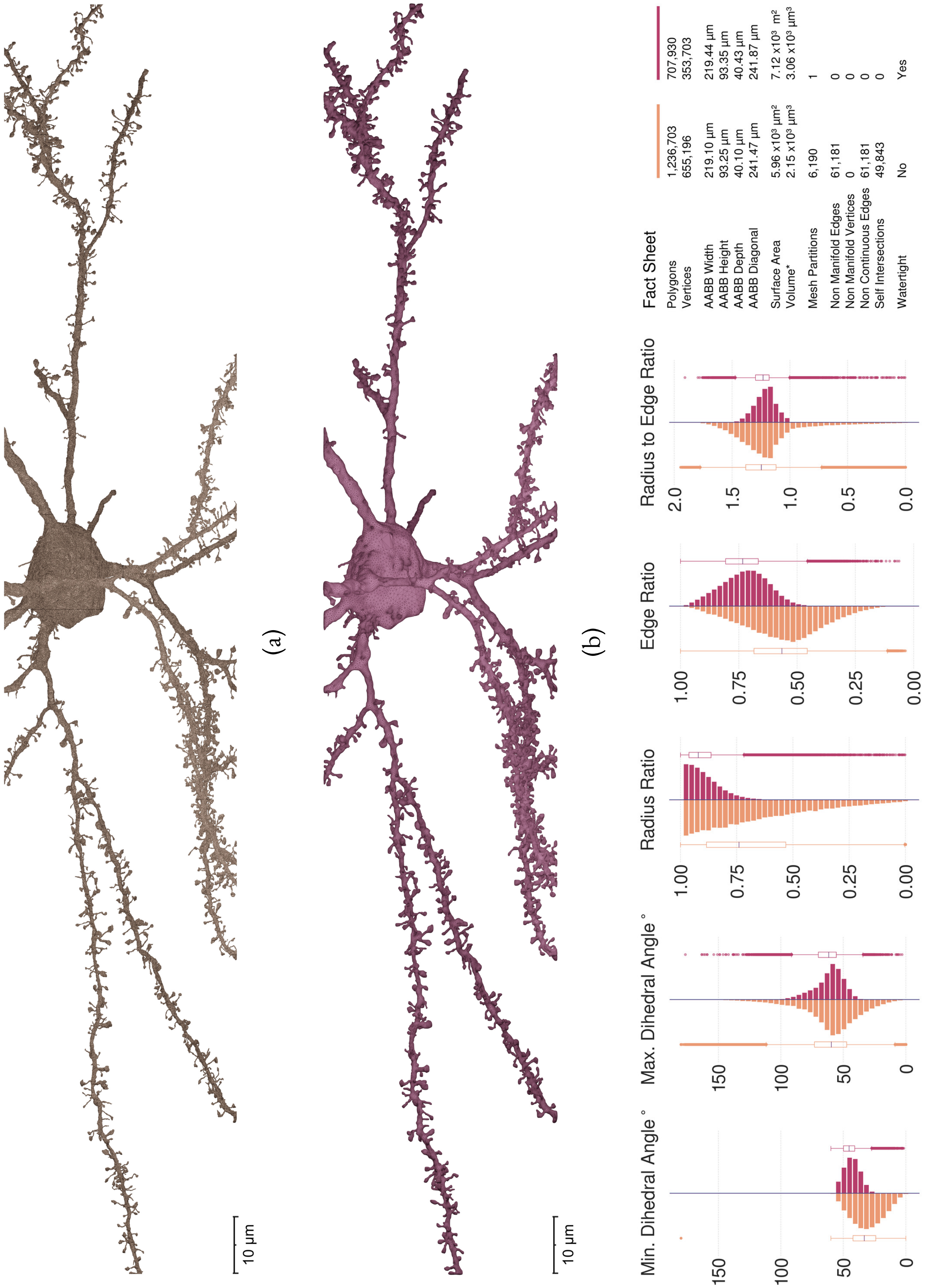

Figure S60: Comparative visual and quantitative analysis between input (in orange) and resulting (in salmon) meshes of Neuron 6.

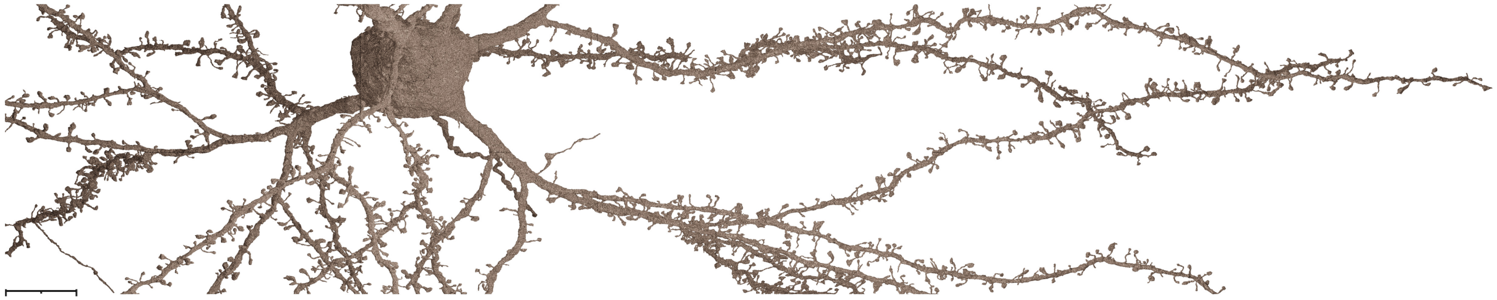

(a)

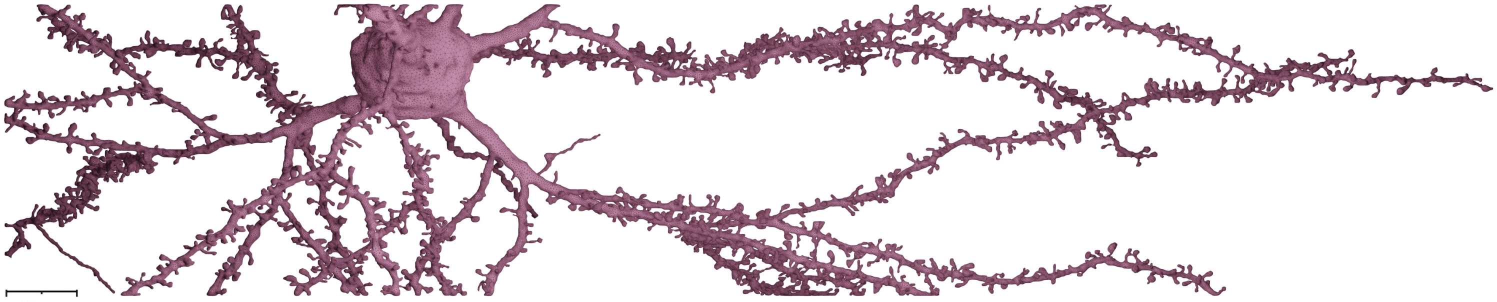

(b)

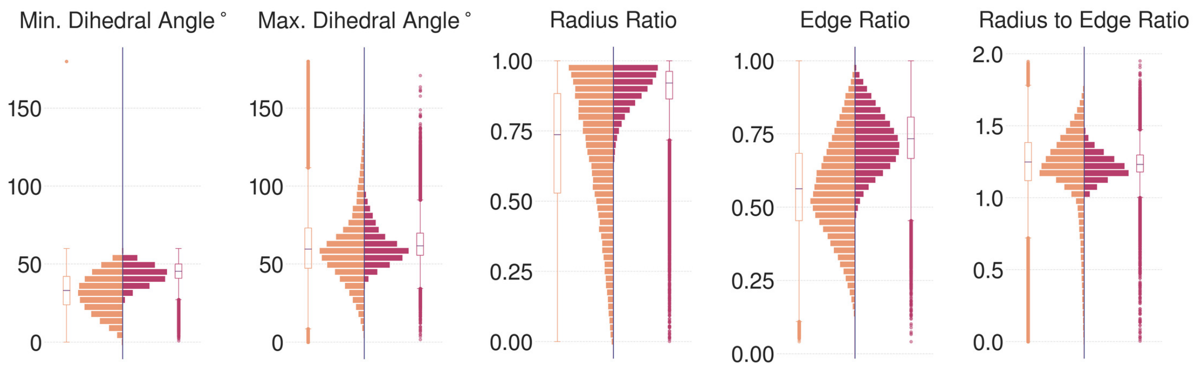

Fact Sheet

| | | |
|---|---|---|
| Polygons | 1,538,180 | 938,164 |
| Vertices | 809,426 | 468,744 |
| AABB Width | 205.74 µm | 205.98 µm |
| AABB Height | 111.92 µm | 112.02 µm |
| AABB Depth | 40.10 µm | 40.38 µm |
| AABB Diagonal | 237.62 µm | 237.92 µm |
| Surface Area | 7.42 x10³ µm² | 9.06 x10³ m² |
| Volume* | 2.49 x10³ µm³ | 3.60 x10³ µm³ |
| Mesh Partitions | 6,460 | 1 |
| Non Manifold Edges | 67,576 | 0 |
| Non Manifold Vertices | 0 | 0 |
| Non Continuous Edges | 67,576 | 0 |
| Self Intersections | 56,426 | 0 |
| Watertight | No | Yes |

(c)

Figure S61: Comparative visual and quantitative analysis between input (in orange) and resulting (in salmon) meshes of Neuron 7.

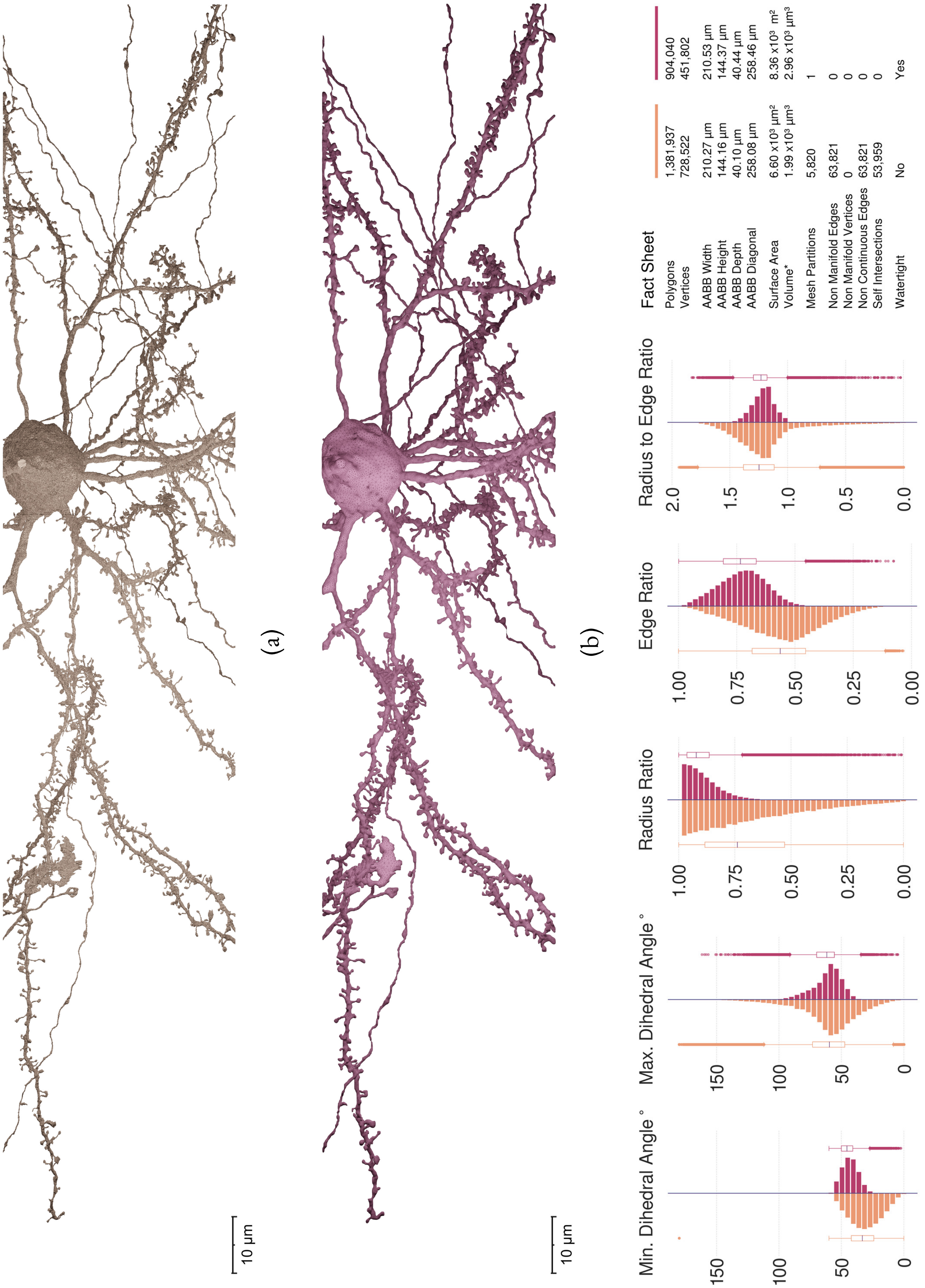

Figure S62: Comparative visual and quantitative analysis between input (in orange) and resulting (in salmon) meshes of Neuron 8.

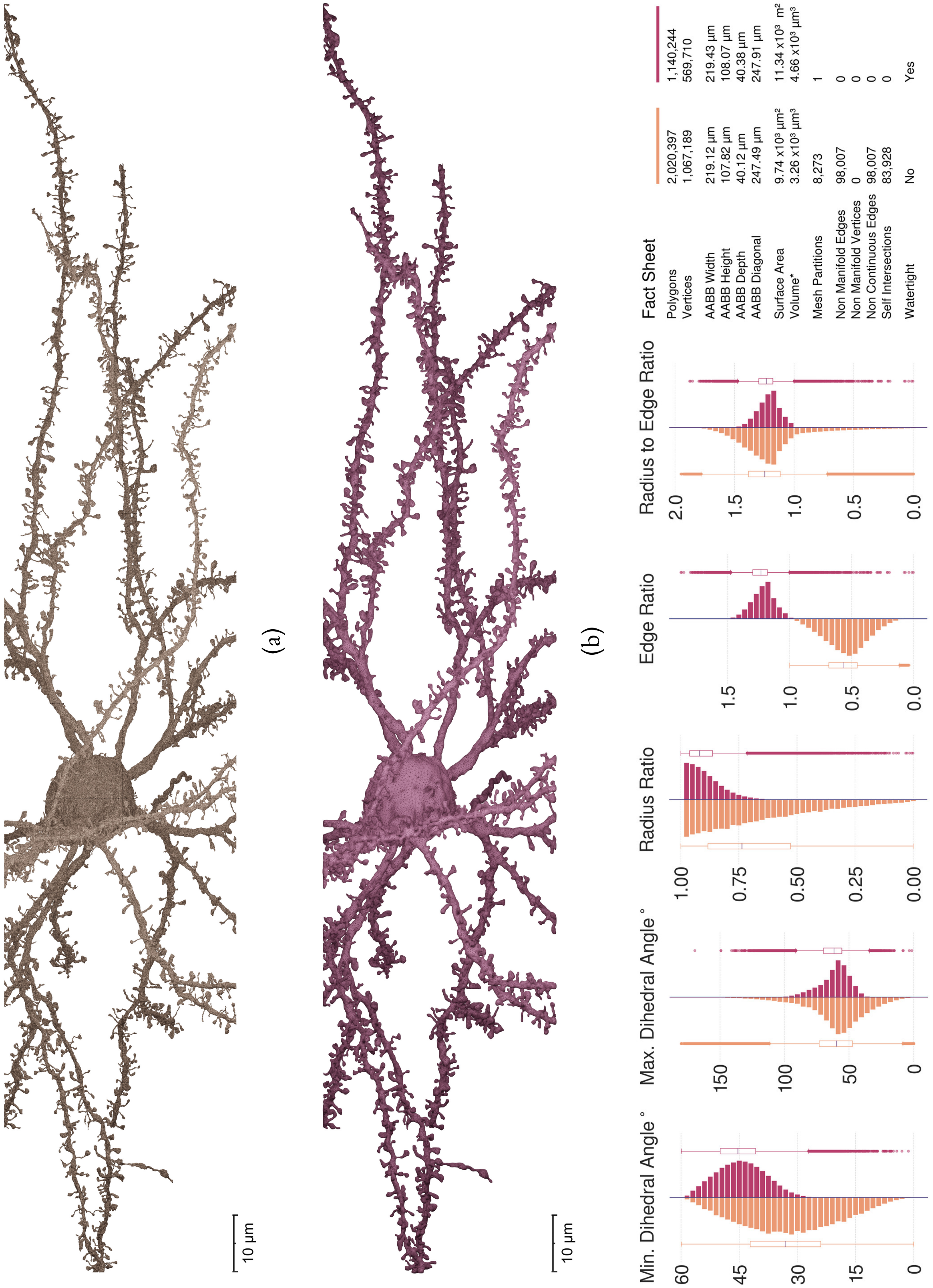

Figure S63: Comparative visual and quantitative analysis between input (in orange) and resulting (in salmon) meshes of Neuron 9.

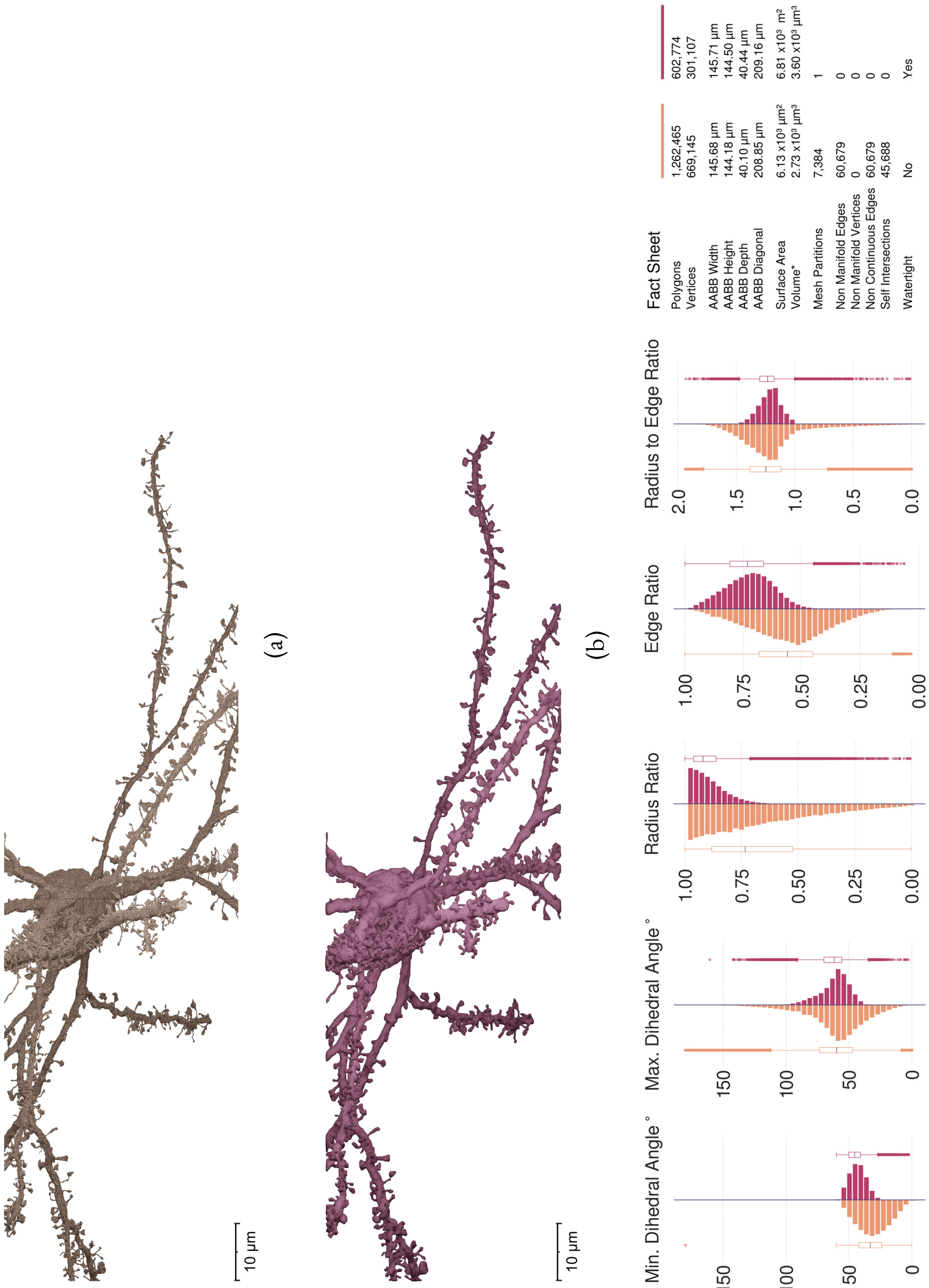

Figure S64: Comparative visual and quantitative analysis between input (in orange) and resulting (in salmon) meshes of Neuron 10.

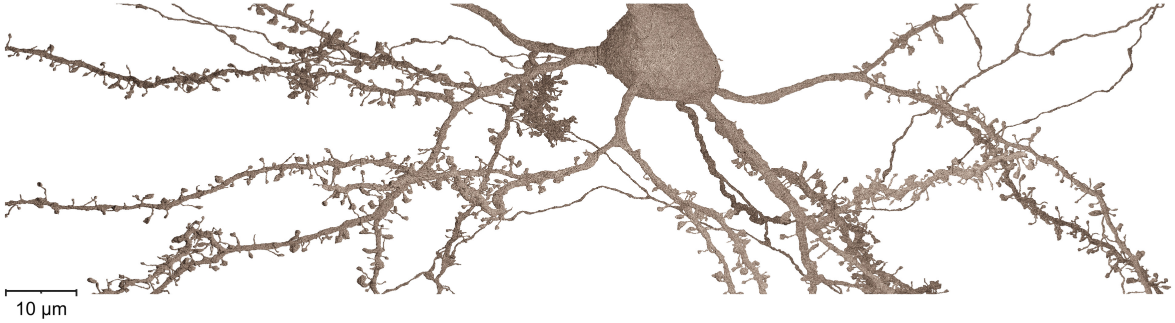

(a)

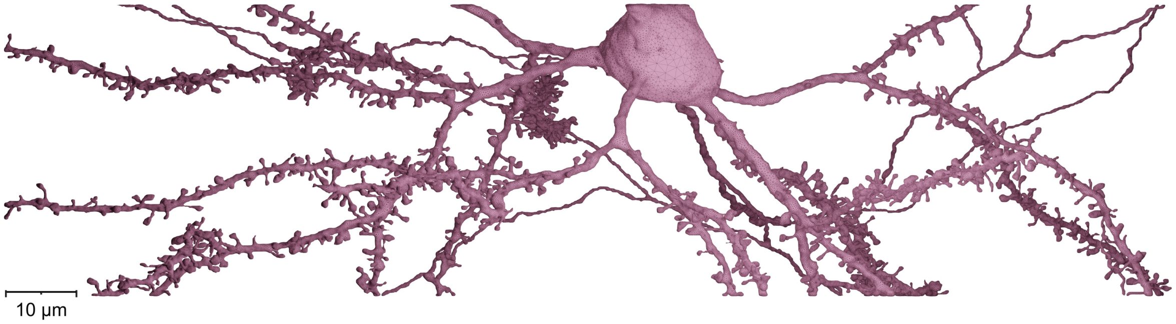

(b)

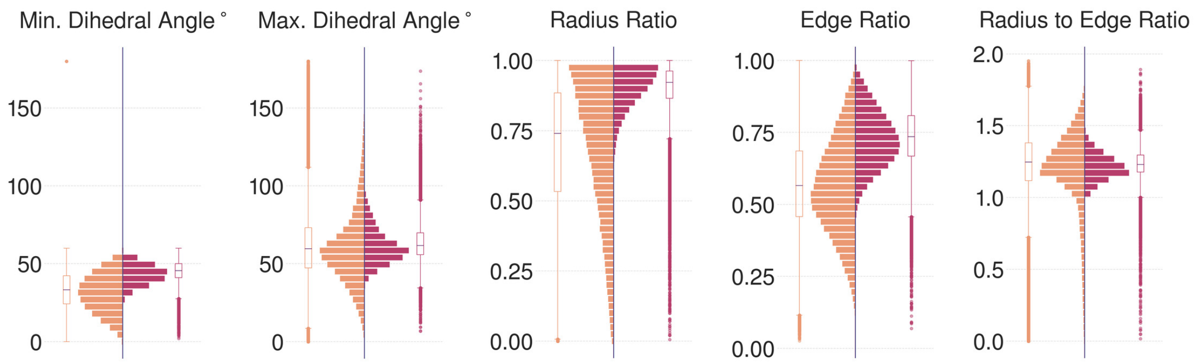

| Fact Sheet | | |
|---|---|---|
| Polygons | 1,375,305 | 868,326 |
| Vertices | 716,777 | 434,003 |
| AABB Width | 161.30 µm | 161.49 µm |
| AABB Height | 144.18 µm | 144.42 µm |
| AABB Depth | 40.10 µm | 40.61 µm |
| AABB Diagonal | 220.03 µm | 220.43 µm |
| Surface Area | 6.70 x10³ µm² | 8.33 x10³ m² |
| Volume* | 2.28 x10³ µm³ | 3.21 x10³ µm³ |
| Mesh Partitions | 4,289 | 1 |
| Non Manifold Edges | 49,753 | 0 |
| Non Manifold Vertices | 0 | 0 |
| Non Continuous Edges | 49,753 | 0 |
| Self Intersections | 43,947 | 0 |
| Watertight | No | Yes |

(c)

Figure S65: Comparative visual and quantitative analysis between input (in orange) and resulting (in salmon) meshes of Neuron 11.

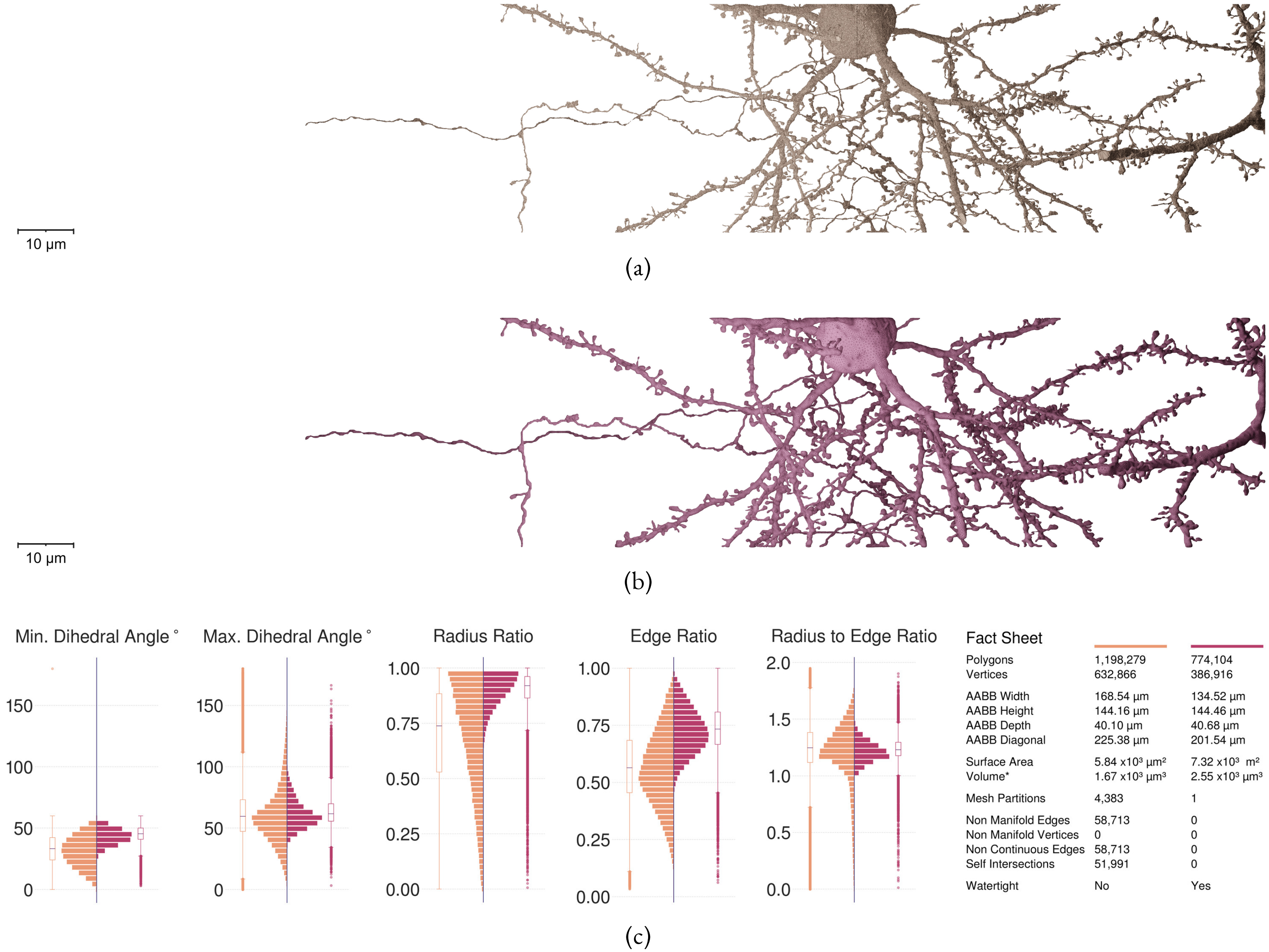

Figure S66: Comparative visual and quantitative analysis between input (in orange) and resulting (in salmon) meshes of Neuron 12.

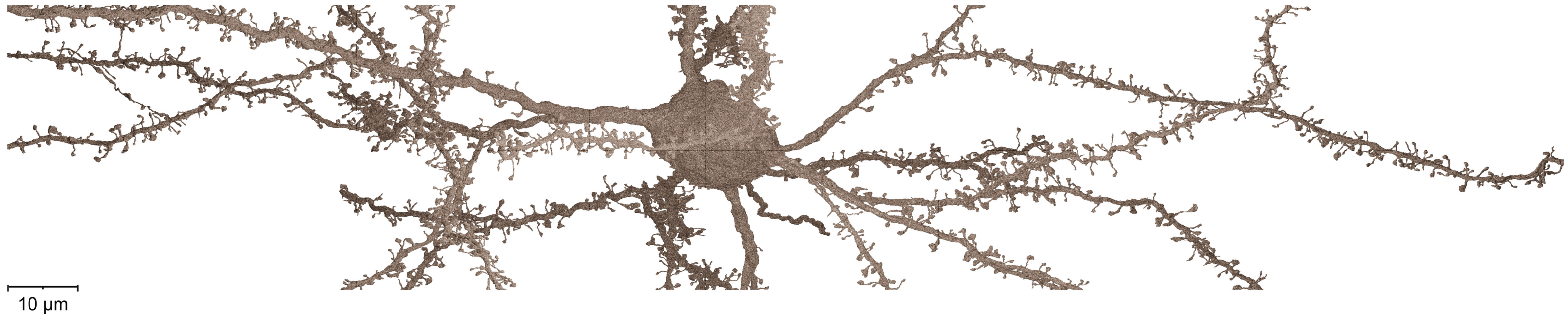

(a)

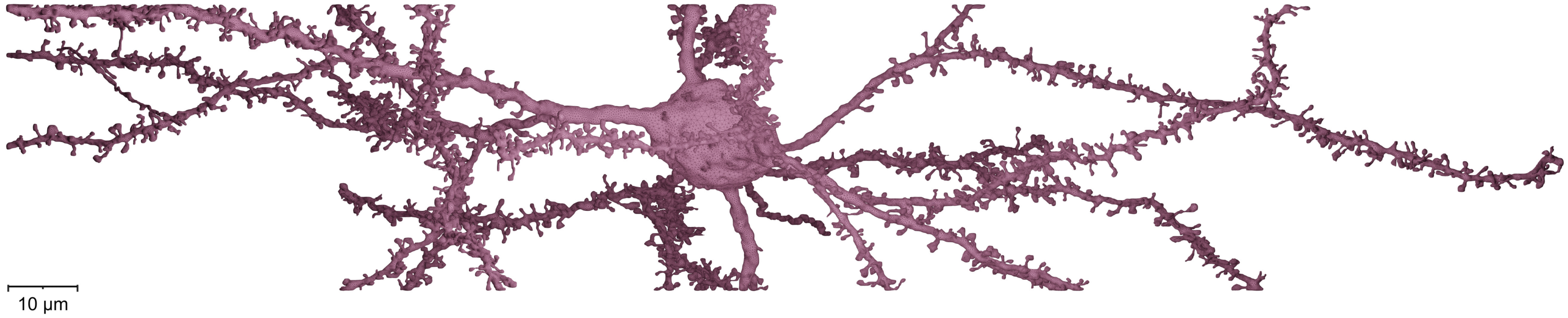

(b)

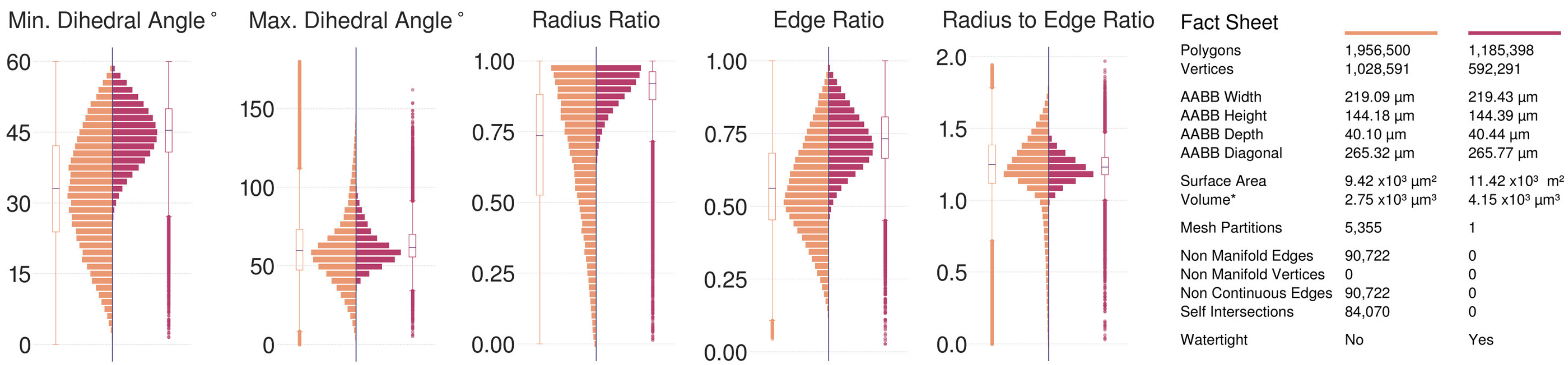

(c)

Figure S67: Comparative visual and quantitative analysis between input (in orange) and resulting (in salmon) meshes of Neuron 13.

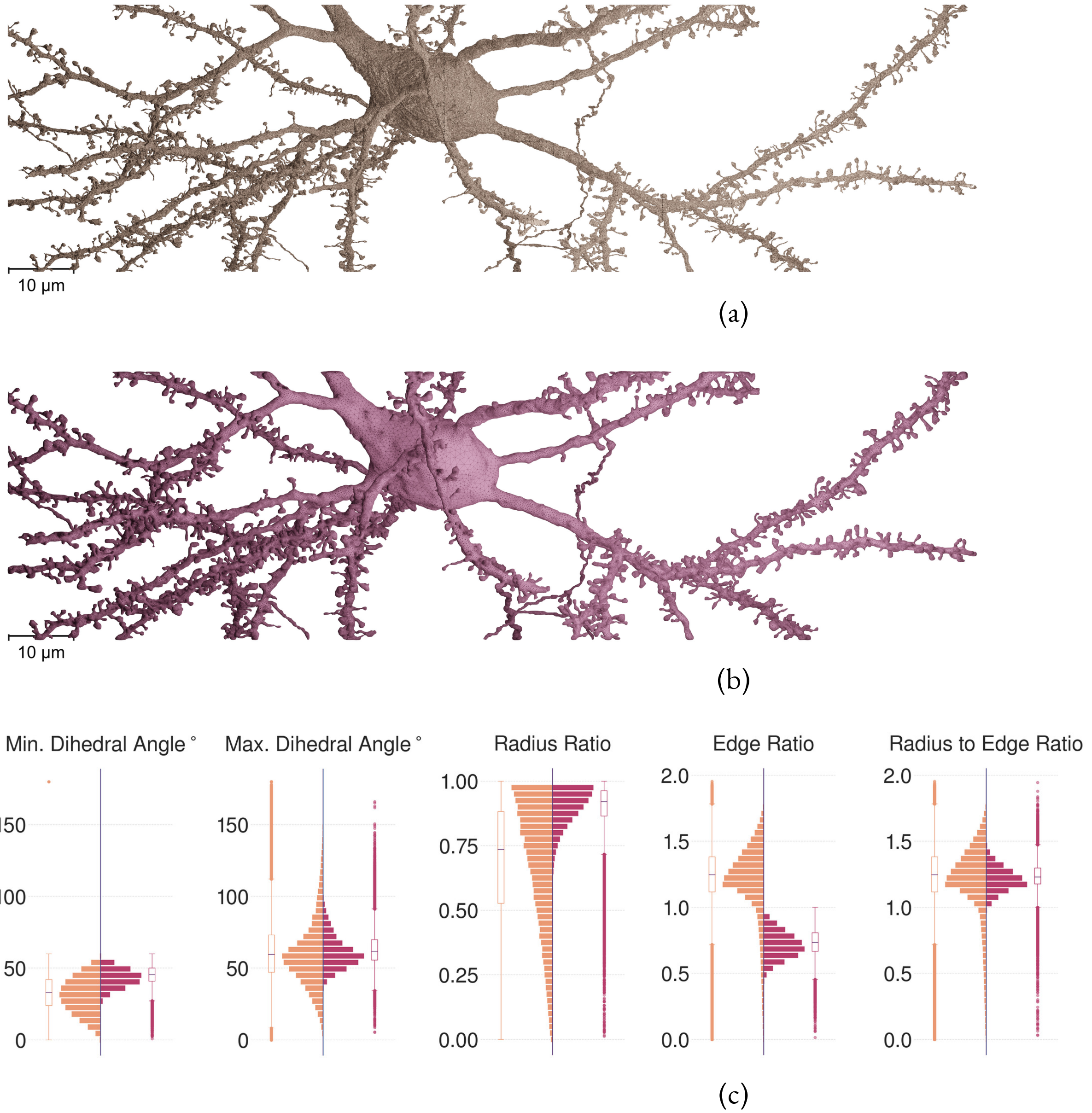

| Fact Sheet | | |
|---|---|---|
| Polygons | 1,622,524 | 927,384 |
| Vertices | 854,478 | 463,406 |
| AABB Width | 145.57 μm | 145.58 μm |
| AABB Height | 108.10 μm | 108.32 μm |
| AABB Depth | 40.10 μm | 40.41 μm |
| AABB Diagonal | 185.70 μm | 185.90 μm |
| Surface Area | 7.85 x10³ μm² | 9.37 x10³ m² |
| Volume* | 2.82 x10³ μm³ | 3.90 x10³ μm³ |
| Mesh Partitions | 6,168 | 1 |
| Non Manifold Edges | 74,198 | 0 |
| Non Manifold Vertices | 0 | 0 |
| Non Continuous Edges | 74,198 | 0 |
| Self Intersections | 64,652 | 0 |
| Watertight | No | Yes |

Figure S68: Comparative visual and quantitative analysis between input (in orange) and resulting (in salmon) meshes of Neuron 14.

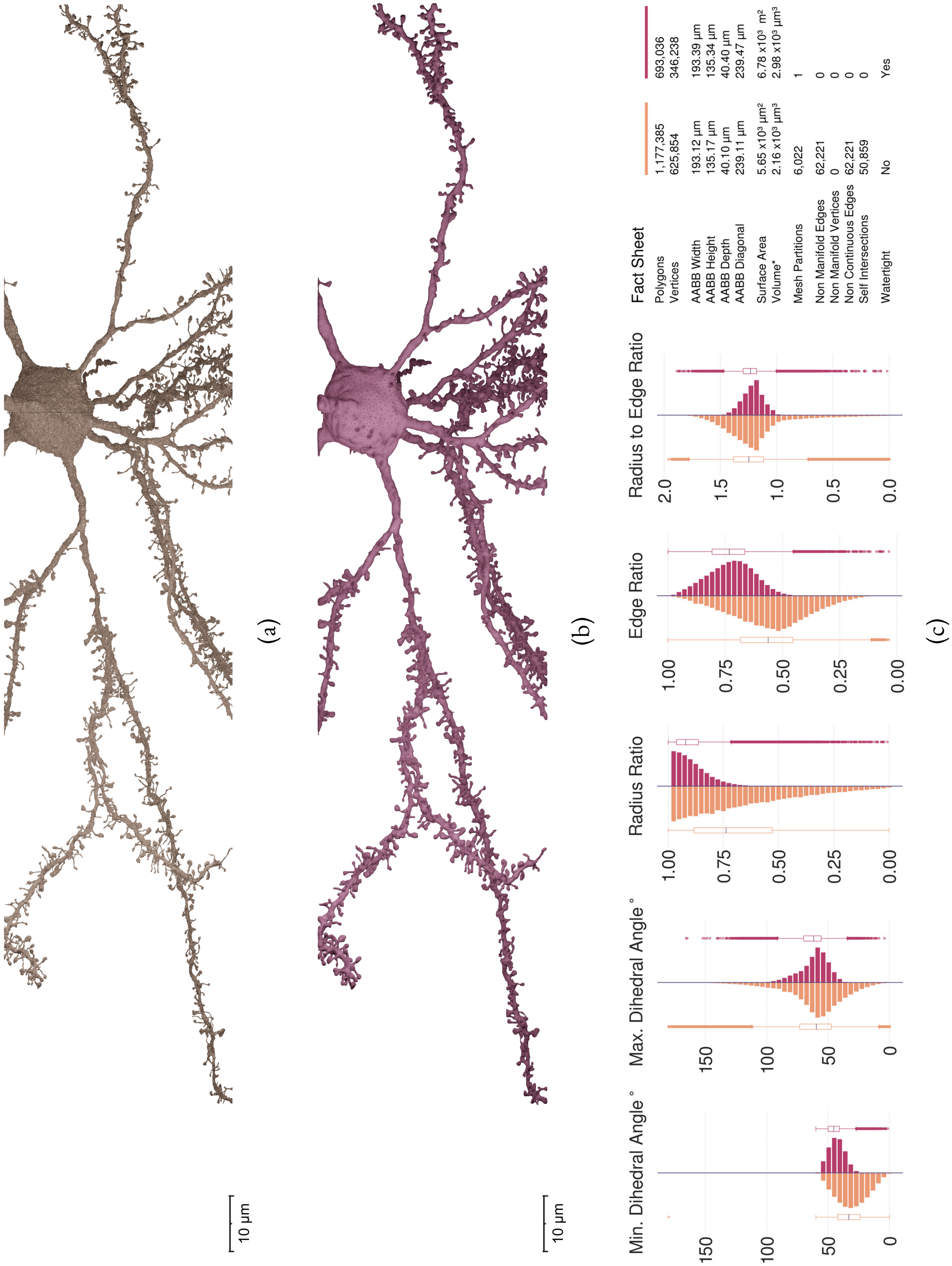

Figure S69: Comparative visual and quantitative analysis between input (in orange) and resulting (in salmon) meshes of Neuron 15.

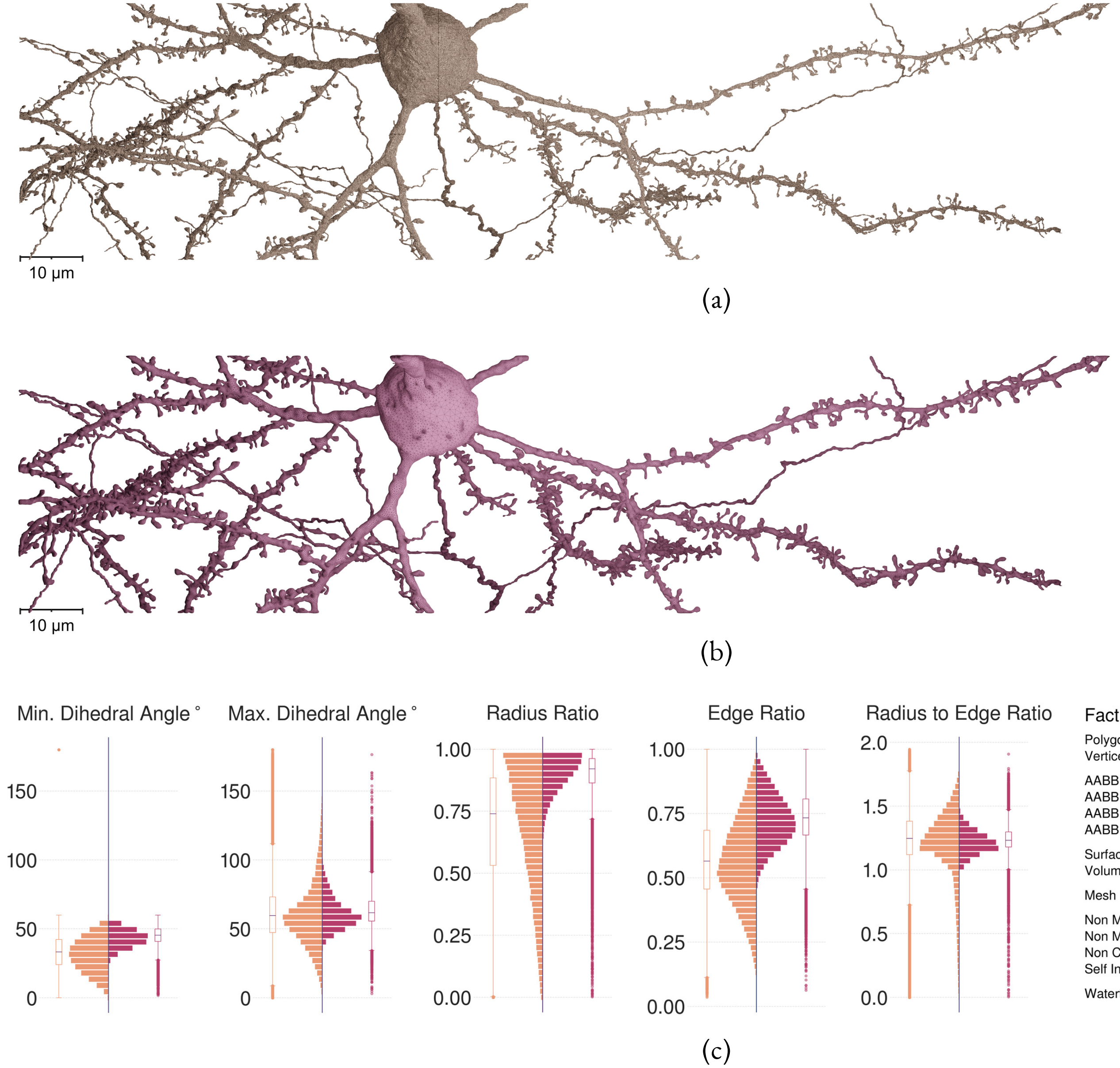

Fact Sheet

| | Input (orange) | Resulting (salmon) |
|---|---|---|
| Polygons | 1,187,921 | 762,754 |
| Vertices | 625,716 | 381,197 |
| AABB Width | 170.61 µm | 170.66 µm |
| AABB Height | 140.14 µm | 140.34 µm |
| AABB Depth | 40.10 µm | 40.40 µm |
| AABB Diagonal | 224.40 µm | 224.61 µm |
| Surface Area | 5.71 x10³ µm² | 7.18 x10³ m² |
| Volume* | 1.89 x10³ µm³ | 2.70 x10³ µm³ |
| Mesh Partitions | 4,288 | 1 |
| Non Manifold Edges | 55,333 | 0 |
| Non Manifold Vertices | 0 | 0 |
| Non Continuous Edges | 55,333 | 0 |
| Self Intersections | 48,790 | 0 |
| Watertight | No | Yes |

Figure S70: Comparative visual and quantitative analysis between input (in orange) and resulting (in salmon) meshes of Neuron 16.

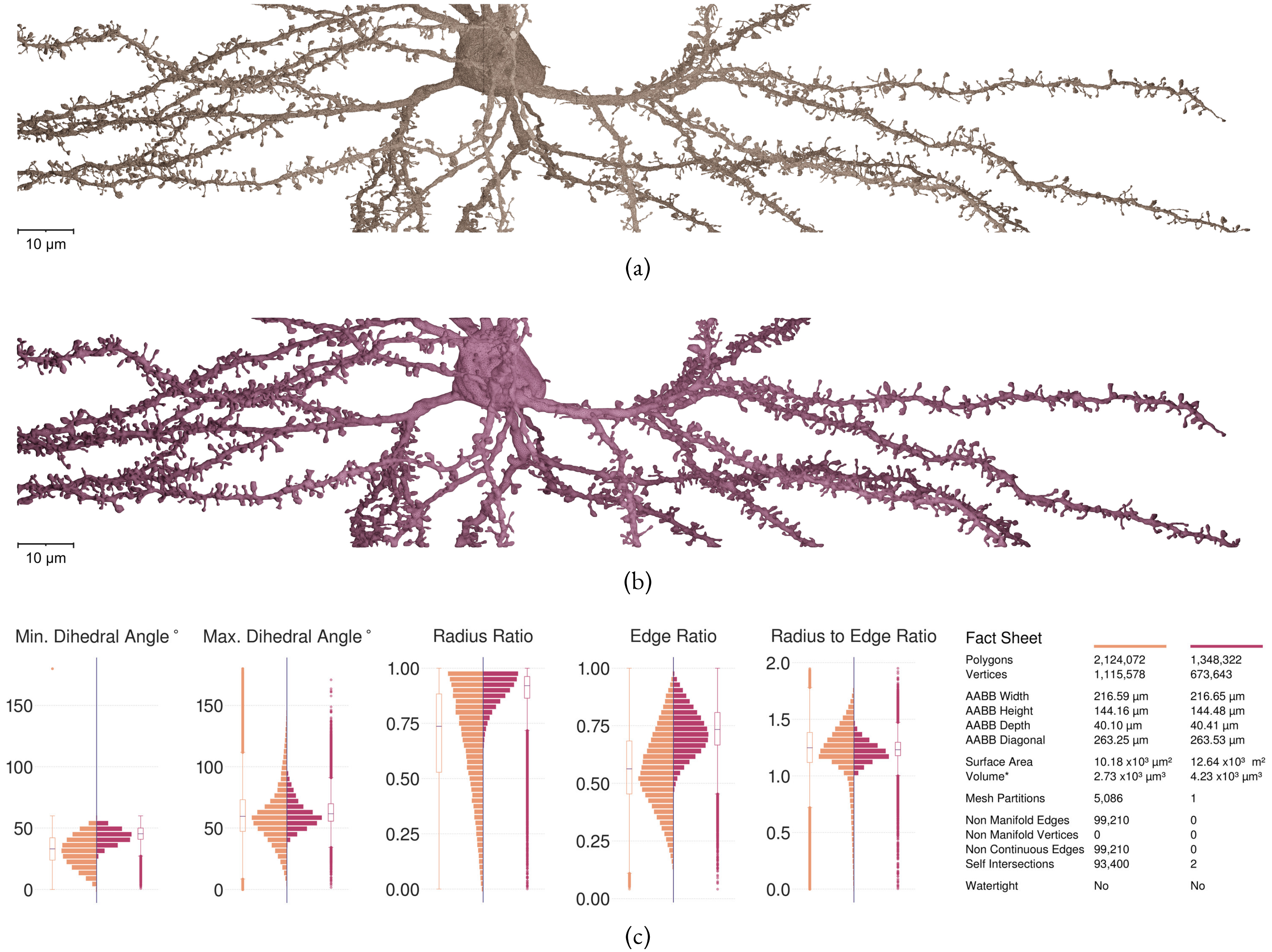

Figure S71: Comparative visual and quantitative analysis between input (in orange) and resulting (in salmon) meshes of Neuron 17.

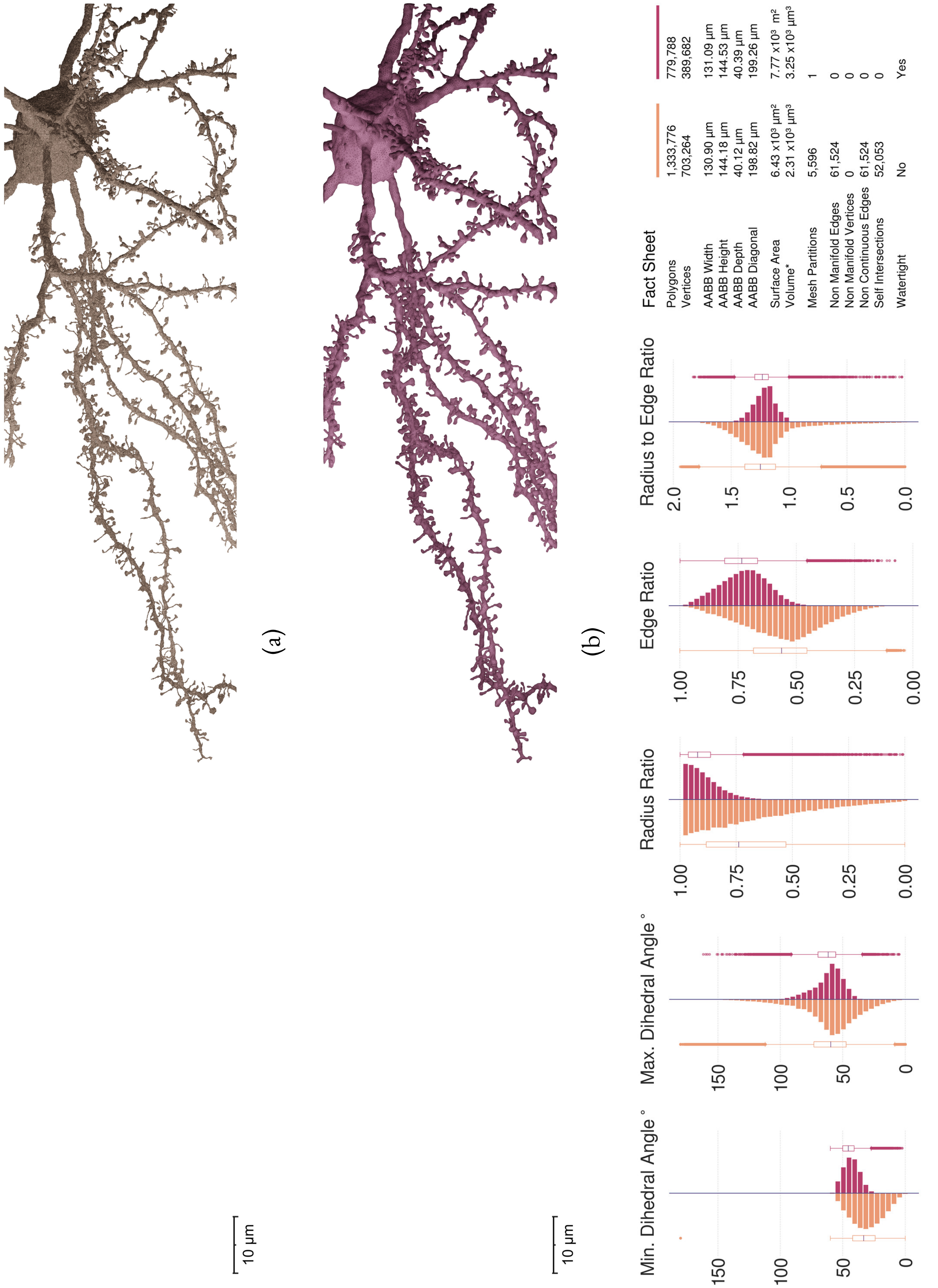

Figure S72: Comparative visual and quantitative analysis between input (in orange) and resulting (in salmon) meshes of Neuron 18.

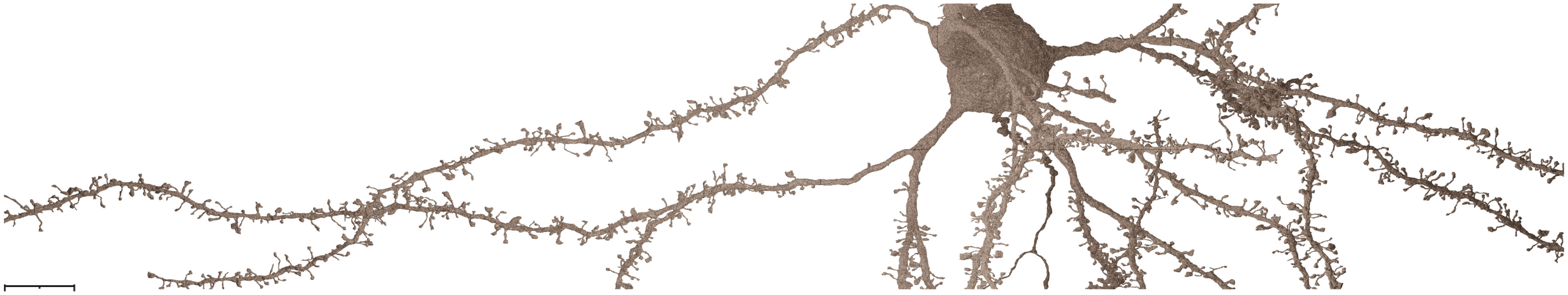

(a)

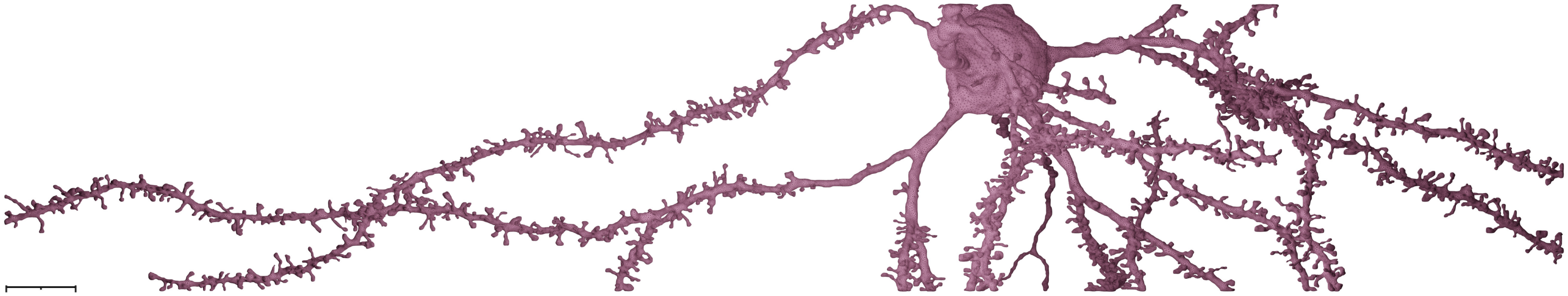

(b)

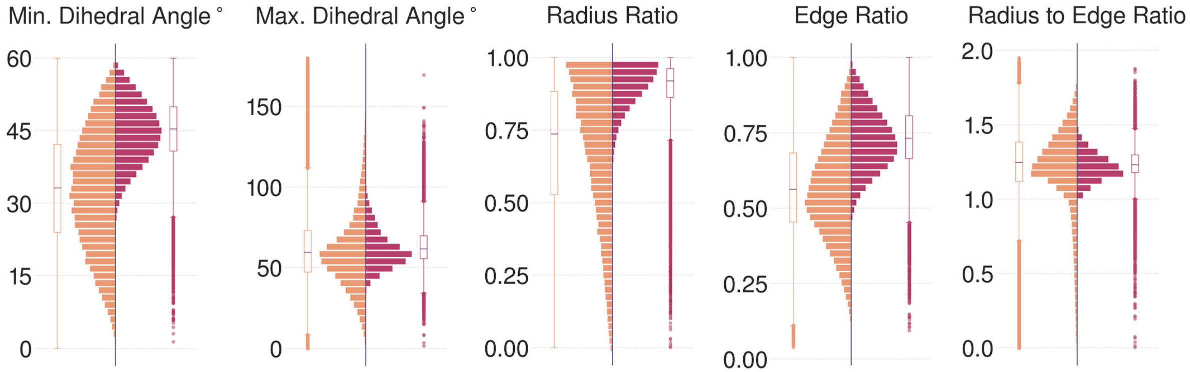

| Fact Sheet | | |
|---|---|---|
| Polygons | 1,345,280 | 803,710 |
| Vertices | 711,478 | 401,623 |
| AABB Width | 219.12 μm | 219.46 μm |
| AABB Height | 144.18 μm | 144.42 μm |
| AABB Depth | 40.10 μm | 40.49 μm |
| AABB Diagonal | 265.35 μm | 265.82 μm |
| Surface Area | 6.43 x10³ μm² | 7.83 x10³ m² |
| Volume* | 2.22 x10³ μm³ | 3.15 x10³ μm³ |
| Mesh Partitions | 5,207 | 1 |
| Non Manifold Edges | 67,814 | 0 |
| Non Manifold Vertices | 0 | 0 |
| Non Continuous Edges | 67,814 | 0 |
| Self Intersections | 59,995 | 0 |
| Watertight | No | Yes |

(c)

Figure S73: Comparative visual and quantitative analysis between input (in orange) and resulting (in salmon) meshes of Neuron 19.

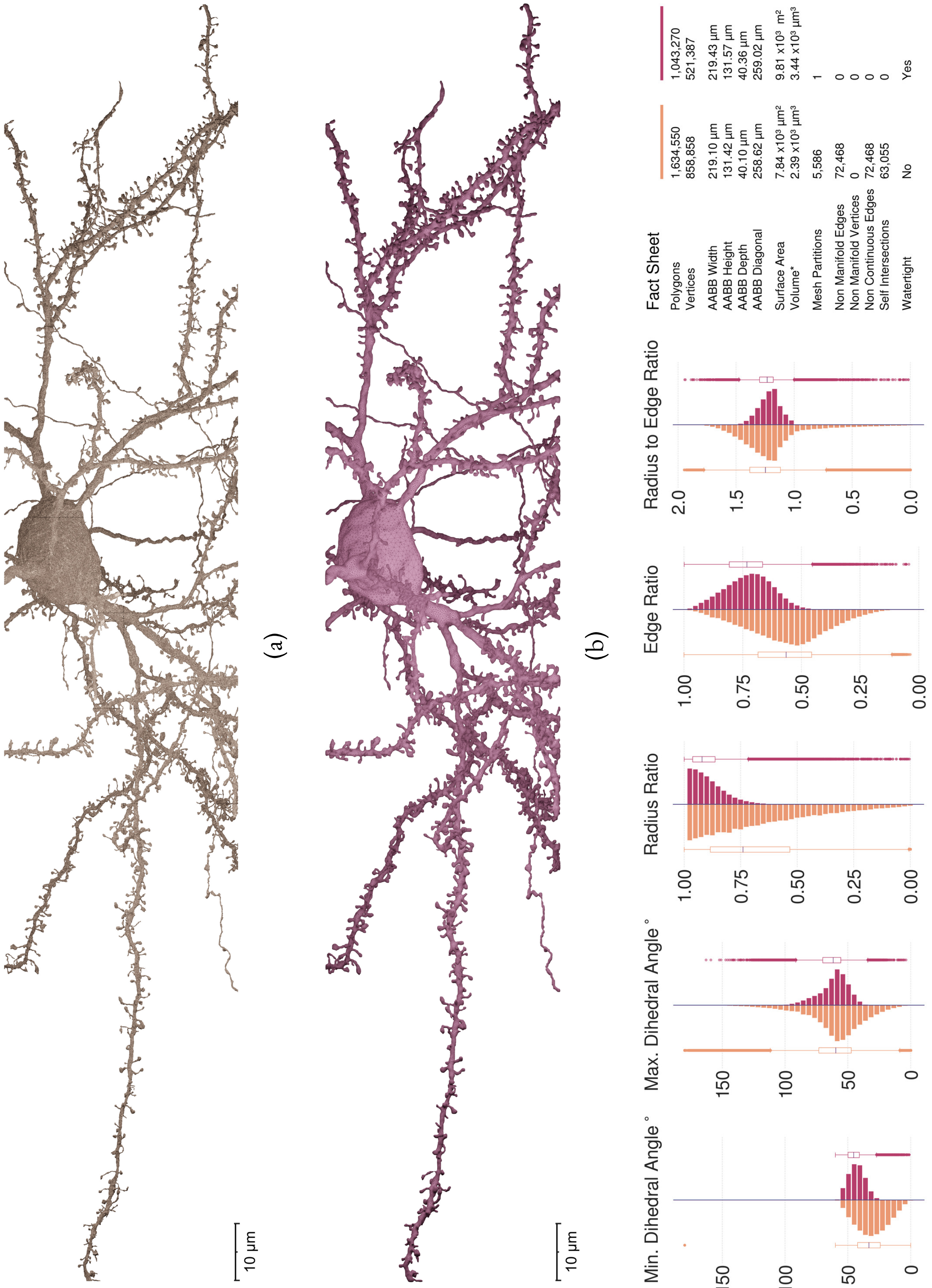

Figure S74: Comparative visual and quantitative analysis between input (in orange) and resulting (in salmon) meshes of Neuron 20.

#### 6.2 Remeshing fragmented meshes with slicing artifacts

In certain cases, segmented meshes might have slicing artifacts, where thin gaps can exist along the dendritic branches. To remove these gaps and create continuous mesh models, we have used ULTRAMESH2MESH with specific voxelization resolution, with which we can connect the fragmented components without changing the structure of the model. Figure S75 demonstrates an example with a fragmented neuronal mesh with slicing artifacts.

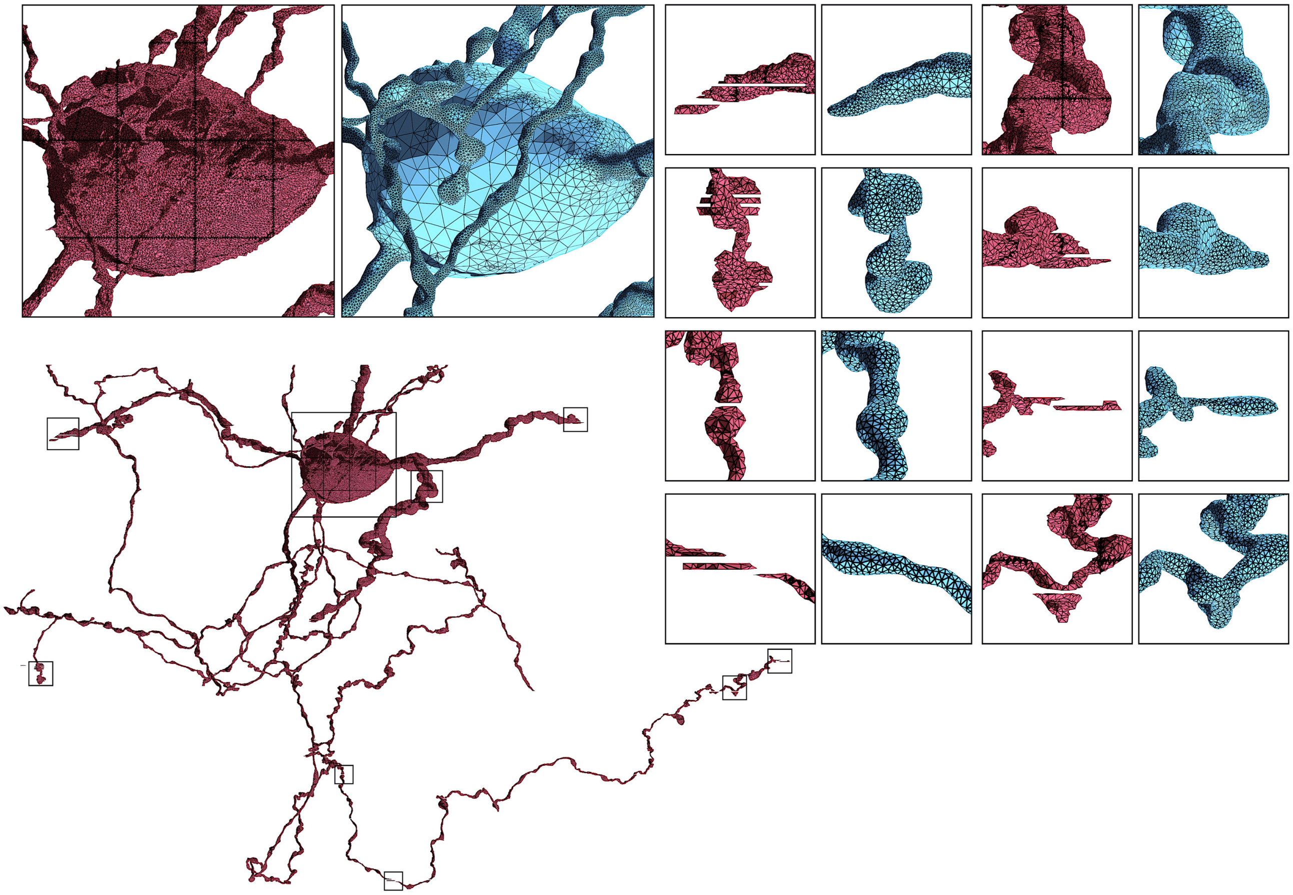

Figure S75: **Re-meshing Fragmented Mesh Model with Slicing Artifacts**, Related to Fig. 3
Reconstruction of a continuous watertight manifold of a neuronal mesh (in blue) from a fragmented input mesh (in red) having multiple partitions.

### 7 Morphology structures

#### 7.1 Neuronal morphology structure

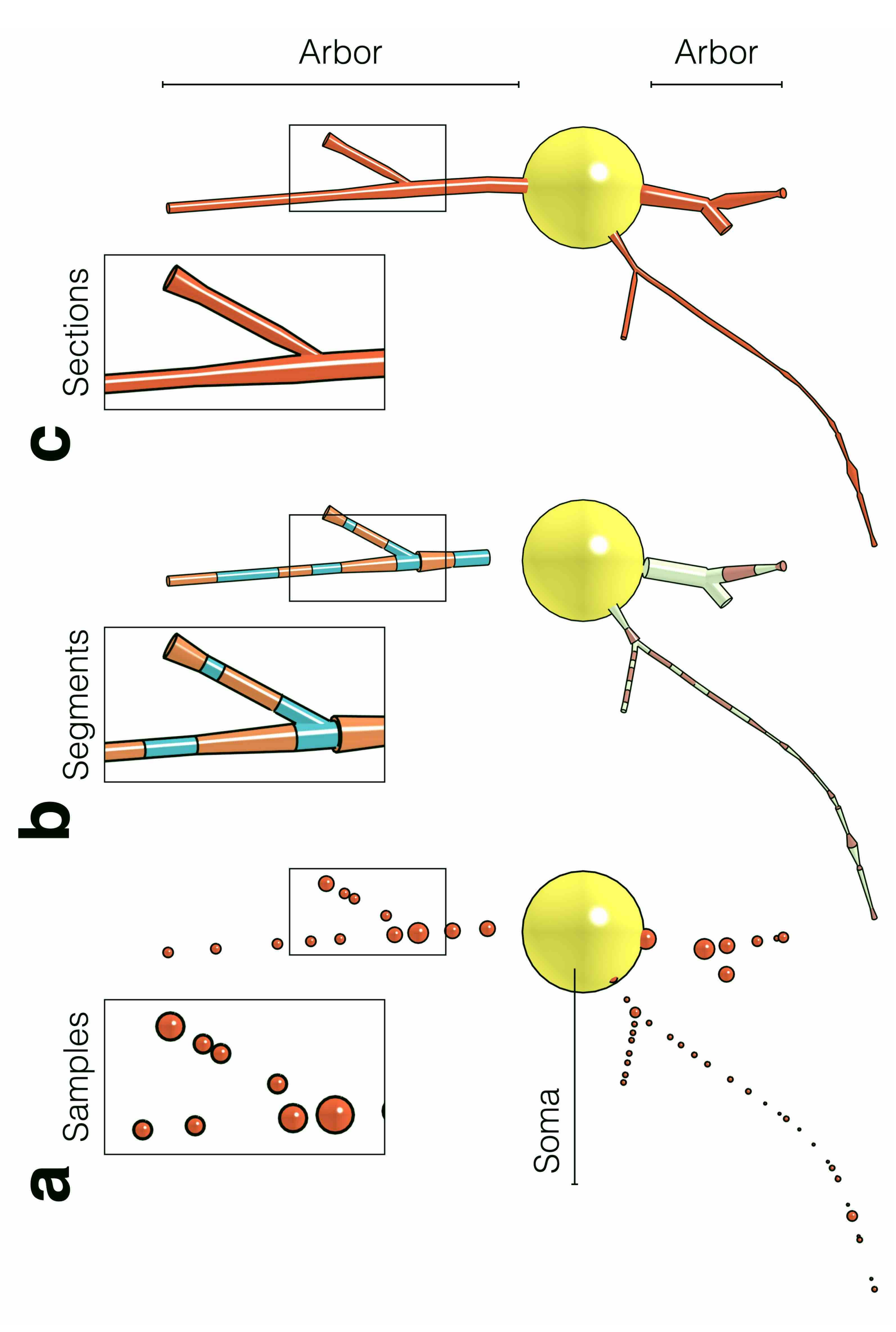

Figure S76: **Neuron Morphology Structure.**
Digitized morphology skeletons are represented by acyclic graphs. The root node represents the cell body (or soma) –in yellow– of the neuron. The morphology is traced and digitized into a list of samples (a). Each sample has a unique index, 3D Cartesian coordinate and radius. A pair of connected samples represents a segment (b). A concatenated list of adjacent segments between two branching points represents a section (c). Neurites or arbors are composed of a hierarchical organization of sections. Note that branching samples might have different radii and dendritic spines are not shown in this illustration.

#### 7.2 Astrocyte morphology structure

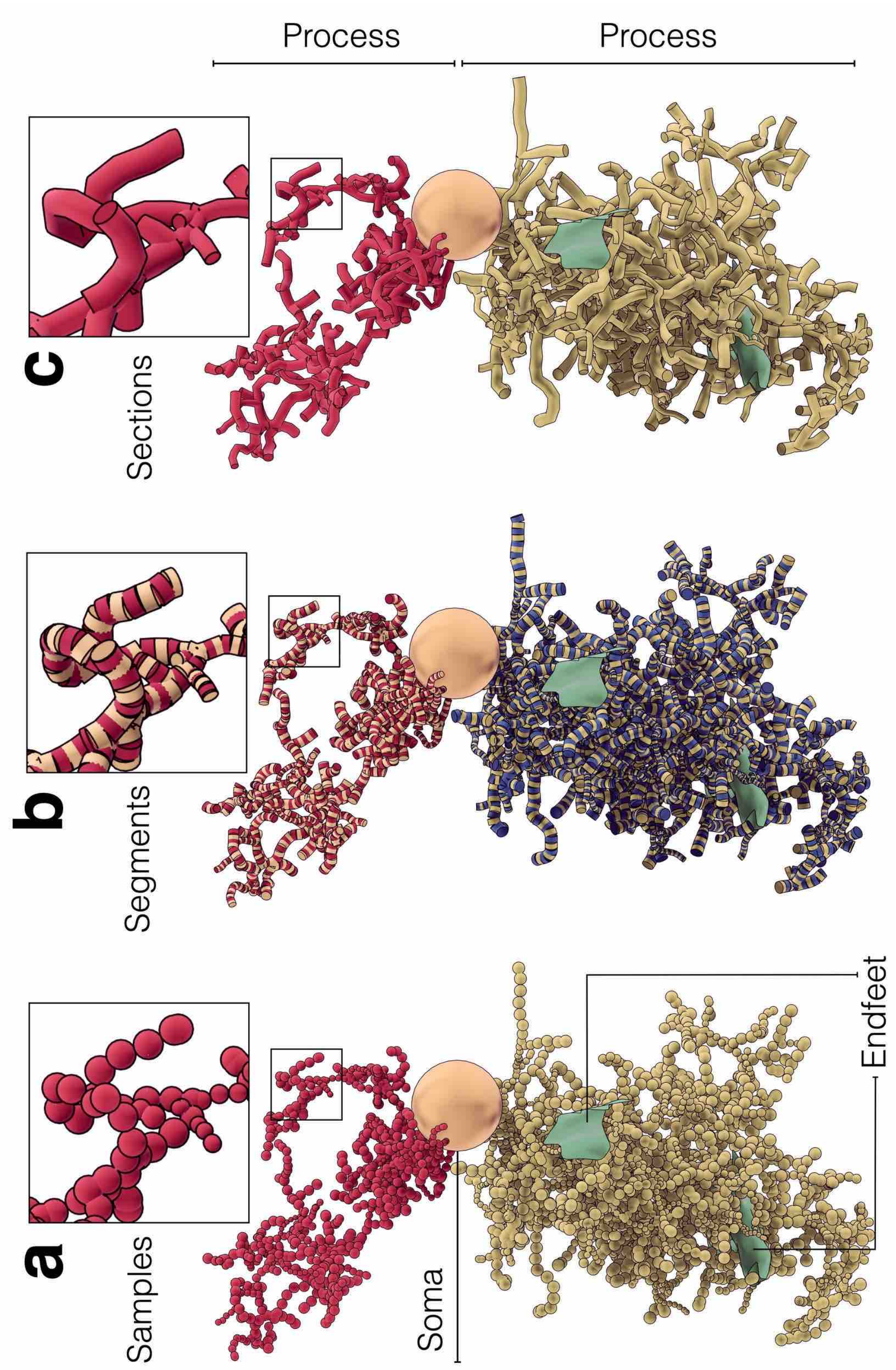

Figure S77: **Astrocyte Morphology Structure**.
Astrocytic morphologies are similar to the neuronal ones shown in Figure S76, but additionally, they have multiple endfeet –shown in green. These endfeet are represented by triangular patches (list of connected triangles), where each vertex along the surface has a thickness. Further details on the exact structure of astrocytic morphologies are available[13,23]. Note that astrocytic processes have sponge-like spatial structure in contrast to neurons, but they are both represented by acyclic graphs.

#### 7.3 Vascular morphology structure

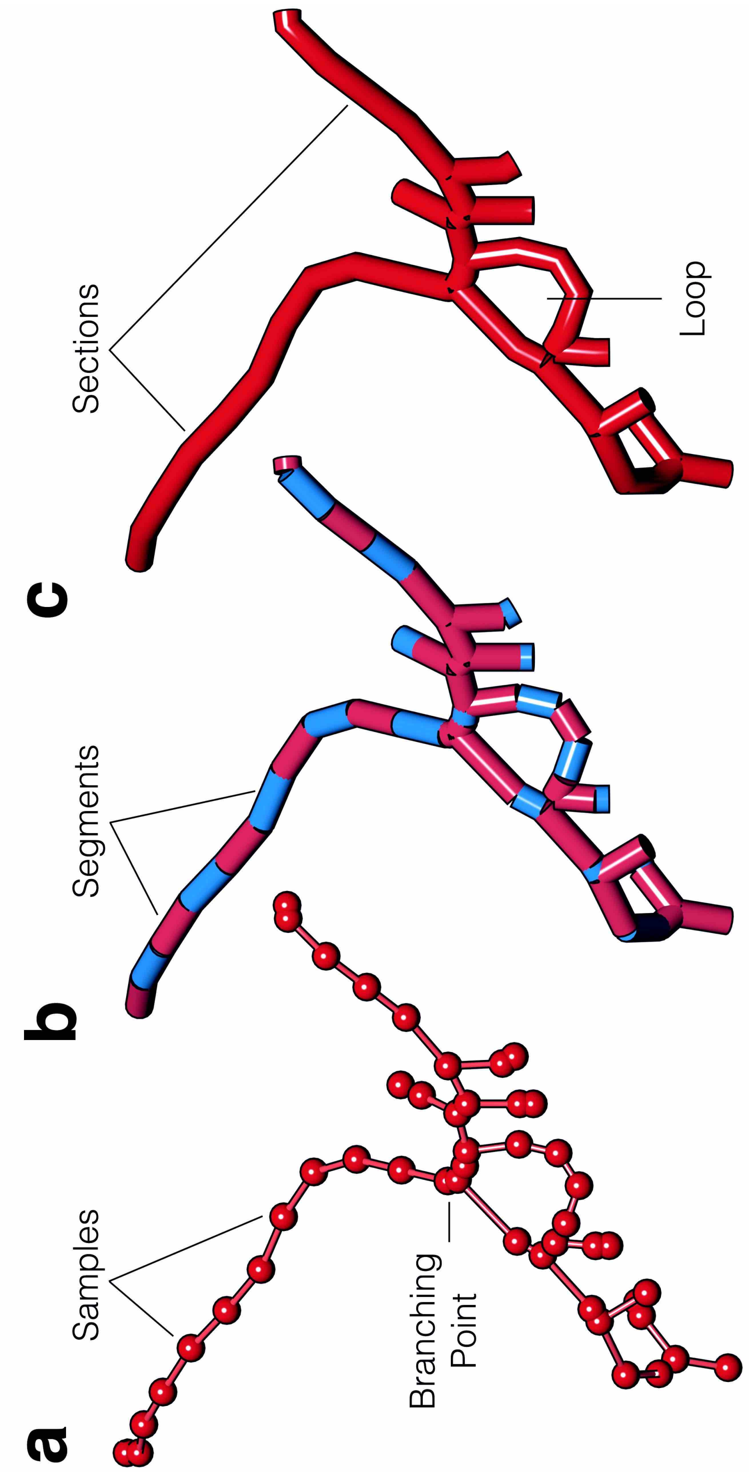

Figure S78: **Vascular Morphology Structure**.
Vascular morphologies are represented with cyclic graphs to account for loops. The branching structure is similar to the arborizations of neurons and astrocytes. The skeleton is digitized into a list of connected samples (a), where every adjacent pair of samples represents a segment (b) and a group of connected segments between two branching points represents a section. Each section in the morphology has a unique index, and references to its parent and child sections, with which we can reconstruct the hierarchical organization of the graph. The loops are identified in the morphology when two sections have the same references to parent and child sections.

### 8 Spine geometries extracted from cortical EM volumes

To enhance the realism of the resulting neuronal meshes, we extracted ∼50 spine meshes from the four neurons shown in Figures S10– S13. Each neuronal mesh is loaded in Blender, where spine geometries are visually identified. Afterwards, and for each spine, we created a bounding box covering its spatial extent and overlapping with the dendritic section it emanates from. We then applied, per spine, a mesh intersection operator to extract its geometry as an independent object. The spines are oriented along the Y-axis after identifying their base and apex. Spine geometries are then processed to clean any self-intersecting facets along their surfaces, optimized and finally exported as independent mesh objects. A collage of a set of exemplar spines with various types and geometries is shown in Figure S79.

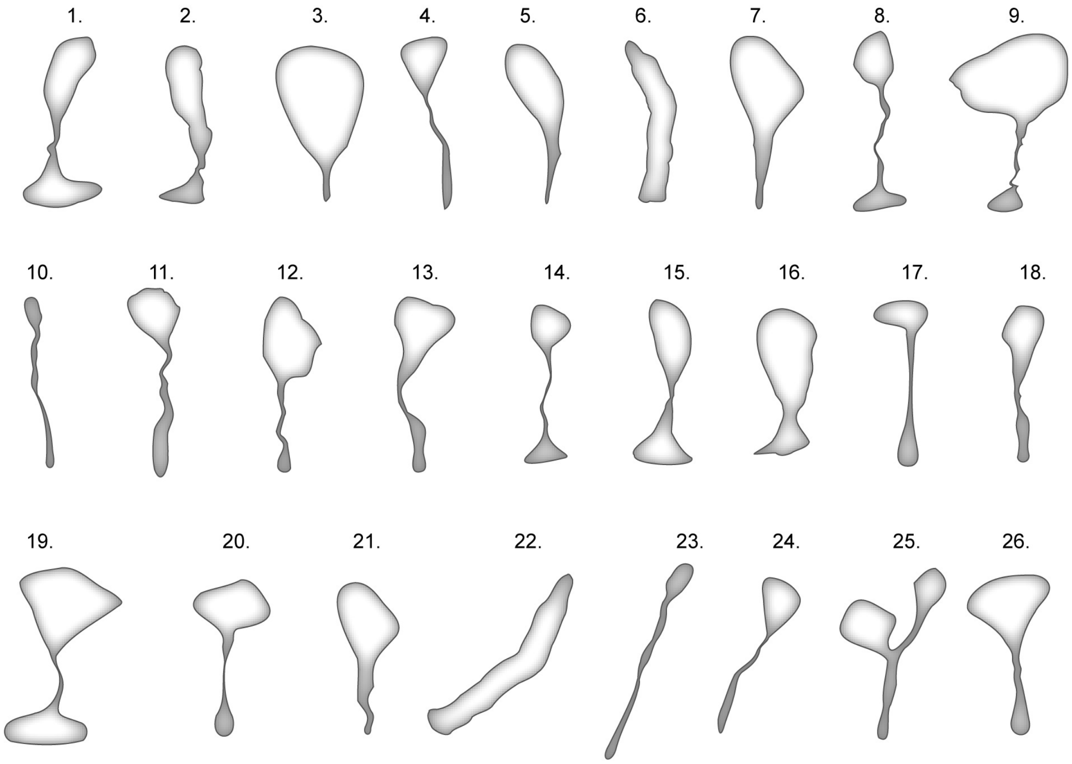

Figure S79: **Realistic Geometries of Neuronal Spines with Various Shapes.**
The geometries are manually extracted from the neuronal meshes shown in Figures S10, S11, S12 and S13.

### 9 Generating biologically realistic neuronal meshes from digitized morphologies

The proxy mesh generation that represents each section or computed path is approached in two steps. Firstly, the positions and directions along the path, where a sectional geometry in the form of circumference is placed and oriented are computed. Finally, and once this geometry is properly emplaced, we obtain a tubular mesh by the subsequent connection of the geometry following the path order.

Table S8: Symbols

| Symbol | Definition |
| --- | --- |
| $p$ | Point |
| $h$ | Interpolation step $[0,1]$ |
| $p_1$ and $p_2$ | Positions of the original segment nodes |
| $m_1$ and $m_2$ | Tangents of the original segment nodes |
| $p_0$ | Position of the previous node |
| $p_3$ | Position of the next node |

To obtain a smooth reconstruction for each path, the sectional geometries are not only emplaced in the existing samples, but new positions are computed in between the segments determined by the samples. For each segment, we make use of the Cubic Hermite spline interpolation[24] to compute the new positions according to Equation 15.

$$p(h) = (2h^3 - 3h^2 + 1)p_1 + (h^3 - 2h^2 + h)m_1 + (-2h^3 + 3h^2)p_2 + (h^3 - h^2)m_2 \tag{15}$$

The directions at each new point are computed using the derivative with respect to the interpolation step, $h$, of the same Hermite spline formulation according to Equation 16.

$$m(h) = (6h^2 - 6^h)p_1 + (3h^2 - 4h + 1)m_1 + (-6h^2 + 6h)p_2 + (3h^2 - 2h)m_2 \tag{16}$$

To avoid loops or self-intersections between the generated points, the tangent at each original segment point is computed using the Centripetal Catmull-Rom spline formulation, that uses the positions of the previous and next samples to the current segment:

$$m_1 = (t_2 - t_1)\left(\frac{p_1 - p_0}{t_1 - t_0} - \frac{p_2 - p_0}{t_2 - t_0} + \frac{p_2 - p_1}{t_2 - t_1}\right) \tag{17}$$

$$m_2 = (t_2 - t_1)\left(\frac{p_2 - p_1}{t_2 - t_1} - \frac{p_3 - p_1}{t_3 - t_1} + \frac{p_3 - p_2}{t_3 - t_2}\right) \tag{18}$$

where: $t_i = ||p_i - p_{i-1}||^{\alpha} + t_{i-1}$ and $t_0 = 0, i = 1, 2, 3$ and $\alpha \in [0, 1]$.

Afterwards, all the generated proxy mesh (including the somatic mesh created from the FEM simulation) are rasterized in a volume grid, where a continuous manifold is obtained, smoothed, optimized and verified to be watertight (**Fig.** **S80**).

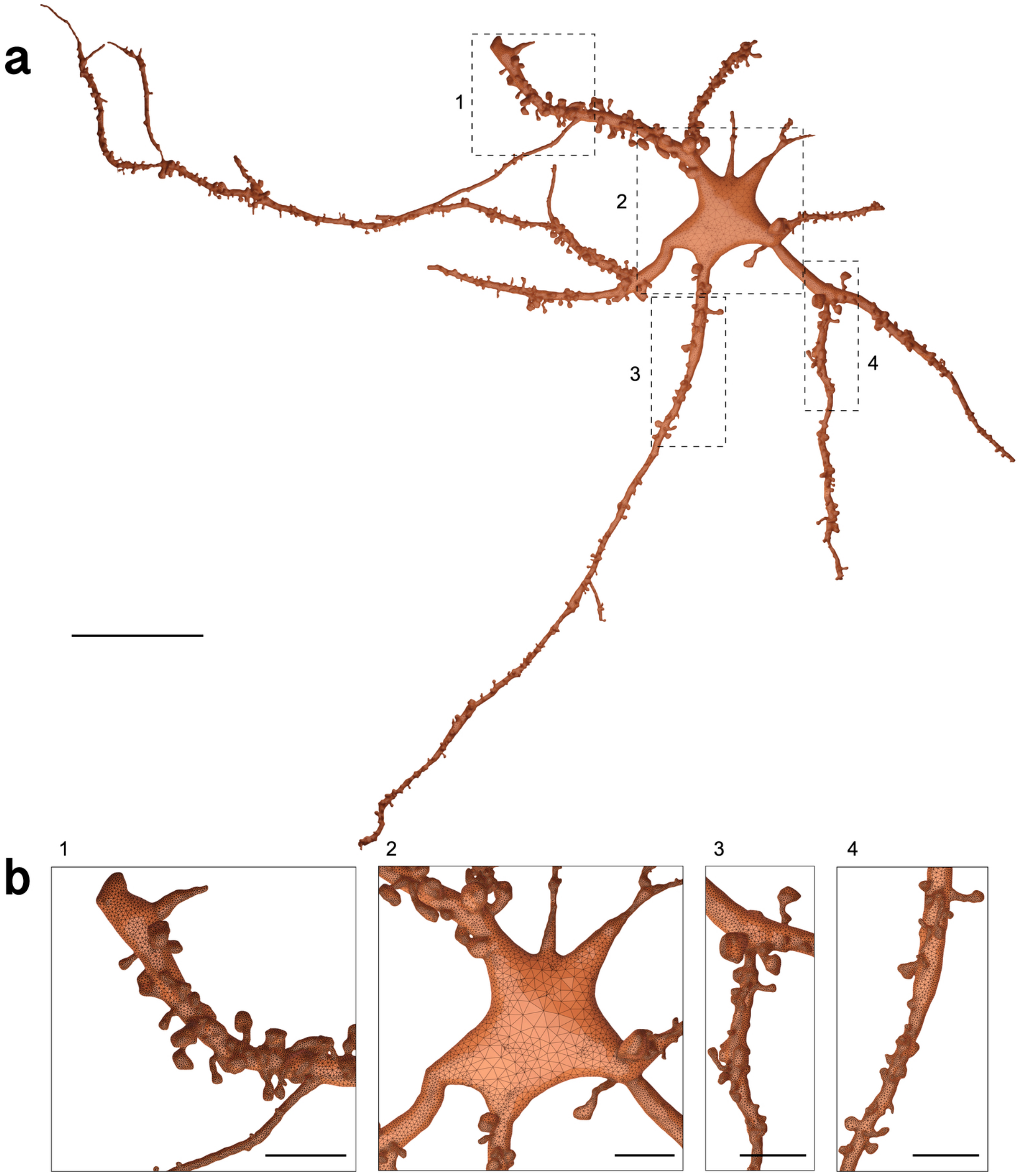

Figure S80: **Reconstruction of a Spiny Neuronal Mesh from its Corresponding Morphology.** Related to Fig. 4. The mesh is reconstructed using ultraNeuroMorpho2Mesh with watertight manifold surface. (**a**) High resolution wireframe rendering of the mesh model shown in Figure 8e. (**b**) The closeups highlight the adaptive topology of the mesh (1-4), the integration of the spines along the dendrites (1, 3 and 4) and the seamless continuity between the soma and dendritic arborizations (2). The morphology is publicly available from NeuroMorpho.Org[25] (Cell ID: 08872, Cell Name: KO-1-DIV-TTb). Spine geometries are manually segmented from the neuropil[22] shown in Figure 3, and spines are randomly distributed using the Meshing Toolbox in NeuroMorphoVis[12]. Scale bars, 20 $\mu$m (**a**) and 5 $\mu$m (**b**).

### 10 Generating continuous cellular meshes from fragmented components

We have used ULTRAMESHES2MESH to build a continuous, optimized and watertight mesh model of a spiny neuron from a set of fragmented meshes. The somatic mesh is created with the Soma generation toolbox of NeuroMorphoVis[12]. The somatic profile is reconstructed on a physically-plausible basis relying on Hooke's law and mass-spring systems to simulate the growth of the soma from an initial icosphere. The arbors are created using the piecewise-watertight method of the meshing toolbox, where each branch in the morphology corresponds an independent mesh partition. The spine meshes are segmented as described in Section 8. Their distributions are computed based on the locations of the neurons in a digitally reconstructed circuit[26]. The meshes corresponding to all the components of the neuron are loaded from a single directory, voxelized in the same grid, where a continuous membrane surface is directly reconstructed. This surface is then optimized and gets verified to be watertight. The process is illustrated in Figure S81.

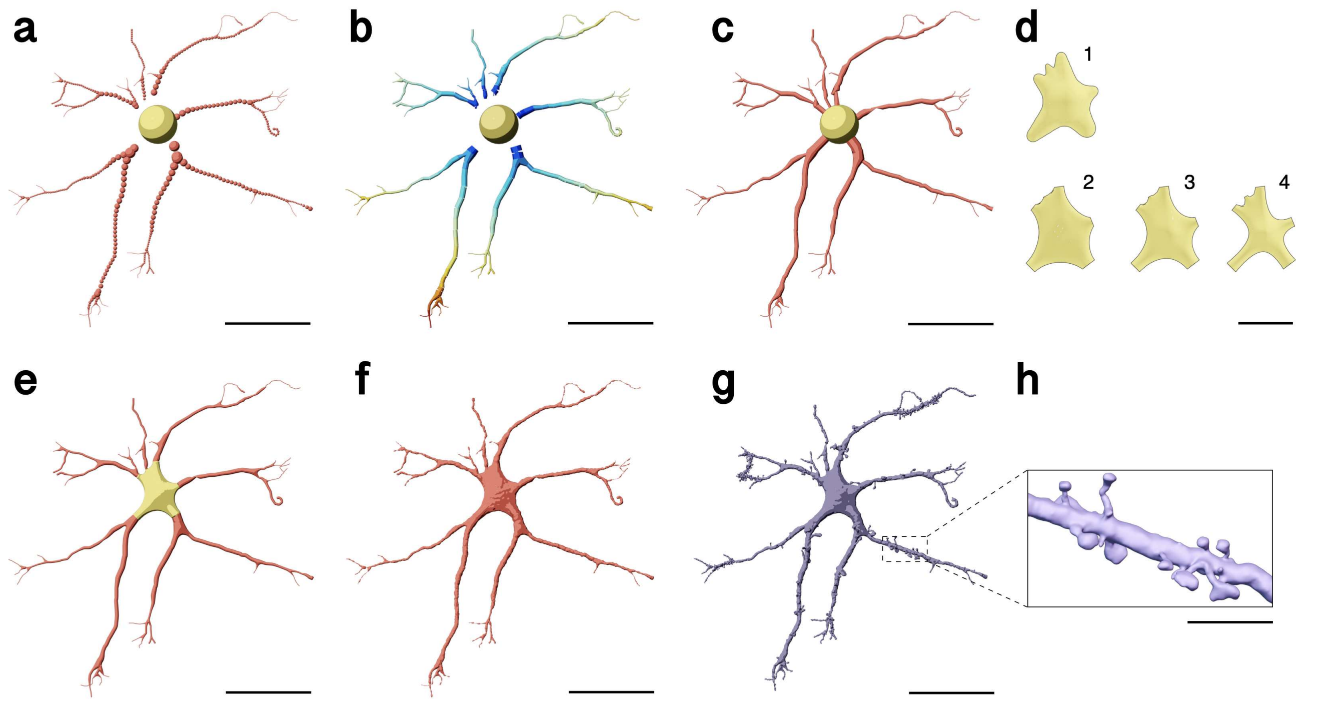

Figure S81: **Reconstruction of an Ultrarealistic Synthetic Watertight Mesh Model of a Spiny Neuron.**
(a) The structure of the morphology is shown as (a) a set of digitized samples, (b) set of segments, where each segment has a different color, (c) set of independent sections. (d) The soma in the top row (1) is reconstructed using implicit surfaces[13]. The somata in the bottom row (2, 3 and 4) are reconstructed on a physically plausible basis using Hooke's law and mass spring models[12] using different simulation parameters. (e) The skin modifiers are used to reconstruct high fidelity neuritic arborizations with accurate branching geometries. (f) The soma and arbors reconstructed in (d) and (e) are integrated in a single mesh. (g) An integrated mesh of the spiny neuron with a continuous and watertight surface is created using ULTRAMESHES2MESH. (h) A closeup showing the dendritic spines. Scale bars, 50$\mu$m (**a, b, c, e, f, g**), 20$\mu$m (**d**), 10 $\mu$m (**h**).

### 11 Generating astroglial meshes from complete synthetic morphologies

We have used ultraAstroMorpho2Mesh to create highly realistic watertight mesh models of astroglial cells from their corresponding synthetic morphologies. This application extends ultraNeuroMorpho2Mesh and loads the endfeet data to build –per endfoot– a proxy mesh that is gets voxelized in the volume grid before the application of the marching cubes kernel. Figure S82 illustrates the arborizations of the astrocytic morphology and the reconstructed mesh after inclusion of the endfeet data along the surface of the resulting mesh.

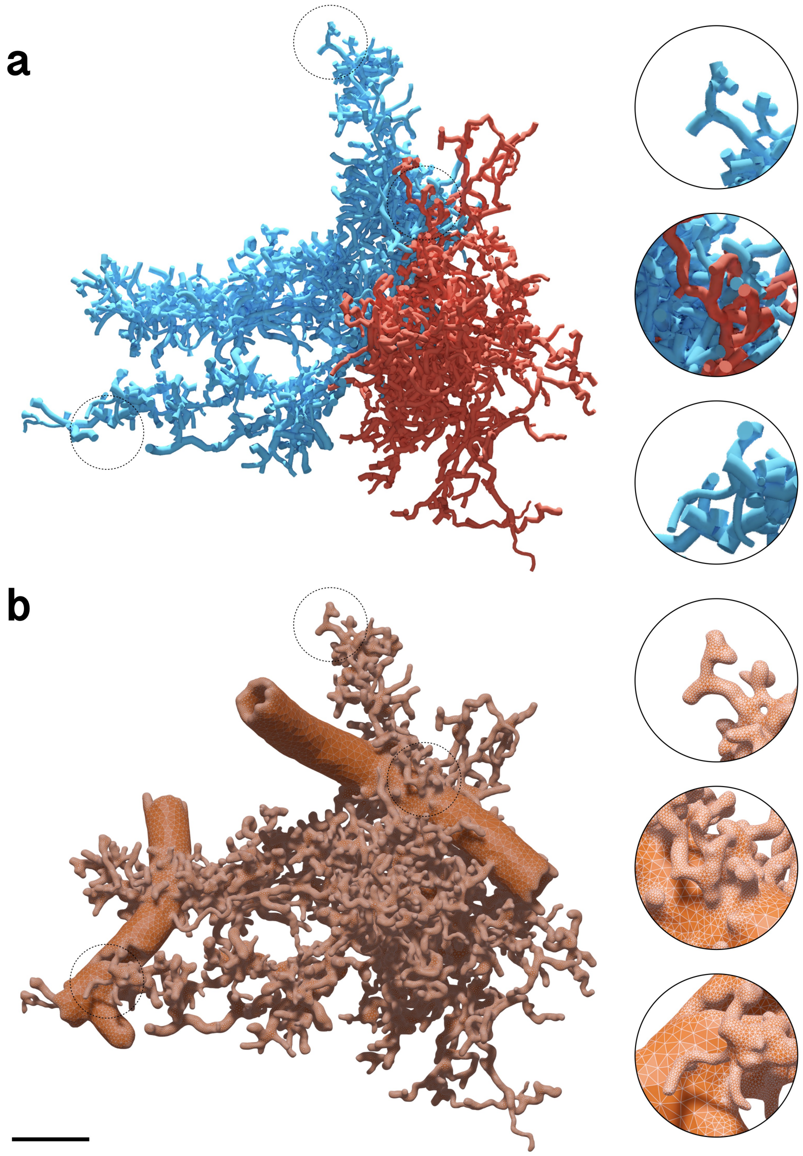

Figure S82: **Astrocyte Mesh Generation from Morphology.** Related to Fig. 5.
Generation of a watertight mesh (**b**) of a synthetic astroglial morphology (**a**). The red and blue branches of the morphology correspond to perisynaptic and perivascular processes respectively. The closeups highlight the clean topology and adaptive tessellation of the resulting mesh. Scale bar, 25 $\mu$m.

#### 12 Generating vasculature meshes from corresponding graph networks

We have used ultraVessMorpho2Mesh to create multi-partitioned watertight meshes of cerebral vasculature from their corresponding morphological networks. In certain cases, the input morphology might have some artifacts resulting from the segmentation process, for example: a zero-radius sample. Such artifacts can potentially affect the quality of the resulting mesh or even lead to the failure of the mesh generation process. We therefore apply a list of pre-processing and morpholometric analysis kernels, with which we can completely validate the structural aspects of the morphology. The analysis results for the vascular network shown in Figure 6 is summarized in Table S9. The sequence of the mesh generation process starting from a large-scale network until the creation of a watertight mesh is shown in Figure S83. The intermediate voxelization step is illustrated in Figure S84.

Table S9: Quantitative morphometric analysis results of the exemplar vascular network shown in Figure 6a.

| **Quantity** | **Value** | **Unit** |
|---|---:|:---:|
| Number of samples | 233,469 | - |
| Minimum sample radius | 0.34997 | $\mu$m |
| Maximum sample radius | 25.46540 | $\mu$m |
| Average sample radius | 1.47667 | $\mu$m |
| Number of zero-radius samples | 0 | - |
| Number of segments | 233,468 | - |
| Minimum segment length | 0.00105 | $\mu$m |
| Maximum segment length | 1.99360 | $\mu$m |
| Average segment length | 1.01871 | $\mu$m |
| Number of sections | 10,009 | - |
| Number of short sections | 1597 | - |
| Number of sections with two samples | 542 | - |
| Minimum section length | 0.009242 | $\mu$m |
| Maximum section length | 327.318 | $\mu$m |
| Average section length | 22.531 | $\mu$m |
| Number of loops | 400 | - |
| Number of components (or partitions) | 16 | - |
| Morphology total length | 227,548.875 | $\mu$m |

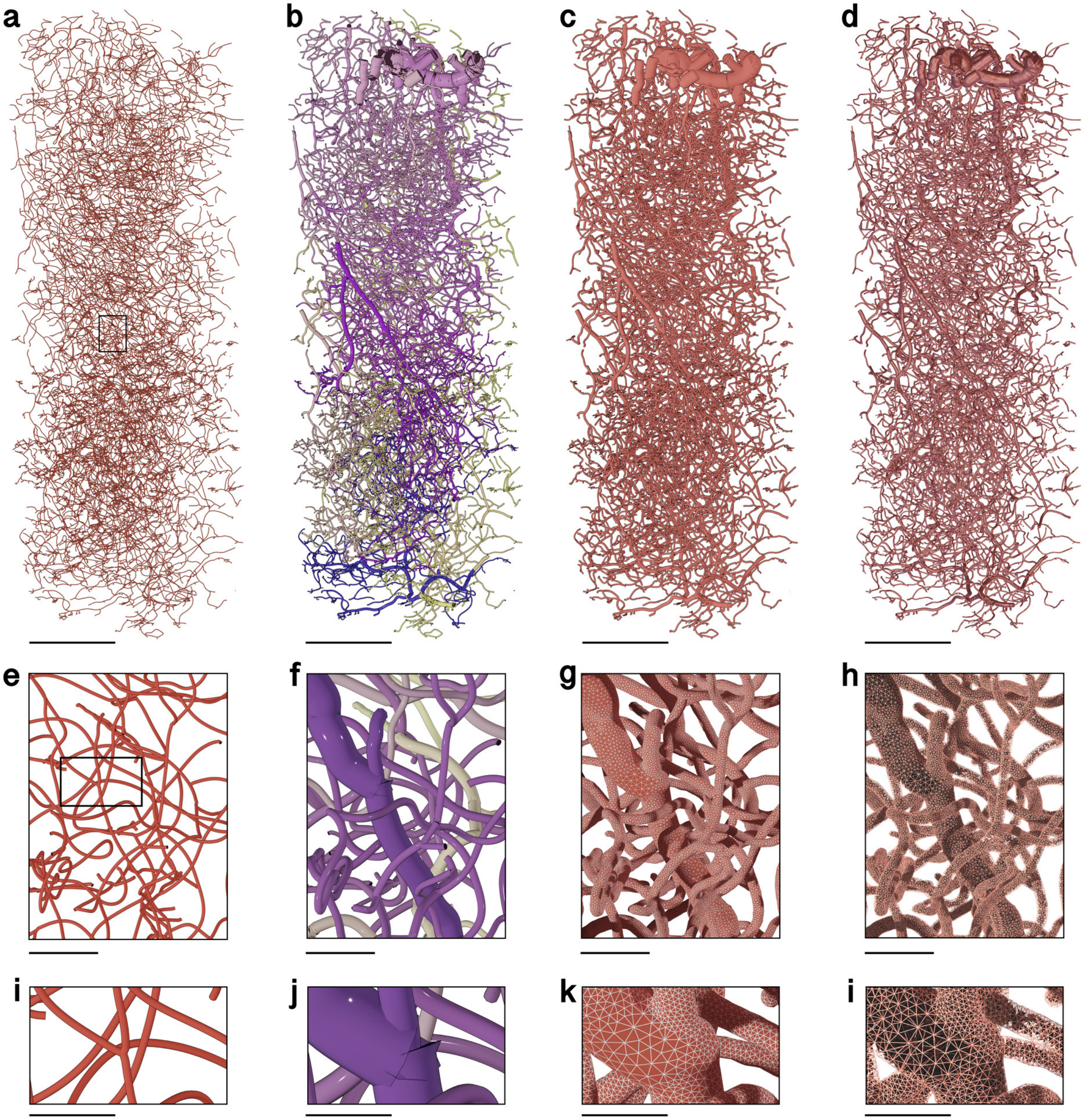

Figure S83: **Vasculature Mesh Reconstruction.** Related to Fig. 6.
Steps showing the conversion of a large-scale, highly detailed vascular graph into a corresponding tetrahedral volumetric mesh tailored to perform blood transport simulations. (**a**) Thin-line representation of the input vascular graph showing its connectivity. (**b**) Polyline representation of the graph showing the actual radii at each vertex in the graph. (**c**) The polyline representation is converted with ultraVessMorpho2Mesh into a watertight manifold having multiple mesh partitions. (**d**) Reconstruction of a tetrahedral volumetric mesh from the watertight mesh reconstructed with Ultraliser. TetGen is used to reconstruct this tetrahedral mesh. The closeups in (**e** - **l**) focus on a branching point. (**k**) Wireframe rendering of the surface mesh showing the accurate and adaptive topology of the manifold. (**l**) Wireframe rendering of the tetrahedral mesh visualizing the internal structure of the volume.
Scale bars, 200 $\mu$m (**a-d**), 25 $\mu$m (**e-h**), 10 $\mu$m (**i-l**).

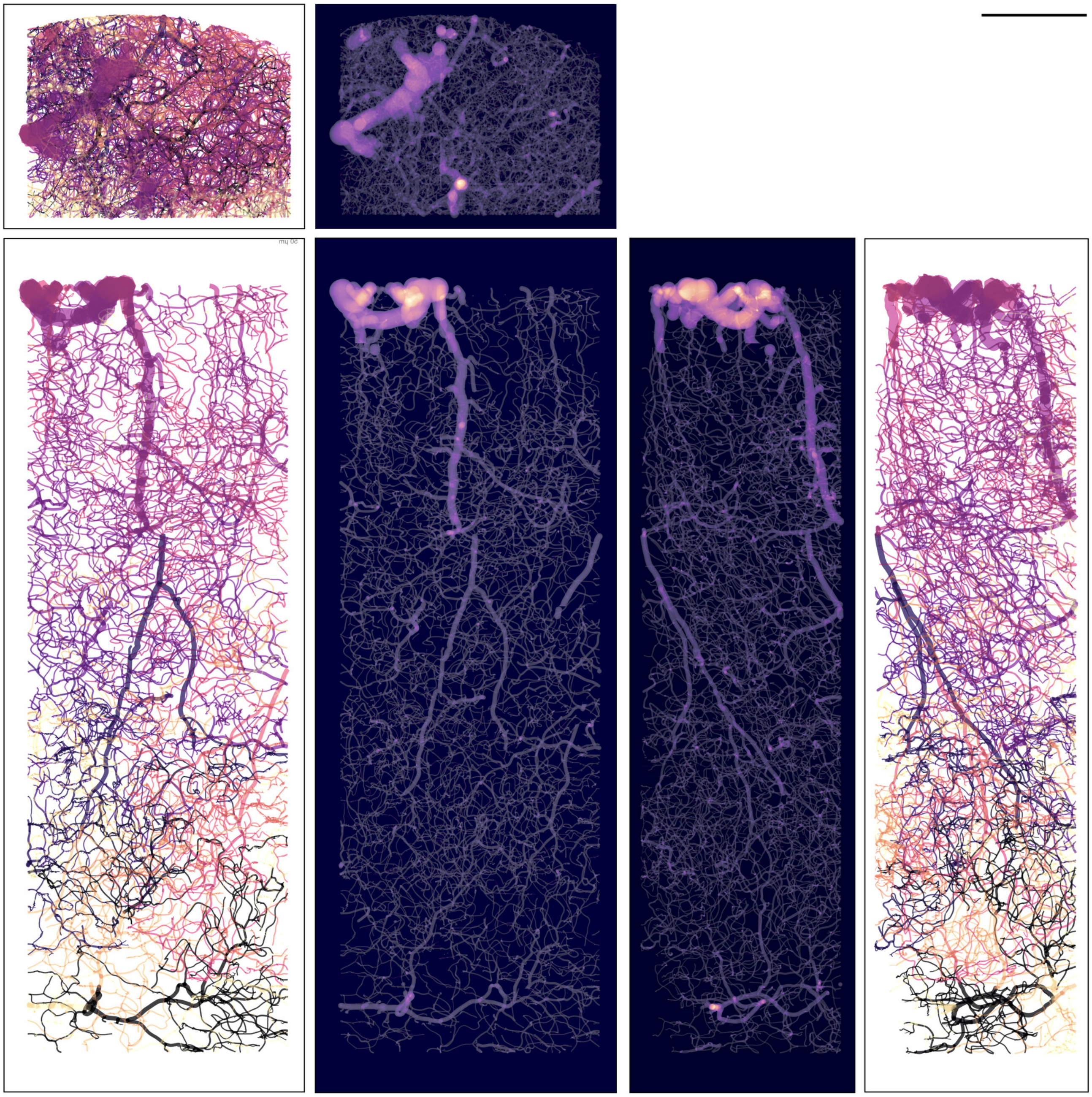

Figure S84: **Vasculature Volume Reconstruction.** Related to Fig. 6.
A side by side comparison between the three principal projections of a large-scale vascular network (**Fig. S83**) in its morphological format (with white background) and its volumetric format (with black background). The morphology skeleton is color-coded based on the vessel radius (smallest radius in black, and largest radius in violet). Scale bar, 200 $\mu$m.

### 13 Comparative analysis with existing frameworks and applications

To demonstrate the critical significance of Ultraliser, we performed detailed quantitative and qualitative comparisons with similar state-of-the-art open source applications that are used for remeshing and mesh reconstruction from morphologial skeletons of neurons (Table S1) and vascular networks (Table S2).

#### 13.1 Remeshing non-watertight mesh models

We *remeshed* a fragmented non-watertight mesh (shown in **Fig. S85b,f**) of a cortical neuron segmented from layer II/III of the visual cortex (refer to **Fig. S54**) with the following applications: VolRoverN[3], GAMer[2,27] and ManifoldPlus[4]. VolRoverN was capable of loading and visualizing the mesh, but it was unable to process it. We then used the Blender[28] add-on of GAMer to facilitate loading and visualizing the mesh. After the application of the surface smoothing function, a significant amount of triangles were removed from the mesh leading to a further fragmented mesh with missing structures (as shown in **Fig. S85c,g**). ManifoldPlus was capable of creating a watertight mesh, but it has destroyed the mesh topology by adding new facets that connect the different spines and small structures together (**Fig. S85d,h**). In contrast, ultraMesh2Mesh was capable of creating an optimized watertight manifold with clean topology (as shown in **Fig. S85e,i**) with a voxelization resolution of four voxels per micron, i.e. 0.25 microns of spatial resolution.

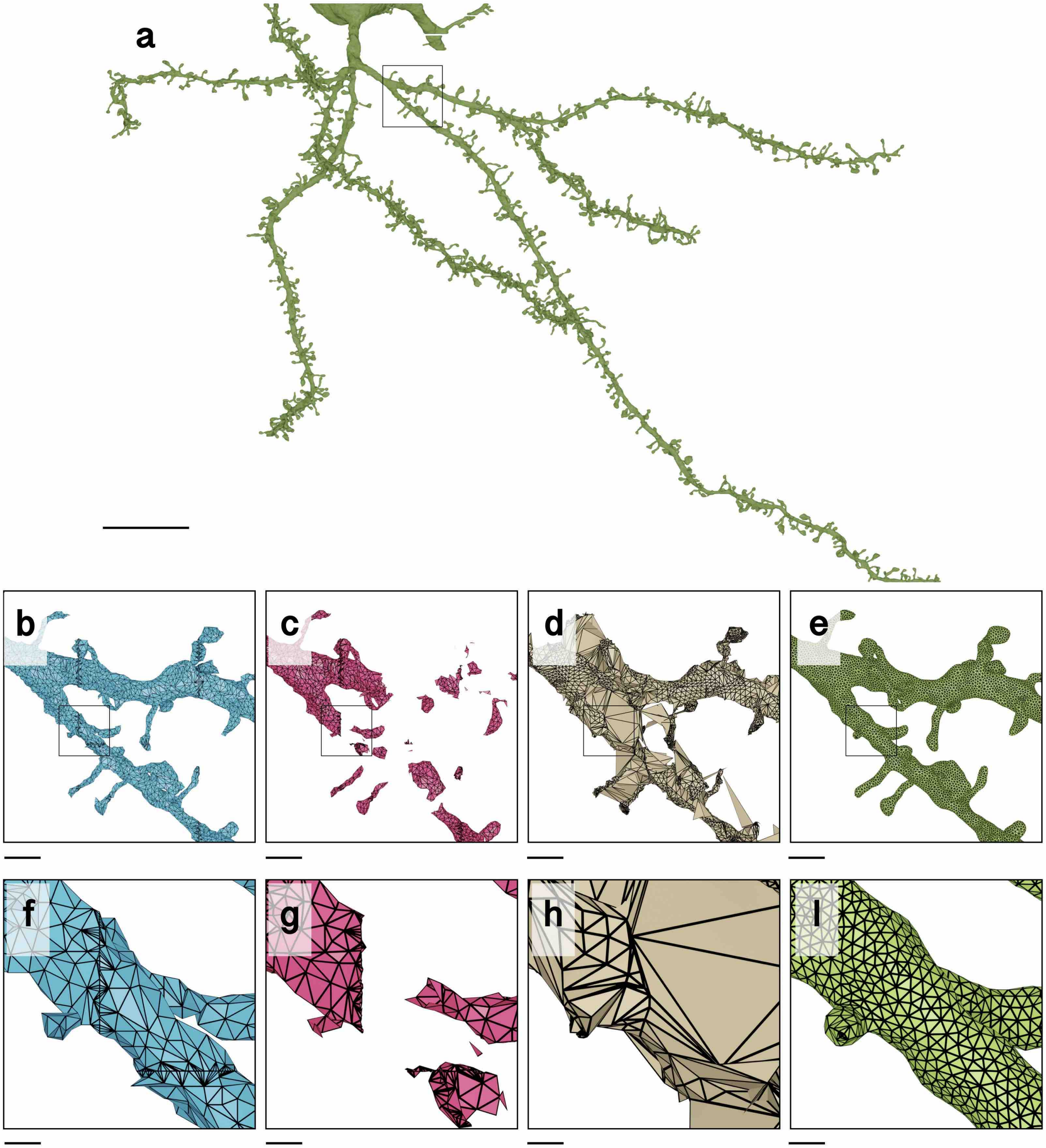

Figure S85: **Remeshing a Non-watertight Fragmented Mesh of a Cortical Neuron**.
Wireframe visualizations comparing the input mesh of a cortical neuron (**a**, **b**, **f**) and the generated meshes with GAMer (**c**, **g**), ManifoldPlus (**d**, **h**) and ultraMesh2Mesh (**e**, **i**) respectively. The original mesh is downloaded from the MICrONS program (microns-explorer.org). Scale bars, 20 $\mu$m (**a**), 5 $\mu$m (**b**, **c**, **d**, **e**) 5 $\mu$m, $\mu$m (**f**, **g**, **h**, **i**) 5 $\mu$m.

##### 13.2 Meshing of neuronal and astroglial morphology skeletons

We used an exemplar neuronal morphology downloaded from NeuroMorpho.Org[25] to compare the performance of ultraNeuroMorpho2Mesh with respect to other open source neuron mesh generation applications (summarized in Table S1) including: AnaMorph[7], NeuroTessMesh[11], Neuronize[9,10] and multiple meshing implementations in NeuroMorphoVis[12]. A visual comparison between the resulting meshes is shown in Figure S86. Comparative qualitative and quantitative analysis of the topology of the meshes is shown in Figure S87.

AnaMorph was incapable of generating a mesh due to a disjoint issue[1], therefore it was excluded from the comparison. None of the other method was capable of creating a watertight mesh except the Metaballs method[13] in NeuroMorphoVis[12]. Nevertheless, this mesh was limited in two aspects: (i) it has poor topology (**Fig. S87d**) and (ii) it also has massive tessellation (∼1.5 million triangles, refer to **Fig. S86d**). Meanwhile, the mesh created with ultraNeuroMorpho2Mesh has superior topology and only ∼172 thousands triangles.

In addition to their incapability to create a manifold watertight surface, the other methods have suffered from further artifacts that can even limit their usability for visual analysis purposes. For example, the meshes created with Neuronize[9] and NeuroTessMesh[11] have major artifacts around the branching points (**Fig. S86e** and **Fig. S86f**) despite the realistic 3D profiles of their somata that are created on a physically-plausible basis using Hooke's law and the finite-element method respectively. The Union-operator-based method[14] in NeuroMorphoVis[12] suffers from non-organic branching shapes and inhomogeneous tessellation, i.e. dense triangulation around the joint locations and less triangulation elsewhere (**Fig. S86b**). Realistic branching is accomplished with the Skinning modifier method[29], but the resulting mesh is highly tessellated (**Fig. S86c**). The resulting mesh with the Piecewise-watertight method[30] contains 163 partitions, which makes it also limited for visualizing electrophysiological simulations, where transparency is needed to visualize the variations from depolarization to hyperpolarization (**Fig. S86a**).

In contrast, the mesh created with ultraNeuroMorpho2Mesh is watertight, optimized ($\sim 172 \times 10^3$ triangles), has optimum topology (**Fig. S87g**), has highly realistic somatic profile simulated with the finite-element method and natural-looking branching structures (**Fig. S86g**). These features makes this mesh optimum for usage in multiple applications including: performing reaction-diffusion simulations, visual analytics and visualization of physiological parameters, skeletonization and voxelization with traditional applications as shown earlier in Figure S2.

1 The error message is: `RedBlue_Algorithm()`: no red edge intersect a blue face and vice versa => red and blue meshes disjoint.

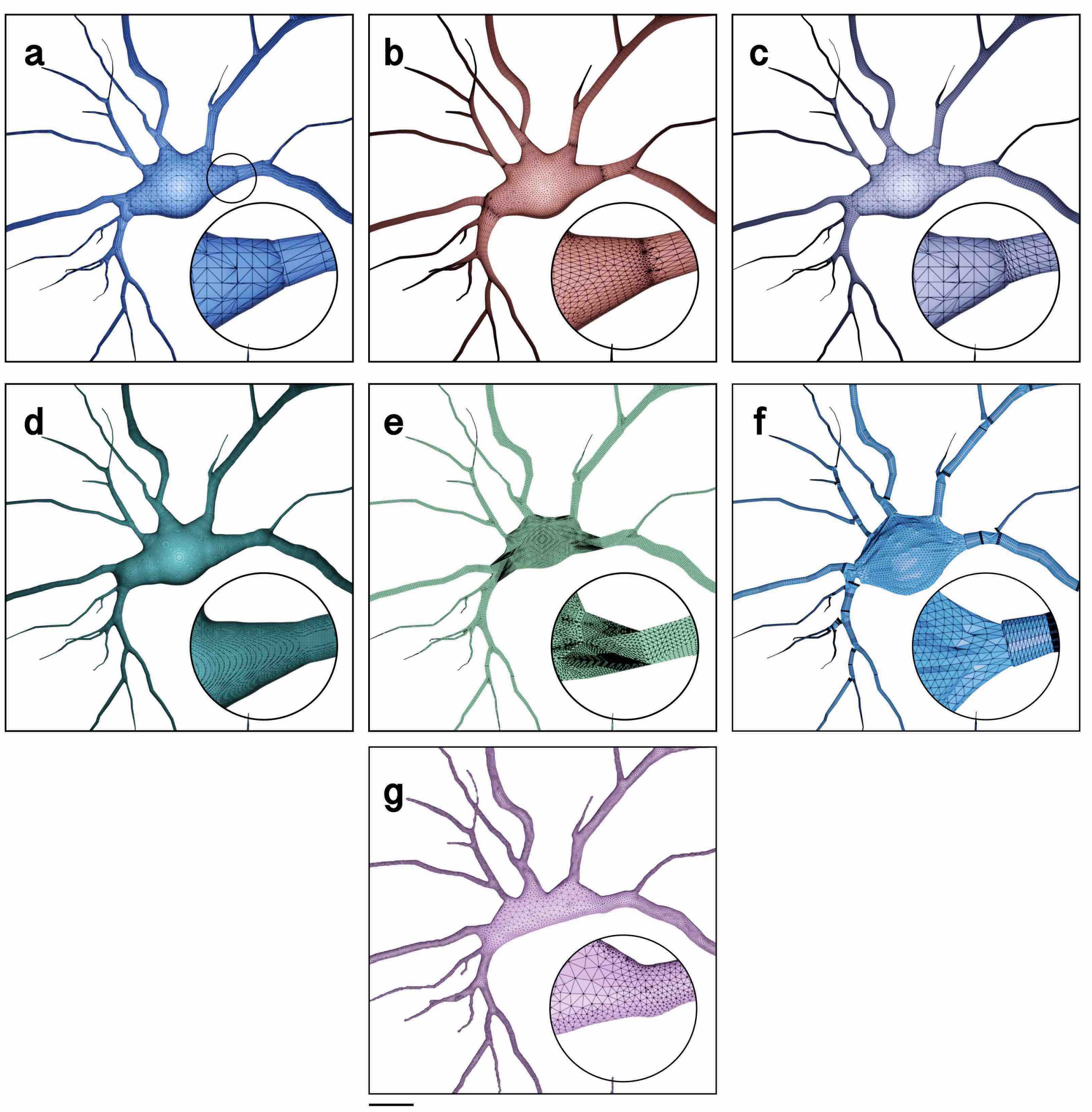

Figure S86: **Reconstruction of Neuronal Meshes from a Morphology Skeleton.**
Visual comparison between the resulting meshes generated by the different meshing methods and applications listed in Table S1 and ultraNeuroMorpho2Mesh. Comparative qualitative and quantitative analysis is shown in Figure S87. The morphology is downloaded from NeuroMorpho.Org[25]. Scale bar, 5 $\mu$m.

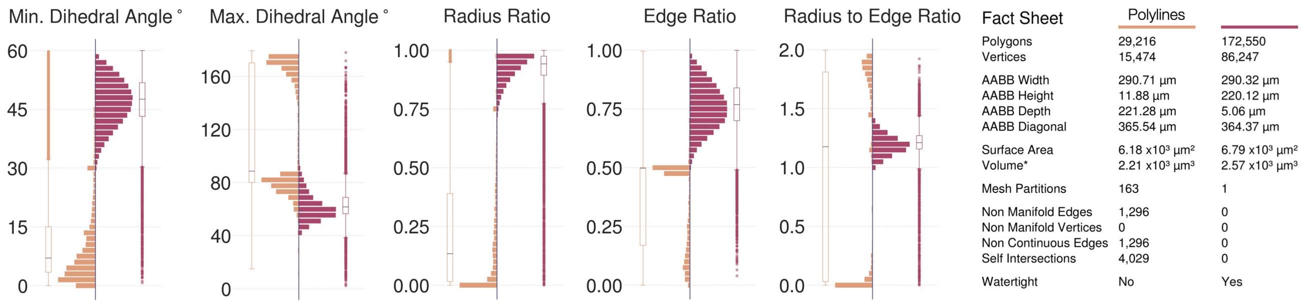

(**a**) Mesh created using the Piecewise Watertight (or Polylines) method[30] in NeuroMorphoVis[12], Visualization in Figure S86a.

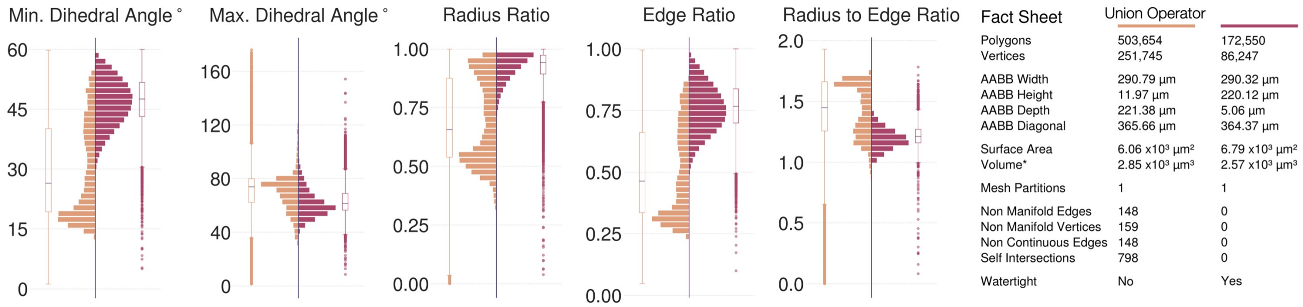

(**b**) Mesh created using the Union Operators method[14] in NeuroMorphoVis[12], Visualization in Figure S86b.

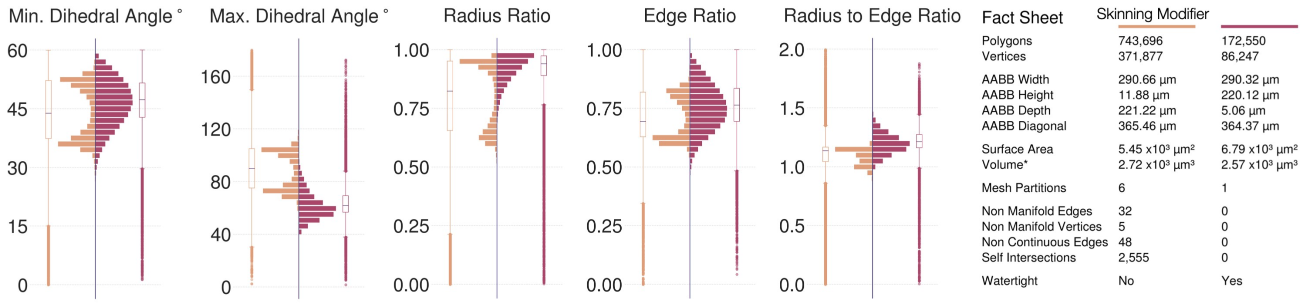

(**c**) Mesh created using the Skinning Modifier method[29] in NeuroMorphoVis[12], Visualization in Figure S86c.

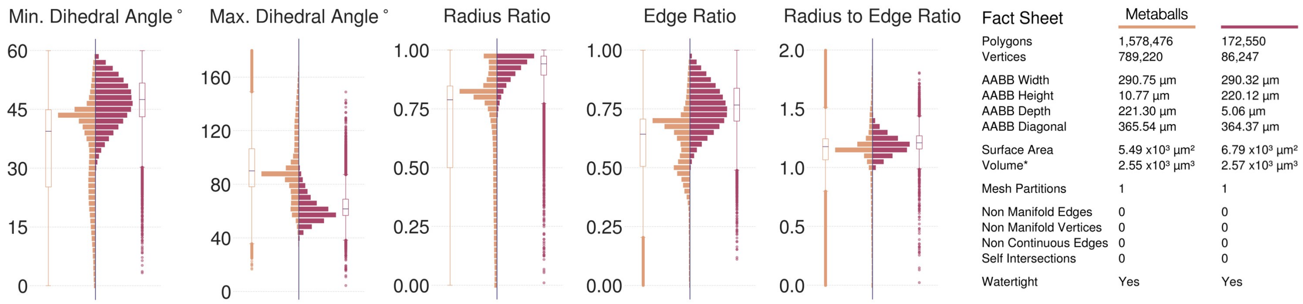

(**d**) Mesh created using the Metaballs method[13] in NeuroMorphoVis[12], Visualization in Figure S86d.

Figure S87: Continued in the next page.

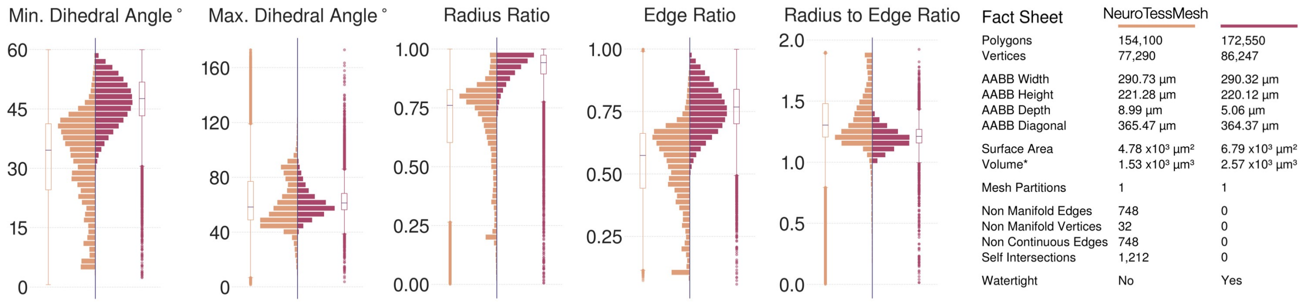

(**e**) Mesh created using NeuroTessMesh[11], Visualization in Figure S86e.

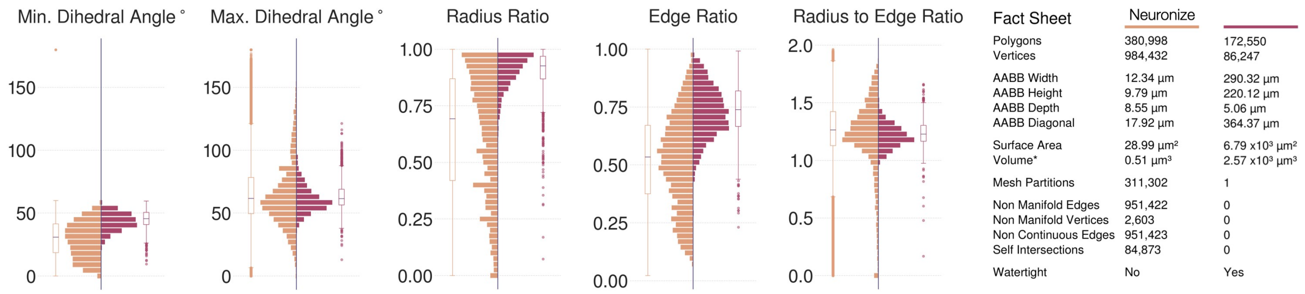

(**f**) Mesh created using Neuronize[9], Visualization in Figure S86f.

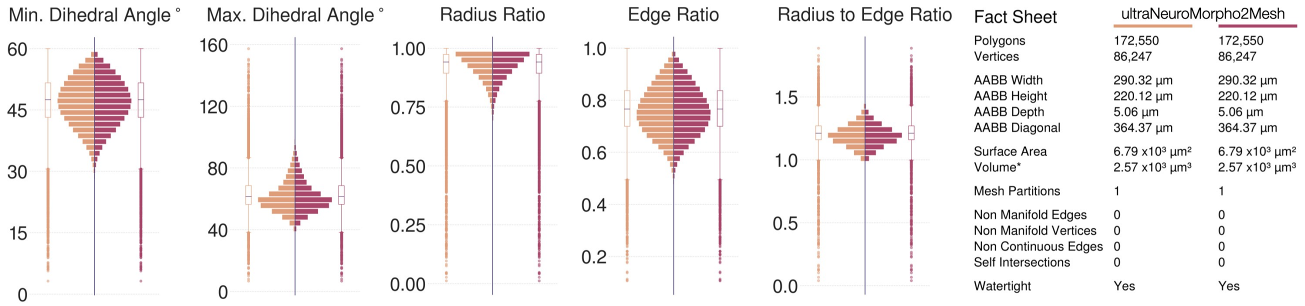

(**g**) Mesh created using ultraNeuroMorpho2Mesh, Visualization in Figure S86g.

Figure S87: Comparative qualitative and quantitative analysis of the neuronal meshes generated by the different meshing methods and applications listed in Table S1 and ultraNeuroMorpho2Mesh. Wireframe models of the resulting meshes are displayed in Figure S86.

##### 13.3 Meshing of vascular morphology skeletons

We also used an exemplar vascular morphology downloaded from the Brain Vasculature (BraVa)[31] database (cng.gmu.edu/brava) to compare the performance of ultraVessMorpho2Mesh with respect to other available vascular mesh generation applications (Table S2). A visual comparison between the resulting meshes for the exemplar morphology is shown in Figure S88. Comparative qualitative and quantitative analysis of the topology of the meshes is shown in Figure S89. Similarly, the Metaballs method[13] in VessMorphoVis was also capable of creating a watertight mesh, but with poor topology and high tessellation ($\sim 1.2 \times 10^6$ triangles) compared to the mesh created with ultraVessMorho2Mesh ($\sim 8.7 \times 10^3$ triangles).

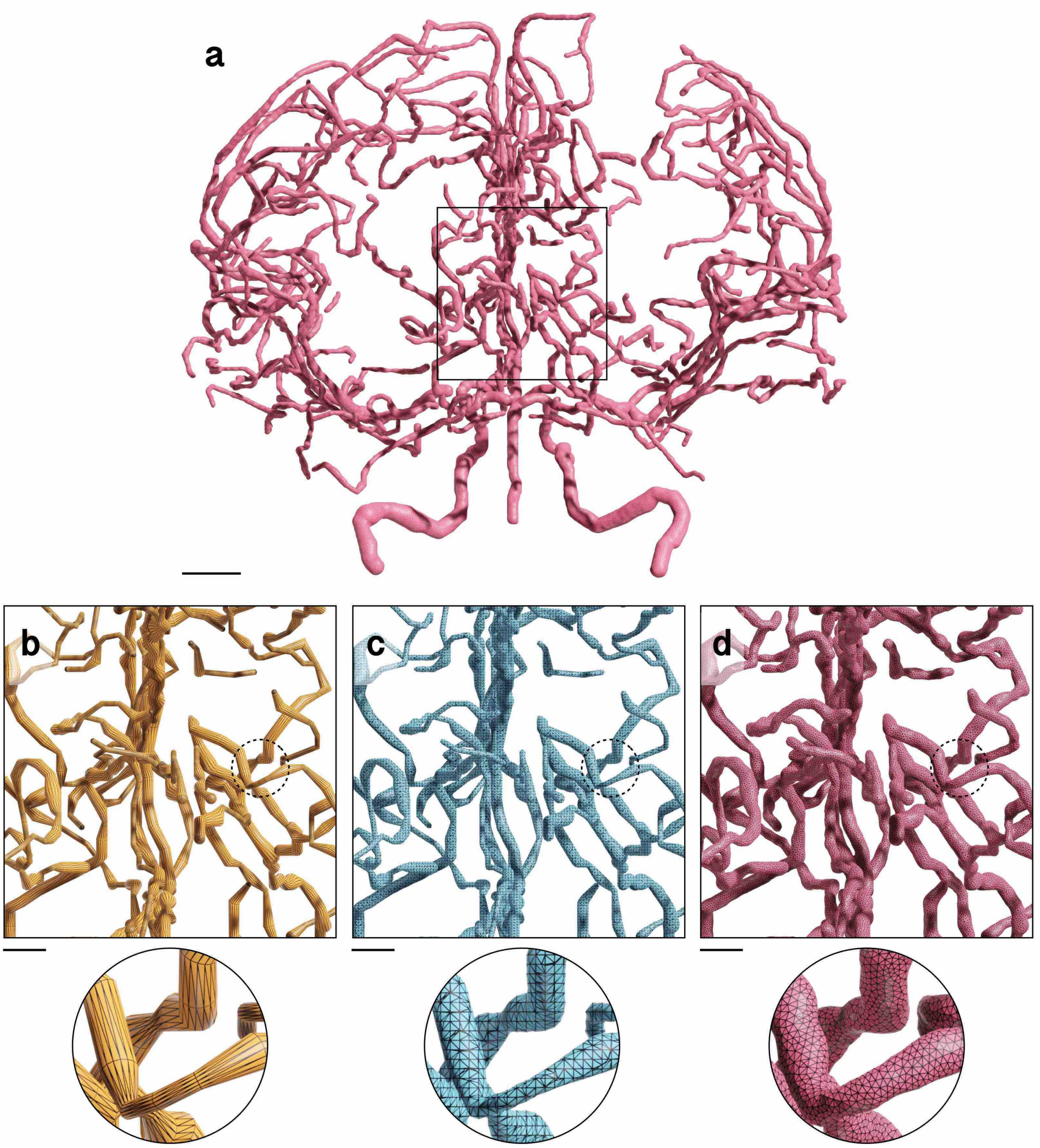

Figure S88: **Mesh Reconstruction of Human Arterial Vasculature from its Morphology Network.**
Visual comparison between the resulting vascular meshes created from an exemplar network of human arterial arborizations (**a**) using (**b**) the piecewise-watertight, (**c**) the metaballs methods in VessMorphoVis and (**d**) the ultraVessMorpho2Mesh application that is integrated into Ultraliser. The closeups highlight the topology of the resulting meshes. Comparative qualitative and quantitative analysis is shown in Figure S89. The vascular morphology network is available from the Brain Vasculature (BraVa)[31] database (cng.gmu.edu/brava). Scale bars, 20 $\mu$m (**a**), 5 $\mu$m (**b**,**c**,**d**).

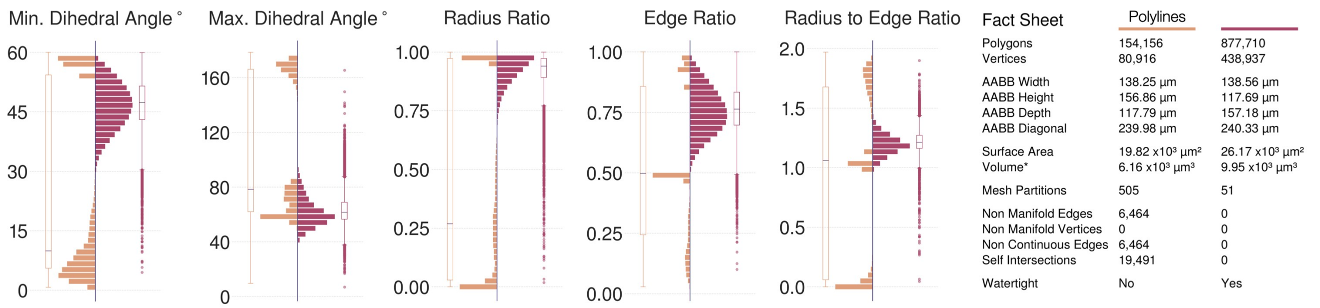

(**a**) Mesh created using the Piecewise Watertight (or Polylines) method in VessMorphoVis, Visualization in Figure S88b.

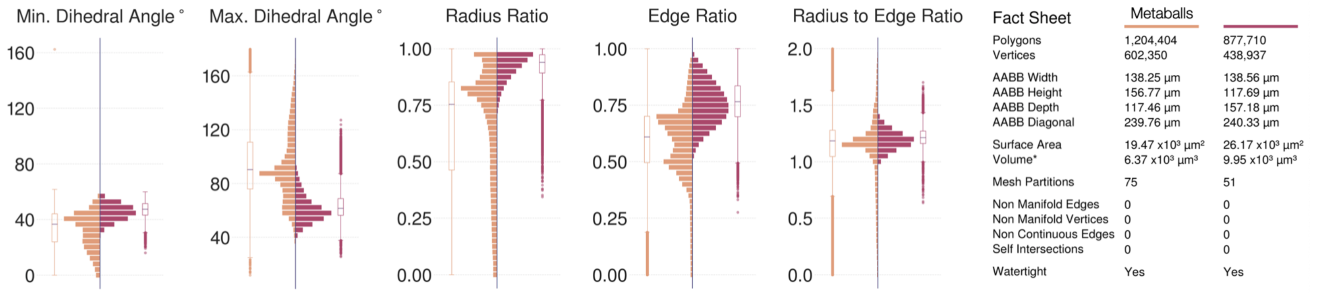

(**b**) Mesh created using the Metaballs method in VessMorphoVis, Visualization in Figure S88c.

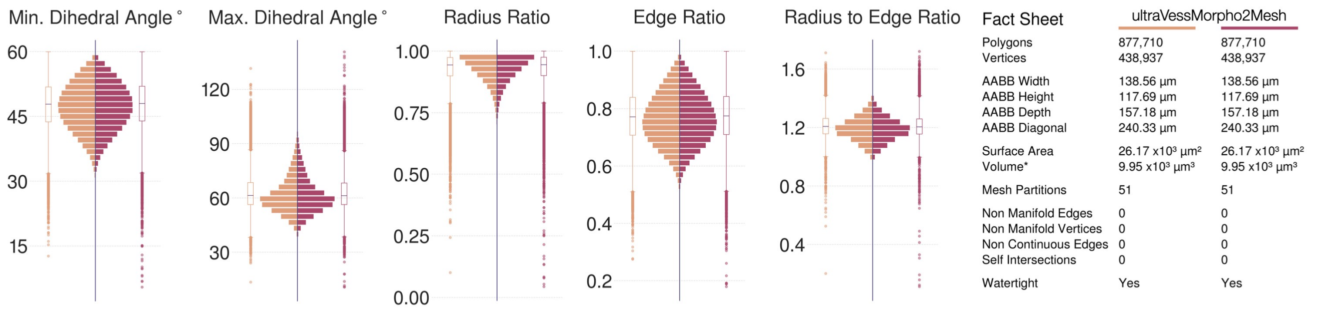

(**c**) Mesh created using ULTRAVESSMORPHO2MESH, Visualization in Figure S88d.

Figure S89: Comparative qualitative and quantitative analysis of the three vascular meshes created by (**a**) the piecewise-watertight, the metaballs methods of VessMorphoVis and (**c**) ULTRAVESSMORPHO2MESH. The wireframe models of the three meshes are shown in Figure S88.

#### References

1. Abdellah, M. In Silico Brain Imaging: Physically-plausible Methods for Visualizing Neocortical Microcircuitry, 400. http://infoscience.epfl.ch/record/232444 (2017).
2. Lee, C. T. *et al.* 3D mesh processing using GAMer 2 to enable reaction-diffusion simulations in realistic cellular geometries. *PLoS computational biology* **16,** e1007756 (2020).
3. Edwards, J. *et al.* VolRoverN: enhancing surface and volumetric reconstruction for realistic dynamical simulation of cellular and subcellular function. *Neuroinformatics* **12,** 277–289 (2014).
4. Huang, J., Zhou, Y. & Guibas, L. Manifoldplus: A robust and scalable watertight manifold surface generation method for triangle soups. *arXiv preprint arXiv:2005.11621* (2020).
5. Eilemann, S. *et al. From big data to big displays high-performance visualization at blue brain* in *International Conference on High Performance Computing* (2017), 662–675.
6. Eilemann, S. *et al. Parallel rendering on hybrid multi-gpu clusters* in *Eurographics Symposium on Parallel Graphics and Visualization* (2012), 109–117.
7. Mörschel, K., Breit, M. & Queisser, G. Generating neuron geometries for detailed three-dimensional simulations using anamorph. *Neuroinformatics* **15,** 247–269 (2017).
8. McDougal, R. A., Hines, M. L. & Lytton, W. W. Water-tight membranes from neuronal morphology files. *Journal of neuroscience methods* **220,** 167–178 (2013).
9. Brito, J. P. *et al.* Neuronize: a tool for building realistic neuronal cell morphologies. *Frontiers in neuroanatomy* **7** (2013).
10. Velasco, I. *et al.* Neuronize v2: bridging the gap between existing proprietary tools to optimize neuroscientific workflows. *Frontiers in neuroanatomy* **14,** 585793 (2020).
11. Garcia-Cantero, J. J., Brito, J. P., Mata, S., Bayona, S. & Pastor, L. NeurotessMesh: A tool for the Generation and Visualization of Neuron Meshes and Adaptive on-the-Fly Refinement. *Frontiers in neuroinformatics* **11,** 38 (2017).
12. Abdellah, M. *et al.* NeuroMorphoVis: a collaborative framework for analysis and visualization of neuronal morphology skeletons reconstructed from microscopy stacks. *Bioinformatics* **34,** i574–i582 (2018).
13. Abdellah, M. *et al.* Metaball skinning of synthetic astroglial morphologies into realistic mesh models for visual analytics and in silico simulations. *Bioinformatics* **37,** i426–i433 (2021).
14. Abdellah, M. *et al. Meshing of Spiny Neuronal Morphologies using Union Operators* in *Computer Graphics and Visual Computing (CGVC)* (eds Vangorp, P. & Turner, M. J.) (The Eurographics Association, 2022). ISBN: 978-3-03868-188-5.
15. Abdellah, M. *et al.* Interactive visualization and analysis of morphological skeletons of brain vasculature networks with VessMorphoVis. *Bioinformatics* **36,** i534–i541 (2020).

16. Botsch, M., Kobbelt, L., Pauly, M., Alliez, P. & Lévy, B. *Polygon mesh processing* (CRC press, 2010).

17. Aspert, N., Santa-Cruz, D. & Ebrahimi, T. *Mesh: Measuring errors between surfaces using the hausdorff distance* in *Proceedings. IEEE international conference on multimedia and expo* **1** (2002), 705–708.

18. Coggan, J. S. *et al.* A Process for Digitizing and Simulating Biologically Realistic Oligocellular Networks Demonstrated for the Neuro-Glio-Vascular Ensemble. *Frontiers in neuroscience* **12** (2018).

19. Cardona, A. *TrakEM2: an ImageJ-based program for morphological data mining and 3d modeling* in *Proc. ImageJ User and Developer Conference* **4** (2006).

20. Cardona, A. *et al.* TrakEM2 software for neural circuit reconstruction. *PloS one* **7,** e38011 (2012).

21. Holst, G., Berg, S., Kare, K., Magistretti, P. & Cali, C. *Adding large EM stack support* in *2016 4th Saudi International Conference on Information Technology (Big Data Analysis)(KACSTIT)* (2016), 1–7.

22. Cali, C. *et al.* 3D cellular reconstruction of cortical glia and parenchymal morphometric analysis from Serial Block-Face Electron Microscopy of juvenile rat. *Progress in neurobiology* **183,** 101696 (2019).

23. Zisis, E. *et al.* Digital Reconstruction of the Neuro-Glia-Vascular Architecture. *Cerebral Cortex* **31,** 5686–5703 (2021).

24. Mettke, H. Convex cubic Hermite-spline interpolation. *Journal of Computational and Applied Mathematics* **9,** 205–211 (1983).

25. Ascoli, G. A., Donohue, D. E. & Halavi, M. NeuroMorpho. Org: a central resource for neuronal morphologies. *Journal of Neuroscience* **27,** 9247–9251 (2007).

26. Markram, H., Muller, E., Ramaswamy, S., Reimann, M. W., Abdellah, M., *et al.* Reconstruction and simulation of neocortical microcircuitry. *Cell* **163,** 456–492 (2015).

27. Lee, C. T. *et al.* An open-source mesh generation platform for biophysical modeling using realistic cellular geometries. *Biophysical journal* **118,** 1003–1008 (2020).

28. Foundation, B. *Blender - 3D modelling and rendering package* Blender Foundation (Blender Institute, Amsterdam, 2018). www.blender.org.

29. Abdellah, M., Favreau, C., Hernando, J., Lapere, S. & Schürmann, F. *Generating High Fidelity Surface Meshes of Neocortical Neurons using Skin Modifiers* in *Computer Graphics and Visual Computing (CGVC)* (eds Vidal, F. P., Tam, G. K. L. & Roberts, J. C.) (The Eurographics Association, 2019). ISBN: 978-3-03868-096-3.

30. Abdellah, M. *et al.* Reconstruction and visualization of large-scale volumetric models of neocortical circuits for physically-plausible in silico optical studies. *BMC bioinformatics* **18,** 402 (2017).

31. Wright, S. N. *et al.* Digital reconstruction and morphometric analysis of human brain arterial vasculature from magnetic resonance angiography. *Neuroimage* **82,** 170–181 (2013).
